# A Unified 3D Generative Model for Synthesizable Structure-Based Drug Design

**DOI:** 10.64898/2026.09.15.751537

**Authors:** Ilia Igashov, Arne Schneuing, Adrian W. Dobbelstein, Irina Morozova, Rebecca M. Neeser, Kara Zielinski, Luciano Andres Abriata, Aaron S. Petruzzella, David R. Pavel Iosub, Olivia Gampp, Artem Y. Lyubimov, Evgenia Elizarova, Isabella Ferrara, Pedro M. F. Sousa, Ana R. Lemos, Fabio Testori, Pierre A. Miranda Herrera, Laurin Kanis, Joseph Schmidt, Mac Kevin E. Braza, Rommie E. Amaro, Nicolas Thomä, Davide M. Ferraris, Roland Riek, James S. Fraser, Philippe Schwaller, Michael Bronstein, Bruno Correia

## Abstract

Traditional screening-based drug discovery is inherently limited by the astronomical scale of the chemical space. Generative modelling offers a compelling alternative to the classical search paradigm and enables rational, bottom-up design of novel and target-specific small molecules. However, its impact has been hampered by challenges in synthetic accessibility of the designed compounds and lack of large-scale experimental validation. Here, we introduce LDDM (Large Drug Discovery Model), a generative framework that supports a range of drug discovery tasks, including constrained and unconstrained docking, fragment linking and growing, and *de novo* design. We further introduce a programmable design algorithm that enables accurate design of synthetically accessible compounds satisfying various fine-grained objectives. We experimentally validated the designed or optimised ligands for five therapeutically relevant protein targets. In all cases, LDDM achieved high success rates, allowing us to identify molecules with confirmed binding affinity while synthesizing only a small number of generated compounds. The best designs were structurally characterised through NMR spectroscopy and X-ray crystallography, demonstrating high prediction accuracy. Overall, LDDM provides a scalable and flexible platform for the rapid and tailored design of small molecules and non-natural peptides for therapeutic applications.

## Introduction

Computational drug discovery has traditionally been dominated by virtual screening and scoring methods that aim to identify potential small molecule ligands through computational docking and ranking of available compound libraries [1, 2]. Virtual screening substantially exceeds the throughput of experimental screening methods [2], and empirical docking-based approaches have been successfully scaled to more than 100 million compounds, yielding novel chemotypes with confirmed biological activity [3]. Despite these advances, the chemical space coverage remains minuscule compared with the number of all possible drug-like compounds which is estimated to be at least a billion times larger [4, 5]. Because computational cost scales linearly with library size, the chemical space accessible by exhaustive screening is fundamentally limited.

Generative deep learning offers an alternative route to exploring chemical space and increasing chemical diversity. Most existing approaches operate with SMILES [6–9] or graph representations [10, 11]. High abundance of 2D molecular data and the low computational cost of training and inference make such models easy to integrate with goal-directed optimisation techniques [8, 12–19] to guide generation toward desired molecular properties and regions of the chemical space. However, because target-specific information is provided indirectly through external docking tools or activity oracles [15, 17, 18], these models capture the geometry and physics of protein–ligand interactions only implicitly.

To overcome this limitation, several recent methods operate directly on 3D molecular data, enabling explicit reasoning about the geometry and physics of molecular conformations and protein-ligand interactions. These include autoregressive 3D models [20–22] and score-based generative frameworks [23–25], both of which are beginning to show strong potential for structure-based drug design. However, several limitations hinder their adoption in early-stage drug discovery programmes. First, binding affinity remains notoriously hard to predict [26], as are many other developability properties that are required to turn a mere ligand into a viable drug candidate [27]. In addition, the ability of generative models to explore vast chemical spaces often comes at the expense of reduced synthetic tractability of the generated molecules [28]. Unlike virtual screening [29, 30], which typically evaluates compounds that are already available or readily synthesizable, *de novo* generated molecules are often difficult or impossible to synthesize, hindering their experimental validation and ultimately slowing down method improvement.

Here, we introduce LDDM (Large Drug Discovery Model), a unified generative framework that jointly models atom types, coordinates and covalent bonds of small molecules, and samples all these data modalities conditioned on the structure of the target protein pocket. By construction, LDDM supports *de novo* ligand generation as well as substructure- and fragment-based design, covalent and non-covalent docking, and peptide design with non-canonical amino acids. Crucially, the fragment-centric approach allows us to introduce a new programmable design algorithm for the iterative generation of new molecules with optimised properties of interest within a given virtual chemical space [32, 33].

Encouraged by competitive *in silico* benchmarking results across various docking and design tasks, we conducted five experimental case studies. First, redesigning the two CDO1-contacting flanks of a known VHL-CDO1 molecular glue yielded new analogs that recruited CDO1 with nanomolar affinity in 67% of cases. Second, modifying a reported peptide inhibitor by docking non-canonical amino acid side chains into the oncogenic cathepsin S protein led to a 3.5-fold improvement in IC_50_. Finally, we present several fully *de novo* designed micromolar ligands for phosphoglycerate kinase 1 (PGK1), BRD4, and SARS-CoV-2 Nsp3, all with experimental evidence supporting the predicted binding poses.

Together, these results position LDDM as a broadly applicable tool for structure-based drug discovery across a wide range of targets and modalities. Moreover, by constraining proposed compounds to on-demand chemical spaces, the framework facilitates rapid and cost-effective experimental validation enabling efficient design-testing cycles to define useful target-specific chemical spaces for ligand development. As a result, our method enhances computational design capabilities and reduces practical barriers to translating computationally generated molecular designs into testable compounds.

## Results

### A unified framework for computational drug discovery

The LDDM architecture was devised to enhance applicability across various drug discovery tasks. To do so, we introduce a fragment-based masked modelling strategy inspired by self-supervised learning techniques in natural language processing [34]. Similar to language models that learn to complete missing tokens in a sentence, we trained LDDM to recover masked portions of input molecules conditioned on a protein pocket, enabling efficient self-supervised learning and modelling the probabilistic structure of the chemical space. To define chemically meaningful units for masking, we decompose each molecule into fragments, as illustrated in Figure 1A. During training, we randomly mask different subsets of fragments and task the model to recover the missing parts. Additionally, we perform masking across data modalities. We either remove the entire fragment (design regime) or mask only its coordinates (docking regime), as shown in Figure 1B.

**Fig. 1:**
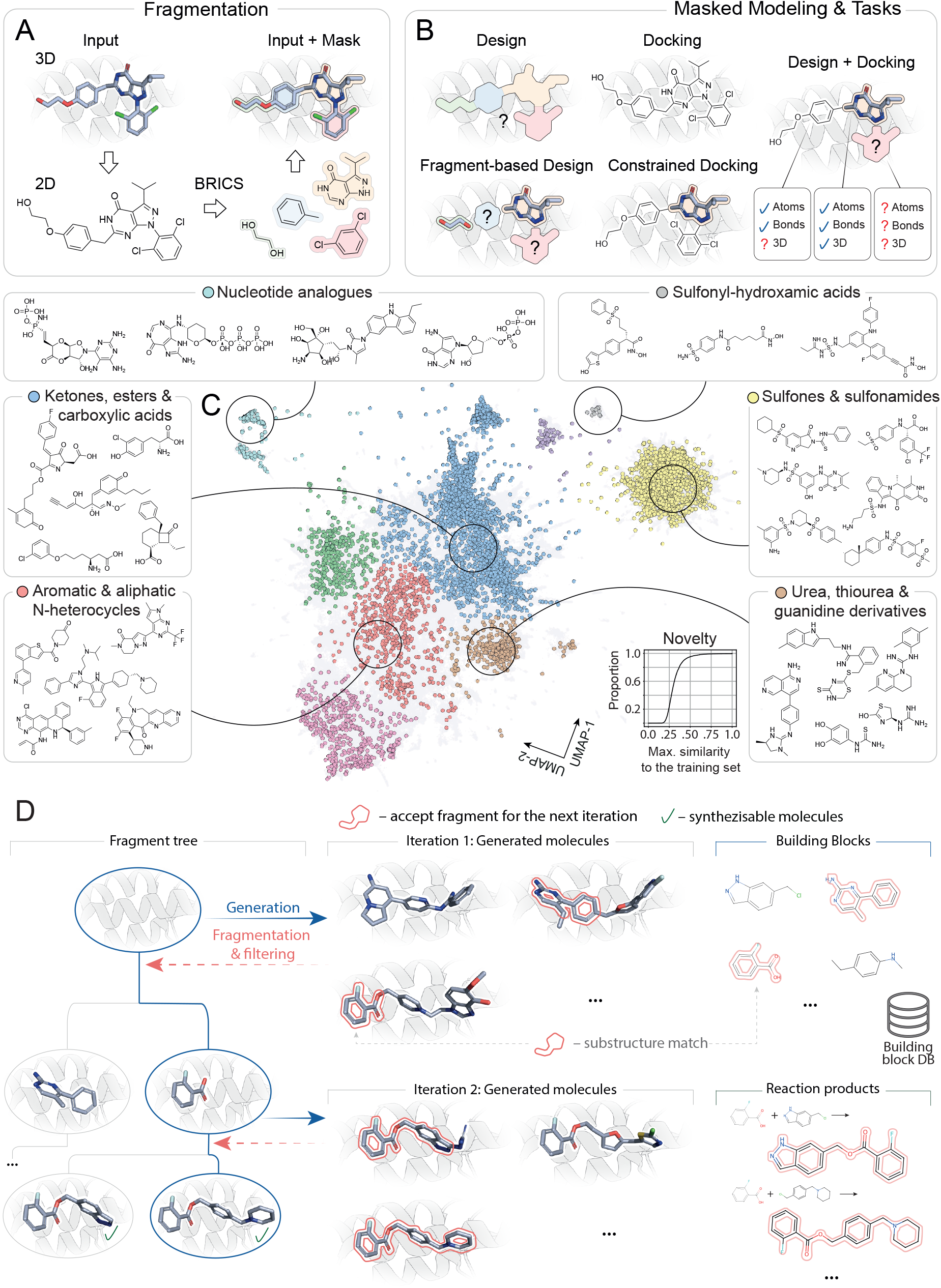
Method overview. **(A)** To prepare the data for training, we identify retrosynthetically-interesting substructures using the BRICS [31] method, and assign fragment labels to different parts of each molecule. **(B)** During training, each fragment can either be masked out completely, partially given as a molecular graph without coordinates or fully provided as additional context for the denoising network. The trained model can then be applied naturally to diverse tasks by specifying the appropriate amount of information given for each fragment. **(C)** Chemical space of LDDM samples (*n* = 10 000). Each generated molecule is represented as a point corresponding to the two-dimensional UMAP vector computed on the PCA-projection (with 100 components) of the molecule’s ChemNet embedding (with 512 components). Different colors of LDDM samples correspond to clusters computed on the UMAP vectors using a spectral clustering algorithm with a predefined number of clusters (*n* = 10). Six panels around the map provide examples of LDDM samples from different clusters. For each cluster, we show several manually selected designs. While LDDM entirely covers the chemical space of the training set (gray background, *n* = 454 662), it produces novel chemical matter as shown in the cumulative histogram of novelty. To quantify novelty of a generated molecule, we compute its maximum Tanimoto similarity (using Morgan fingerprints with 2048 bits and radius 3) to the training set. As shown in the plot, approximately 90% of samples have less than 0.5 similarity to the training set. **(D)** Overview of the programmable and synthesizable generation algorithm. The programmable generation algorithm iteratively grows molecules within a fragment tree (left), guided by predefined global and local criteria. The synthesizable extension constrains this process to fragments and molecules that are synthetically accessible within predefined combinatorial chemical spaces. Starting from an empty protein pocket, LDDM generates an initial set of candidate molecules (center). Sampled molecules are screened against a building block database (right, blue); substructures that match building blocks and satisfy additional constraints are accepted as starting fragments for the next iteration. For each accepted fragment, accessible molecules are generated by applying reaction templates in combination with the building block database (right, green). Newly generated molecules are subsequently evaluated by substructure matching against the reaction product library.

To recover the masked information, we follow Schneuing et al. [35] and train a multi-domain generative model that combines equivariant flow matching [36] for sampling three-dimensional (3D) atom coordinates and Markov bridge models [37] for discrete atom and bond type generation. The generation process starts with sampling the missing coordinates and atom and bond types from the corresponding prior distributions. The noisy data is then iteratively refined resulting in a molecule, where all the denoised variables follow the data distributions learned by the model. LDDM is endowed with the uncertainty estimation mechanism introduced in Schneuing et al. [35], which assigns a confidence score to each atom in a generated molecule.

One of the open challenges in computational drug discovery is the limited availability of experimental three-dimensional data. A sufficient amount of chemical diversity in the training data is crucial for generative models to generalise across drug-like molecules. To train LDDM, we therefore curated a large synthetic dataset of three-dimensional protein-ligand complexes. Our dataset combines multiple data sources including CrossDocked [38], BindingNet v2 [39], and BigBind [40] and consists of 836 259 data points, substantially exceeding the fewer than 50 000 non-redundant experimentally determined protein–ligand complexes represented in the PDB [41]. We ensure high geometric, chemical, and physical quality of our data through various computational filters, as discussed in Methods Section 1.2.1, and demonstrate that training on our synthetic dataset results in improved molecular quality and diversity, compared to training only on the experimental data from the PDB (see Supplementary Information A.1). Figure 1C shows the underlying chemical space of the training set and its coverage with LDDM samples. Notably, LDDM samples, obtained on the diverse held-out test set, uniformly cover the entire chemical space while remaining structurally novel: over 90% of LDDM samples have less than 0.5 Tanimoto similarity to the training data. Examples of LDDM samples from different clusters of the chemical space are shown in the corresponding panels of Figure 1C.

LDDM is a prediction and design framework that enables various drug discovery scenarios. To evaluate its capabilities, we conducted several benchmarks across different tasks and compared LDDM’s performance with that of other specialized methods. First, we benchmarked the local docking capabilities of our model. As shown in Figure 2A, LDDM achieves state-of-the-art performance on the PoseBusters test set [42], reaching a docking root mean squared deviation (RMSD) below 2 Å in 79.5% of valid test cases. This is substantially higher than other popular local docking methods, although a perfectly fair comparison is difficult to achieve in redocking experiments due to different binding site definitions that affect the difficulty of the task. For LDDM, we follow the common strategy of selecting pocket residues using an 8 Å distance cutoff. Interestingly, AlphaFold 3 achieves comparable performance without pocket residues or even a protein *holo* structure specified, as shown in Supplementary Information A.4. At the same time, AlphaFold 3 exhibits stronger signs of memorisation compared to LDDM (Supplementary Figure A3), suggesting weaker generalisation to unseen receptor families.

**Fig. 2:**
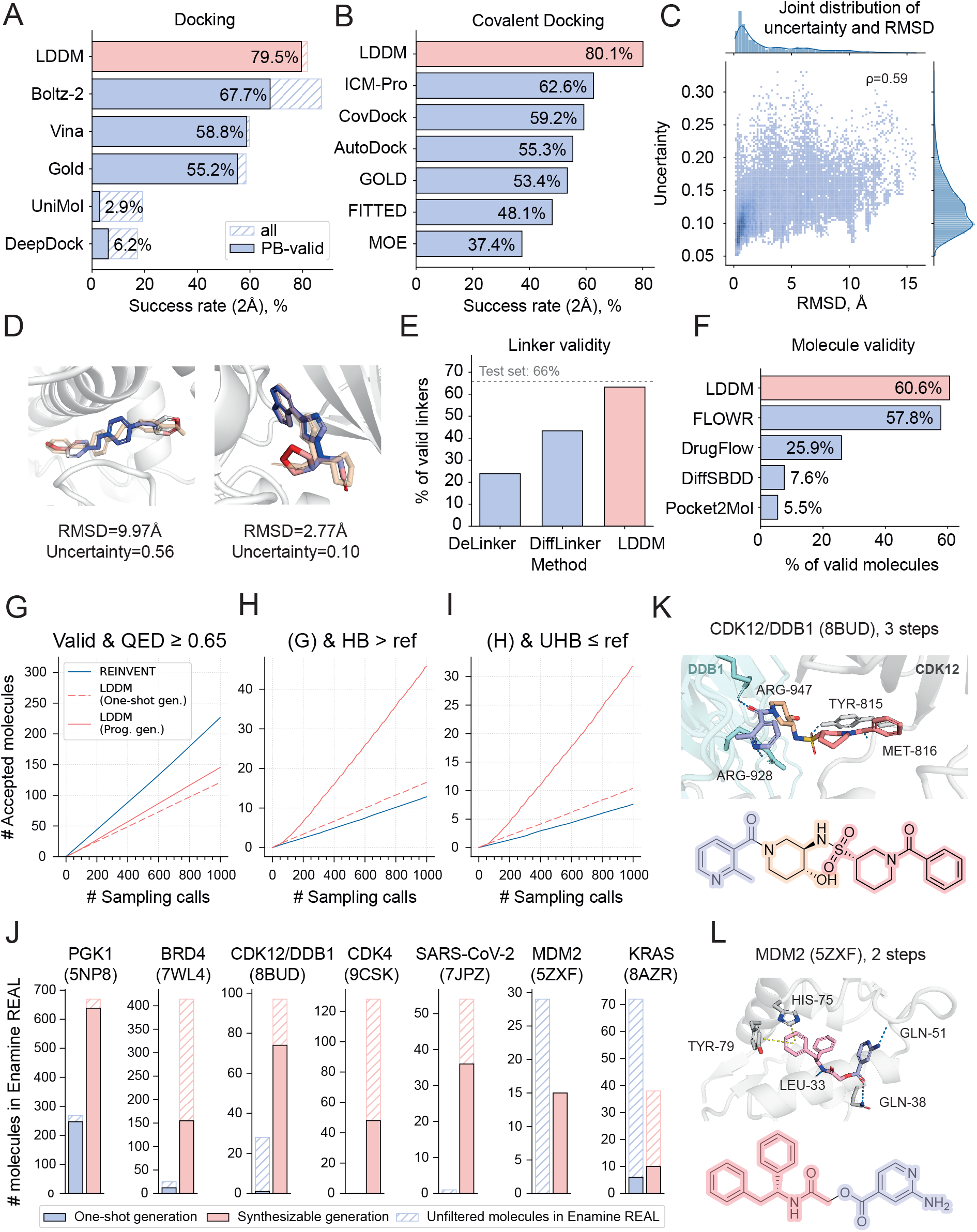
LDDM is a unified framework for computational drug discovery. **(A)** Docking results on the PoseBusters test set. For LDDM, we define a pocket as residues within 8 Å from the reference molecule, and report top-1 success rates for a 2 Å RMSD cutoff. Ranking of LDDM samples is based on the model’s uncertainty. **(B)** Covalent docking results on the test set curated by Scarpino et al. [43]. We report top-1 success rates for a 2 Å RMSD cutoff. Results for all baselines are obtained from Scarpino et al. [43]. **(C)** LDDM uncertainty is a good proxy for scoring docked poses as it correlates well with the RMSD values (Spearman’s *ρ* = 0.59). **(D)** Examples of molecules docked by LDDM (in red-blue) overlaid with ground-truth poses (beige). LDDM samples are coloured according to the per-atom uncertainty values. **(E)** Quality of the generated molecular linkers. Here, we use the Pockets test set from Ref. [44] and report the fraction of designs that pass both PoseBusters and 3D validity filters and connect all fragments. The dashed line shows the possible upper limit for the Pockets test set. **(F)** Quality of the *de novo* generated molecules. We report the percentage of designs that pass all PoseBusters filters and 3D validity filters on the PoseBusters test set. **(G–I)** Programmable generation efficiently optimises local properties. Sampling efficiency on the PoseBusters test set for REINVENT (blue), LDDM (one-shot; red dashed), and LDDM with programmable generation (red solid). Plots show the cumulative number of accepted unique molecules (averaged across targets) as a function of sampling calls. **(G)** PoseBusters-valid and 3D-valid molecules with QED ≥ 0.65. **(H)** Criterion (G) plus a greater number of hydrogen bonds (HB) than the reference ligand. **(I)** Criterion (G) plus fewer unsatisfied hydrogen bonds (UHB) than the reference. While REINVENT excels at improving global properties (QED), controlled sampling in LDDM is more effective at discovering molecules with desirable local properties, such as hydrogen bonding. **(J)** Synthesizable generation outperforms one-shot generation in producing make-on-demand molecules that satisfy both local and global filters. Bar plots show the number of molecules available in the Enamine REAL Space for each of seven protein targets (PDB codes indicated in parentheses), generated using either one-shot (blue) or synthesizable (red) generation, with an equal sampling budget of 4000 molecules. Filled bars denote the number of compounds that satisfy all filtering criteria: PB-valid, 3D-valid, at least two hydrogen bonds or more than the reference ligand, and a Vina minimization score within 2 kcal/mol of the reference ligand. Hatched bars indicate the total number of Enamine REAL matches, without downstream filtering. Synthesizable generation was performed with hybrid docking and the following fragment filters: PB-valid, 3D-valid (all targets) and at least one hydrogen bond (all targets except for PGK1 and BRD4). Only the synthesizable generation strategy consistently yields molecules that meet all criteria, while *de novo* sampling produces significantly fewer or no acceptable candidates. **(K-L)** Selected examples of molecules generated using synthesizable generation for CDK12/DDB1 and MDM2. Molecules are colour-coded based on their constituent building blocks.

Next, we benchmarked covalent docking by constraining the position of the ligand atom covalently attached to the target protein. We test our model on the dataset introduced by Scarpino et al. [43], and compare its performance to other state-of-the-art covalent docking methods studied in the same work. As shown in Figure 2B, LDDM substantially outperforms competing methods in top-1 success rate for a 2 Å RMSD cutoff. Notably, LDDM achieves this within the same general framework, performing covalent docking out of the box, while competing methods are specialized and often cumbersome to deploy.

In both docking experiments, we sampled 100 conformations for each target, and selected only one based on our model’s uncertainty score, which correlates well with the docking RMSD (Spearman’s *ρ* = 0.59), enabling reliable scoring of docking poses (Figure 2C). Notably, LDDM applies uncertainty estimation at the atomic level, identifying specific molecular substructures that may have been docked imprecisely. Figure 2D showcases two selected examples where atomic uncertainties strongly align with RMSD values, comparing ground-truth poses with LDDM-generated samples.

By design, LDDM can be conditioned on known molecular parts, making it well-suited for fragment-based design scenarios. Here, we focus on molecular linker design and compare our approach to specialised generative methods DiffLinker [44] and DeLinker [45]. Similar to benchmarking the *de novo* design capabilities of LDDM, we report the fraction of samples that pass molecular quality filters and additionally check whether all input fragments are connected. We use the same test set as Igashov et al. [44]. As shown in Figure 2E, nearly 65% of the LDDM designed linkers pass all filters, exceeding the success rate of DiffLinker by approximately 22% and resulting in validity scores similar to those of the test set itself. Finally, we consider the *de novo* design task. Here, we used LDDM to design 100 molecules for each of the 308 targets from the PoseBusters test set [42] and assessed the quality of the generated compounds. We use a strict definition of molecular validity including the entire set of PoseBusters filters [42] as well as GenBench3D validity [46]. This allows us to cover a broad range of common issues such as incorrect valency, steric clashes with the protein target, and unrealistic geometries (see Methods Section 1.5). As shown in Figure 2F, over 60% of LDDM designs pass all validity filters, slightly outperforming FLOWR [25] and representing approximately a 2-fold improvement over LDDM’s predecessor, DrugFlow [35], with an even greater improvement over other widely-used generative methods such as Pocket2Mol [20] and DiffSBDD [24].

### Programmable and synthesizable ligand design

While effectively learning the underlying chemical space is a necessary condition for successful generative drug design, it is not sufficient on its own. Candidate molecules must typically satisfy multiple criteria, including solubility, chemical stability, favourable interaction profiles, and high binding affinity. Moreover, to enable experimental validation, it is essential to guide molecular generation toward synthetically accessible regions of chemical space. Our programmable design mechanism, illustrated in Figure 1D and detailed in Methods Section 1.1.3, enables us to address these challenges.

Starting from a protein pocket, LDDM generates an initial set of candidate compounds. Molecules passing basic global filters are then fragmented and evaluated at the local level for properties such as molecular validity and number of hydrogen bonds formed with the target. Accepted fragments are used as inputs for the next round of generation, allowing the model to iteratively build more refined compounds. Final molecules are evaluated against a set of predefined *global* criteria, and those that meet all requirements are retained.

We performed this sampling procedure over 1000 iterations on the PoseBusters test set and assessed the number of unique final molecules that (a) pass molecular quality filters and have a quantitative estimate of drug-likeness (QED) [47] above 0.65, (b) form more hydrogen bonds than the reference compound, and (c) have less unsatisfied hydrogen bond donors and acceptors than the reference. Analogous to the one-shot design evaluation above, we assess the molecular quality using PoseBusters validity [42] and GenBench3D validity [46]. We compare our method with REINVENT [48], a reinforcement learning framework that is tasked with optimising QED and the relative number of satisfied versus unsatisfied hydrogen bonds between the designed molecule and the target protein. We conduct three experiments, gradually increasing the number of local and global validation filters applied.

Figures 2G-I compare the average cumulative number of unique valid molecules per target for REINVENT, one-shot generation with LDDM, and programmable generation. The median and interquartile range per target is shown in the Supplementary Figure A5A. Although REINVENT effectively optimises global properties such as QED, our controlled sampling method substantially outperforms it in local and structure-based optimisation tasks, producing over four times more compounds with improved interaction profiles. This demonstrates the strength of our approach in optimising decomposable, structure-dependent objectives that are difficult to enforce using structure-implicit methods. Furthermore, programmable generation leads to a substantial increase in sampling efficiency compared to one-shot sampling with LDDM.

Building on the programmable design framework, we then added a synthesizable generation strategy that creates molecules according to predefined chemical synthesis rules (Figure 1D). Instead of fragmentation, we perform substructure searches against a library of synthetically accessible compounds. In the initial step, this library consists of a predefined set of building blocks. For each added fragment, the library enumerates all reachable derivatives by applying a set of reactions between the fragment and the initial building block set. Further methodological details are provided in Methods Section 1.1.4. In contrast to existing hierarchical synthon-based docking methods [49], which iteratively assemble molecules through the docking of incomplete R-groups, LDDM generates complete molecules at each step and subsequently prunes them according to the reaction rules. This approach enables us to efficiently explore the entire learned chemical space, but select only the solutions which are reachable within our defined synthesis framework.

In our experiments, we restrict the synthesizable design scope to the Enamine REAL Space [50] and compare it with one-shot sampling (Figures 2J-L). To assess the efficiency of both methods, we report the numbers of designed molecules that are found in Enamine REAL Space and meet a predefined set of quality criteria. Using reactions and building blocks from Enamine, we generated 4000 molecules for seven selected target proteins using our synthesizable design algorithm and standard one-shot generation. The resulting compounds were tested for Enamine REAL Space membership and additional filter criteria assessing molecular quality, predicted Vina scores [51], and molecular interaction counts. As shown in Figure 2J, and detailed in Supplementary Figure A5B, only synthesizable design consistently yields molecules that meet all criteria, while one-shot sampling produces substantially fewer acceptable candidates. Figures 2K-L show examples of two synthetically available compounds, designed by LDDM for CDK12-DDB1 complex and MDM2, respectively.

### Generative redesign and molecular optimisation

LDDM was first tested in constrained generation and pose prediction tasks through two experimentally validated design campaigns. First, we explore new chemistry for an established VHL-CDO1 molecular glue. Second, we optimise a cathepsin S peptide inhibitor with non-natural amino acids.

#### Redesign of a VHL-CDO1 molecular glue

The von Hippel–Lindau protein (VHL) is an E3 ligase substrate receptor that targets prolyl-hydroxylated HIF1*α* for ubiquitin-dependent degradation [52] and has become commonly used for targeted protein degradation. Since Buckley et al. [53] established (4R)-hydroxyproline as the core moiety for binding that has underpinned subsequent VHL ligand and PROTAC development [54, 55], Tutter et al. [52] recently showed that a hydroxyproline-based ligand can also act as a molecular glue, recruiting cysteine dioxygenase 1 (CDO1) to VHL for selective degradation. In their work, the authors showed that modifications on both the t-butyl cap and the aryl group strongly affect CDO1 recruitment and degradation. The ternary co-crystal structure with compound 1 revealed that the hydroxyproline core retains its canonical VHL contacts while the two flanks dock into separate CDO1 pockets, with Q99 making key hydrogen bonds to the central amide (Figure 3A).

**Fig. 3:**
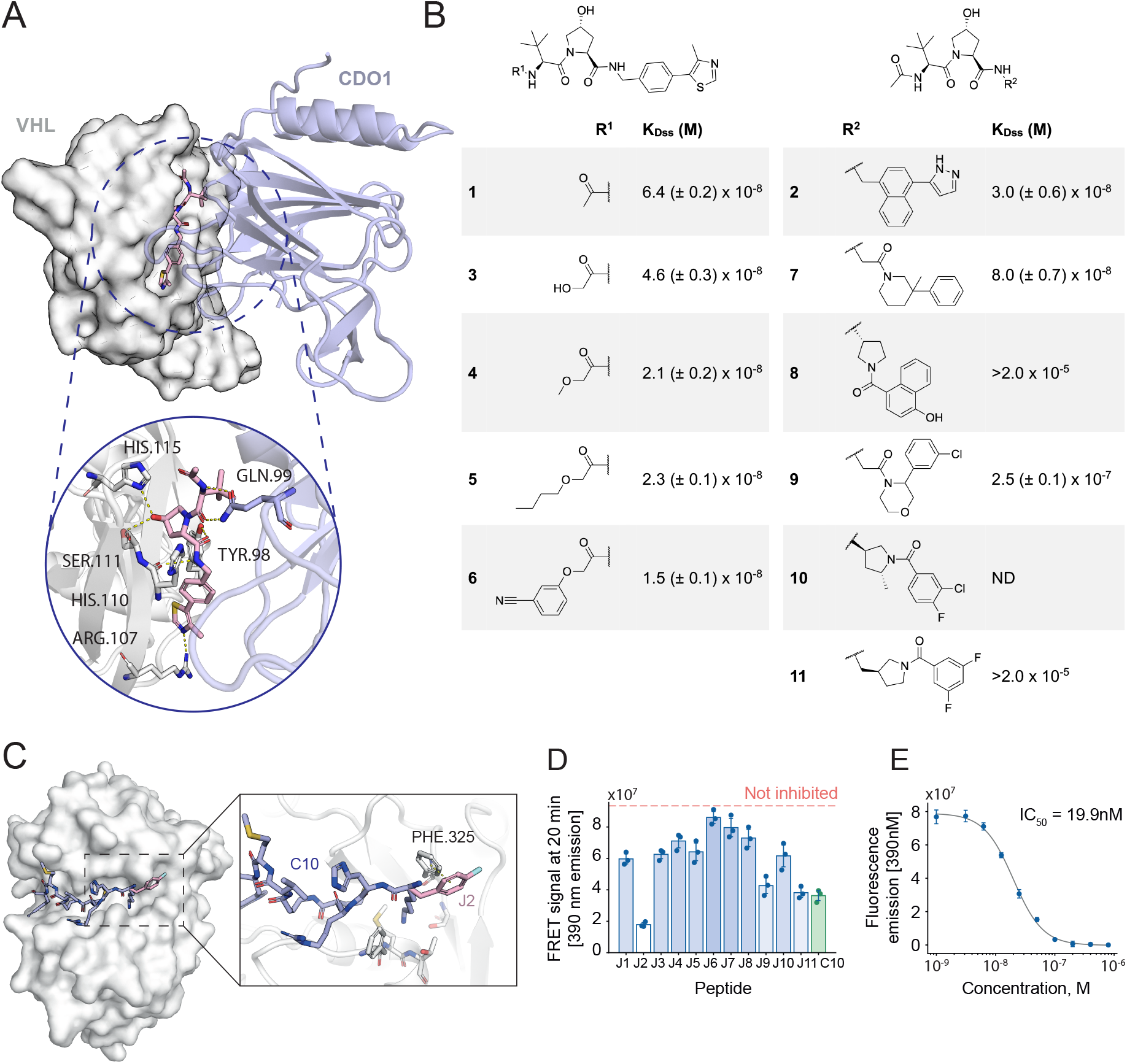
Generative redesign of VHL-CDO1 molecular glues (A-B) and cathepsin S peptide inhibitors (C-E). **(A)** Crystal structure of the VHL:CDO1:compound 1 ternary complex (PDB: 8VLB). **(B)** Previously reported compounds 1 and 2 and new modifications 3-11 generated by LDDM. For each compound, we provide the SPR-determined binding affinity (K_Dss_) on VHL-immobilized surfaces in presence of 1 µM CDO1 (*n* = 3). **(C)** The structure of the original NNPI-C10 (blue) bound to cathepsin S (PDB ID: 8PI3), overlaid with the most potent optimised candidate J2 (pink). The possible *π*-*π*-stacking interaction with F325 is represented by the dashed yellow line. **(D)** Out of eleven tested NNPI designs, one (J2) performed better than the previously optimised inhibitor C10. **(E)** Dose–response curves quantifying the inhibitory potency in FRET assays of NNPI-J2 on CTSS.

Using this structure as a template, we held the hydroxyproline-tert-leucine VHL anchor fixed and used LDDM to explore the two flanks contacting CDO1, profiling the formation of ternary complex by surface plasmon resonance (SPR). As reference, we characterised binary and ternary interactions of compounds 1, 2, and CDO1 with VHL (Figure A25 and Table A5). Using LDDM, we first grew the N-acyl cap occupying the CDO1 hydrophobic patch formed by residues P44, W47, A48, A51 and V152, which Tutter et al. [52] showed to be important for CDO1 recruitment. Of the 35 candidates retained after generation, filtering, and scoring (see Methods Section 1.7), four were synthesized and showed a dose-dependent SPR response comparable to that of compound 1 (Figures 3B and A26, and Table A6), demonstrating that LDDM can predict new modifications while retaining recruitment of CDO1 and ternary complex formation.

We next probed the opposing flank, re-designing the aryl-containing moiety that projects into the CDO1 pocket bounded by residues F53-Q55, G78 and P150. We generated 70 candidates, of which five were selected for synthesis and validation, and two reached nanomolar binding affinity (Figures 3B and A26, and Table A6). The active hits introduced scaffolds not represented in the published chemical space, broadening the accessible options at this hotspot. Across both campaigns, 67% of LDDM-introduced modifications retained nanomolar affinity for ternary complex formation.

In this study, we demonstrated how LDDM can efficiently explore new chemistry of a molecular glue without compromising its baseline potency. Although LDDM successfully redesigned both flanks contacting CDO1, (4R)-hydroxyproline itself remains the only chemotype reported to productively engage the HIF1*α*-binding groove [53, 54]. In future work, we will leverage programmable design with explicit VHL pharmacophore constraints to discover novel chemotypes engaging this site.

#### Optimisation of a cathepsin S peptide inhibitor

Cysteine cathepsins are a class of lysosomal proteases known to be therapeutically relevant targets in cancer and other diseases [56, 57]. In recent work, Petruzzella et al. [58] developed a novel non-natural peptide inhibitor (NNPI) of cathepsin S (CTSS) with reported half-maximal inhibitory concentration (IC_50_) of 70.4 nM. A crystal structure of the NNPI:CTSS complex was also reported in the study. The CTSS inhibitor, called NNPI-C10, contains a Michael acceptor which forms a covalent bond with the catalytic cysteine of CTSS upon binding and inhibits its enzymatic activity, as shown in Figures 3C and A14A. The peptidic structure of NNPI-C10 was originally optimised by saturation mutagenesis screening using all natural amino acids.

Here, we set out to further optimise NNPI-C10 by incorporating additional non-natural amino acids. To this end, we analysed the bound conformation of NNPI-C10 in complex with CTSS and identified five amino acid positions most suitable for substitution (Figure A14B). Using a library of 54 available non-natural amino acids, we generated 270 single-point mutants of NNPI-C10. For each mutant, we used LDDM to generate rotamers of the newly introduced non-natural amino acid side chain in the context of the known protein–peptide complex, taking advantage of LDDM’s capability for constrained pose prediction. We generated 200 side-chain conformations per mutant, which were evaluated using various quality metrics, as described in Methods Section 1.8.1. Out of the designs passing all the filters, the top-scoring pose was finally selected for each mutant based on the LDDM confidence score. An overview of the design workflow using LDDM is shown in Figure A14A, and more details are provided in Methods Section 1.8.1.

To shortlist the most promising candidates for experimental validation, we leveraged site-saturation mutagenesis (SSM) data for NNPI-C10 reported in the original study [58]. First, we replicated the computational single-point mutagenesis with LDDM using only natural amino acids, and trained an ensemble of linear regression models to predict mutant activity values from the original SSM matrix. For simplicity, each mutant was represented as a vector of descriptors, including the LDDM confidence score, various docking scores, and counts of polar and non-polar interactions (see Methods Section 1.8.2). As shown in Figure A16, for each of the four targeted positions, the best performing mutation was consistently ranked among the top five predictions. We then applied the trained ensemble to rank the non-natural mutants by predicted potency, and selected top designs for experimental testing (Figure A15).

Out of eleven new NNPIs tested in a FRET assay, one performed better than the experimentally optimised NNPI-C10 (Figure 3D), improving CTSS inhibition 3.5-fold from 70.4 nM to 19.9 nM (Figure 3E). In this case, the terminal leucine of the original peptide NNPI-C10 was replaced by the non-natural amino acid 4-fluorophenylalanine. As modelled by LDDM (Figure 3C), this exchange of the aliphatic side chain of leucine for a rigid aromatic ring may engage F325 of the binding site through an aromatic edge-to-face *π*-*π* stacking.

This result highlights how structure-based computational modelling can drive potency optimisation in a single iteration using only a small number of synthesized molecules. As LDDM’s training set did not include peptides, this case study also demonstrates that the model’s applicability extends beyond the class of molecules it was trained on.

### One-shot *de novo* design of PGK1 ligands

To probe LDDM’s *de novo* design capabilities, we first present a case study using the one-shot regime, i.e., without the programmable sampling algorithm. The generated designs were filtered, and only those available in Enamine’s on-demand catalog were experimentally tested.

Phosphoglycerate kinase 1 (PGK1) is a well-known enzyme that plays a crucial role in various cellular processes, including energy metabolism, angiogenesis, DNA replication and repair, and has emerged as a promising drug target [59]. In humans, PGK1 is known to be involved in oncogenesis, Parkinson’s disease and amyotrophic lateral sclerosis [60, 61]. The structure of PGK1 consists of two domains connected by a hinge region. In our experiments, we selected the nucleotide-binding pocket of PGK1’s C-terminal domain as the target site due to its functional importance and highly conserved structure across all PGK isoforms [62].

For this pocket, we designed a series of compounds using LDDM one-shot generation, and shortlisted the most promising candidates using various computational filters (see Methods Section 1.9). The top six purchasable molecules shown in Figure 4A were selected for experimental validation using grating-coupled interferometry (GCI) (Supplementary Table A2).

**Fig. 4:**
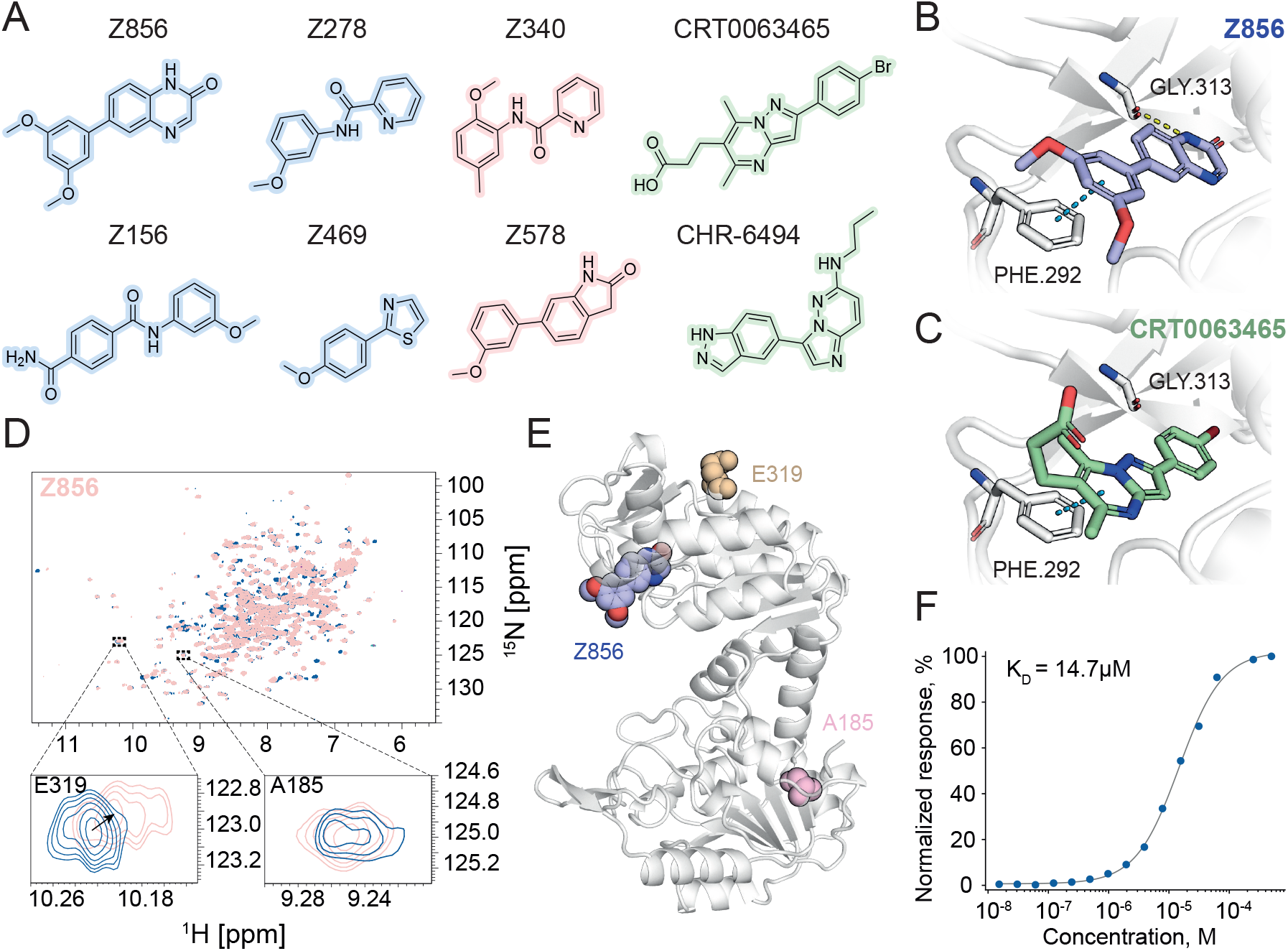
One-shot *de novo* design of PGK1 ligands. **(A)** Chemical structures of CRT0063465, CHR-6494 and the six compounds tested experimentally against PGK1. Four identified hits are highlighted in blue, and two candidates with no observed binding are highlighted in red. **(B)** The predicted bound pose of compound Z856. Amino acids involved in interactions are shown in sticks. The blue dashed line represents *π*-*π* stacking and the yellow dashed line a hydrogen bond. **(C)** Bound pose of CRT0063465 (PDB ID: 5NP8). The dashed line shows *π*-stacking. **(D)** Overlaid [^15^N,^1^H]-HSQC spectra of apo-PGK1^G185A^ (dark blue) with 0.5 mM of compound Z856 (light pink) indicates its binding to PGK1. Chemical shift of E319, with no shift observed for A185, indicates that Z856 binds to the C-terminal domain of PGK1. **(E)** Structure of PGK1 (PDB ID: 5NP8) with the binding pose of Z856 predicted by LDDM and highlighted residues E319 and A185. **(F)** Grating coupled interferometry dose-response curve for Z856 binding to PGK1 with the estimated affinity of 14.7 µM.

The initial rapid kinetic screening revealed four successful hits highlighted in blue in Figure 4A, with determined binding affinities starting from low-micromolar (Supplementary Figure A6). The best binding affinity of 14.7 µM was measured for Z856 (Figure 4F), surpassing the affinity of 30 µM determined for the positive control CHR-6494 [59] (Supplementary Figure A6).

The predicted 3D binding pose of Z856 is shown in Figure 4B. While there is no available crystal structure of CHR-6494 in complex with PGK1, we analysed the binding pose of CRT0063465, another known PGK1 ligand with the reported affinity of 24 µM [63]. As shown in Figures 4B and 4C, Z856 and CRT0063465 adopt a similar planar binding orientation and engage Phe-292 through *π*-*π* stacking interactions. Additionally, our design Z856 forms a hydrogen bond with Gly-313.

To confirm whether Z856 binds to the targeted nucleotide site of PGK1, we additionally performed NMR spectroscopy. As shown in Figure 4D, the NMR spectrum of PGK1 exhibits measurable chemical shifts upon binding of Z856. In particular, we observed a large chemical shift for Glu-319, located in the C-terminal domain of PGK1 near the nucleotide-binding site (Figure 4E), which indicates that Z856 binds to the nucleotide binding site inducing the conformational change (Figure 4E). Similarly, we validated binding of Z469, as shown in Supplementary Figure A8, and saw a similar pattern, albeit with a slightly less pronounced shift of the Glu-319 peak.

While none of the candidates was found among the previously reported PGK1 binders or in the training set (Supplementary Figure A7), our best candidate contains a quinoxalinone scaffold, similar to quinoxaline and quinazoline motifs previously explored against PGK1 [60, 64]. With only eight quinoxalinone-containing compounds in the training set, all bound to targets non-homologous to PGK1, this result suggests that LDDM can extrapolate previously seen chemistry to a new protein context.

### Synthesizable *de novo* design

Finally, in this section, we report *de novo* designs with experimental results for two additional targets. In these case studies, we used the programmable sampling algorithm to ensure synthesizability in a bottom-up fashion instead of relying on post-hoc searches of full molecules against Enamine REAL.

#### Design of BRD4 ligands

Bromodomain-containing protein 4 (BRD4) is one of the key epigenetic readers involved in the regulation of cell cycle progression, gene expression and DNA damage repair [65]. Aberrant BRD4 activity is associated with a diverse range of diseases, including different types of cancers and inflammation [66]. It has emerged as a promising pharmaceutical target, and several inhibitors are in clinical trials, but no drug has been approved by the FDA yet [67]. The well-known BRD4 inhibitor (+)-JQ1 possesses a very short half-life of 0.9 hours, which requires frequent dosing and hinders prolonged *in vivo* target engagement [68]. Other known inhibitors lack selectivity and can cause unintended side effects such as severe thrombocytopenia [69].

In our study, we targeted the hydrophobic acetyl-lysine binding pocket of BRD4, the same binding site as that of the (+)-JQ1 inhibitor. For this pocket, we designed a series of compounds using the synthesizable design algorithm restricted to the Enamine REAL Space, and shortlisted the most promising candidates using different computational filters (see Methods Section 1.10). The top eight molecules (Figure 5A) were synthesized and screened at 100 µM for binding using GCI rapid kinetics. Among them, three ligands showed binding (Supplementary Figure A17, Supplementary Table A4), and we observed a clear dose-dependent signal for compound Z967, with the estimated binding affinity in the 40 µM to 60 µM range (Supplementary Figure A17B). Details on the screening procedure and affinity measurement are provided in Methods Section 1.10.

**Fig. 5:**
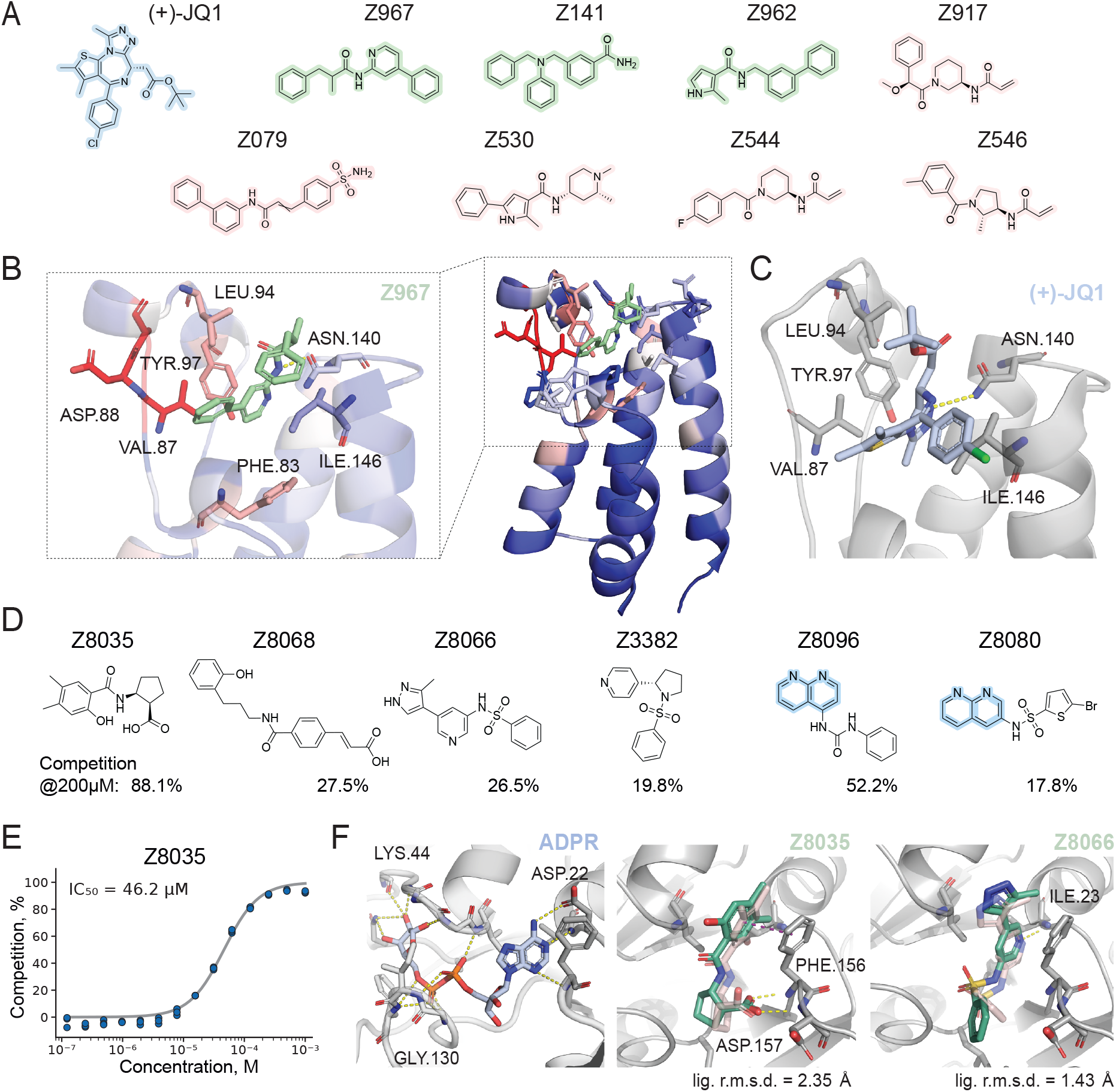
Synthesizable *de novo* design of BRD4 ligands and Sars-Cov-2 Nsp3 inhibitors. **(A)** Chemical structures of JQ1 and eight designs tested experimentally against BRD4. Three identified hits are highlighted in green, and five candidates with no observed binding are highlighted in red. **(B)** Full view and zoomed view of BRD4 (PDB ID: 7WL4) with Z967 in its predicted binding pose (green). Carbon atoms in the protein residues are colored by quantitative NMR chemical shift perturbations (CSP). Blue means 0 ppm and the gradient goes up to red for 0.15 ppm. Substantial chemical shift perturbations at VAL.87, ASP.88, TYR.97, and LEU.94 among others, indicate binding to the predicted pocket. **(C)** Crystal structure of (+)-JQ1 in complex with BRD4 (PDB ID: 3MXF). The amino acids involved in the interactions are shown in sticks, and hydrogen bonds are shown with yellow dashes. **(D)** Chemical structures of four *de novo* and two fragment-based HTRF screening hits against the SARS-CoV-2 Nsp3 Mac1 macrodomain that showed greater than 15 % competition with ADP-ribose binding at 200 µM ligand concentration. The fragment substructure is highlighted in blue. **(E)** Dose–response curve for Z8035 (HTRF ADP-ribose-peptide displacement), with an estimated IC_50_ of 46.2 µM. **(F)** Crystal structures of ADPR (PDB ID: 6W02), Z8035 (PDB ID: 38HB), and Z8066 (PDB ID: 38HC) bound to Mac1. Hydrogen bonds are shown as yellow dashed lines, and the *π*–*π* stacking interaction with F156 is shown as a purple dashed line. The designed ligands Z8035 and Z8066, based on templates PDB IDs 5SRL and 7HQL, respectively, are superimposed on the resolved crystal structures. The backbones match the crystal structures (C*α* RMSD *<* 0.3 Å). Designed ligand poses are shown in light red and have ligand RMSDs of 2.35 Å and 1.43 Å relative to the crystallographic poses.

The binding pose of compound Z967, predicted by LDDM (Figure 5B), suggests that it adopts a similar planar binding orientation to that of (+)-JQ1 in the hydrophobic pocket (Figure 5C), and that it engages the crucial residue Asn-140 through a hydrogen bond that also anchors the binding of (+)-JQ1. We further validated the binding site of Z967 with NMR spectroscopy. First, we confirmed that binding of (+)-JQ1 to BRD4 induces large chemical shifts on multiple crosspeaks of the ^1^H,^15^N HSQC spectrum (Supplementary Figure A19). Titration of Z967 into BRD4 induces chemical shift perturbations of comparable magnitudes for a similar set of residues (Supplementary Figure A22), especially for 14 residues consistent with the predicted binding pose (Figure 5C, Supplementary Figure A21). In particular, we observed large chemical shifts for Val-87, Asp-88, and Tyr-97 and smaller shifts for Phe-83, Leu-94, Asn-140 and Ile-146 for compound Z967, confirming predicted binding to the acetyl-lysine pocket.

#### Design of SARS-CoV-2 Nsp3 inhibitors

The macrodomain (Mac1) of non-structural protein 3 (Nsp3) of severe acute respiratory syndrome coronavirus 2 (SARS-CoV-2) is a mono-ADP-ribosylhydrolase that reverses host antiviral ADP-ribosylation, making it an attractive antiviral target [70]. Because there are only few inhibitors of the macrodomain enzyme family, the first potent chemotypes have only recently emerged from large-scale virtual and crystallographic screening [71]. Mac1 was also the target of the third Critical Assessment of Computational Hit-finding Experiments (CACHE #3), a community benchmark for computational inhibitor design [72]. Using previously published 3D structures of Mac1 bound to various ligands [71] and fragments [73], we designed a set of compounds targeting the ADP-ribose binding cavity of Mac1 via the synthesizable design algorithm restricted to the Enamine REAL Space [50]. After computational design and filtering (see Methods Section 1.11), 57 *de novo* and 31 fragment-based ligand candidates were successfully synthesized and screened.

Binding to Mac1 was assessed with a homogeneous time-resolved fluorescence (HTRF) competition assay in which test compounds displace an ADP-ribose-conjugated peptide from the protein. Six unique compounds (four *de novo* and two fragment-based designs) reached at least 15 % competition as shown in Figure 5D and Figure A23A. A subsequent dose-response series has confirmed concentration-dependent competition for all six hits, as shown in Supplementary Figure A23B. The most potent molecule, the *de novo* design Z8035, displayed an IC_50_ of 46.2 µM (Figure 5E).

We subsequently determined crystal structures of Mac1 in complex with Z8035 and Z8066 (Figure 5F). Both compounds occupy the adenine sub-pocket of the ADP-ribose site. The crystallographic pose of Z8035 resembles the designed pose, with a ligand RMSD of 2.35 Å. This compound forms hydrogen bonds to the backbone amines of Phe156 and Asp157, which is a stereotyped interaction defining the “oxyanion” subsite observed frequently in the initial Mac1 fragment screen [73]. Z8066 was resolved in two alternative conformations, both similar to the designed binding mode and recapitulating the predicted hydrogen bond to Ile23, with ligand RMSDs of 1.43 Å and 1.65 Å (Supplementary Figure A23D). Both compounds feature chemically distinct scaffolds, with maximum Tanimoto similarities to previously reported Mac1 ligands of 0.32 and 0.24, respectively [71] (Supplementary Figure A24). We also obtained a crystal structure of Z4996 despite its inactivity in the primary screen. Although Z4996 occupies the ADP-ribose-binding site, its crystallographic pose deviates substantially from the designed pose, with a ligand RMSD of 5.07 Å (Supplementary Figure A23C).

CACHE #3 participants collectively screened 1739 candidates across 23 workflows, among them 1592 were chemically novel; the best novel hits achieved binding affinities of approximately 18 µM [72]. By contrast, we designed and validated only 57 *de novo* candidates resulting in a chemically distinct inhibitor with an IC_50_ of 46.2 µM.

### Conclusion

Early-stage drug discovery is largely performed by various forms of experimental or computational screening methods, which are inherently limited to testing only a small fraction of all theoretically possible drug-like molecules. Computational *de novo* design methods, such as generative models, hold promise to efficiently access and explore larger chemical spaces, finding new solutions for currently undruggable target proteins and reducing early-stage discovery timelines.

This study introduces a new training framework for 3D-generative models inspired by unsupervised pre-training of large language models via masking of input tokens. In the context of molecule generation, masking is performed on the level of fragments and across modalities by removing coordinates only (docking regime) or also the molecular graph (design regime) of randomly selected fragments of the training molecules. By construction, LDDM is directly applicable to a wide range of docking and design applications, and our benchmarks show that it performs on par or better than specialized algorithms across all tested tasks.

With programmable design, we address the need for greater control over local, decomposable and structure-dependent properties of generated molecules. By leveraging the model’s fragment-based design capabilities, we iteratively place and grow building blocks within predefined chemical spaces. We demonstrate that this approach optimises local properties more efficiently than unconstrained one-shot generation and another established structure-free baseline.

Crucially, our framework explicitly addresses the bottleneck of synthetic accessibility by restricting generation to synthetically tractable virtual spaces such as the Enamine REAL Space, thereby facilitating downstream experimental validation. Although this constraint may appear to reduce the key advantage of generative modelling over screening, we view it as a pragmatic compromise required to enable experimental testing of designed compounds. The approach offers a new alternative to existing combinatorial docking strategies [49] and provides an efficient route to exploring virtual spaces that would be prohibitively costly or infeasible to screen exhaustively. In contrast to combinatorial docking, where computational cost increases with the number of building blocks and reactions, our pipeline can get more efficient with larger virtual spaces, as it requires fewer sampling calls to find a matching substructure.

In this study, we could experimentally verify binding for various generated molecules across diverse design scenarios. We managed to find and verify *de novo*-designed micromolar hits across three targets: PGK1, BRD4, and the Mac1 domain of SARS-CoV-2 Nsp3. Additionally, using the docking capabilities of LDDM, we optimised a peptide ligand for cathepsin S, even though our model has not been trained on this class of molecules. In a generative R-group exploration experiment, we showcased how LDDM can be used for diversifying the available chemistry for a VHL-CDO1 molecular glue while retaining similar levels of affinity of the ternary complex formation.

In addition to these successful experimental case studies, we also attempted to design new ligands for two targets, for which we could not yet conclusively confirm the accuracy of LDDM’s computational predictions. Three of the five *de novo*-designed ligands for KRAS (Supplementary Section A.7) show signal in GCI experiments, but the unusually high response amplitude does not allow us to rule out artifacts such as aggregation or unspecific binding (Supplementary Figure A10). We also designed covalent ligands for Pin1, which showed binding signal on GCI (Supplementary Section A.8). However, a crystal structure we managed to obtain for one of our designs, shows a significantly different binding mode than predicted. We believe that the compound’s low target affinity was not sufficient to outcompete a more favourable contact with a neighbouring subunit in the crystal (Supplementary Figure A12).

Even in the successful experiments, many of our validated ligands do not yet reach the affinities necessary for a drug candidate, probably due to their small size and limited number of interactions. These challenges can be overcome in the future by pairing the programmable design algorithm with improved ligand pose scoring and property predictors that will allow us to enforce high affinity and drug-like properties already at the hit identification stage.

Overall, our results demonstrate the practical relevance and versatile applicability of generative modelling across diverse drug discovery scenarios. While end-to-end computational design of high-affinity ligands remains challenging, LDDM consistently finds hit-level ligands with correctly predicted binding poses, requiring only a low number of synthesized compounds and a single design cycle. We anticipate that the greatest potential for streamlining the subsequent drug discovery stages lies in the iterative application of LDDM, supported by intermediate wet-lab validation and dedicated synthesis efforts. While such campaigns are beyond the scope of the present study, the hit rates and pose accuracy reported here provide a strong foundation for future work.

## Supporting information

Supplementary Information (Chemistry)

## Acknowledgements

We would like to thank Beat Fierz and Yuji Kamei for providing access to their library of non-natural amino acids, and Martin Buttenschoen for providing the baseline data for the docking benchmark. We are grateful to Kelvin Lau for his help with the biophysical assays presented in this study. We acknowledge the ESRF (Grenoble, France) for providing synchrotron radiation beamtime on beamline BM07-FIP2, supported by the French ANR PIA3 (France 2030) EquipEx+ project MAGNIFIX under grant agreement ANR-21-ESRE-0011. We thank Eric Mathieu for assistance and support as local contact during the experiment. We thank Andrea Unzue Lopez, Ingo Kober, Ingo V. Hartung, and Jörg Bomke for their valuable input that helped conduct the experiment on redesigning VHL-CDO1 molecular glues. Finally, we acknowledge Jeff Guo, Andrew Leach, Christian Dallago, and Jessica Lanini for their fruitful discussions. A.W.D acknowledges funding from the SwissAI PhD Fellowship 2025 and the AITHYRA Research Institute for Biomedical Artificial Intelligence of the Austrian Academy of Sciences. R.M.N. thanks Proxima (USA) for their support. M.B. is partially supported by the EPSRC Turing AI World-Leading Research Fellowship No. EP/X040062/1 and EPSRC AI Hub on Mathematical Foundations of Intelligence: An “Erlangen Programme” for AI No. EP/Y028872/1. D.M.F. acknowledges the Italian Space Agency (ASI) for funding the PhD fellowship of F.T. Parts of this work were performed in the context of an Innovation project supported by Innosuisse (131.374 IP-LS).

## Author Contributions Statement

I.I., A.S., A.W.D. and B.C. conceived and coordinated the project. I.I., A.S., and A.W.D. developed the machine learning model and overall methodology. I.I., A.S., A.W.D., and R.M.N. performed the computational benchmarks. I.I., A.S., A.W.D., and E.E. designed and evaluated molecules for the experimental case studies. I.M. developed and coordinated experimental validation workflows and performed the biophysical binding assays. K.Z. ran the competition assays and performed crystal soaking experiments for SARS-CoV-2 Nsp3. A.Y.L. performed structural modelling and refinement of the SARS-CoV-2 Nsp3 crystal structures. L.A.A. performed the NMR spectroscopy experiment for BRD4. A.S.P. performed the FRET assays for the cathepsin S experiments. D.R.P.I. performed various experimental validation workflows. O.G. ran the NMR experiments and analysed results for PGK1. I.F. conceived and coordinated the VHL-CDO1 redesign project, and performed chemical synthesis of all related compounds. P.M.F.S. and A.R.L. performed biophysical characterisation of VHL:small molecule:CDO1 tricomplex formation. L.K. and P.A.M.H. expressed and purified CDO1. F.T. and D.M.F. solved the Pin1 crystal structure. M.K.E.B. helped with the analysis of the BRD4 compounds. J.S. purified BRD4 protein. R.E.A., N.T., R.R., J.S.F., P.S., M.B. and B.C. supervised the work and acquired the necessary funding. I.I., A.S., A.W.D., and I.M. wrote the manuscript with input from all authors.

## Code Availability

The source code used in this study is available at https://github.com/lpdi-epfl/lddm.

## 1 Methods

### 1.1 LDDM

#### 1.1.1 Architecture

We use the geometric heterogeneous graph neural network architecture from DrugFlow [35], including self-conditioning, confidence head, and virtual nodes. Furthermore, we modify the input layer to accept three additional binary flags indicating whether a bond type, atom type or atom coordinate is known. Small molecule atoms are featurised as one-hot vectors representing 13 different types, {C, N, O, S, B, Br, Cl, P, I, F, NH, N+, O-} where charges and explicit hydrogens are included in selected cases following [35]. Protein residue nodes carry a feature encoding their type (one of the 20 standard amino acids) as well as vector-valued features indicating the positions of all atoms belonging to that residue relative to its *C_α_* position. Bond types are either single, double, triple, aromatic, or “None”. We remove all edges between pocket residues or between residues and ligand atoms that are more than 10 Å apart for computational efficiency.

#### 1.1.2 Fragment-based masked modelling

LDDM is trained as a generative model with varying degrees of conditioning. Randomly sampled parts of the training molecules are masked out and the model is tasked to recover the missing chemical moieties similar to masked language modelling. Before training, all molecules are fragmented using the breaking of retrosynthetically interesting chemical substructures (BRICS) method [31]. At each training iteration, we then select a role for each of the resulting fragments before noise is added. These roles are sampled with equal probability. Fragments can be designed, docked, or provided purely as context. If selected for design, both the molecular graph and the coordinates are noised and subsequently predicted by the model. For docking, coordinates are noised but the model has access to the molecular graph of the fragment. Context fragments are not noised at all, and the model can use their full geometric and chemical information to infer the structures of other parts of the ligand. To predict missing parts of the molecular system, we employ a generative modelling framework that integrates continuous flow matching [36] for atom coordinates and Markov bridge models [37] for categorical atom and bond types.

#### 1.1.3 Programmable generative design

Building on the substructure-conditioning ability of LDDM, the programmable generation algorithm iteratively samples candidate molecules, analyses their properties, and selects substructures that meet predefined criteria for subsequent iterations. At each iteration, two filtering functions are applied:

- *Global Filtering* discards invalid or undesired molecules based on global, non-decomposable molecule properties like QED. For the sampling efficiency benchmark (Figures 2G-I), we discard molecules with QED below the median QED of the most recent 100 sampled molecules.
- *Local Filtering* evaluates fragments individually. For the benchmark shown in Figure 2, we discard fragments with implausible 3D poses, without interactions with the target, and fragments that introduce unsatisfied hydrogen bond donors and acceptors.

Each valid fragment is stored as a node in a *fragment tree*, where each parent node is a substructure of its children. We manage the sampling budget at each iteration using a scheme inspired by the Upper Confidence Bound (UCB) for Monte Carlo tree search [74]:

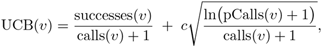

where successes(*v*) and calls(*v*) track the number of accepted molecules derived from node *v* and the total sampling attempts at *v*, respectively. pCalls(*v*) stands for the total number of samples generated from the parent of the considered node. The hyperparameter *c* controls the exploration-exploitation trade-off and was set to 1 in our experiments. The number of samples allocated to each node is proportional to its UCB score, balancing exploration of under-sampled fragments with exploitation of promising ones. Full algorithmic details are provided in Appendix (Algorithm 1).

#### 1.1.4 Synthesizable generative design

To generate molecules that are synthetically accessible, we replace the BRICS fragmentation with sub-structure searches in a building block database. Each fragment node *v* in the tree is linked to a library of reachable reaction products, precomputed from reaction templates *R* and building blocks *B* that are compatible with the current molecule. Here, we obtained a mapping of 7 053 400 synthon–reaction pairs from Enamine, comprising 306 663 unique synthons.

Newly generated molecules must contain a substructure match with one of these precomputed products to create a new child node that contains the matching substructure. This ensures that expansion pathways align with tentative synthetic routes.

To enable sampling tree expansion also for similar reaction products (i.e. partial substructure matches), we additionally employ a hybrid design/docking strategy: the maximum common substructure (MCS) is kept fixed, while unmatched atoms of the reaction product are docked flexibly. For partial docking, at most 25% of sampled molecules per iteration are selected based on MCS size, requiring ≥ 0.65 of atoms in the MCS and ≥ 0.25 Morgan fingerprint similarity. Partially docked molecules are then deduplicated based on the LDDM uncertainty score. All newly generated building blocks are subjected to the same global and local checks used in programmable generation. By combining fragment-based masked modelling with reaction-driven expansions, synthesizable generation systematically explores combinatorial fragment spaces while optimising for local properties (see Algorithm 2 in the Appendix).

#### 1.1.5 SDE sampling

For the computational benchmarks in Figure 2, we injected stochasticity into the coordinate sampling process, which we empirically found to improve the overall molecular quality. To this end, we embedded the trained flow matching model in a stochastic differential equation (SDE) following Mark et al. [75]:

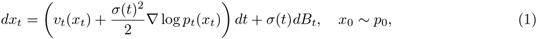

where *x_t_* is the atom coordinate at time *t*, *v_t_*(*x*) is the learned velocity field, ∇ log *p_t_*(*x_t_*) is the score, *B_t_* denotes the Wiener process, and *σ*(*t*): [0, 1] → ℝ_≥0_ is a noise schedule that can be chosen at sampling time. For our experiments, we used *σ*(*t*) = *c*(1 − *t*) with *c* = 5. When all distributions are Gaussian, the score can be directly expressed as a function of the velocity field:

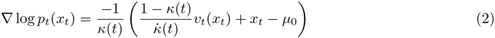

where *κ*(*t*) denotes the interpolation schedule used to train the model, here *κ*(*t*) = 1 − *t*, and *µ*_0_ is the mean of the prior *p*_0_. Note that *µ*_0_ = 0 in most standard flow matching implementations, but we center the prior in the geometric center of the pocket C*_α_* atoms, which is not necessarily located in the origin. Finally, the resulting SDE can be approximately solved using the Euler-Maruyama method.

### 1.2 Datasets

#### 1.2.1 Training dataset

To train LDDM, we curated a large synthetic dataset combining data from CrossDocked [38], BindingNet v2 [39], and BigBind [40]. While both CrossDocked and BindingNet v2 provide protein-ligand complexes with 3D information, BigBind only contains mapping between proteins and small molecules according to ChEMBL assay records. Therefore, for BigBind we first applied Gnina [51] to dock all the molecules. To ensure high quality of the training data, we then applied PoseBusters and GenBench3D validity filters. Besides, we filtered out all examples with extreme values of Gnina efficiency score, strain energy, logP, and molecular size. To detect such outliers, we used the 1st and the 99th percentiles computed over the entire data. Finally, we removed all examples with molecules similar to the molecules from the PoseBusters test set. We detected such examples using Tanimoto similarity (Morgan fingerprints with 2048 bits and radius 3) with the cutoff 0.5. The resulting dataset includes 836 259 protein-ligand complexes with 454 662 unique chemical structures.

#### 1.2.2 PoseBusters test set

Our primary test set is the PoseBusters benchmark set [42] with 308 unique ligands and proteins from the Protein Data Bank released since 2021.

#### 1.2.3 Covalent docking test set

We use the dataset curated by Scarpino et al. [43] for the covalent docking benchmark. This dataset contains 207 high-quality covalent protein–ligand complexes from the PDB with a resolution of 2.5 Å or better. The authors retained only entries with drug-like ligands that are covalently bound to the protein via a cysteine residue, as specified by the connectivity information available in the PDB files. The dataset covers 207 unique ligands with multiple different warhead chemistries as well as 54 distinct targets from various protein classes.

#### 1.2.4 DiffLinker pockets dataset

We follow the dataset curation procedure described by Igashov et al. [44] to benchmark LDDM against DiffLinker. Specifically, we build upon the protein–ligand dataset curated by Schneuing et al. [24] and apply the same linker–fragment fragmentation scheme introduced in DiffLinker, yielding a test set of 566 examples. To compare LDDM with DeLinker [45], we further exclude the 146 examples that fail DeLinker preprocessing, leaving a final test set of 420 examples. Most of these excluded cases contain phosphorus, which cannot be featurized by the DeLinker model.

### 1.3 Baselines

#### 1.3.1 Docking

Boltz-2 [76] is a co-folding model and typically predicts bound poses based on the ligand’s chemical structure and the target sequence alone. For this experiment, however, we provided a structural template of the target and specified pocket residues, thereby operating Boltz-2 in a comparable regime to the other local, pocket-conditioned docking methods. Its dependency on the degree of conditioning information is discussed in Supplementary Section A.4 together with additional methodological details. The reported success rates were obtained by computing the RMSD between the predicted ligand binding mode and the closest crystallographic ligand using RDKit’s function CalcRMS. Binding poses predicted by Vina, GOLD, UniMol, and DeepDock were kindly shared by Buttenschoen et al. [42].

#### 1.3.2 Covalent docking

Results for all baselines were extracted from the data accompanying the paper that originally introduced this covalent docking benchmark [43].

#### 1.3.3 Linker design

DeLinker [45] is a graph-based generative model that iteratively designs new linkers for two given input fragments and their relative 3D positions and orientations. It is not directly conditioned on the binding pocket structure. We use the code available at https://github.com/oxpig/DeLinker, running the ZINC-pretrained checkpoint. For each MOAD test example we cast the two reference fragments into fragment SMILES with two dummy exit atoms, plus the distance and angle between the fragment exit vectors, fix the linker length to that of the reference linker, and draw 64 samples. As DeLinker emits SMILES, we generate a 3D conformer for each sample with RDKit constrained to the reference fragment geometry, adapting https://github.com/oxpig/DeLinker/blob/master/analysis/rdkit_conf_parallel.py.

DiffLinker [44] is a 3D diffusion model that can design linkers for an arbitrary number of fragments. DiffLinker can generate linkers using the input fragments alone, but it is also available as a pocket-conditioned version. For this benchmark, we sampled molecules using a pocket-conditioned DiffLinker checkpoint that we took from the original publication.

#### 1.3.4 One-shot molecule design

FLOWR [25] and DrugFlow [35] are recent 3D target-conditioned flow matching models for small molecule design. Similar to LDDM, both models simultaneously generate molecular graphs (atom and bond types) and 3D coordinates. DiffSBDD [24] is another closely related diffusion generative model for structure-based drug design. Unlike the other baselines, it does not generate chemical bonds end-to-end but relies on a post-processing step that infers them based on the predicted coordinates and atom types. Pocket2Mol [20] is an autoregressive model that samples new atoms and bonds conditioned on the pocket context and previously placed atoms.

For each baseline, we used the published reference implementations and pre-trained checkpoints:

- FLOWR: https://github.com/jule-c/flowr
- DrugFlow: https://github.com/LPDI-EPFL/DrugFlow
- DiffSBDD: https://github.com/arneschneuing/DiffSBDD
- Pocket2Mol: https://github.com/pengxingang/pocket2mol

For a fair comparison between the similarly performing models LDDM and FLOWR, we sampled new molecules with the same size as the known reference ligand in both cases. This is not exact for LDDM, which has an end-to-end size adaptation mechanism, but can be approximately achieved by adding 5 atoms (half the maximal number of virtual nodes added during training) to the reference size. All other baselines use their own internal size sampling mechanisms.

#### 1.3.5 Property optimisation

##### REINVENT

We compare our method to the popular Reinforcement Learning (RL) framework REINVENT [48]. REINVENT employs a Recurrent Neural Network to generate molecules as SMILES (text representation) and implements goal-directed learning through RL. It thus can only be implicitly conditioned on the target protein through the RL reward. To assess REINVENT’s ability to optimise molecular interactions with a target protein, our reward function is designed to maximize the number of hydrogen bonds formed, as identified by ProLIF [77]. The reward is transformed according to:

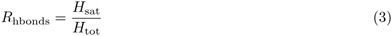

where *H*_sat_ is the number of satisfied hydrogen bonds and *H*_tot_ is the total number of unsatisfied and satisfied hydrogen bonds in the molecule. Unsatisfied hydrogen bonds are hydrogen bond acceptors and donors that do not fulfill their potential while satisfied ones do form an interaction with the protein. This transformation ensures that molecules where all potential hydrogen bond sites are involved in a hydrogen bond to the target receive a maximum reward of 1. We limit the number of oracle calls (successful docking and interaction profiling) to 1000 to ensure a fair and computationally feasible comparison with a batch size of 32. Additionally, we incorporate the QED score as a secondary objective, where molecules with a QED score above 0.65 receive a reward of 1, while those below this threshold receive a reward of 0. The combined reward is the product of the transformed QED and interaction score. Finally, to promote structural diversity among generated molecules, we apply REINVENT’s built-in diversity filter (bucket size of 25 based on identical Murcko scaffold and minimum similarity of 0.4).

### 1.4 Chemical space visualization

To visualize the chemical space covered by LDDM samples, we used ChemNet embeddings [78], mapped on the first 100 principal components using PCA and computed 2D projections with UMAP [79]. Examples for the panels were selected manually. Tanimoto similarity was measured using RDKit Morgan fingerprints with 2048 bits and radius 3.

### 1.5 Metrics

#### 1.5.1 Molecule validity

We consider molecules as valid if they pass all PoseBusters (PB) and Validity3D checks. PoseBusters filters include basic chemical sanitization, intramolecular geometry, and intermolecular, clash-related rules [42]. Validity3D evaluates whether all bond lengths, angles and torsion angles lie within the empirical distributions collected over protein-ligand complexes from PDB and is based on the GenBench3D suite [46]. Additionally, we validate the planarity of aromatic rings and assess geometry of non-aromatic rings using the same reference set of protein-ligand complexes from PDB. We typically report the fraction of molecules that passes all these filters.

#### 1.5.2 Linker validity

For the fragment linking benchmark, we also check whether the generated linkers connect all input fragments in addition to the PoseBusters and Validity3D filters discussed above. The reported number is the percentage of designs that meet all three criteria.

#### 1.5.3 Interaction profiling

We identify molecular interactions using ProLIF [77]. To count unsatisfied hydrogen bonds, we first collect all potential hydrogen bond donors and acceptors using SMARTS patterns and remove those that form hydrogen bonds with the protein as identified by ProLIF. To obtain a more conservative estimate, we also discard cases in which there is enough empty space for a water molecule to enter the binding pocket and form a hydrogen bond with the ligand atom based on idealized hydrogen bonding geometry. All remaining candidates are classified as unsatisfied hydrogen bond donors or acceptors, respectively.

#### 1.5.4 Strain energy

Ligands can adopt strained poses upon binding to adjust to the geometry of their binding pocket. To assess the internal strain of a molecule’s bound conformation, we relax generated poses without pockets using RDKit’s implementation of the Merck Molecular Force Field (MMFF94) [80]. We do not account for non-bonded interactions as these depend on context such as chemical groups in the binding pocket or surrounding solvent molecules. We compute strain energy as the energy difference Δ*E* = *E*_before_ − *E*_after_.

#### 1.5.5 Other molecular properties

Other common estimated properties, such as the quantitative estimate of drug-likeness (QED) [47], the synthetic accessibility (SA) score [81] and lipophilicity (logP) [82], are all computed with RDKit.

### 1.6 Grating-coupled interferometry

Grating-coupled interferometry (GCI, Creoptix) was used as the primary hit screening tool for BRD4, PGK1, and KRAS. The measurements were performed on a Creoptix WAVE system (Malvern Panalytical) using Creoptix WAVE control software (Malvern Panalytical, v. 4.5.18). All protein targets were immobilised on a 4PCH chip (Malvern Panalytical) by amine coupling to achieve a surface mass of 10 000–15 000 response units. The chip surface was activated with a 1:1 mixture of 100 mM N-hydroxysuccinimide and 400 mM 1-ethyl-3-(3-dimethylaminopropyl)-carbodiimide, and subsequently passivated with 1 M ethanolamine at pH 8.5 after immobilization of the protein in immobilization buffer (10mM Sodium Acetate, pH 4.5). The running buffer consisted of 20 mM HEPES, 150 mM NaCl, 1 mM DTT, 1 % DMSO, 0.005 % (v/v) surfactant P020 (GE Healthcare) for BRD4, PGK1, and PIN1. For KRAS, we also added 5 mM MgCl_2_. All compounds were tested using the Waverapid kinetic assay (repeated analyte pulses of increasing duration). The flow rate was 400 µL/min, with association pulses spanning 5 s, followed by a dissociation of 20 s. All measurements were fitted with a 1:1 binding kinetic model.

Hit identification was based on the kinetic parameters from Waverapid fitting, applying two criteria: (1) dissociation and association errors ≤ 100%, and (2) maximum response *R*_max_ ≥ 50% of the value of the positive control. The activity of all protein targets was verified by binding of positive control compounds.

Finally, full multicycle kinetics were performed for selected top hits to determine their *K*_D_ values.

### 1.7 Redesign of VHL-CDO1 molecular glues

#### 1.7.1 Computational redesign

First, we used LDDM to grow the N-acyl cap of the compound 1 occupying the CDO1 hydrophobic patch formed by residues P44, W47, A48, A51 and V152. As input, we used the crystal structure of the VHL:CDO1:compound 1 ternary complex (PDB code: 8VLB). We first designed 10 000 compounds, which were subsequently filtered using the following criteria: passed sanitization, PoseBusters, 3D validity and PAINS filters, 0≤logP≤4, 500-step constrained relaxation in RDKit has converged with RMSD≤2Å. After this filtering, the 424 remaining molecules were further scored using a simple linear model that was separately trained to discriminate between true positive and false positive bound poses. After applying a 0.95 cutoff to the computed score, we retained 35 candidates. Finally, we manually reviewed these compounds and shortlisted four candidates that were synthesized and validated in SPR.

Next, we redesigned the aryl-containing moiety of compound 1 that projects into the CDO1 pocket bounded by residues F53-Q55, G78 and P150. In this case, we designed 50 000 compounds, which were filtered using the same criteria, resulting in 7000 valid candidates. Similar scoring and subsequent selection of designs with score above 0.95 resulted in 70 compounds. We then manually reviewed these compounds and shortlisted five candidates that were synthesized and validated in SPR.

#### 1.7.2 Chemistry

Reactants, reagents, and anhydrous solvents were of commercial quality and used as purchased. Reactions were magnetically stirred and were monitored via LC-MS analysis. Nuclear magnetic resonance spectra (NMR) were recorded using Bruker AV-400, AV-500 or AV-700 spectrometers in DMSO-d6 if not indicated otherwise. The spectra were analyzed using MestReNova v15.0.1-35756 (2024 Mestrelab Research S. L.). The chemical shift (*δ*) in ppm of the reference solvent peak in the 1H NMR spectra was set according to the values described by Fulmer et al. [83].

Removal of solvents under reduced pressure was carried out at 35-50°C water bath temperature using a rotary evaporator or the V-10 solvent evaporator system from Biotage (methods: very high boiling point: 6000 rpm, 0 mbar, 56°C; volatile: 6000 rpm, 30 mbar, 36°C; high volatile: 6000 rpm, 100 mbar, 36°C). Purification by automated flash column chromatography was carried out on Isco Combiflash systems (CombiFlash NextGen 300+, Teledyne Isco) using Redisep columns using the indicated solvent systems and gradients.

Purification by preparative RP HPLC was carried out on Isco Combiflash EZ Prep systems (CombiFlash, EZ Prep, Teledyne Isco) using a SunFire Prep C18 OBD column (30 mm x 150 mm; 5 µm) or, where indicated, using a prepacked Interchim Puriflash C18 50 µm F0040 flash column.

HPLC-MS analyses were carried out on Agilent 1260 Infinity II LC-systems with Infinity Lab LC/MSD systems using one of the following methods as indicated:

**Method A**: Cortecs T3 2.7 µm 50-4.6 mm; flow: 3.5 mL/min; 45°C; MS: 85-1300 amu positive; mobile phase A: water + 0.05% HCOOH; mobile phase B: MeCN+ 0.04% HCOOH; gradient: 1-99% B (0-2.5 min), 99%B (2.5-2.9 min).

**Method B**: Chromolith HR RP-18e 50-4.6 mm; flow: 3.3 mL/min; 45°C; MS: 85-1300 amu positive; mobile phase A: water + 0.05% HCOOH; mobile phase B: MeCN+ 0.04% HCOOH; gradient: 1-99% B (0-2.0 min), 99%B (2.0-2.5 min).

#### 1.7.3 Expression and purification of CDO1

The coding sequence of CDO1 (residues 125–323) was subcloned into a pET24a vector encoding an N-terminal 6×His-SUMO-TEV-SpyTag-HRV3C fusion tag. The construct was transformed into E. coli Rosetta(DE3) cells by heat shock. Cultures were grown in LB medium supplemented with kanamycin (50 µg/mL) at 37°C with shaking until an OD_600_ of 0.8 was reached. Protein expression was induced by addition of IPTG to a final concentration of 0.5 mM, and cultures were incubated overnight at 18°C. Cells were harvested by centrifugation and lysed by sonication in lysis buffer (50 mM Tris pH 7.5, 500 mM NaCl, 25 mM imidazole, 10% (v/v) glycerol, 0.5 mM TCEP, 2 mM DTT). The lysate was clarified by ultracentrifugation, and the supernatant was incubated with Ni-NTA agarose resin (Qiagen) for 1 h at 4°C. The resin was washed with wash buffer (50 mM Tris pH 7.5, 500 mM NaCl, 25 mM imidazole, 10% glycerol, 0.5 mM TCEP), and bound protein was eluted with elution buffer (50 mM Tris pH 7.5, 500 mM NaCl, 500 mM imidazole, 10% glycerol, 0.5 mM TCEP). Eluate fractions were pooled and diluted to a final NaCl concentration of 50 mM before loading onto a HiTrap Q HP anion exchange column (Cytiva). Protein was eluted over a linear gradient of 20 column volumes from buffer A (50 mM HEPES pH 8.0, 50 mM NaCl, 10% glycerol, 0.5 mM TCEP) to buffer B (50 mM HEPES pH 8.0, 1 M NaCl, 10% glycerol, 0.5 mM TCEP). Fractions containing CDO1 were pooled and incubated overnight at 4°C with TEV protease to remove the affinity tag. The cleaved protein was concentrated and further purified by size-exclusion chromatography on a HiLoad 16/600 Superdex 200 column (Cytiva) equilibrated in SEC buffer (20 mM HEPES pH 7.5, 500 mM NaCl, 10% glycerol, 0.5 mM TCEP).

#### 1.7.4 Surface Plasmon Resonance characterisation of binary and ternary complexes

Surface plasmon resonance (SPR) experiments were performed on a Biacore 4000 instrument (Cytiva, Uppsala, Sweden) to characterise the binary interaction affinity of compounds 1 and 2 or CDO1 (125-323) toward immobilized VCB complex (VHL (54-213)-Elongin B (1-104)-Elongin C (17-112)).

Recombinant human VCB complex (10 µg/mL in 10 mM sodium acetate, pH 5.0) was immobilized on a CM5 sensor chip by amine coupling at 25°C in the presence of 1 µM ARV-771 (MedChemExpress), a known VCB-binding PROTAC, to preserve the functional binding state of the complex and improve immobilization homogeneity. Prior to protein immobilization, the chip surface was activated with a mixture of 20 mM 1-ethyl-3-(3-dimethylaminopropyl)-carbodiimide (EDC) and 5 mM N-hydroxysuccinimide (NHS) for 1.5 min. Protein was injected at 10 µL/min for 16 or 32 min, resulting in immobilization levels of approximately 1,500 to 4,500 response units (RU). The remaining activated groups were blocked with a 5 min injection of 1.0 M ethanolamine, pH 8.5. The background buffer during immobilization consisted of 10 mM HEPES (pH 7.4), 150 mM NaCl, 0.05% Tween-20 (HBS-P+).

Compounds 1 and 2, as well as CDO1 (125-323) were prepared in running buffer containing 10 mM HEPES (pH 7.4), 150 mM NaCl, 3 mM DTT, and 0.05% Tween-20. Compounds were tested at concentrations up to 1 µM, while CDO1 was tested up to 2 µM. Ten concentrations were generated using a two-fold serial dilution in running buffer supplemented with 2% DMSO.

Ternary complex formation on VCB surfaces was additionally evaluated for compounds 1-11 in the presence of a constant concentration of 1 µM CDO1. All compounds were initially tested at concentrations up to 1 µM. For compounds showing weak binding (compounds 8 and 11) or no detectable interaction (compound 10), the concentration range was extended up to 20 µM.

A DMSO solvent correction (1 – 3%) was performed to compensate for bulk refractive index variations. Binding analysis consisted of a 120 s analyte injection at 30 µL/min (association phase), followed by a 600 s dissociation phase under continuous buffer flow. Sensorgrams were double referenced by subtracting both the response from the reference surface and the buffer blank injection. Data were fitted using a 1:1 Langmuir binding model to determine kinetic and steady-state affinity parameters. A global Rmax parameter was applied to interactions with resolved binding kinetics.

### 1.8 Optimisation of cathepsin S inhibitors

#### 1.8.1 Computational single-point mutagenesis with LDDM

As our starting point, we used the crystal structure of NNPI-C10 in the complex with CTSS (PDB: 8PI3) [58]. We selected five amino acid positions for computational mutagenesis based on the profile of interactions between the starting NNPI-C10 and CTSS (Figure A16A).

Using our library of 54 non-natural amino acids, we generated 270 single-point mutants. Next, we applied LDDM to generate the coordinates of the introduced amino acid side chains. During generation, we kept the coordinates of the entire peptide backbone and other amino acids unchanged and treated them as known context.

For each mutant, we generated 200 side chain poses and selected only the ones that reduced the number of unsatisfied hydrogen bonds, increased the number of hydrophobic interactions and increased either the number of hydrogen bonds or number of *π*-stacking interactions compared to NNPI-C10. Additionally, we removed all designs with the added strain energy exceeding the added strain energy of NNPI-C10 in complex with CTSS by more than 1.5 times. Finally, we selected the best docking pose for each mutant based on the LDDM’s uncertainty score, resulting in 67 candidates in total.

#### 1.8.2 Scoring of mutants

To rank the selected candidates, we trained a separate model to predict the inhibition potency of the mutants. To train such a model, we used the site saturation mutagenesis screening performed by Petruzzella et al. [58] for the set of 20 natural amino acids.

To this end, we repeated the computational mutagenesis procedure from 1.8.1 with natural amino acids. For each of these mutants, we computed a vector of seven high-level descriptors including:

- LDDM’s uncertainty score,
- number of hydrogen bonds,
- number of unsatisfied hydrogen bonds,
- number of hydrophobic interactions,
- added strain energy,
- Vina efficiency score, and
- Gnina efficiency score.

Using these descriptors as inputs, we trained an ensemble of linear regression models to predict the inhibition potency from the mutagenesis screening reported by Petruzzella et al. [58].

Since the starting peptide in the experiment was different from NNPI-C10, we could not use the reported potency values all together across amino acid positions. Instead, we assumed independence between the values from different sequence positions, and trained four separate linear models for each position, as shown in Figure A14C. We note that we excluded the screening results for MET-4 due to the very low signal and high homogeneity of the potency values in this position.

To validate this approach, we performed cross-validation. For each position, we predicted potency using an ensemble of all models except the one corresponding to this position (as shown in Figure A14D). Despite the low overall correlation between the real and predicted potency values, our approach demonstrated high recall, always ranking the best mutation among the top 5, as shown in Figure A16B. We therefore used an ensemble of all four linear models to score 67 shortlisted NNPIs designed with LDDM.

#### 1.8.3 FRET assay

The inhibitory activity of non-natural peptide inhibitors (NNPIs) on recombinant human cathepsin S (CTSS) was assessed by FRET assay, as previously described [58]. In short, CTSS was diluted to 10 µg/mL in Assay buffer (50 mM NaOAc, 5 mM DTT, 250 mM NaCl), at pH 4.5 and incubated for 90 minutes at 37 °C to activate it. The protease was then diluted to 1 µg/mL in Assay buffer and plated in a 96-well plate (Corning, cat #3474) (50 µL/well). NNPIs were incubated with CTSS for 30 minutes at the indicated concentration. The substrate peptide DNP-KLRMKLPKP-MCA (produced by the PPCF Facility of UNIL, Switzerland) was then added to the activated CTSS at a final concentration of 20 µM (total volume per well: 100 µL). A substrate blank containing 50 µL of Assay Buffer and 50 µL of substrate peptide without any CTSS was included (final concetration of substrate: 20 µM). Fluorescent emission of the MCA dye was measured with SpectraMax iD3 (Molecular Devices) in kinetic mode for 90 minutes (excitation/emission: 330/390 nm).

### 1.9 Design of PGK1 ligands

#### 1.9.1 Computational design

To design PGK1 binders, we used the crystal structure with PDB ID 5NP8 and applied an LDDM version trained on the CrossDocked dataset in the one-shot design regime. We discarded molecules that did not pass PoseBusters validity filters [42], REOS filters [84], or had at least one ring system that was not found in ChEMBL [85, 86]. Additionally, we filtered by lipophilicity (1 *<* logP *<* 5) and Gnina re-docking consistency (RMSD *<* 0.3 Å/atom), and kept only molecules with less than 5 rotatable bonds. Eventually, we obtained 422 unique samples, which were searched in the Enamine REAL Space. We further refined the selection of 198 molecules that were directly purchasable from Enamine based on their interaction profiles, focusing on interactions with key residues such as Phe-292, Gly-313 and Ala-215 as well as other face-to-face pi-stacking interactions and hydrophobic contacts. We ordered the 6 most promising candidates for experimental validation.

#### 1.9.2 PGK1^G185A^ protein expression and purification for NMR

A preculture was grown for 5 h at 37 °C, 110 rpm in 1× Luria Broth with kanamycin, and transferred to M9 Minimal Medium: 1 L (3 g KH_2_PO_4_, 8.4 g Na_2_HPO_4_·2H_2_O, 0.5 g NaCl, 7 g D-glucose, 1 g ^15^NH_4_Cl, 10 mL MEM vitamin mix, 1 mL kanamycin, 2 mL 1 M MgSO_4_). Protein expression was induced at OD_600_ = 0.4 by addition of 1 mM IPTG, followed by overnight incubation at 25 °C. Cells were harvested by centrifugation (4000 g, 20 min, 4 °C) and resuspended in 35 mL lysis buffer: 50 mM Tris, 500 mM NaCl, 2 mM MgCl_2_, and one EDTA-free protease inhibitor cocktail tablet. Cells were lysed by three passes through a microfluidizer. Lysate was centrifuged (35 000 g, 20 min, 4 °C) and the supernatant filtered before loading onto a 5 mL HisTrap FF Ni-column (GE Healthcare), equilibrated with Buffer A: 50 mM Tris, 500 mM NaCl, 5 mM MgCl_2_, 10 mM imidazole, 2 mM BME, pH 7.4, with a flow rate of 2 mL/min using a peristaltic pump. The column was washed with 50 mL of Buffer A, and protein was eluted with 150 mM imidazole. Dialysis was performed overnight at 4 °C in 20 mM Tris, 50 mM NaCl, 1 mM MgCl_2_, 2 mM BME, pH 8.0, to remove imidazole. The His-tag was cleaved with TEV protease over two days at 4 °C. The protein solution was filtered, passed over the HisTrap column to remove the tag, and concentrated in a 10 kDa cutoff concentrator (Merck Millipore Amicon® Ultra). A PD10 column was used to exchange into NMR buffer: 20 mM HEPES, 100 mM NaCl, 5 mM MgCl_2_, 2 mM TCEP, 10 % D_2_O.

#### 1.9.3 Hit finding GCI PGK1

Screening of PGK1 designed compounds for hits was performed at 500 µM concentration using the rapid kinetic mode of the GCI method (see Section 1.6). The hit calling criteria are summarized in Supplementary Table A2. For PGK1, CHR-6494 was used as positive control with a literature *K*_D_ range of 37 µM [59]. The immobilised protein was active, with an observed *K*_D_ of 30 µM for CHR-6494.

#### 1.9.4 NMR experiments

All NMR experiments were performed on a Bruker Avance III HD 600 MHz spectrometer equipped with a TCI cryoprobe and a SampleXpress. Two-dimensional [^15^N,^1^ H]-HSQC experiments for determining binding affinities were recorded with 128 complex points in the indirect dimension 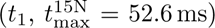 and 1024 complex points in the direct dimension 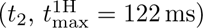. Each increment was acquired with 8 scans and an interscan delay of 0.8 s. The concentrations of ^15^N-labelled PGK1^G185A^ were 100 µM with the ligand concentration at 0.5 mM.

### 1.10 Design of BRD4 ligands

#### 1.10.1 Computational design

To design BRD4 binders, we used the crystal structure with PDB ID 7WL4 and applied synthesizable algorithm restricted to the Enamine REAL Space. In this experiment, we used the model trained on the entire synthetic dataset, and at every generation step we controlled logP between 0 and 4.0, logP normalized by the number of heavy atoms between 0 and 0.15, ensured that the 500-step UFF-relaxation with RDKit has converged with the resulting structure deviating by less that 2 Å from the designed one, and filtered out designs which introduce steric clashes or fail PoseBusters and GenBench3D validity filters. We also filtered out compounds with unsatisfied hydrogen bond donors and acceptors, as well as compounds with no non-hydrophobic interactions. Similarly to designing Mac1 binders, we ensured 3D pose consistency with GNINA, and selected top compounds by the amount of hydrogen bonds and strain energy.

#### 1.10.2 Hit finding GCI BRD4

Screening of BRD4 designed compounds for hits was performed at 100 µM concentration using the rapid kinetic mode of the GCI method (see Section 1.6). Hit calling criteria are summarized in Supplementary Table A4. The activity of the BRD4 protein target was verified by binding of positive control compound (+)-JQ1. The immobilised BRD4 protein was active, with an observed experimental *K*_D_ of 30 nM for (+)-JQ1, and with a literature *K*_D_ range of 50 nM to 90 nM [87].

#### 1.10.3 Protein expression and purification for NMR

The BRD4 construct was cloned into the LM670 expression vector by Golden Gate assembly. Plasmids LM670 was kindly provided by Lukas Milles and were previously described in Wicky et al. [88]. The resulting plasmid was transformed into BL21 T7 Express E. coli cells for protein expression. For isotopically labelled protein production, transformed cells were grown in 2 L of M9 minimal medium supplemented with ^15^NH_4_Cl and ^13^C-glucose as the sole nitrogen and carbon sources, respectively, to obtain uniformly ^15^N/^13^C-labelled protein. Cultures were grown at 37 °C until an OD_600_ of approximately 0.8 was reached, after which protein expression was induced with 0.5 mM IPTG. Expression was carried out overnight at 18 °C with shaking at 200 rpm. Cells were harvested by centrifugation and lysed by sonication under native conditions. The clarified lysate was loaded onto a His-affinity column packed with Qiagen Ni-NTA Superflow resin. The resin was washed to remove non-specifically bound proteins, and His-tagged BRD4 was eluted using imidazole-containing buffer. The eluted protein was incubated overnight with TEV protease in PBS supplemented with 2 mM TCEP to remove the affinity tag. Following cleavage, the sample was passed over a second His-affinity column to remove uncleaved protein, His-tagged TEV protease, and the cleaved His tag. The flow-through containing cleaved BRD4 was concentrated using Amicon centrifugal concentrators. Final buffer exchange was performed by repeated dilution and concentration in Amicon concentrators into 10 mM sodium phosphate, 100 mM NaCl, pH 7.5. The purified protein was concentrated to the desired final concentration for downstream experiments.

#### 1.10.4 NMR experiments

NMR experiments on BRD4 were performed on a Bruker Avance Neo 800 MHz spectrometer equipped with a TCI Cryoprobe. Two-dimensional [^15^N,^1^H]-HSQC experiments were recorded with 128 complex points in the indirect dimension and 1024 complex points in the direct dimension Each increment was acquired with 8 scans and interscan delays of 0.8 s to 1 s. The concentrations of ^15^N-labelled proteins were all around 90 µM to 100 µM, with the ligand concentrations spanning 0, 0.25, 0.5, 1, 1.5, 2.5, 4, and 8 equivalents. To interpret the BRD4 titrations, we assigned the backbone resonances by means of HNCA, HN(CO)CA, CBCA(CO)NH, HNCACB and HNCO triple-resonance experiments, aided by NMRtist [89] and checked and completed by manual inspection in the software CARA [90]. Spectral acquisition and processing were carried out with Bruker TopSpin version 4.5, and titrations were analyzed with NMRFAM-SPARKY [91] plus tools from the PDB manipulation suite [92].

### 1.11 Design of SARS-CoV-2 Nsp3 ligands

#### 1.11.1 Computational design

##### Preparation of pocket templates

As starting points for design, we assembled an ensemble of experimentally determined SARS-CoV-2 Nsp3 macrodomain (Mac1) structures from crystallographic fragment- and ligand-screening campaigns [71, 73], comprising 256 ligand-bound and 291 fragment-bound complexes. All structures were superimposed and their binding pockets compared pairwise with USALIGN [93]. Using the resulting pocket-RMSD matrix, we clustered the pockets with *k*-medoids, selecting the number of clusters by silhouette score, and used the cluster medoids as non-redundant pocket templates spanning the conformational diversity of the adenosine sub-pocket.

For *de novo* design, we used the cluster representatives as pocket templates, retaining 12 ligand-bound structures (5sp6, 5spy, 5sqr, 5srl, 5srv, 7bf5, 7h0t, 7h17, 7hhw, 7hql, 8azm, 8tv7) as design contexts. For fragment-based design, we used crystallographic fragments as starting points, keeping their bound pose as the seed for growing (templates 5s4f, 5s3l and 7hpk).

##### Synthesizable de novo and fragment-based design with LDDM

We generated ligand candidates with LDDM and the synthesizable design algorithm (see Section 1.1.4), restricted to the Enamine REAL synthon space [50]. In the *de novo* mode, LDDM generated complete ligand poses inside each pocket template. In the fragment-based mode, the bound fragment was retained as a starting seed in the first iteration of the algorithm.

##### Scoring and selection

Candidates were filtered on the fly to retain only drug-like molecules (0 ≤ logP ≤ 4, normalized logP ≤ 0.15, ESOL ≥ −4) with a force-field-relaxed pose (RMSD ≤ 2 Å), low internal strain (strain energy ≤ 2 per heavy atom), valid 3D geometry, no steric clashes, and no unsatisfied hydrogen bonds.

We pooled all *de novo* and fragment-based candidates, excluded molecules below 20 heavy atoms, and scored the remainder along several orthogonal metrics. First, we re-docked each design with GNINA [51] and retained poses that were self-consistent, i.e. the re-docked pose lay within 3 Å of the energy-minimised pose with a comparable Vina score. Second, we co-folded each protein-ligand complex with AlphaFold3 [94] and Boltz [76] and flagged designs whose designed pose was reproduced by co-folding (RMSD *<* 3 Å with interface confidence ipTM *>* 0.7 for AlphaFold3 or ligand-ipTM *>* 0.85 for Boltz). Third, we retained designs with a high Boltz-predicted binding probability (*>* 0.30 at a co-folded pose within 3 Å). Finally, we applied a learned scorer: a gradient-boosting ensemble trained on a SARS-CoV-2 Mac1 cross-docking dataset to distinguish native-like from decoy binding poses, using physicochemical descriptors, pharma-cophoric hydrogen-bond and *π* counts, surface shape and electrostatic complementarity, Vina/GNINA ligand efficiencies, and interaction counts as features. To avoid information leakage, training and validation complexes were split by ligand similarity (Tanimoto cutoff 0.3), and designs scoring above 0.7 were retained.

A design was shortlisted if it satisfied at least one of these criteria. We then removed near-duplicates with a greedy Tanimoto filter (cutoff 0.5), prioritising designs that satisfied more criteria and had stronger interaction profiles. This yielded 102 candidates (61 *de novo* and 41 fragment-based), of which 88 (57 *de novo* and 31 fragment-based) were successfully synthesised from the Enamine REAL collection and experimentally tested.

#### 1.11.2 Experimental methods

##### Inhibition experiments

The inhibitory activity of the designed compounds against Mac1 was assessed by the displacement of an ADPr-conjugated biotinylated peptide (ARTK(Bio)QTARK(Aoa-RADP)S; Cambridge Peptides, Birmingham, UK) from His6-tagged Mac1 using an established HTRF screening assay [73]. Compounds were dispensed into white ProxiPlate-384 Plus (PerkinElmer) assay plates using an Echo 525 liquid handler (Labcyte). Binding assays were conducted in a final volume of 16 µL with 12.5 nM Mac1, 200 nM peptide, 1:20 000 anti–His6-Eu^3+^ cryptate (PerkinElmer AD0402), and 1:500 streptavidin-XL665 (PerkinElmer 610SAXLB) in assay buffer [25 mM HEPES (pH 7.0), 20 mM NaCl, 0.05% bovine serum albumin, and 0.05% Tween-20]. Assay reagents were dispensed into plates using a Dragonfly Discovery liquid handler (SPT Labtech). Protein and peptide were dispensed and preincubated for 30 min at room temperature. The HTRF reagents were then added for a 1-hour incubation. Fluorescence was measured on a PerkinElmer EnVision multimode microplate reader using a dual-emission protocol (A = excitation 320 nm, emission 665 nm; B = excitation 320 nm, emission 620 nm). Raw data were processed to give an HTRF ratio (channel A/B × 10 000). Data are presented as mean ± SD of three technical replicates. All 88 compounds were first screened at two concentrations (200 µM and 50 µM) to assess inhibitory activity. The 6 compounds with greater than 15% competition were subsequently profiled in a 14-point dose-response series ranging from 0.122 µM to 1000 µM. The data were fit to a four-parameter Hill equation constrained to [0, 100]% to extract an IC50. ADPr was included as a positive control to confirm assay performance.

##### Protein crystallization and structure determination

SARS-CoV-2 Nsp3 Mac1 (residues 3–169) was expressed and purified as previously described [73]. Purified protein was concentrated to 40 mg/mL in 20 mM Tris-HCl (pH 8), 150 mM NaCl, 5 % glycerol, and 2 mM DTT, flash-frozen in liquid nitrogen, and stored at −80 °C.

Crystals for soaking were grown as previously described [73]. Briefly, crystals were grown by sitting-drop microseeding in 96-well SwissCI three-well plates by combining, in order, 200 nL protein (40 mg/mL), 100 nL reservoir solution (28 % PEG 3000 and 100 mM CHES, pH 9.5), and 100 nL seed stock (1:100 000). Crystals reached full size within 12 h to 24 h at 20 °C. Fragments (100 mM in DMSO) were dispensed onto drops with an Echo 650 acoustic liquid handler to a final concentration of 10 mM (10 % DMSO) and soaked for 2 h to 4 h. Crystals were cryo-cooled directly in liquid nitrogen without added cryoprotectant. Diffraction data were collected at 100 K at Stanford Synchrotron Radiation Lightsource (SSRL) beamline BL12-1 and processed with xia2 [95]. Structures were solved by molecular replacement using PDB ID 7KQO as the search model. Ligand-bound states were identified and modeled using PanDDA [96] and Bandicoot, a macOS-native fork of Coot 0.9.8.95 [97] (https://github.com/fraser-lab/bandicoot), followed by refinement with phenix.refine [98].

## Appendix A Extended Results

### A.1 Dependency of Molecular Quality on the Training Dataset

We additionally trained LDDM on Protein Data Pank (PDB). To curate this dataset, we selected all protein-ligand complexes released before 30 September 2019 and used the Ligand Expo database to assign covalent bond types to all ligands and filtered out the ones from which we could not match the chemical structures with the corresponding Ligand Expo [99] representatives. Additionally, we removed all molecules that did not pass the RDKit sanitization procedure. Besides, we filtered out all ligands that were labeled as invalid in the latest BindingMOAD release [100]. Next, we curated our own additional list of compounds that were abundant in our collection but yet of little interest (typically sugars, organophosphates and fatty acids). We also excluded pairs in which less than 50% of ligand heavy atoms are in contact (i.e. within 5Å) with the protein chain. Finally, we applied the same filters as to our synthetic dataset, to ensure equally high quality of the final training dataset, derived from PDB. We then sampled molecules on the PoseBusters test set, using two models with identical configurations (both training and generation), but trained on different datasets. As shown in Table A1, the model trained on the synthetic dataset generates substantially higher-quality and more diverse molecules.

**Table A1:**
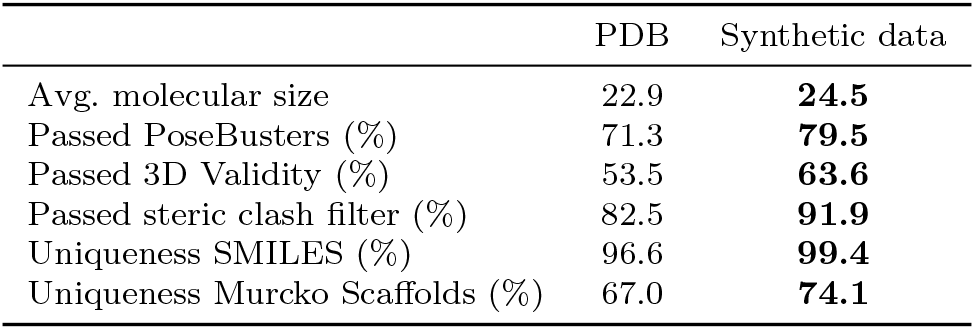
Dependency of the molecular quality on the training dataset. Quality metrics of molecules sampled on PoseBusters test set (100 molecules per target, 308 targets) using two identical models trained on different datasets.

### A.2 Recovery Experiments

To showcase the benefits of structure-based generative design over other alternative computational drug discovery strategies, we analyse the recovery of chemical features from known ligands and benchmark LDDM against docking-based virtual screening (VS) and the reinforcement learning framework REINVENT [48]. To this end, we compared all molecules with the reference ligands from the PoseBusters test set using a shape and colour similarity metric SC_RDKit_ [101, 102]. The colour similarity function compares two 3D conformers based on the overlap of their pharmacophore features, while the shape similarity measure is a simple volumetric comparison between the two conformers [45]. For each target, we select the candidate with the highest similarity to the reference compound, and provide the cumulative success rate for the whole range of similarity values (i.e. between 0 and 1) over all PoseBusters test targets. In the absence of a universal similarity cutoff, we report the area under the curve (AUC) of the cumulative success rate (Figure A1A). As positive and negative controls, we report shape-colour similarities of re-docked reference ligands and randomly selected molecules, respectively. All docking experiments are conducted using variants of AutoDock Vina [103]. Further details on baseline configurations and evaluation metrics are provided in Section B.4. We additionally report the same metrics on the subsets PostBusters test set clustered based on the pocket and ligand shape similarity to the training set, demonstrating high generalization capabilities of LDDM (see Supplementary Section A.3). As shown in Figure A1A, LDDM demonstrates a considerably higher ability to recover shape and pharmacophore patterns of the reference compounds.

**Fig. A1:**
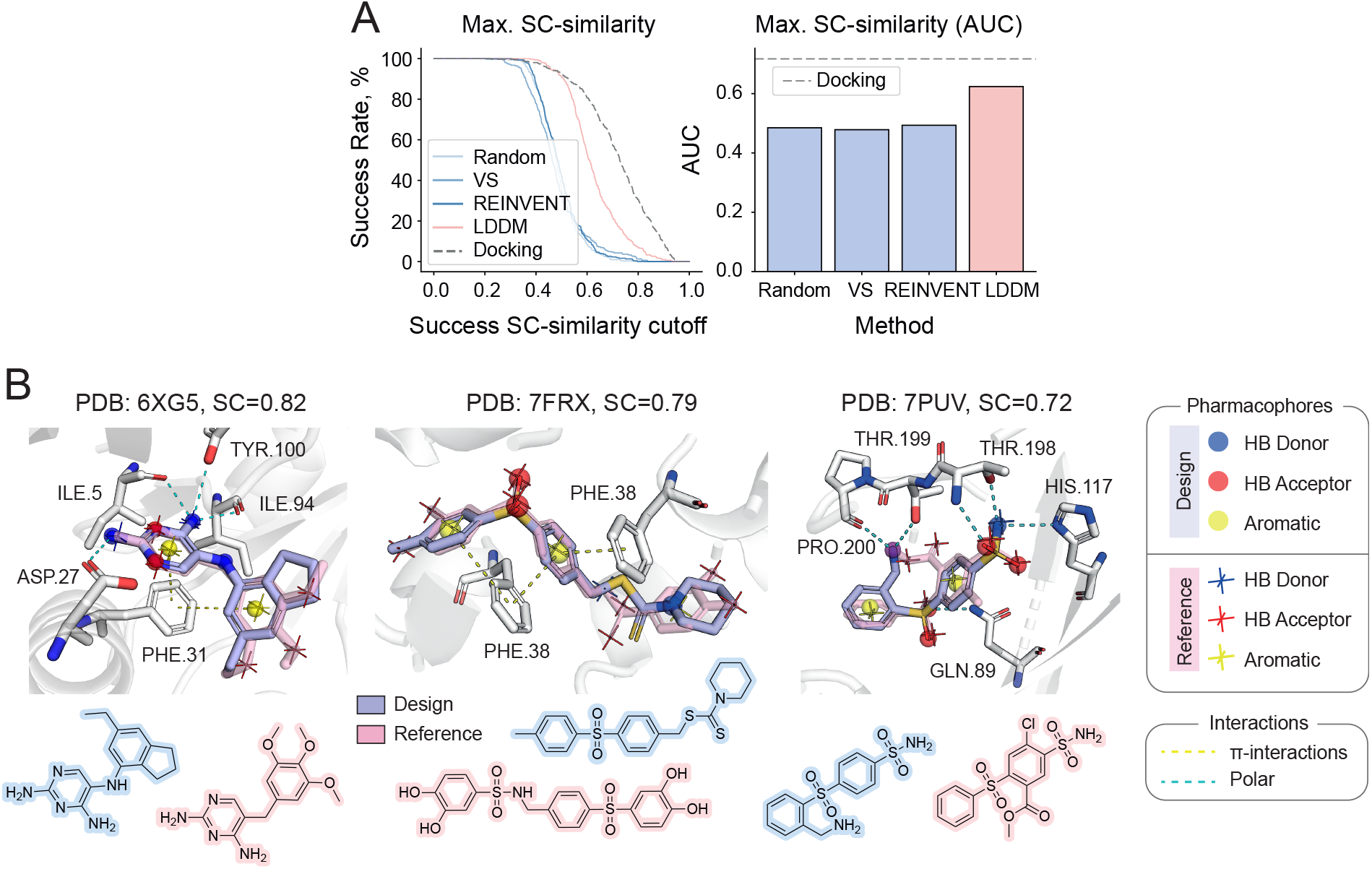
Recovery of the reference binding signal. **(A)** We use SC_RDKit_ to asses shape and colour (i.e. pharmacophore-based) similarity between generated 3D molecules and reference compounds from the PoseBusters test set (308 targets). In the absence of a universal similarity cutoff, we report the area under the curve (AUC) of the cumulative success rate for the whole range of similarity values (i.e. between 0 and 1). As a positive control, we evaluate docking poses of the reference molecules (100 conformations per target) obtained using AutoDock Vina (dashed gray lines). **(B)** Three selected examples of LDDM samples (red) with high similarity to the reference molecules (blue). All three examples demonstrate high volumetric (shape) and pharmocophore (colour) overlap. Pharmacophores are shown as spheres (for designed molecules) and crosses (for reference molecules) with colors assigned according to the panel on the right. Polar and *π*-interactions are represented as blue and yellow dashed lines, respectively. At the bottom we provide chemical structures of the designed and reference compounds with blue and red backgrounds, correspondingly.

To illustrate the meaning of the chosen similarity metric, we selected several examples, where LDDM designs achieve high shape-colour similarity to the reference molecules (Figure A1B). In all cases, the designed and ground-truth molecules have similar coarse-grained topologies and high volumetric overlap. Notably, heteroatoms with similar properties are often placed in the same regions of the pockets, yielding higher colour similarities, ultimately making intuitive chemical sense.

### A.3 Performance of LDDM Based on the Similarity to the Training Set

To estimate the level of memorization by LDDM, we computed the pocket and ligand shape similarity between the PoseBusters test set and our synthetic training set, following the Runs’N’Poses benchmark [104]. For each PoseBusters test example, we then selected the most similar training data point and used the corresponding similarity value to define the cluster of the test example. Overall, we created five clusters corresponding to the following similarity intervals: (0.0, 0.2], (0.2, 0.4], (0.4, 0.6], (0.6, 0.8], and (0.8, 1.0]. We then evaluated the performance of LDDM on all design and docking tasks independently for each of these clusters.

As shown in Figures A2A and A2B, LDDM achieves the highest performance on the subset of examples most similar to the training set in both docking and recovery benchmarks. However, the performance degradation on less similar examples is minor in both tasks, which demonstrates a low level of memorization and a high level of generalizability of LDDM to new examples. The quality of the designed molecules also does not strongly depend on the similarity of the input pocket to the training pockets, as shown in Figure A2C.

**Fig. A2:**
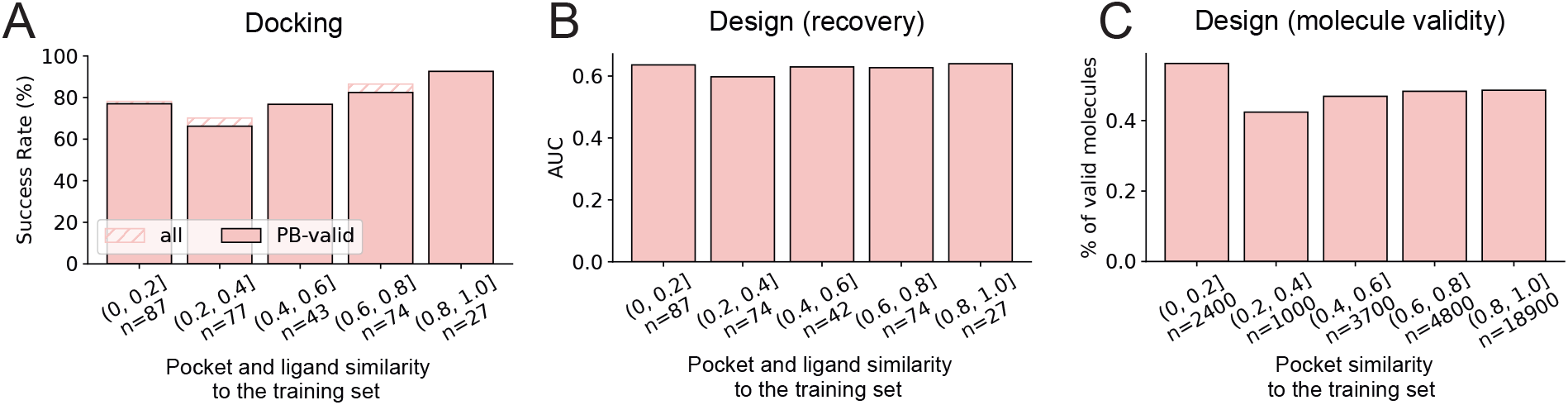
Docking (A) and design (B-C) results based on the similarity to the training set.

Additionally, we compare the performance of LDDM and AlphaFold 3 (AF3) depending on the presence of the protein-ligand complexes in each method’s training data. We note that AF3 was trained on a different training set, and therefore our test set was split on similarity bins for AF3 and LDDM independently. In this experiment, we used the co-folding results published by Abramson et al. [94]. As shown in Figure A3, the results we have obtained for AlphaFold 3 on the PoseBusters test set are in agreement with the previously reported observations on the larger Runs N’ Poses benchmark [104]. Notably, LDDM exhibits substantially weaker signs of memorisation.

**Fig. A3:**
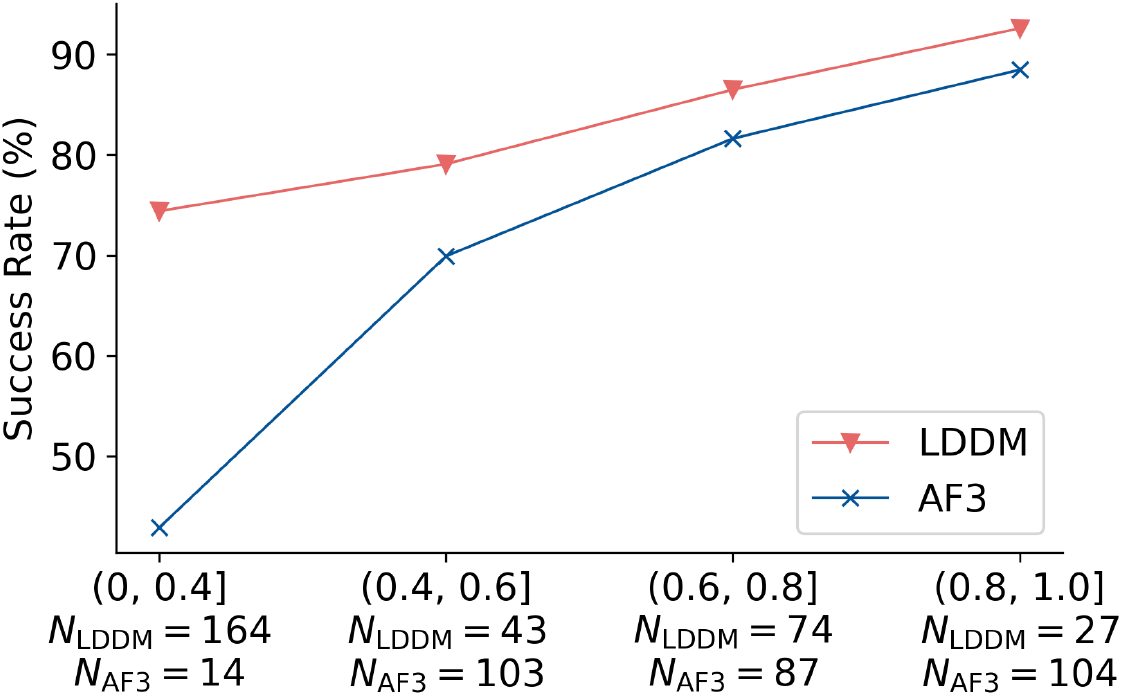
Memorisation effects of LDDM and AlphaFold 3 on the PoseBusters test set. Since the methods were trained on different datasets, we computed the similarity between the PoseBusters test set and both training sets independently. The size of the similarity bin is provided on the x-axis for both methods.

### A.4 Additional Results for Co-folding Methods on the PoseBusters Test Set

We benchmarked the co-folding methods AlphaFold 3 [94], Boltz-1 [105] and Boltz-2 [76] on the Pose-Busters test set under input-conditioning regimes of increasing information content: single sequence, MSA only, MSA with structural templates, and (for Boltz-2) MSA with templates and specified pocket residues. For every target we kept only the chains with at least one heavy atom within 6 Å of a ligand heavy atom, and report the top prediction selected by the model’s own confidence ranking. Following the PoseBusters protocol [42], we computed the ligand RMSD to the crystal pose after aligning the predicted and reference pockets, and counted a prediction as a success if its ligand RMSD is at most 2 Å. We additionally report the PoseBusters-valid success rate, which counts only successful poses that simultaneously pass all PoseBusters physical validity checks. All methods are evaluated on the 303 targets for which every method and conditioning regime produced a prediction; five targets (7K0V VQP, 7ROU 66I, 7SUC COM, 8C5M MTA, 8DP2 UMA) were excluded because the pocket-conditioned Boltz-2 runs failed to produce a prediction for them.

Figure A4 summarises the docking performance. Success rates increase strongly with the amount of conditioning information: single-sequence inputs are essentially unusable for AlphaFold 3 (1.3% at 2 Å) and weak for Boltz-2 (33.1%), the latter likely reflecting data leakage from its later training cutoff. Adding an MSA (computed following the Boltz protocol) raises the 2 Å success rate to 70.5% for AlphaFold 3 and 76.9% for Boltz-2. Richer conditioning likewise shifts the cumulative RMSD curves towards lower thresholds, and a large fraction of the sub-2 Å poses additionally satisfy the full set of PoseBusters validity checks.

**Fig. A4:**
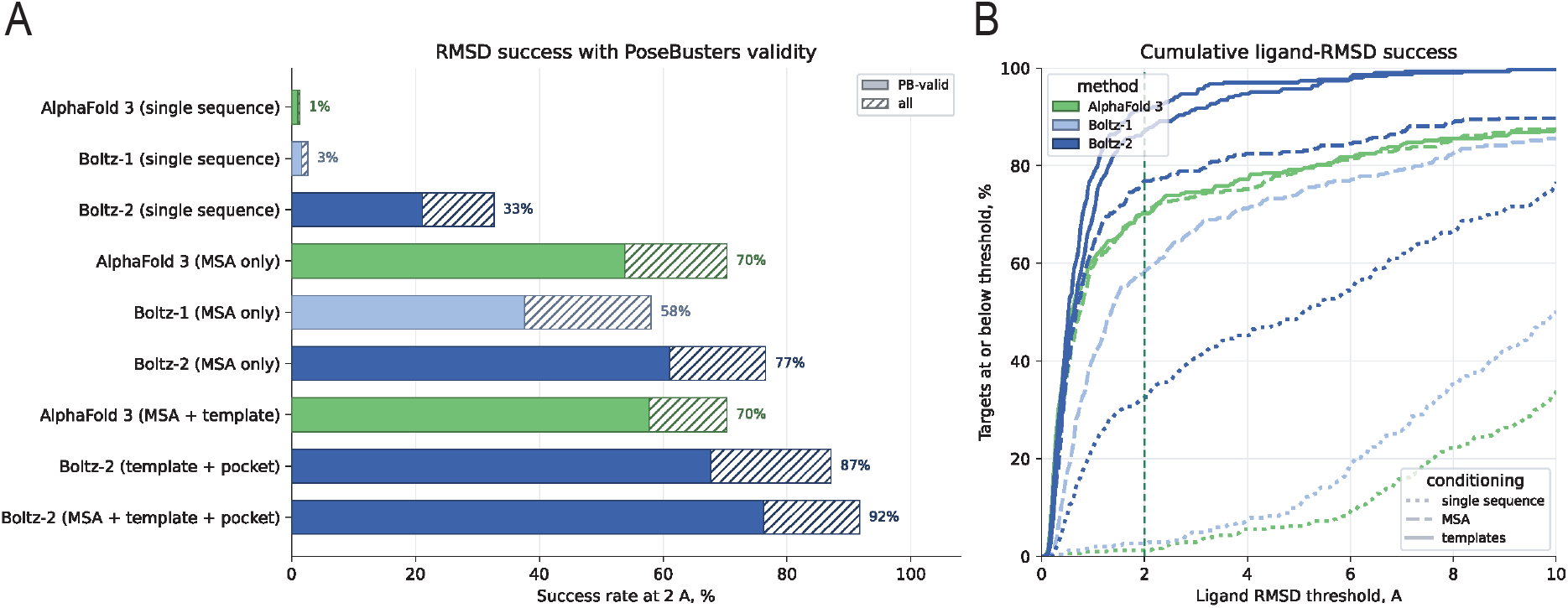
Docking performance of co-folding methods on the PoseBusters test set. **(A)** Success rate at the 2 Å ligand-RMSD threshold for each method and conditioning regime, computed on the confidence-ranked top prediction after pocket alignment. Solid bars indicate poses that are both within 2 Å and pass all PoseBusters validity checks; hatched extensions add the remaining poses within 2 Å that fail at least one validity check, so the full bar gives the unconditioned 2 Å success rate. **(B)** Cumulative fraction of targets whose confidence-ranked prediction has a ligand RMSD at or below a given threshold. Colour encodes the method (AlphaFold 3, Boltz-1, Boltz-2) and line style the conditioning regime (single sequence, MSA, templates); the dashed vertical line marks the 2 Å threshold.

### A.5 Extended Programmable Design Results

**Fig. A5:**
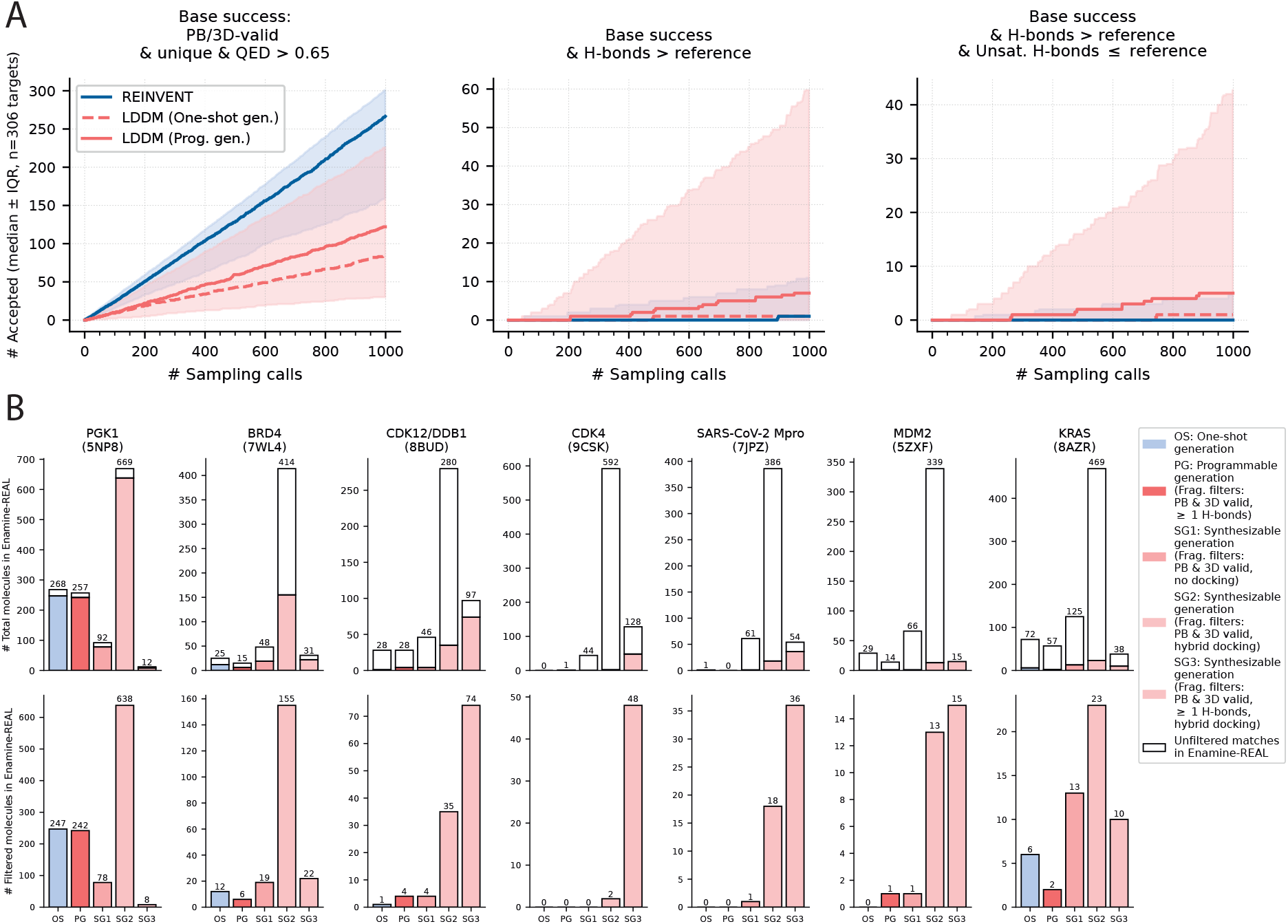
Extended Programmable Design Results. **(A)** Extended programmable generation results. Median sampling efficiency on the PoseBusters test set is shown for REINVENT (blue), LDDM *de novo* (red dashed), and LDDM with programmable generation (red solid). Shaded regions indicate interquartile ranges across targets. Plots report sampling success rates under increasingly strict criteria, as defined in Figure 2G–I. While programmable generation consistently outperforms REINVENT in optimizing local properties, sampling success varies substantially across targets. **(B)** Extended synthesizable generation results. Bar plots show the number of molecules available in the Enamine REAL Space that additionally satisfy constraints defined in Figure 2J. The lower row provides an enlarged view of the upper row, displaying only the fraction of molecules that meet all criteria. For seven selected targets, 4,000 molecules were generated using *de novo* generation (blue), programmable generation with the same filter setting as in panel (A) (red), and three different settings for synthesizable generation (pink): (SG1) fragments filtered by 3D validity and PoseBusters validity; (SG2) same fragment filters as SG1, with hybrid docking of similar reaction products enabled; (SG3) same as SG2, with the additional requirement that fragments form at least one hydrogen bond with the target. Synthesizable generation produces the largest number of molecules meeting all criteria compared to other methods, although the optimal filter setting depends on the target.

### A.6 Design of PGK1 Ligands

**Fig. A6:**
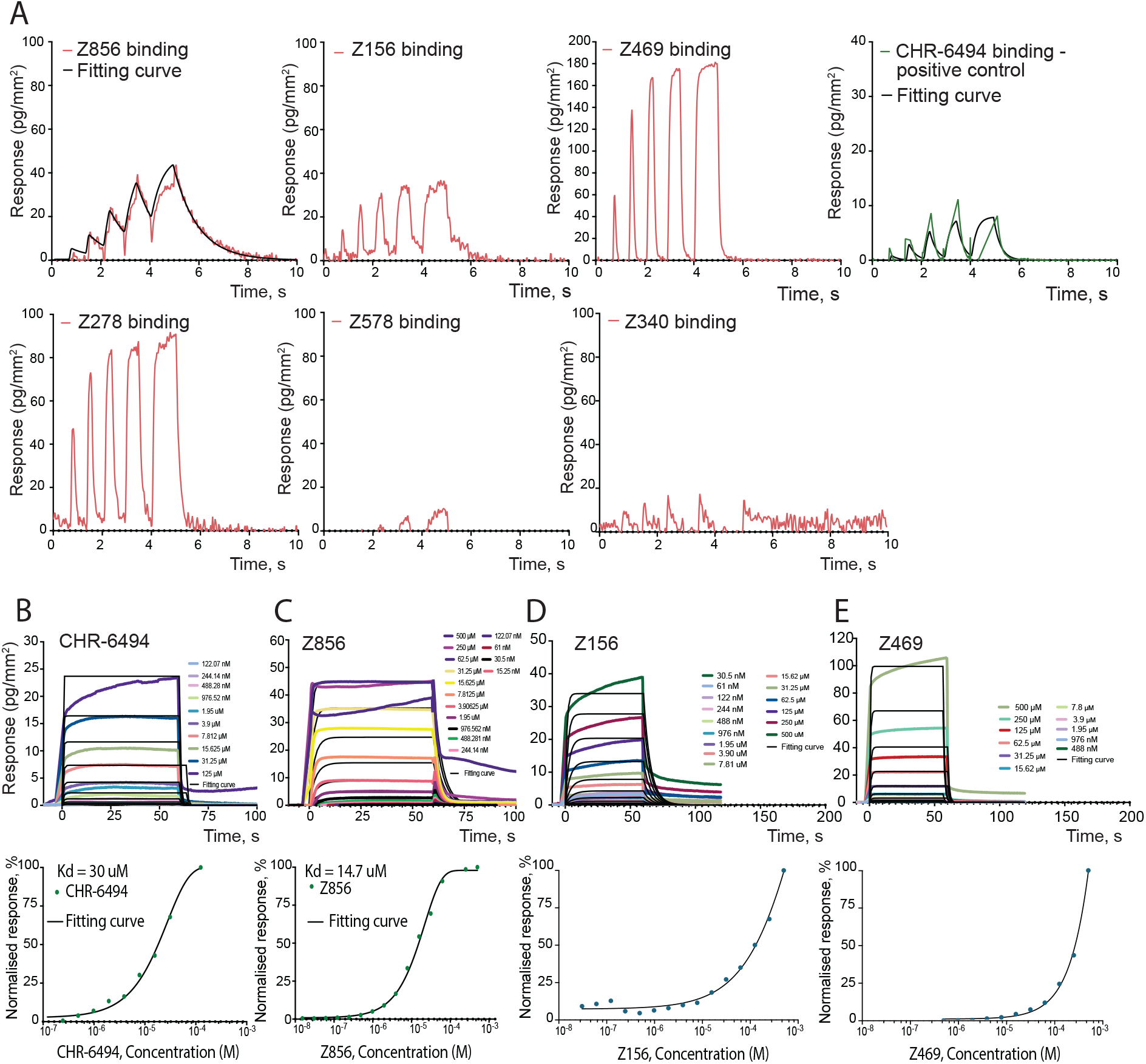
Experimental GCI Data for PGK1. **(A)** Rapid kinetics sensograms for all six tested compounds at 500 µM. **(B-E)** dose-response curves for K_D_ determination, and multicycle kinetic sensograms for varying compound concentrations for positive control CHR-6494 **(B)** and our designs Z856 **(C)**, Z156 **(D)** and Z469 **(E)**.

**Fig. A7:**
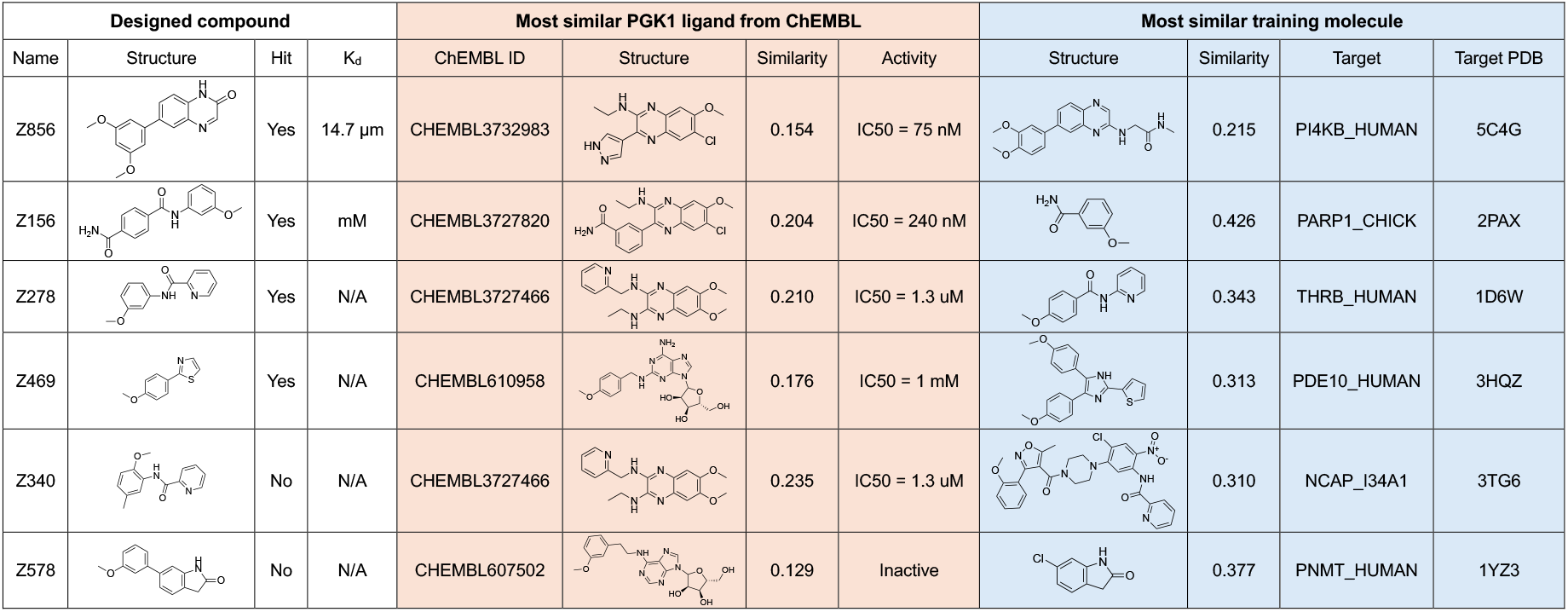
Similarities between PGK1 designs, known PGK1 ligands and training data. We report Tanimoto similarity between RDKit Morgan fingerprints with radius 3 and 2048 bits.

**Table A2:** Hit identification criteria for PGK1. We defined experimental hits based on two criteria: (1) Observed R_max_ *>* 6 pg*/*mm^2^ and (2) dissociation and association errors ≤ 100%. The theoretical R_max_ value was computed as 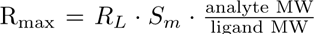 where *R_L_* is the amount of immobilized ligand in pg*/*mm^2^ and *S_m_*is the stoichiometry (number of binding sites for the analyte on the ligand). The relative diss. and ass. errors are calculated as the 95% confidence intervals of the dissociation rate (*k_d_*) and association rate (*k_a_*) estimates divided by their values.

| Compound | Theoretical $R_{\max}$,<br>$\text{pg/mm}^2$ | Observed $R_{\max}$,<br>$\text{pg/mm}^2$ | Diss. Error,<br>% | Ass. Error,<br>% | Hit? |
| --- | --- | --- | --- | --- | --- |
| CHR-6494 | 97 | 12 | 5.1 | 16 | yes |
| Z856 | 94 | 41 | 7 | 27 | yes |
| Z156 | 90 | 35 | 9.8 | 29 | yes |
| Z469 | 63 | 178 | 24 | 74 | yes |
| Z278 | 76 | 89 | 7.2 | 17 | yes |
| Z578 | 79 | 7 | $1.7 \times 10^8$ | $3.3 \times 10^3$ | no |
| Z340 | 80 | 0 | 99 | 73 | no |

**Fig. A8:**
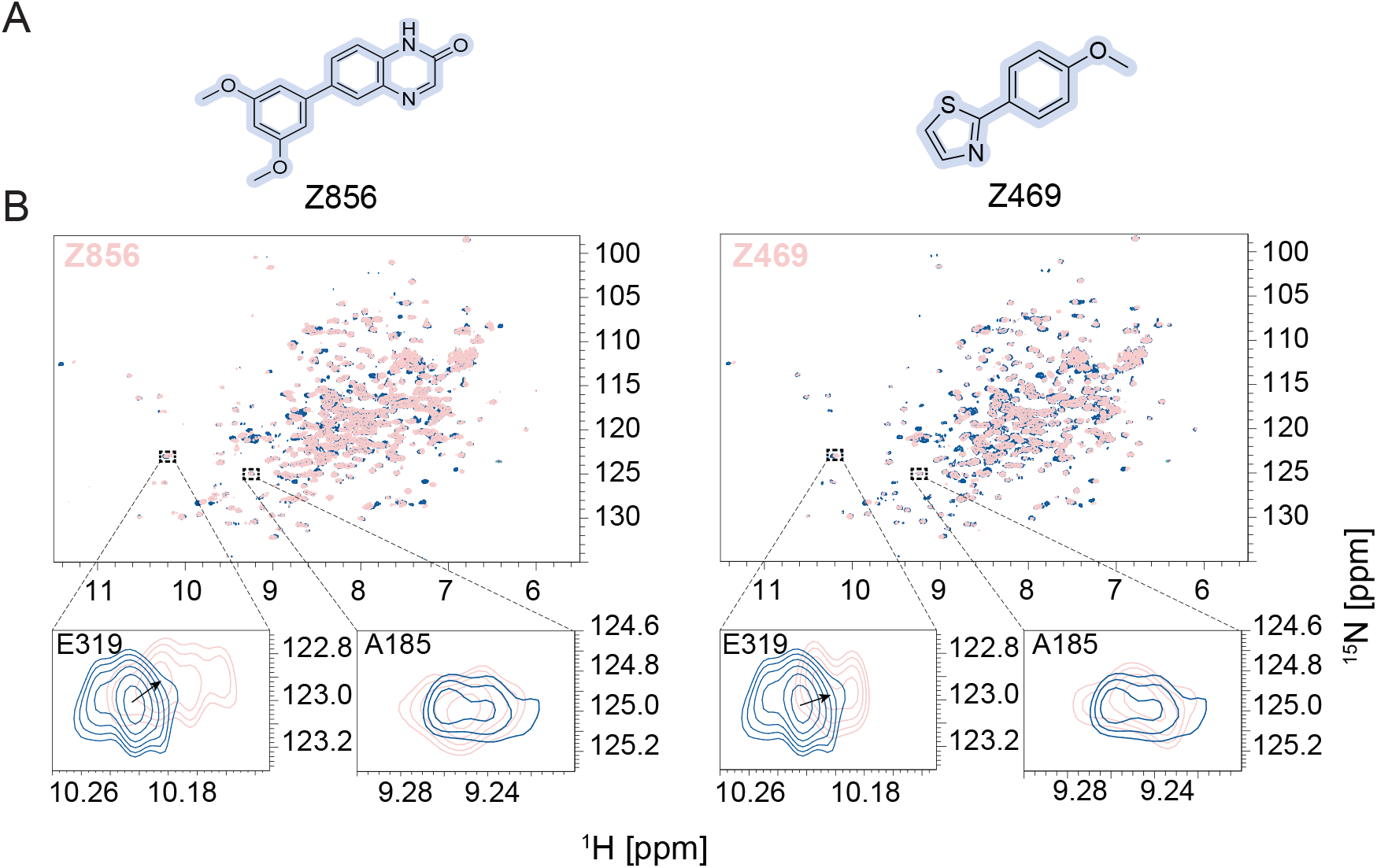
NMR spectroscopy data for PGK1 compounds Z856 and Z469. **(A)** Structures of Z856 and Z469 hit compounds. **(B)** [^15^N, ^1^H]-HSQC overlay of 100 µM apo-PGK1 G185A (dark blue) bound to 0.5 mM of the specified ligand (light red). Chemical shift of E319, with no shift observed for A185, indicates that Z856 and Z469 bind to the nucleotide binding domain of PGK1.

### A.7 Design of KRAS Ligands

Kirsten rat sarcoma virus GTPase (KRAS) is one of the key therapeutic targets in cancer research, whose hyperactivation due to mutations such as G12C leads to tumorigenesis [106]. Despite significant progress over the past decades in identifying chemical compounds targeting KRAS, the development of potent inhibitors remains challenging. This difficulty arises from several factors including the structural flexibility of KRAS, its rapid GTP hydrolysis, the emergence of resistance through secondary mutations, and others [107].

In our study, we focused on the Switch II pocket, the same site targeted by the pan-KRAS inhibitor BI-2865 [108]. For this pocket, we designed a series of compounds using LDDM, and shortlisted the most promising candidates using different computational filters (see Section B.5). The top five purchasable molecules (Figure A9A) were synthesized and screened for binding using GCI (Supplementary Table A3). Among three successful ligands, the most potent molecule, Z755, showed an affinity of 85.7 µM (Figure A9C). Details on the screening procedure and affinity measurement are provided in Methods Section 1.6. Figure A9B compares the modelled 3D structure of Z755 with the crystal structure of the positive control BI-2865 bound to KRAS (PDB ID: 5AZV). Although both compounds form hydrogen bonds with multiple residues in the Switch II pocket, their interaction profiles are largely distinct.

To confirm whether Z755 binds to the targeted site, we performed a blocking experiment, in which KRAS^G12C^ was preincubated with the covalent inhibitor Sotorasib [109]. This experiment showed reduced binding signal for Z755 in the presence of Sotorasib (Figure A9D), suggesting that Z755 binds to the targeted Switch II pocket.

**Fig. A9:**
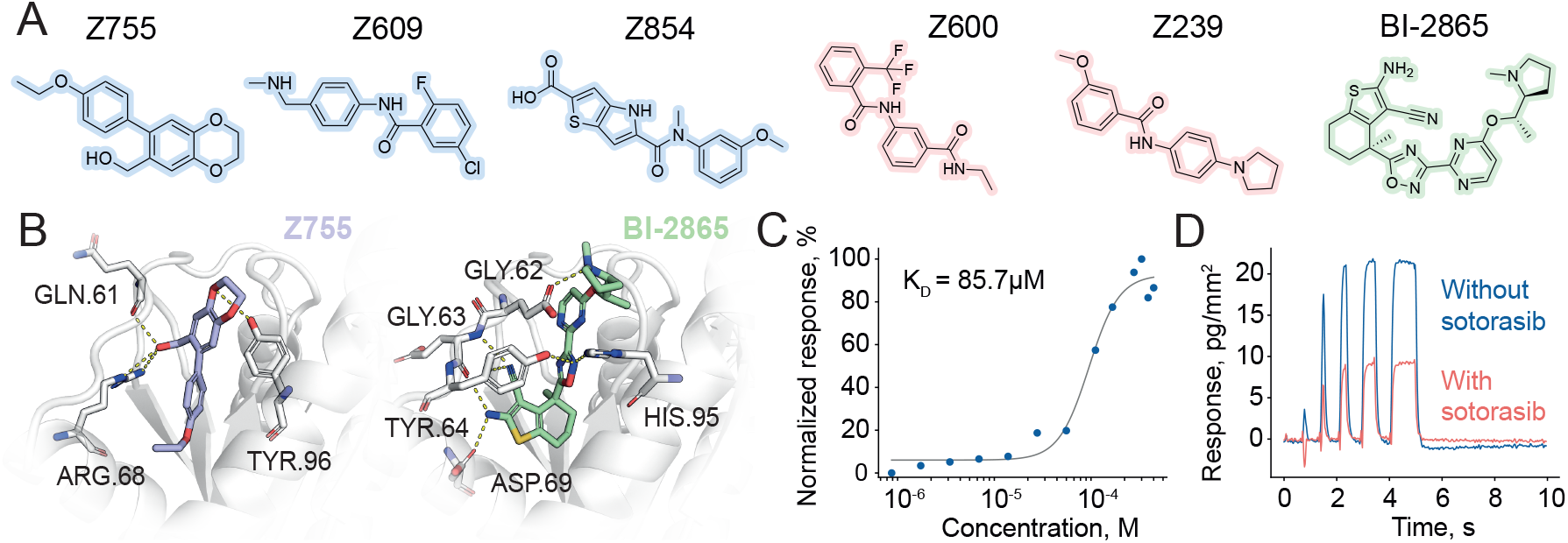
One-shot *de novo* design of KRAS ligands. **(A)** Chemical structures of BI-2865 and five designs tested experimentally against KRAS. Three identified hits are highlighted in blue, and two candidates with no observed binding are highlighted in red. **(B)** Predicted bound pose of Z755 and the crystal structure of BI-2865 in complex with KRAS (PDB ID: 8AZV). Amino acids involved in interactions are shown in sticks. Dashed yellow lines represent hydrogen bonds. **(C)** Grating coupled interferometry dose-response curve for Z755 binding to KRAS with the estimated affinity of 85.7 µM. **(D)** Rapid kinetics sensogram of Z755 binding to KRAS before (blue) and after (red) blocking the target pocket with covalent ligand Sotorasib.

**Fig. A10:**
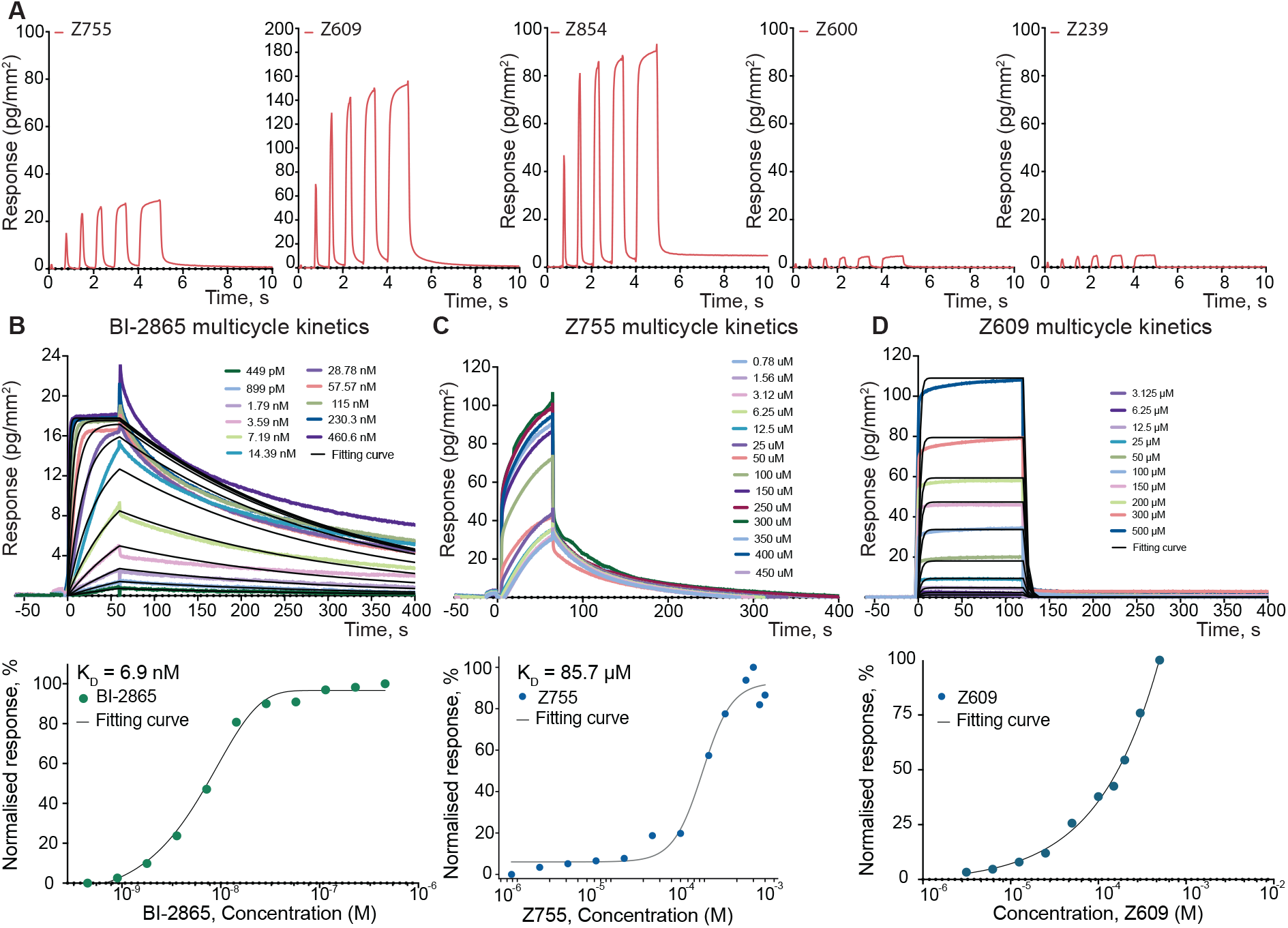
Experimental GCI Data for KRAS. **(A)** Rapid kinetics sensograms for all five tested compounds at 500 µM. **(B-D)** Multicycle kinetics sensograms for varying compound concentrations of the positive control BI-2865 **(B)** and designed compounds Z755 **(C)** and Z609 **(D)**.

**Fig. A11:**
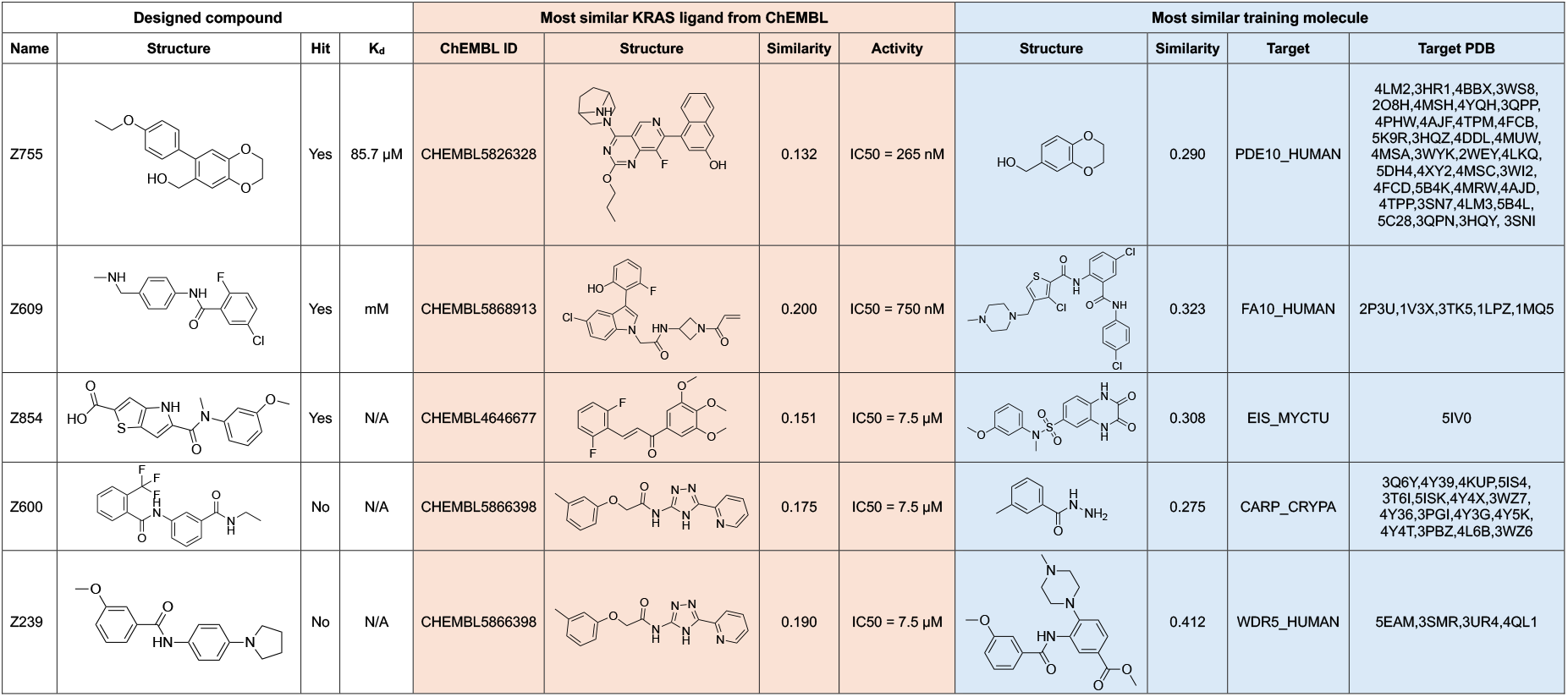
Similarities between KRAS designs, known KRAS ligands and training data. We report Tanimoto similarity between RDKit Morgan fingerprints with radius 3 and 2048 bits.

**Table A3:** Hit identification criteria for KRAS. We defined experimental hits based on two criteria: (1) Observed R_max_ *>* 15 pg*/*mm^2^ and (2) dissociation and association errors ≤ 100%. The theoretical R_max_ value was computed as 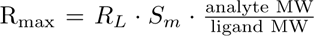 where *R_L_* is the amount of immobilized ligand in pg*/*mm^2^ and *S_m_* is the stoichiometry (number of binding sites for the analyte on the ligand). The relative diss. and ass. errors are calculated as the 95% confidence intervals of the dissociation rate (*k_d_*) and association rate (*k_a_*) estimates divided by their values.

| Compound | Theoretical $R_{\max}$,<br>pg/mm<sup>2</sup> | Observed $R_{\max}$,<br>pg/mm<sup>2</sup> | Diss. Error,<br>% | Ass. Error,<br>% | Hit? |
| --- | --- | --- | --- | --- | --- |
| Z755 | 205 | 29 | 9 | 50 | yes |
| Z609 | 209 | 153 | 9.2 | 32 | yes |
| Z854 | 236 | 90 | 53 | 37 | yes |
| Z600 | 212 | 4 | 120 | 120 | no |
| Z239 | 240 | 4 | $4.8 \times 10^8$ | 170 | no |

### A.8 Fragment-based Design of Covalent Ligands for Pin1

Pin1 is a cis–trans prolyl isomerase that plays a crucial role in tumorigenesis [110]. To design covalent Pin1 ligands, we started from a crystal structure of human Pin1 covalently bound to the validated inhibitor 164A10 (PDB ID: 8VJG) [111]. We removed the reference compound but kept the reacted chloro-acetamide fragment in place, as shown in Figure A12A. We then sampled 385 401 new ligand candidates with LDDM using 100 steps of the Heun sampler (see Supplementary Section B.3). Since the covalent bond between molecule and protein is not explicitly encoded, LDDM sometimes grows the fragment on both sides. In these cases, we only kept the substructures connected to the dedicated attachment atom (the nitrogen of the acetamide fragment). Furthermore, we created a final selection of 703 candidates by filtering out any designs that do not pass all PoseBusters filters, are smaller than 15 heavy atoms, contain ring systems that are found less than 100 times in ChEMBL, have more than 4 rotatable bonds or more than 3 unsatisfied hydrogen bond donors or acceptors, do not form at least one hydrogen bond or *π*-stacking interaction, or are duplicates of other designs. Among the remaining designs, we could find 25 molecules in Enamine’s REAL space and manually selected 5 candidates for experimental validation (Figure A12C).

First, we screened our designs using grating-coupled interferometry (GCI), which showed a binding signal for all five compounds (see Supplementary Figure A13). For compound Z154, we obtained a crystal structure of the complex (PDB code: 9SLI). As shown in Figure A12D, the designed molecule covalently engages Cys-113, albeit in a binding conformation different from the prediction. Upon analysis of asymmetric unit of the crystal, we observed that there were extensive contacts of the compound arising from crystal packing with neighbouring subunits. As shown in Figure A12E, our crystal was formed such that copies of Z154 were in close proximity to each other with aromatic rings lying in parallel planes and potentially forming *π*-*π* stacking interactions, potentially influencing the binding conformation of the ligand.

This experiment shows that LDDM’s general-purpose training strategy enables covalent ligand design out of the box. However, the available data does not allow us to verify the predicted binding mode under native conditions. As a result, further studies will be required to determine the extent to which the designed protein–ligand interactions drive covalent complex formation.

**Fig. A12:**
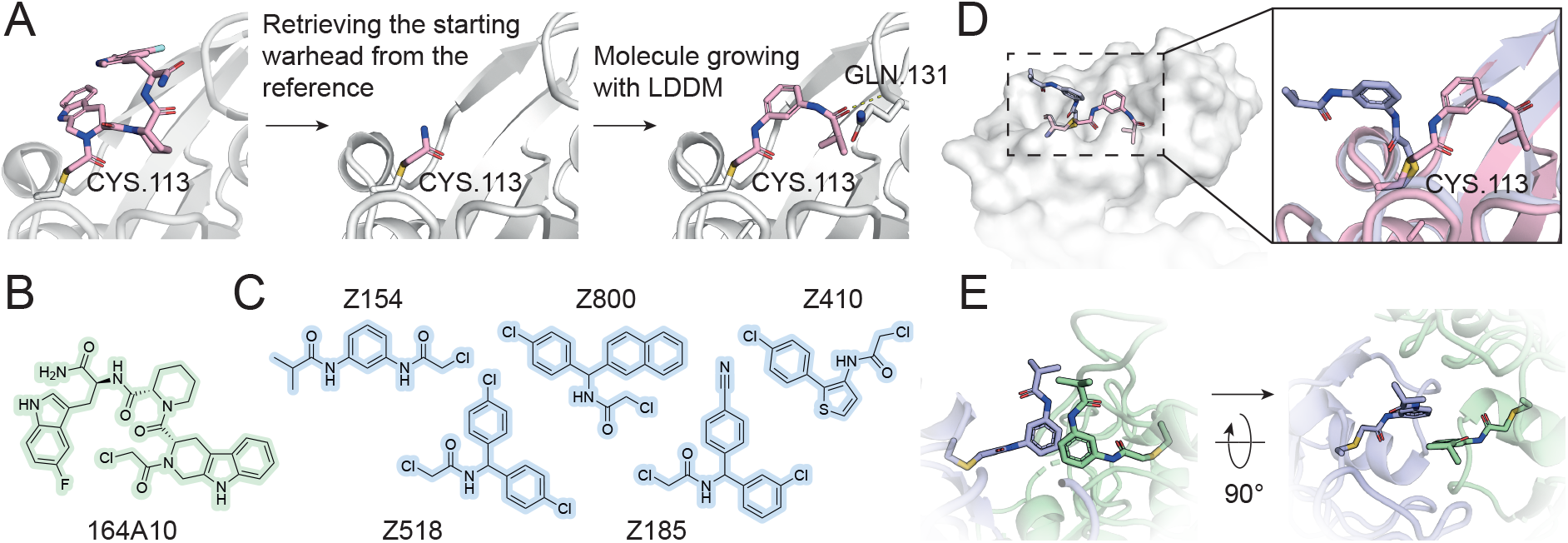
Fragment-based design of Pin1 ligands. **(A)** To design new covalent ligands for Pin1, we used LDDM to grow the starting chloro-acetamide fragment (PDB ID: 8VJG). **(B)** Chemical structure of the known covalent inhibitor 164A10. **(C)** Chemical structures of five LDDM designs. **(D)** Crystal structure of Z154 (blue) in complex with Pin1 overlaid with the modelled pose of Z154 (pink). **(E)** Contacts between Z154 compounds in the crystal.

**Fig. A13:**
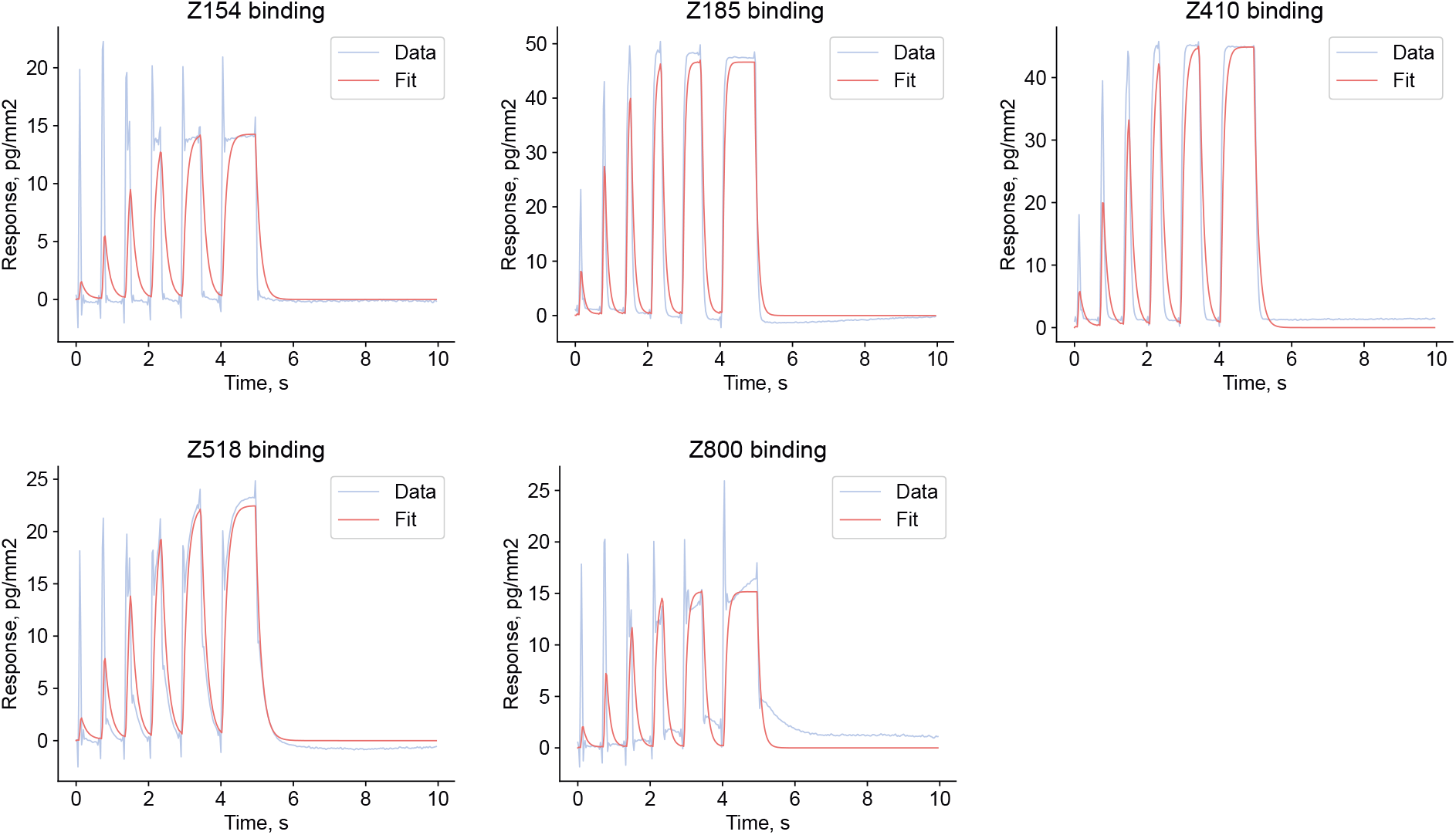
RAPID GCI kinetics sensograms for Pin1 designs, tested at 500 µM. We observed binding signal for all five tested compounds.

### A.9 Optimization of the Cathepsin S Peptide Inhibitor

**Fig. A14:**
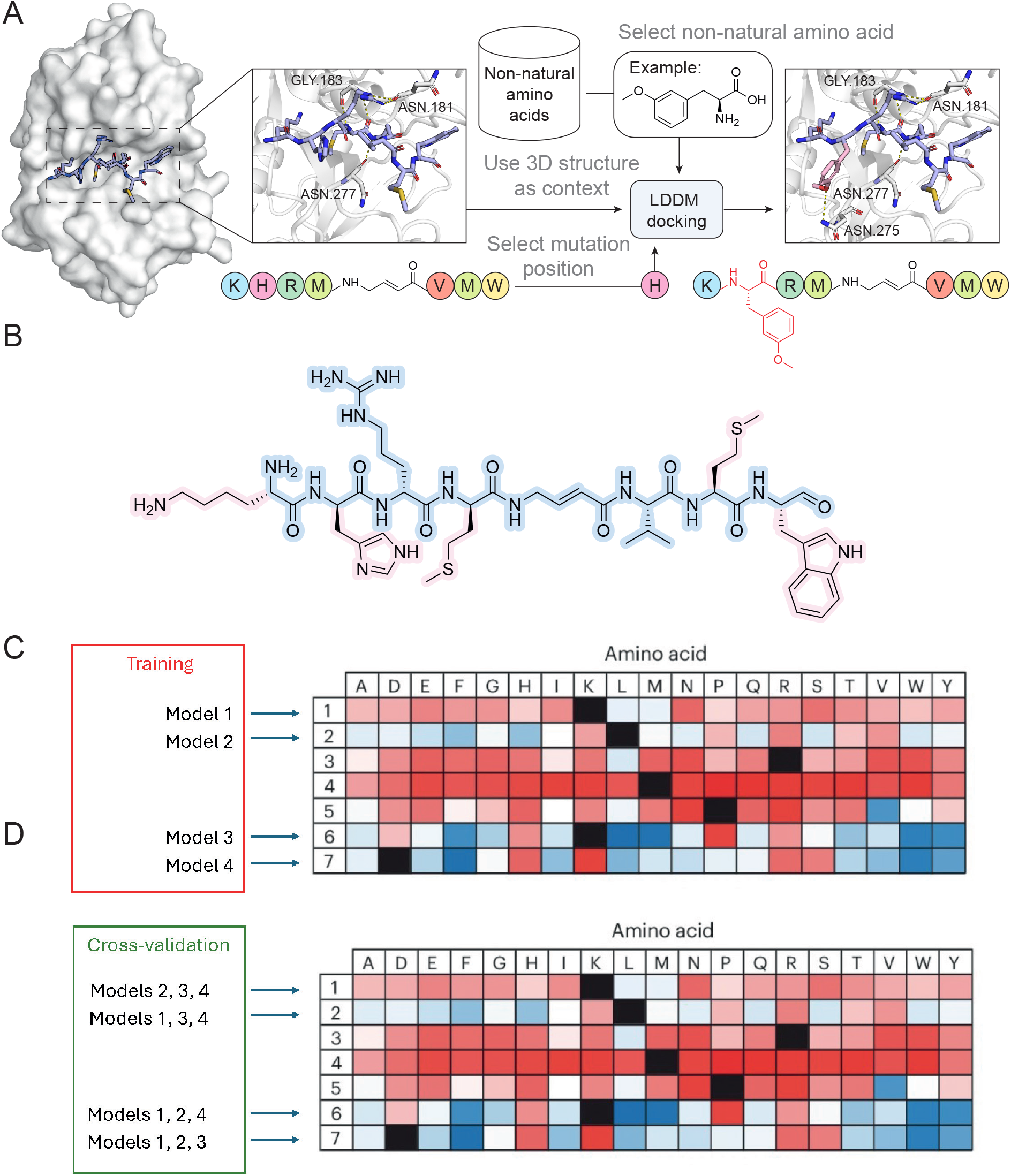
Optimisation and scoring of non-natural peptides. **(A)** Schematic representation of the process. Starting with the structure of NNPI-C10 bound to cathepsin S (on the left, PDB ID: 8PI3), we select a position for mutation and a non-natural amino acid from the database. We then replace the selected residue with the non-natural amino acid, and dock its side chain using LDDM. An example of the resulting structure with the a non-natural amino acid is shown on the right. **(B)** Chemical structure of the starting NNPI-C10. Five side chains subject to the computational mutagenesis are highlighted in pink. **(C)** We train four independent linear regression models, one per PSSM row. **(D)** To evaluate the performance of the trained models, we predicted the mutation effects for each sequence position using an ensemble of all models except the one corresponding to this position. PSSM matrix representation was taken from Petruzzella et al. [58].

**Fig. A15:**
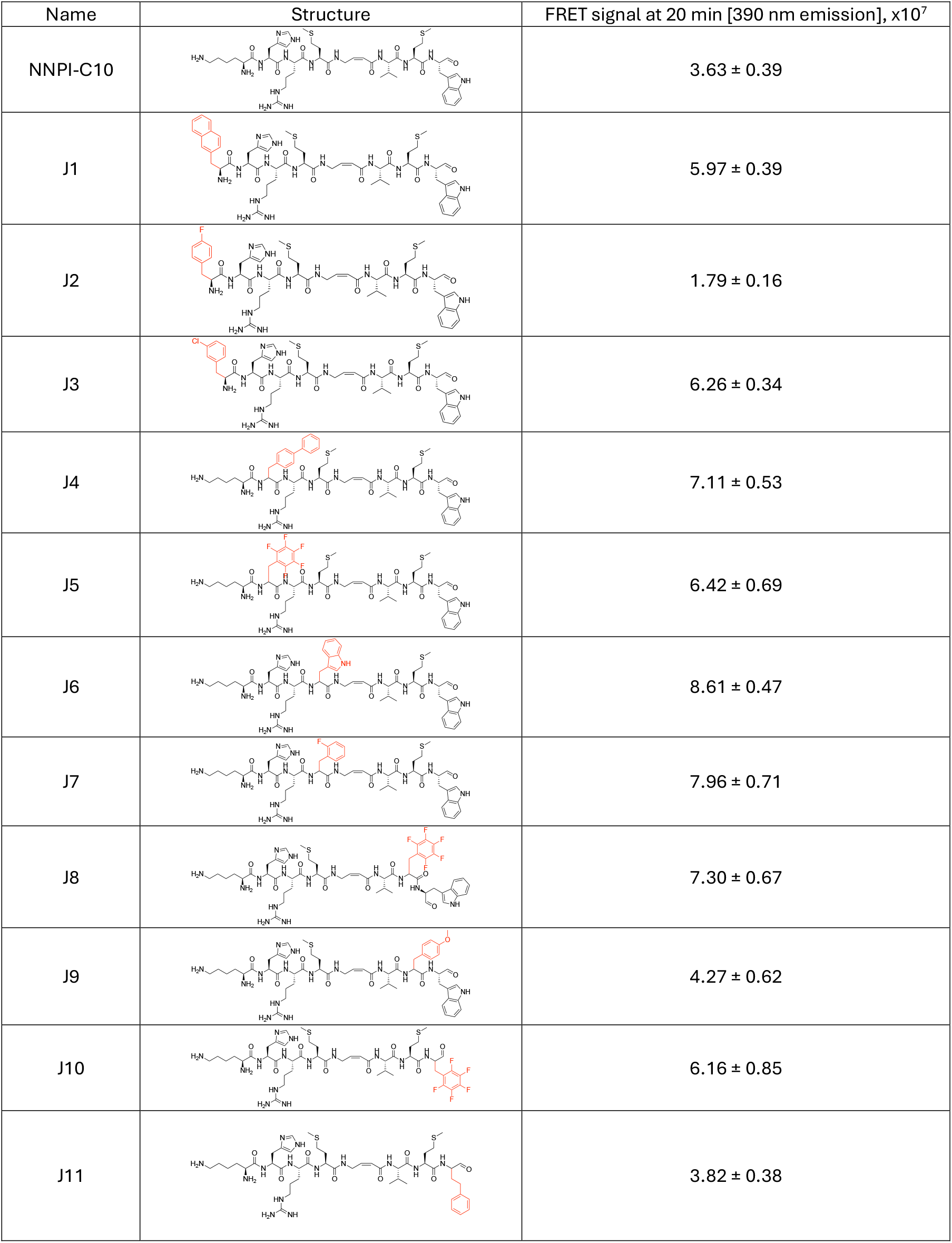
List of synthesized and experimentally tested peptides, together with their measured potency values. In each of the examples, the introduced amino acid is highlighted in red.

**Fig. A16:**
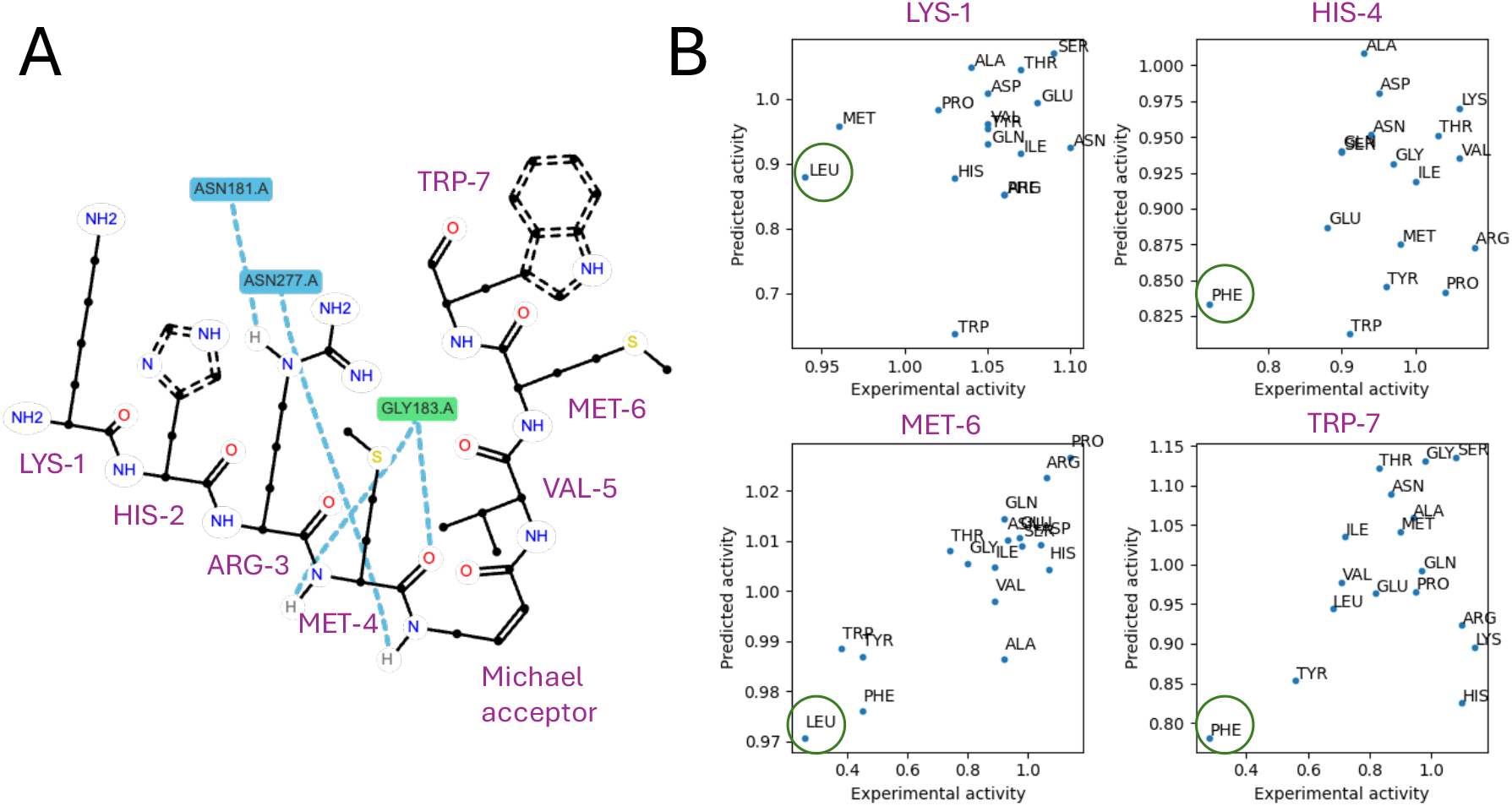
Cross-validation results for the NNPI scoring model. **(A)** Profile of interactions between NNPI-C10 and CTSS. **(B)** Our scoring approach demonstrates high recall, always ranking the best mutation (circled) among the top 5.

### A.10 Design of BRD4 Ligands

**Table A4:**
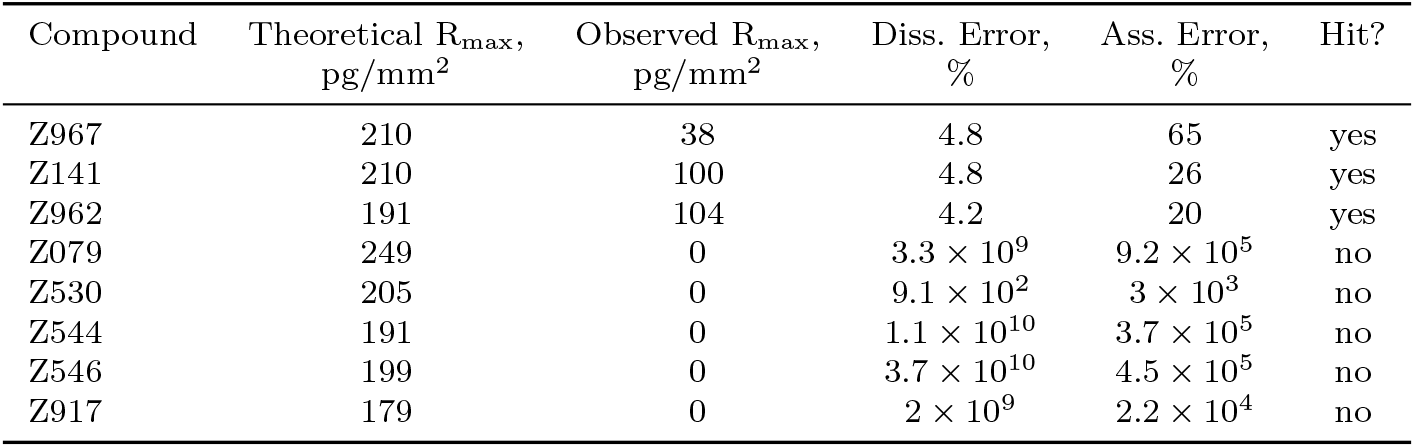
Hit identification criteria for BRD4. We defined experimental hits based on two criteria: (1) Observed R_max_ *>* 6 pg*/*mm^2^ and (2) dissociation and association errors ≤ 100%. The theoretical R_max_ value was computed as 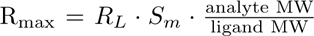 where *R_L_* is the amount of immobilized ligand in pg*/*mm^2^ and *S_m_*is the stoichiometry (number of binding sites for the analyte on the ligand). The relative diss. and ass. errors are calculated as the 95% confidence intervals of the dissociation rate (*k_d_*) and association rate (*k_a_*) estimates divided by their values.

**Fig. A17:**
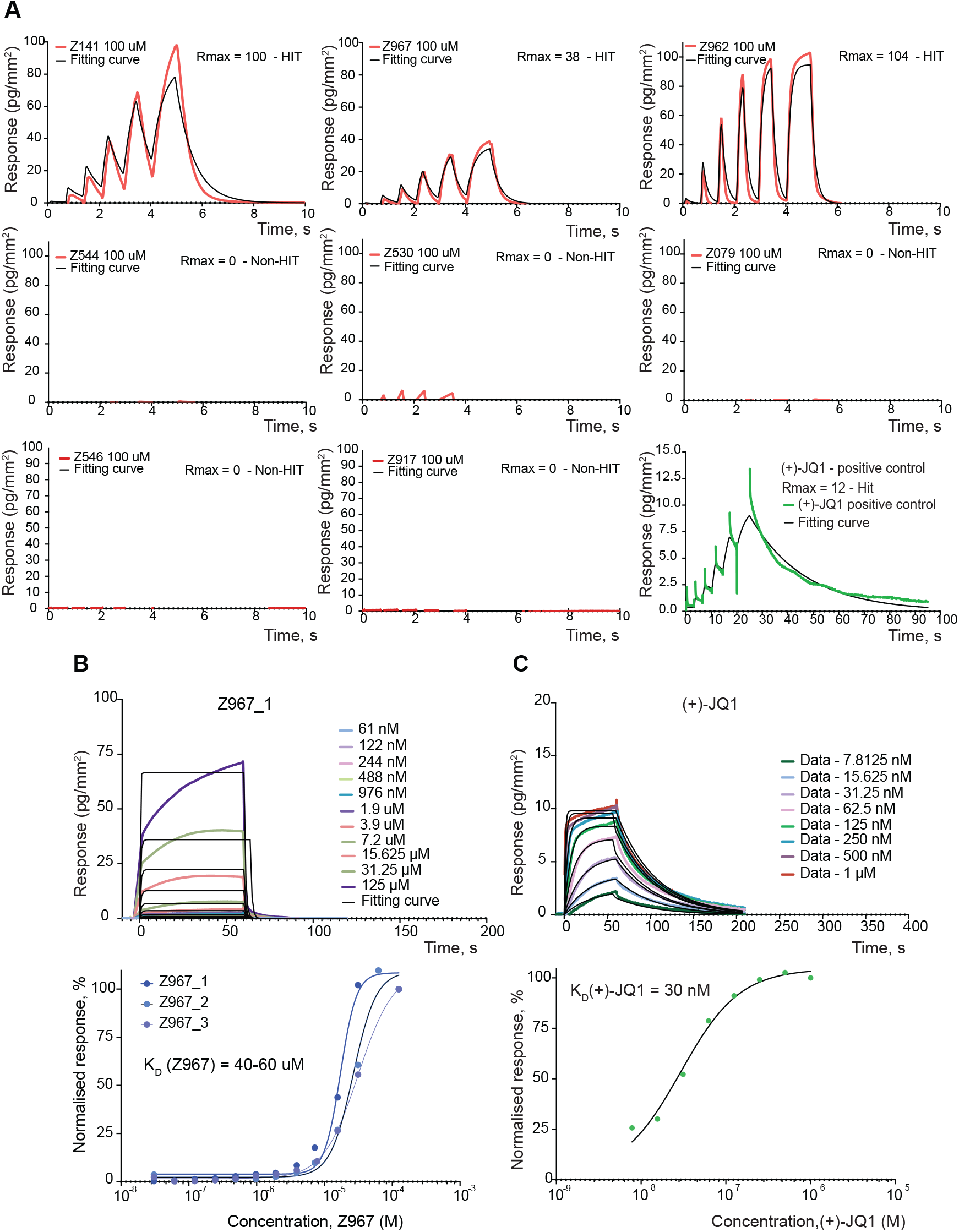
BRD4 GCI kinetic data. **(A)** Rapid kinetics sensograms for all eight BRD4 compounds screened at 100 µM and positive control (+)-JQ1. **(B)** Multicycle kinetic data for BRD4 and compound Z967 with affinity in the range of 40 µM to 60 µM. **(C)** Multicycle kinetic data for BRD4 and (+)-JQ1 positive control with affinity of 30 nM.

**Fig. A18:**
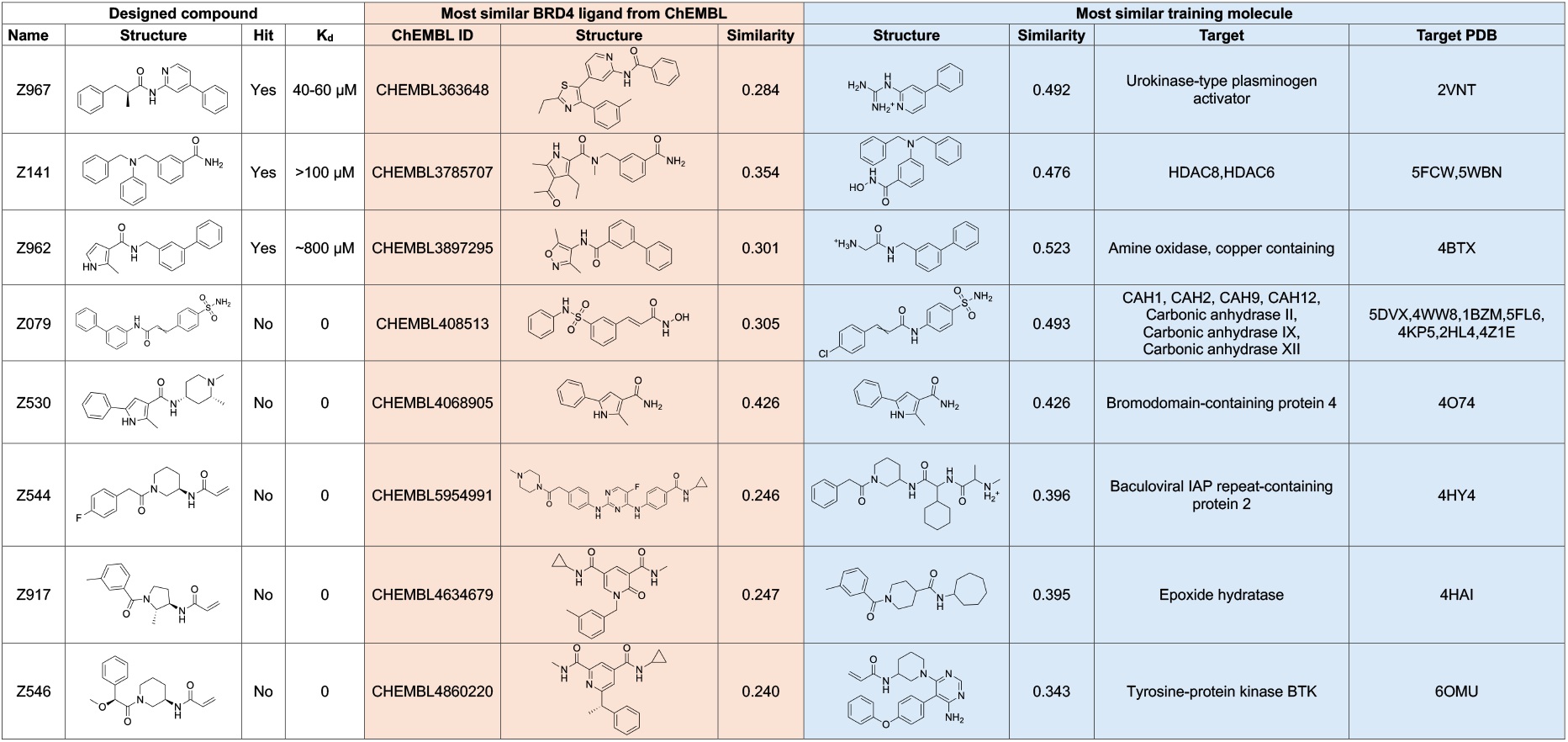
Similarities between BRD4 designs, known BRD4 ligands and training data. We report Tanimoto similarity between RDKit Morgan fingerprints with radius 3 and 2048 bits.

**Fig. A19:**
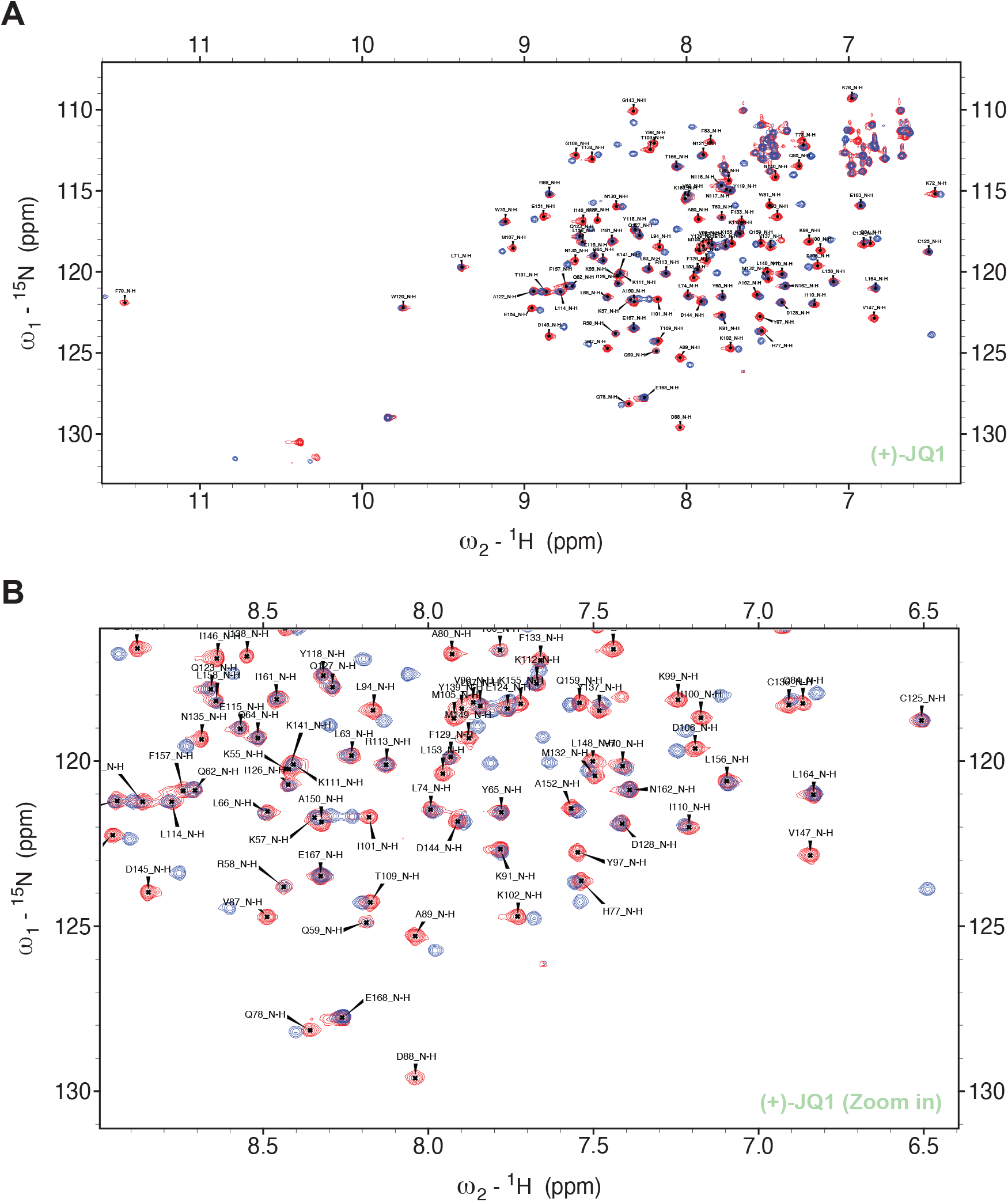
Overlaid [^15^N,^1^H]-HSQC spectra of apo-BRD4 with (+)-JQ1 compound. **(A)** Overlaid [^15^N,^1^H]-HSQC spectra of apo-BRD4 (dark red) with one equivalent of (+)-JQ1 (96 µM) (dark blue) indicates its binding to BRD4. **(B)** Zoom in overlaid [^15^N,^1^H]-HSQC spectra of apo-BRD4 (dark red) with one equivalent of (+)-JQ1 (96 µM) (dark blue) indicates its binding to BRD4.

**Fig. A20:**
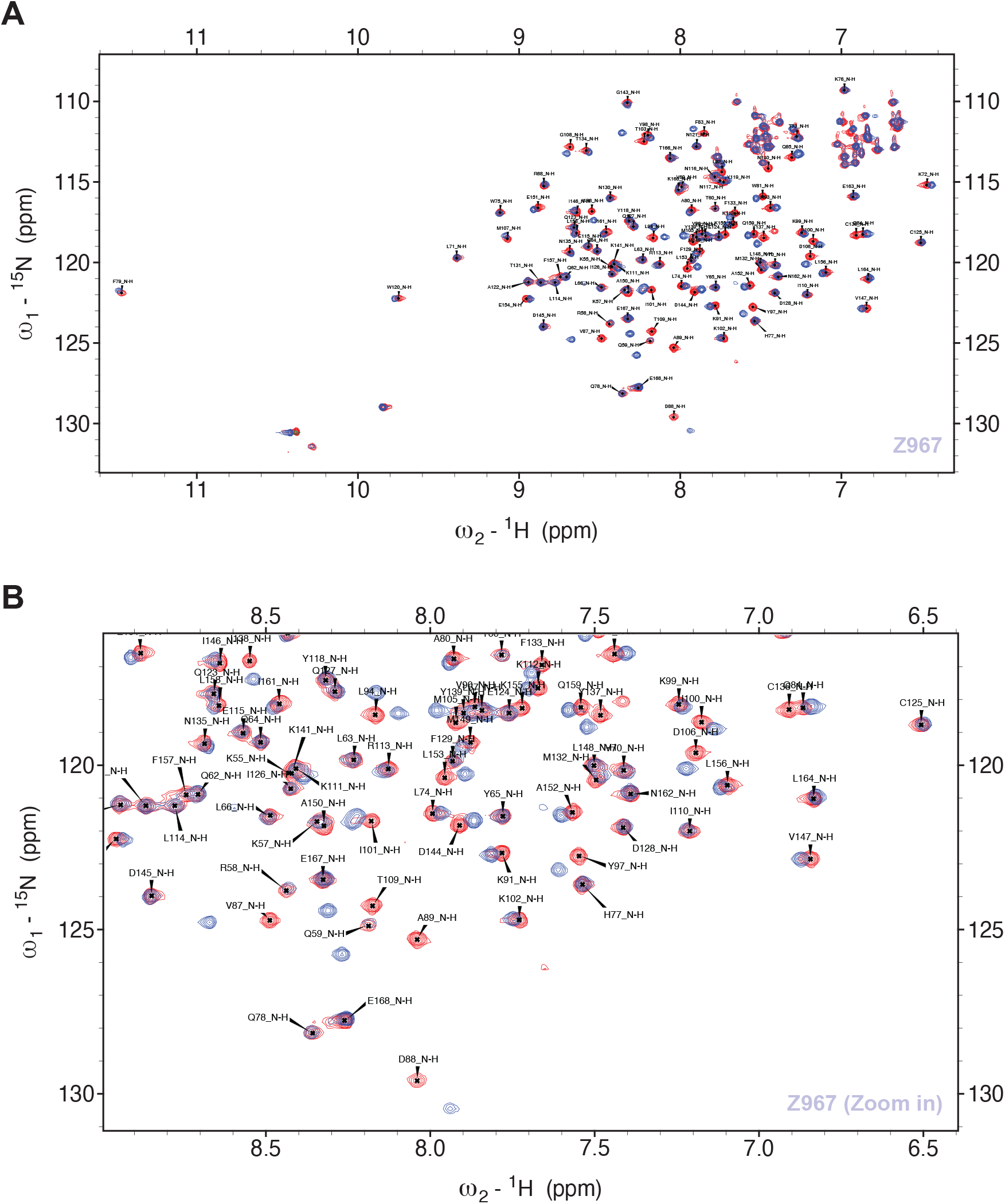
Overlaid [^15^N,^1^H]-HSQC spectra of apo-BRD4 with Z967 compound. **(A)** Overlaid [^15^N,^1^H]-HSQC spectra of apo-BRD4 (dark red) with one equivalent of Z967 (96 µM) (dark blue) indicates its binding to BRD4. **(B)** Zoom in overlaid [^15^N,^1^H]-HSQC spectra of apo-BRD4 (dark red) with one equivalent of Z967 (96 µM) (dark blue) indicates its binding to BRD4.

**Fig. A21:**
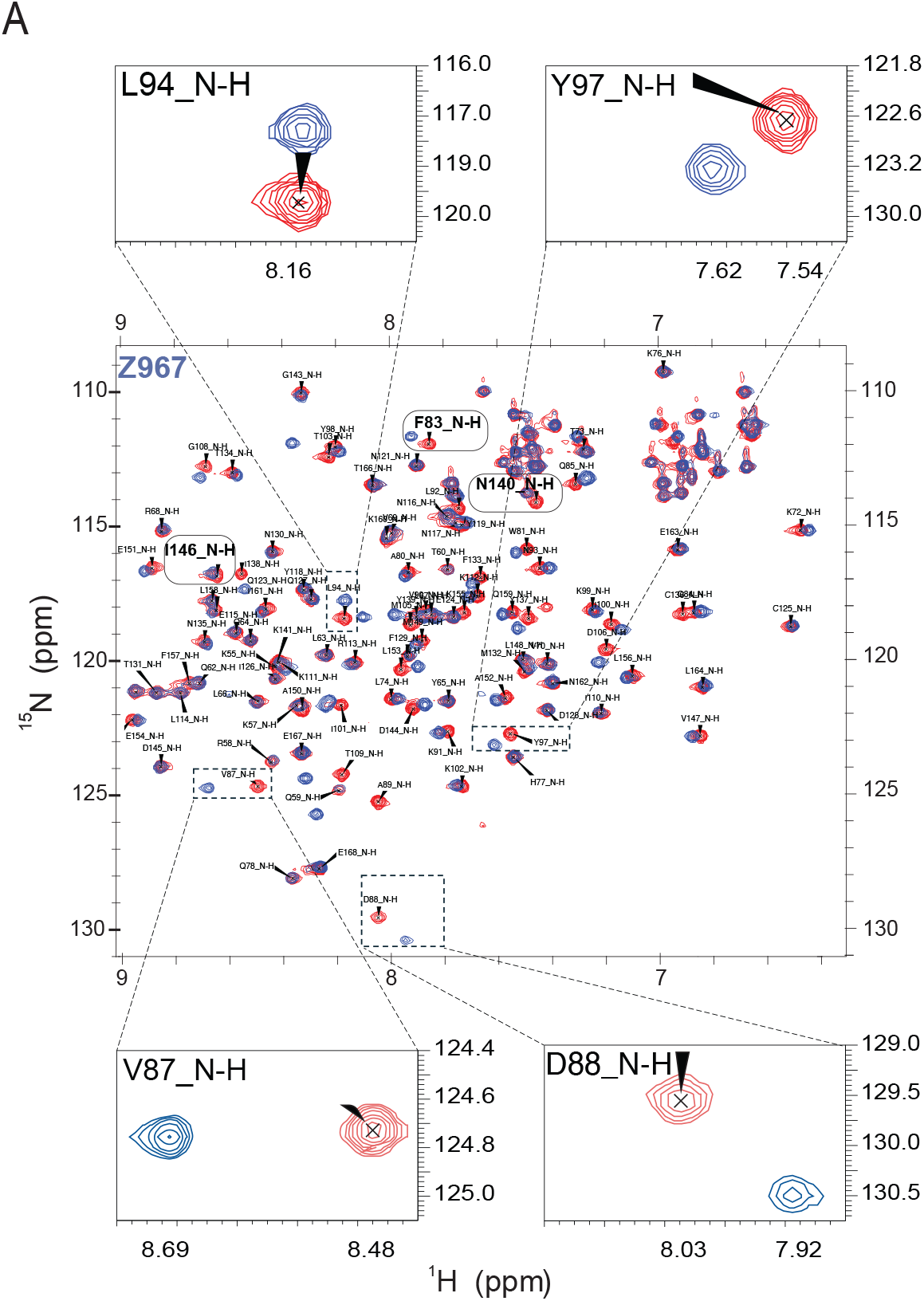
Overlaid [^15^N,^1^H]-HSQC spectra of apo-BRD4 with Z967 compound. **(A)** Overlaid [^15^N,^1^H]-HSQC spectra of of apo-BRD4 (dark red) with Z967 (96 µM) (dark blue) indicates its binding to BRD4. All corresponding shifts are marked and in agreement with the predicted pose.

**Fig. A22:**
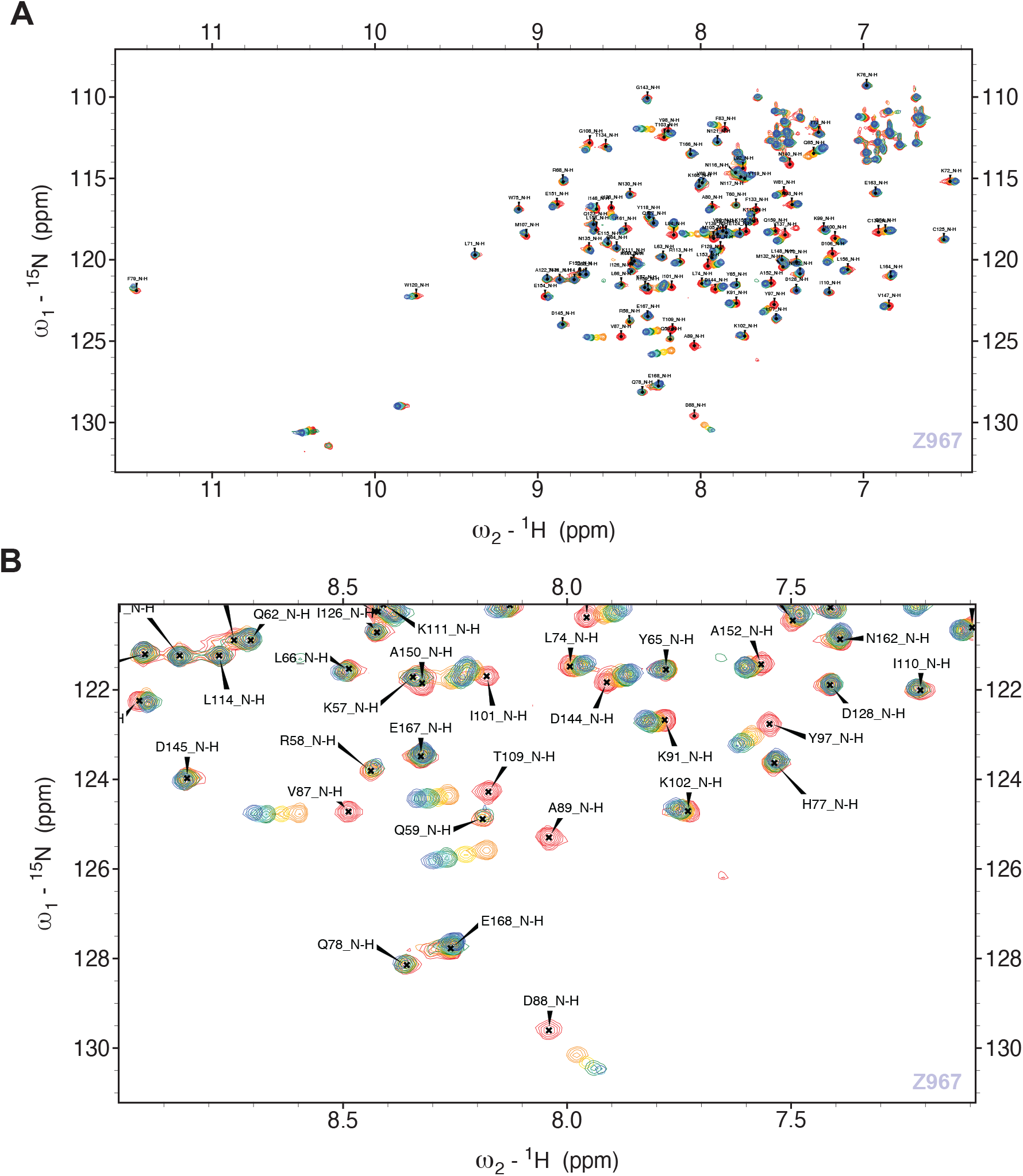
Overlaid [^15^N,^1^H]-HSQC spectra of titrations of apo-BRD4 with Z967 compound. **(A)** Overlaid [^15^N,^1^H]-HSQC spectra of titrations of apo-BRD4 (dark red) with Z967 (96 µM) indicates its binding to BRD4. Z967 compound concentrations for titrations (0, 0.25, 0.5, 1, 1.5, 2.5 equivalents) are shown in different colors from 0 in red to 2.5 in blue. **(B)** Zoom in overlaid [^15^N,^1^H]-HSQC spectra of titrations of apo-BRD4 (dark red) with Z967 (96 µM) indicates its binding to BRD4. Z967 compound concentrations for titrations (0, 0.25, 0.5, 1, 1.5, 2.5 equivalents) are shown in different colors from 0 in red to 2.5 in blue.

### A.11 Design of SARS-CoV-2 Nsp3 Inhibitors

**Fig. A23.**
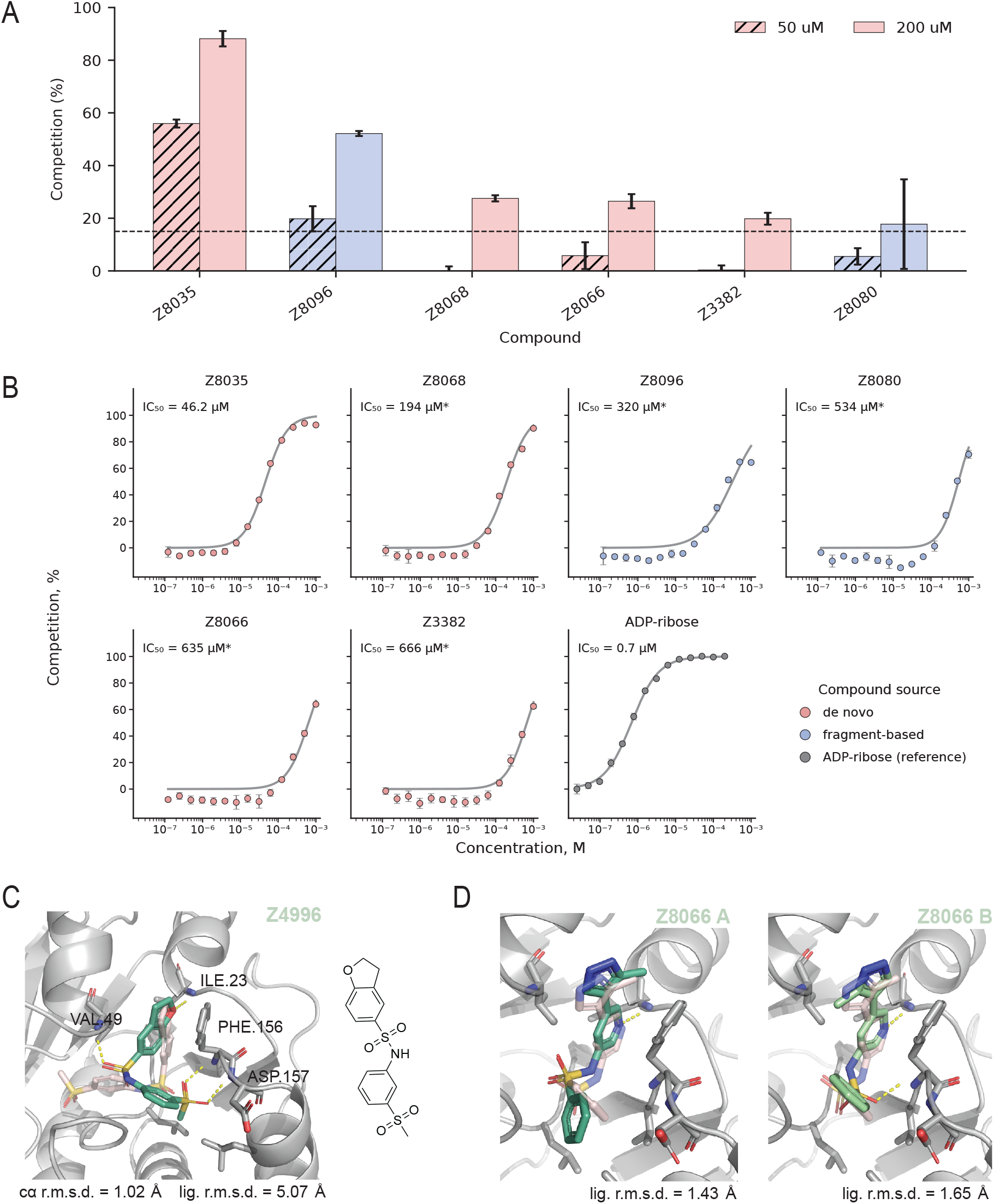
Experimental validation of computationally designed SARS-CoV-2 Nsp3 Mac1 ligands. **(A)** Activity of the six designs that exceeded the 15 % competition hit threshold in the primary screen. Bars show the mean HTRF competition (%) at 50 µM (hatched) and 200 µM (solid); error bars denote the standard deviation across triplicates. Bars are coloured by design origin (*de novo*, pink; fragment-based, blue). **(B)** Dose-response curves for the six hits and the ADP-ribose reference, measured over a 14-point dilution series in triplicates. Points are the mean competition (%) ± standard deviation; the grey line is a four-parameter Hill fit constrained to [0, 100] %, with the fitted IC_50_ indicated in each panel. Asterisks denote curves that did not reach a clear upper plateau within the tested concentration range, so the corresponding IC_50_ estimates should be interpreted with caution. The best *de novo* design (Z9941018035) inhibits with an IC_50_ of 46.2 µM, while ADP-ribose, the natural ligand, serves as a positive-control reference (IC_50_ 0.7 µM). The potency of our fragment-based compounds appears to be lower than the best fragment-based hits from the blinded competition, but the absolute numbers are highly dependent on the selected starting fragments and therefore difficult to compare. **(C)** Crystal structure of Z4996 bound to the SARS-CoV-2 Nsp3 Mac1 macrodomain (PDB ID: 38HY). Although Z4996 was inactive in the primary screen, a crystal structure of the complex was obtained with the ligand binding in the designated binding pocket. The designed complex is superimposed on the crystal structure (blue), with the designed ligand pose (red) differing substantially (ligand RMSD = 5.07 Å). **(D)** Two alternative crystallographic conformations of Z8066 (A and B) bound to Mac1 (PDB ID: 38HC), superimposed on the designed complex. The designed ligand pose is shown in light red; ligand RMSDs relative to conformations A and B are 1.43 Å and 1.65 Å, respectively.

**Fig. A24:**
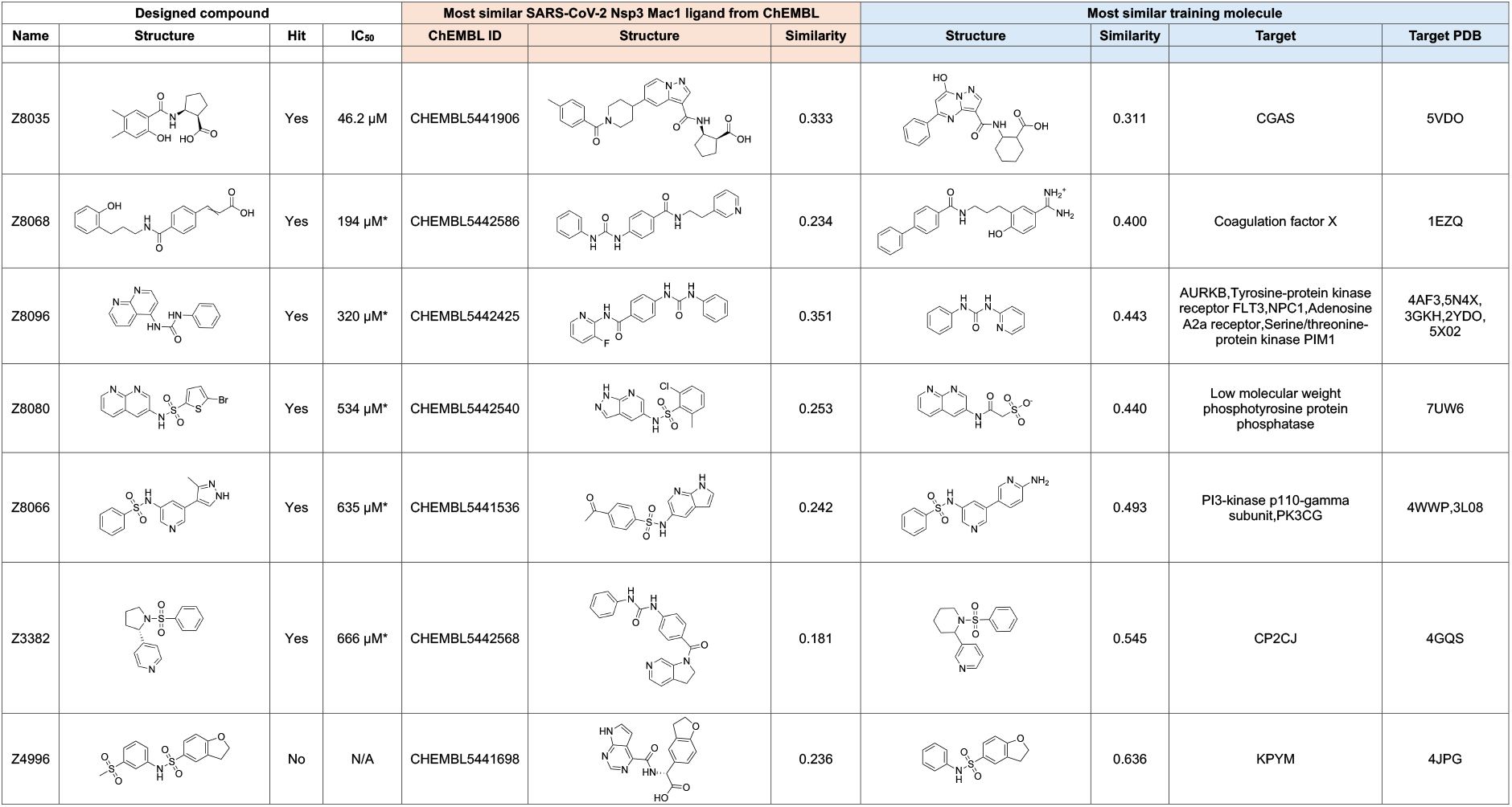
Similarities between SARS-CoV-2 Nsp3 Mac1 designs, known SARS-CoV-2 Nsp3 Mac1 ligands and training data. We report Tanimoto similarity between RDKit Morgan fingerprints with radius 3 and 2048 bits.

### A.12 Redesign of VHL-CDO1 Molecular Glues

**Fig. A25:**
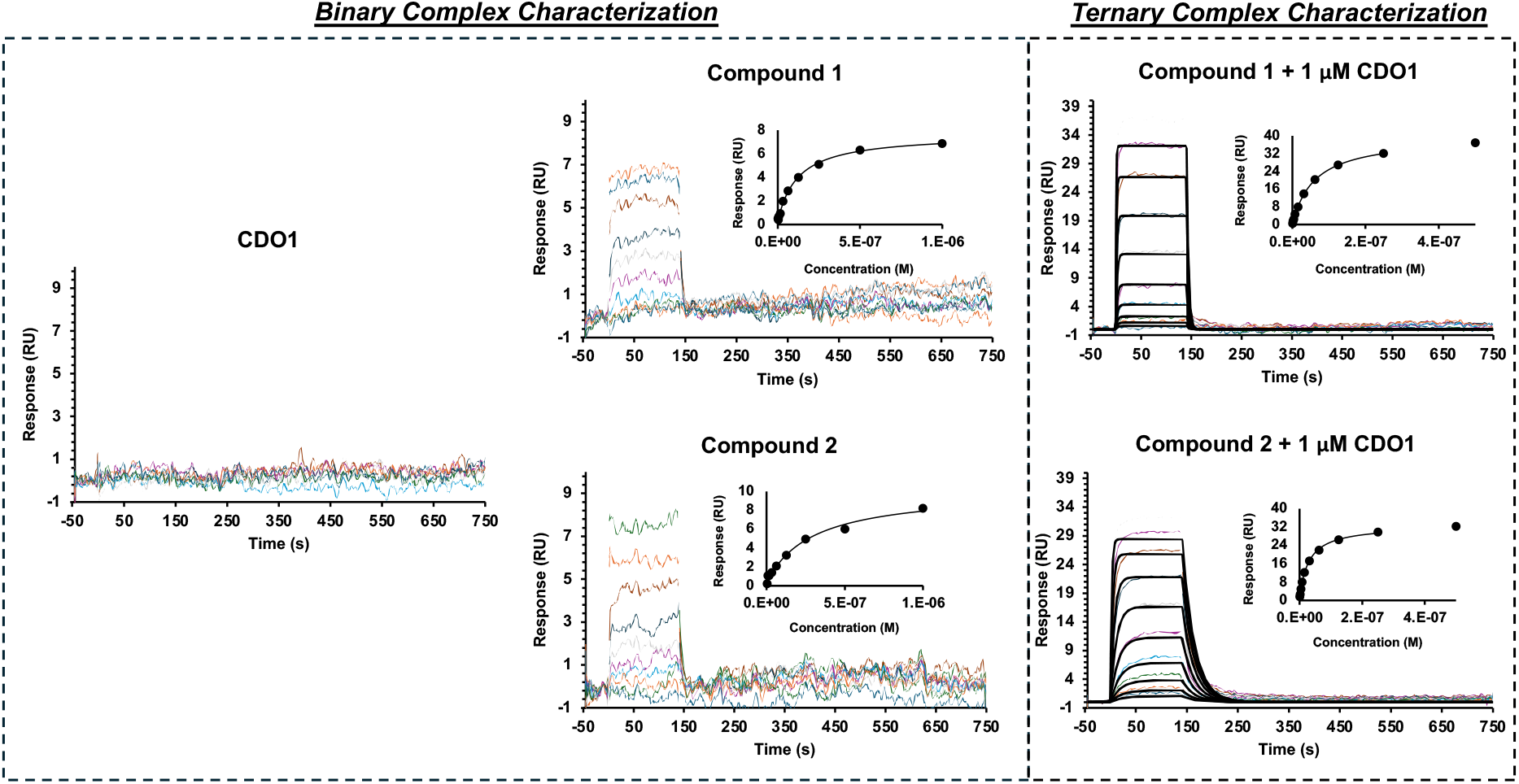
SPR characterisation of binary and ternary interactions of compounds 1, 2, and CDO1 on VCB-immobilized surfaces. Representative sensorgrams showing the interaction of compounds 1, 2, and CDO1 with VCB-immobilized surfaces under binary and ternary conditions. Left panel: binary interaction analysis of compounds 1 and 2 (tested up to 1 µM) and CDO1 (tested up to 2 µM) against VCB surfaces. CDO1 showed no detectable interaction with VCB within the tested concentration range. Compound 1 and 2 displayed transient binding behaviour, and steady-state profiles, recorded at the end of the association phase, are shown in the corresponding insets. Right panel: ternary interaction analysis of compounds 1 and 2 tested against VCB surfaces in the presence of 1 µM CDO1. Under ternary conditions, both compounds exhibited resolved binding kinetics, increased Rmax values, and enhanced affinity relative to the corresponding binary interactions. Complete kinetic and affinity parameters are reported in Table A5 for all tested samples. Colored curves represent recorded data and are overlaid with a 1:1 Langmuir binding model kinetic fit (black). The top concentration and following injections are related by a 2-fold dilution. All analytes were analyzed in triplicate, and one representative sensorgram is shown for each condition.

**Fig. A26.**
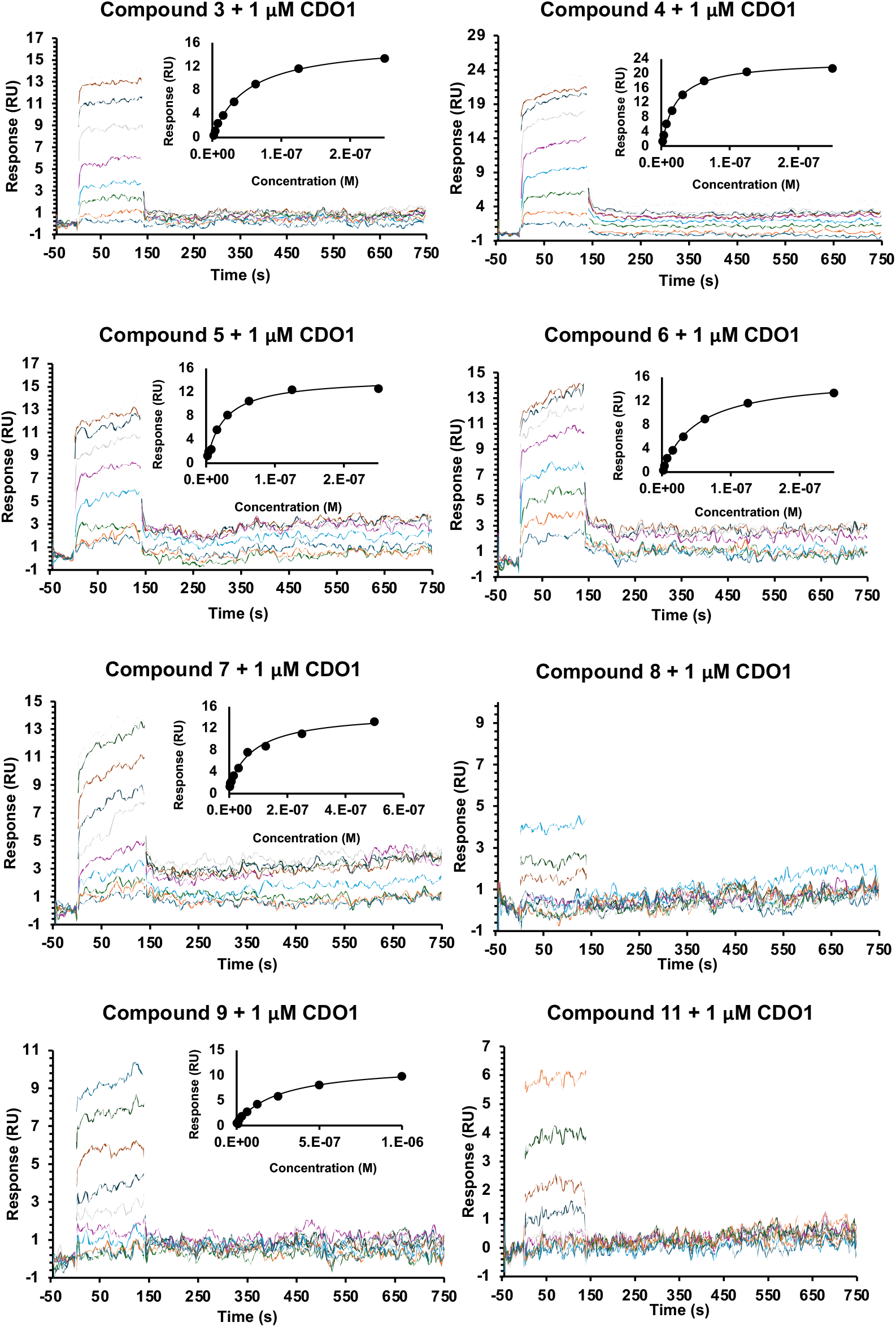
SPR characterisation of ternary interactions of compounds 3-11 on VCB-immobilized surfaces in the presence of CDO1. Representative sensorgrams showing the interaction of compounds 3-11 with VCB-immobilized surfaces under ternary conditions (1 µM CDO1). All compounds were initially tested up to 1 µM. Compounds 8, 10, and 11 were further evaluated up to 20 µM due to weak or undetectable binding, and for these compounds the apparent affinity exceeded 20 µM and could not be reliably determined. Compound 10 showed no detectable interaction under the tested conditions and is therefore not shown. Under ternary conditions, all compounds displaying measurable binding exhibited transient binding behaviour. Steady-state response levels, recorded at the end of the association phase, are shown in the corresponding insets. Colored curves represent experimental sensorgrams. The highest concentration and subsequent injections correspond to a two-fold serial dilution series and complete affinity parameters are reported in Table A6. All compounds were analyzed in triplicate.

**Table A5:** SPR determined affinity (*K*_Dss_) and kinetic (*k*_a_, *k*_d_, *K*_D_) parameters of compounds 1, 2, and CDO1 on VCB-immobilized surfaces in the absence (binary) or presence (ternary) of CDO1 (*n* = 3).

| | $K_{Dss}$<br>(M $\times 10^{-8}$) | $K_D$<br>(M $\times 10^{-8}$) | $k_a$<br>(M$^{-1}$ s$^{-1}$ $\times 10^6$) | $k_d$<br>(s$^{-1}$ $\times 10^{-2}$) |
| --- | --- | --- | --- | --- |
| CDO1 | ND | ND | ND | ND |
| Comp. **1** | 15 ($\pm 2$) | ND | ND | ND |
| Comp. **2** | 28 ($\pm 1$) | ND | ND | ND |
| Comp. **1** + 1 μM CDO1 | 6.4 ($\pm 0.2$) | 6.5 ($\pm 0.7$) | 3.8 ($\pm 0.6$) | 25 ($\pm 4$) |
| Comp. **2** + 1 μM CDO1 | 3.0 ($\pm 0.6$) | 2.6 ($\pm 0.4$) | 1.6 ($\pm 0.1$) | 4.1 ($\pm 0.2$) |

**Table A6:** SPR determined binding affinity (*K*_Dss_) of compounds 3–11 on VCB-immobilized surfaces in presence of 1 µM CDO1 (*n* = 3).

| | $K_{Dss}$ (M $\times 10^{-8}$) |
| --- | --- |
| Compound **3** + 1 μM CDO1 | 4.6 ($\pm 0.3$) |
| Compound **4** + 1 μM CDO1 | 2.1 ($\pm 0.2$) |
| Compound **5** + 1 μM CDO1 | 2.3 ($\pm 0.1$) |
| Compound **6** + 1 μM CDO1 | 1.5 ($\pm 0.1$) |
| Compound **7** + 1 μM CDO1 | 8.0 ($\pm 0.7$) |
| Compound **8** + 1 μM CDO1 | > 2000 |
| Compound **9** + 1 μM CDO1 | 25 ($\pm 1$) |
| Compound **10** + 1 μM CDO1 | ND |
| Compound **11** + 1 μM CDO1 | > 2000 |

## Appendix B Extended Methods

### B.1 Algorithmic Details for Programmable Generation

Programmable generation starts from an empty pocket and proceeds iteratively. In each round, candidate molecules are sampled from LDDM, conditioned on the nodes of a fragment tree *T* (unconditionally at the empty root). Candidates are filtered globally, fragmented using BRICS [31], and filtered locally; accepted fragments become new nodes and thus the conditioning scaffolds of the next round. The procedure stops after *N*_max_ iterations or once *A*_max_ molecules have been accepted. We write *M* for the space of molecules and *A* ⊆ *M* for the set of accepted molecules returned by the procedure. All molecules are represented with atom coordinates inside a fixed protein pocket, which we leave implicit throughout. The procedure has the following components:

- a *global filter* FilterMol: *M* → {0, 1}, which discards invalid or undesired molecules;
- a *local filter* FilterFrag: *M* → {0, 1}, which accepts a fragment under local constraints evaluated in the pocket (e.g., protein–ligand interaction profiles);
- a *fragment tree T* whose nodes *v* store a fragment *v.*mol, with *r.*mol = ∅ at the root *r*. Given a set of fragments *F* ⊆ *M*, the procedure AddChildren(*T, v, F*) attaches every *x* ∈ *F* to *T* as a child of *v*, discarding fragments that do not contain *v.*mol as a substructure, merging duplicates into existing children.

Algorithm 1 outlines the iterative procedure. In each iteration, the allocation function Alloc distributes a *sampling budget b* over the nodes of *T*. To balance exploration of under-sampled fragments against exploitation of promising ones, we adopt an Upper Confidence Bound (UCB) mechanism inspired by Monte Carlo tree search [74]. Each node *v* tracks calls(*v*), the number of molecules sampled at *v*, and successes(*v*), the number of those that passed FilterMol; we write pCalls(*v*) = calls(parent(*v*)), and pCalls(*v*) = calls(*v*) at the root. The UCB score for node *v*is then computed as

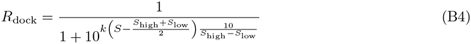

where *c* is a hyperparameter controlling the exploration–exploitation trade-off. In practice, we use *c* = 1 in our benchmarks. Alloc returns integer budgets {*b_v_*}*_v_*_∈*T*_ with Σ*_v_ b_v_* = *b*, proportional to UCB(*v*).

**Algorithm 1.**
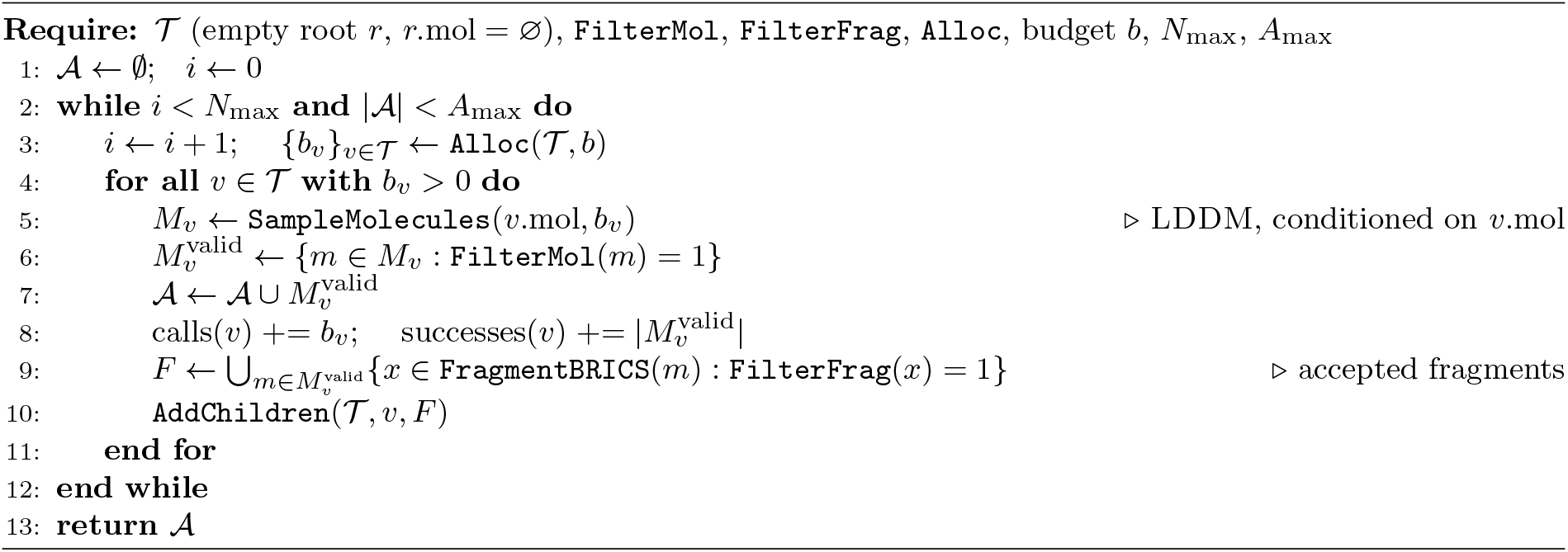
Programmable Generation with LDDM.

### B.2 Algorithmic Details for Synthesizable Generation

Synthesizable generation extends Algorithm 1 by fragmenting molecules along substructures that are reachable in a synthetically accessible product space. Every node *v* additionally stores a product space *P_v_* and a synthon *v.*syn; the root holds the available building blocks, *P_r_* = *B*, and an empty synthon. Sampling at *v* is conditioned on *v.*syn rather than on *v.*mol, so that the atoms consumed by a reaction remain free; the global and local filters act as before. Here successes(*v*) counts the accepted synthesizable fragments obtained at *v* rather than the molecules passing FilterMol. The procedure stops after *N*_max_ iterations or once *T* holds *T*_max_ nodes. MatchProducts(*m, P_v_*) returns the substructures of *m* that match a product in *P_v_*, and the optional DockSimilarProducts(*m, P_v_*) returns products *p* ∈ *P_v_* similar to *m*, partially redocked by preserving the coordinates of the MCS of *m* and *p* and redocking only the unmatched atoms; both are candidate children of *v*. Candidates that pass FilterFrag are attached by AddChildren, which as in Algorithm 1 retains only those containing *v.*mol, so that every root-to-node path is a sequence of reactions applied to *B*. For each newly created node *w*, ComputeReactions(*w, R, B*) enumerates the products obtainable by applying the templates *R* to *w.*mol and *B*, and ComputeSynthons(*w.*mol*, P_w_*) extracts the maximum common substructure (MCS) of *w.*mol and the products in *P_w_*. ComputeReactions stores with each product the template and building block that produced it; the root path of a node therefore yields a complete synthesis route. Candidate building blocks and products are shortlisted by fingerprint similarity before any MCS computation. Algorithm 2 details the procedure.

**Algorithm 2.**
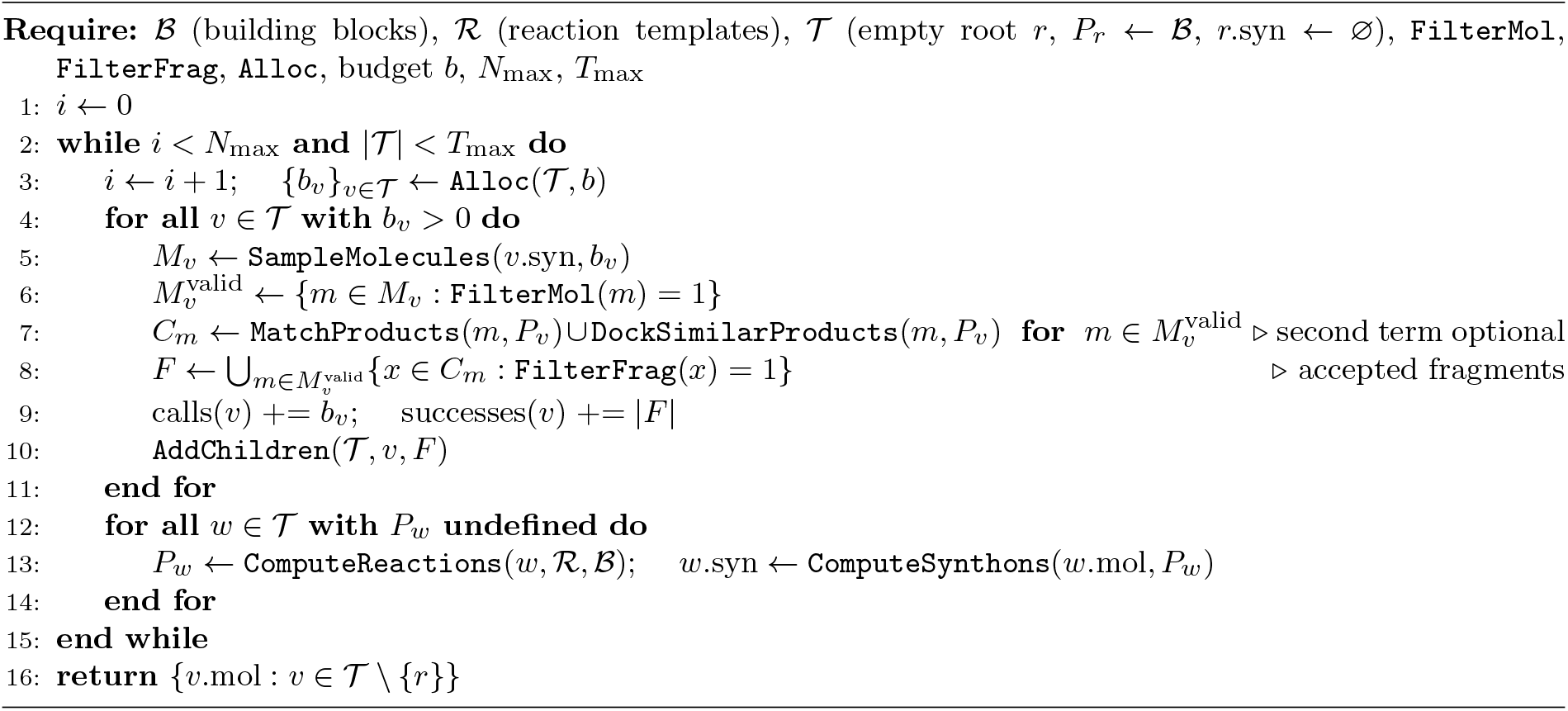
Synthesizable Generation.

### B.3 Heun Sampler

To reduce discretization error and enable sampling in fewer steps, we sometimes employ a second-order numerical integration scheme based on Heun’s method. This method achieves a better numerical approximation of the next state at the cost of an additional neural network call in every step. For continuous variables *x_t_* (i.e. atom coordinates), their value *x_t_*_+Δ*t*_ at the next integration point is computed as

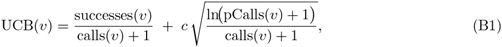

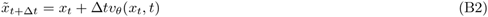

using the predicted vector field *v_θ_*(·, ·).

Analogously, for the categorical atom and bond type variables, we average the logits predicted in the current step and the intermediate step to perform an update in the Markov bridge framework.

### B.4 Recovery Experiment

#### B.4.1 REINVENT

To assess REINVENT’s ability to recover reference ligands, we optimize for the Vina score by docking using Gnina [51] with the binding site specified by the reference ligand (autobox parameter). We transform the raw docking scores using a reverse sigmoid function

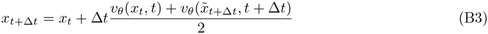

where *S* is the docking score, *S*_low_ and *S*_high_ define the range of scores set to −14 and −2 respectively, and *k* a factor defining smoothness set to 0.25. Other settings are identical to the optimization experiment (see Methods Section 1.3.5).

#### B.4.2 Virtual screening (VS)

To simulate virtual screening, we dock molecules from ChEMBL [112] (release 33) into the test set proteins using a GPU-optimised version of QuickVina [103, 113], and select the top 100 hits according to docking score. For computational tractability, we randomly select a subset of 10 000 ChEMBL molecules as our screening library. For docking, we use target-specific bounding boxes that fully cover the associated reference ligand plus 5Å margin added to all six sides.

#### B.4.3 Random

As a trivial baseline, we include a random selection of docked molecules from the screening library (rather than the top performing ones).

#### B.4.4 Shape-colour similarity

We use a symmetrized version of the shape-colour similarity score proposed in [45]. Shape similarity measures the volumetric overlap of two molecules [101] and colour similarity compares pharmacophores in 3D [102]. Shape-colour similarity is the average of both scores, which outputs values between 0 and 1. We obtain a symmetric score by using both molecules as reference and averaging the outcomes.

### B.5 Design of KRAS ligands

#### B.5.1 Computational design

To design KRAS ligands, we used the crystal structure with PDB ID 8AZR and applied LDDM trained on the CrossDocked dataset in the one-shot design regime. During generation, we discarded molecules that did not pass PoseBusters validity filters [42], REOS filters [84], or had at least one ring system that was not found in ChEMBL [85, 86]. Eventually, we obtained 10 000 unique samples.

#### B.5.2 Filtering Criteria

To increase the likelihood of experimental success, a series of filtering steps were applied. To ensure the solubility of the test compounds, the logP values of the selected molecules were restricted to a range of 1 to 3. Additionally, to reduce molecular flexibility and limit binding degrees of freedom, the number of rotatable bonds was capped at a maximum of 5. After applying these two filters, 2268 molecules remained. These molecules were then subjected to an exact-match search against the Enamine REAL database, which contains 48 billions of compounds. The search was performed using the SpaceLight [114] mode of the InfiniSee tool, based on Tanimoto similarity on fCSFP4 fingerprints. This process identified 207 molecules within the database. To validate the binding mode, molecular docking was conducted using Gnina [51] with an exhaustiveness parameter of 32. A total of 133 molecules were docked with a normalized RMSD of *<* 0.3 Å per atom. For all 133 molecules, an interaction profile was generated based on the predicted binding pose from LDDM, with a focus on recovering the known interaction fingerprint of small-molecule ligands targeting the same pocket. The top five molecules, ranked according to the number and type of recovered interactions, were selected for experimental testing.

#### B.5.3 KRAS^G12C^ protein expression and purification

KRAS gene sequences were cloned into a pet11 vector using NdeI and BlpI restriction sites by Gibson assembly, with an N-terminal 6xHis tag. DNA sequences were purchased from Twist Bioscience or obtained as cloned plasmids from Genscript. Plasmids were transformed into *Escherichia coli* BL21 (DE3) and sequenced. Expression was carried out in Terrific Broth supplemented with ampicillin (100 µg/mL). Cultures were inoculated at OD_600_ = 0.1 from overnight culture and incubated at 37 °C, 220 rpm. Protein expression was induced at OD_600_ = 0.6 by addition of 1 mM IPTG, followed by overnight incubation at 18 °C, 220 rpm. Cells were harvested by centrifugation (5000 g, 25 min, 4 °C) and resuspended in 30 mL lysis buffer: 50 mM Tris-HCl pH 7.5, 500 mM NaCl, 5 mM MgCl_2_, 10 mM imidazole, 5 % glycerol, 1 mM PMSF, 1 mg/mL lysozyme, 4 µg/mL DNase, 0.5× Cell Lytic reagent, and one EDTA-free protease inhibitor cocktail tablet (Roche Diagnostics, Mannheim, Germany). Cells were lysed by sonication using 30 s pulses at power setting 16 with 1 min pauses on ice between pulses. The cell pellet was removed by centrifugation (15 000 *rpm*, 25 min, 4 °C). Ni-NTA purification was performed using a 5 mL HisTrap FF column on an Å KTA Pure system (GE Healthcare). Bound proteins were eluted with up to 500 mM imidazole. Eluted proteins were further purified by size exclusion chromatography on a Hiload 16/600 Superdex 75 pg column (GE Healthcare). KRAS^G12C^ was eluted in buffer: 20 mM HEPES, 150 mM NaCl, 1 mM DTT, 5 mM MgCl_2_, and concentrated. Protein purity was checked by SDS-PAGE. Pure KRAS protein was flash frozen in liquid nitrogen and stored at −80 °C.

#### B.5.4 Hit finding GCI KRAS

Screening of KRAS compounds for hits was performed at 500 µM concentration using the rapid kinetics mode of the GCI method (see Section 1.6). Hit calling criteria are summarized in Supplementary Table A3. For KRAS, BI-2865 was used as positive control with a literature *K*_D_ range of 4.5 nM to 12 nM [108]. The immobilised protein was active, with an observed *K*_D_ of 6.9 nM for BI-2865.

#### B.5.5 Blocking experiment

The binding of the covalent inhibitor Sotorasib was assessed in an independent experiment. This confirmed that cysteine 12 was active in GDP-bound KRAS^G12C^ after immobilization, with Sotorasib binding successfully.

A blocking experiment with Sotorasib was performed to confirm that hit molecules bind to the correct pocket. GDP-bound KRAS^G12C^ was incubated in running buffer with 1 % DMSO, while a parallel sample was incubated with 500 µM Sotorasib in running buffer with 1 % DMSO for 10 min. Both samples (free and Sotorasib-blocked) were immobilised on independent channels and responses measured with the GCI rapid kinetics protocol (see Section 1.6).

### B.6 Experimental Details for Pin1

#### B.6.1 Protein Expression, Purification, and Co-crystallization

The open reading frame encoding the Pin1 PPIase domain, incorporating K77Q and K82Q mutations, was subcloned into the pET-28a(+) vector (GenScript), which contains an N-terminal 6×His tag followed by a thrombin cleavage site. The construct was transformed into E. coli BL21(DE3) cells (Novagen) via the heat shock method. Following transformation, colonies were selected by plating on 2×TY agar supplemented with 50 µg/mL kanamycin and incubated for 16 hours at 37 °C. The resulting colony lawn was scraped and used to inoculate 1 L of sterile 2×TY broth containing kanamycin 50 µg/mL. The culture was grown at 37 °C with vigorous shaking until an optical density at 600 nm (OD_600_) of 0.6 was reached. At this point, the flask was cooled under tap water, and protein expression was induced by adding 0.5 mM IPTG (isopropyl *β*-D-1-thiogalactopyranoside). The culture was incubated for an additional 16 hours at 18 °C. Cells were harvested by centrifugation at 6000 g for 10 minutes at 4 °C. The pellet was washed once in PBS, re-centrifuged, and the final 6 g of biomass was stored at −20 °C. For lysis, the pellet was resuspended in lysis buffer (50 mM HEPES pH 7.5, 500 mM NaCl, 20 mM imidazole, 1 mM DTT) at a ratio of 1 g cells per 9 mL buffer. EDTA-free protease inhibitor cocktail (Merck) and DNase I were added prior to sonication. Cell disruption was performed using a Sonics Vibra-Cell VC 130 ultrasonic homogenizer (10 cycles of 30 seconds on, 1 minute off; 45 % amplitude). Lysates were clarified by centrifugation at 60 000 g for 45 minutes at 4 °C (Beckman Coulter Avanti J-26 XP). The supernatant was incubated with Ni-NTA agarose resin (Qiagen) for 2 hours at 4 °C. The resin was washed with 25 column volumes of wash buffer (lysis buffer supplemented with 40 mM imidazole), and bound protein was eluted with elution buffer containing 500 mM imidazole. After SDS-PAGE analysis, the most concentrated fractions were pooled and subjected to thrombin (Merck) digestion for 16 hours at 4 °C during dialysis against cleavage buffer (50 mM Tris-HCl pH 8.0, 150 mM NaCl, 10 mM CaCl_2_). The digested protein was re-incubated with Ni-NTA resin to remove uncleaved His-tagged protein and passed through a HiTrap Benzamidine FF column (Cytiva) to eliminate residual thrombin. The protein was concentrated to 5 mL using a 10,000 MWCO Vivaspin 20 centrifugal concentrator (Sartorius) and subjected to size-exclusion chromatography using a HiLoad 16/600 Superdex 75 column (Cytiva) equilibrated in buffer (10 mM HEPES pH 7.5, 100 mM NaCl). Eluted fractions were analysed by SDS-PAGE and concentrated to 15 mg/mL. For co-crystallization, the purified protein was pre-incubated with 2 mM of the inhibitor compound and 6 % DMSO at 4 °C for 2 hours. Following centrifugation, 2 µL of the supernatant was mixed with 2 µL of reservoir solution (100 mM HEPES pH 7.5, 200 mM ammonium sulfate, 1.2 M sodium citrate) in a hanging drop vapor diffusion setup. Initial crystal growth occurred within 48 hours. These crystals were used as seeds to improve morphology and diffraction quality in a subsequent hanging drop experiment (4 µL total volume: 2 µL protein-inhibitor complex and 2 µL reservoir solution), yielding well-diffracting crystals within 24 hours.

#### B.6.2 X-ray Data Collection, Structure Determination, and Refinement

Prior to data acquisition, crystals were transferred to a cryoprotectant solution consisting of 100 mM HEPES pH 7.5, 40 % glycerol, 5 % DMSO, and 10 mM inhibitor compound. They were flash-cooled in liquid nitrogen and shipped to the European Synchrotron Radiation Facility (ESRF; Grenoble, France) for analysis. X-ray diffraction experiments were conducted at beamline BM07 using a PIXEL Pilatus 6M detector. Data were indexed, integrated, and scaled to a resolution of 2.1 Å using AIMLESS as part of the CCP4i2 suite [115]. The crystal structure was solved by molecular replacement using Phaser within the Phenix software package [116], with the previously reported Pin1 structure (PDB ID: 4TYO) as the search model. Subsequent manual model building was performed using Coot [97], and structural illustrations were generated with PyMOL [117].

