## Supplementary Information (Chemistry) for "A Unified 3D Generative Model for Synthesizable Structure-Based Drug Design"

#### List of compounds and source

| Compound No. | Name | Structure | CAS | Source / commercial with |
|---|---|---|---|---|
| 1 | VH032 | 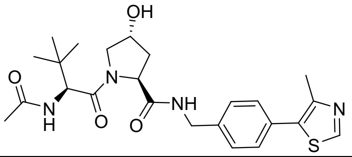 | 1448188-62-2 | Advanced ChemBlocks Inc – Y225646 |
| 2 | NVS-VHL720 | 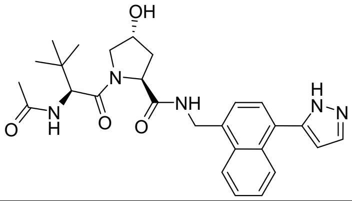 | | See procedure in SI |
| 3 | | 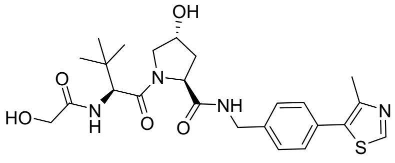 | | See procedure in SI |
| 4 | | 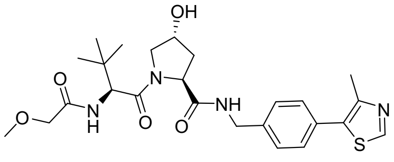 | | See procedure in SI |
| 5 | | 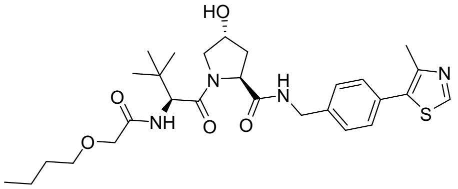 | | See procedure in SI |
| 6 | | 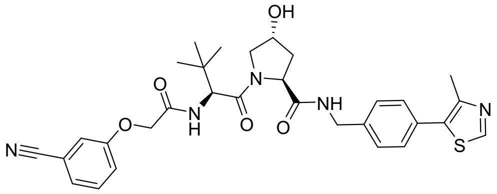 | | See procedure in SI |
| 7 | | 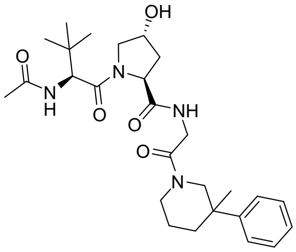 | | See procedure in SI |

| | | | | |
|---|---|---|---|---|
| 8 | | 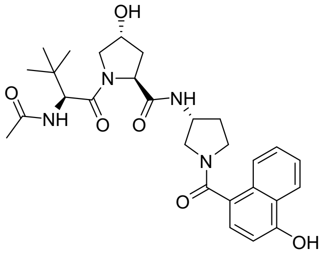 | | See procedure in SI |
| 9 | | 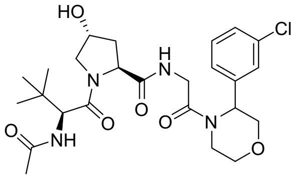 | | See procedure in SI |
| 10 | | 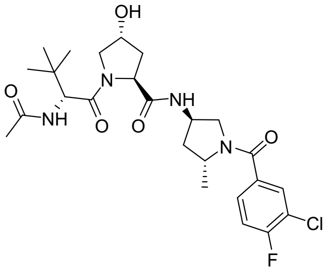 | | See procedure in SI |
| 11 | | 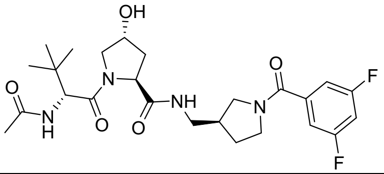 | | See procedure in SI |

#### Synthesis and characterization of compounds

#### Preparation of intermediates for Hyp-OH containing products:

##### 4-(1H-Pyrazol-5-yl)naphthalene-1-carbonitrile (Int-2a)

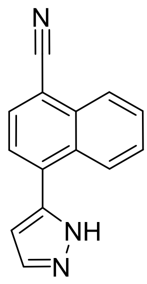


In a 20 mL microwave vial with magnetic stirbar flushed with Ar for 2 min, 4-bromonaphthalene-1-carbonitrile (406.60 mg; 1.72 mmol; 1.00 eq.), (1H-pyrazol-5-yl)boronic acid (595.00 mg; 5.16 mmol; 3.00 eq.), [1,1'-bis(diphenylphosphino)ferrocene]dichloropalladium(II) (193.90 mg; 0.26 mmol; 0.15 eq.), and $Na_2CO_3$ (740.80 mg; 6.99 mmol; 4.07 eq.) were suspended in anhydrous and freshly degassed (sparged with Ar) 1,4-dioxane (10 mL), followed by addition of degassed (sparged with Ar) milliQ water (2 mL). The vial was sealed with a pressure release cap, and sparged with Ar for 5 min. The mixture was then placed in a pre-heated metal block at 90 °C and stirred at 90 °C for 5 days. After 5 days, the mixture was allowed to cool to 25 °C and diluted with water (40 mL). The aqueous layer was extracted with EtOAc (3x 40 mL). The combined organic layer was then washed with brine (1x 120 mL). The organic layer was dried over anhydrous sodium sulfate, filtered, and concentrated in vacuo at 50°C to provide the crude material as a dark brown oil (644.4 mg). The crude material was purified by flash column chromatography using a gradient of 15-35% EtOAc (B) in heptane (A) (LC time program %B: 0.0 min-15%; 3.0 min-15%; 18.0 min-35%; 25.0 min-35%; flow rate: 40 mL/min). Product containing fractions were combined and concentrated in vacuo at 50°C to provide the product as a colorless to pale yellow solid in a purity of 96% (LCMS at 220 nm)and a yield of 74% (292.20 mg; 1.27 mmol).

**LCMS:** Rt: 1.35 min; [M+H$^+$]: 220.1 (Method B).

**$^1$H NMR (700 MHz, DMSO-d$_6$)** δ [ppm] 13.34 (s, 1H), 8.94 (dt, J = 8.6, 1.0 Hz, 1H), 8.21 – 8.15 (m, 2H), 7.98 (dd, J = 2.2, 1.0 Hz, 1H), 7.87 (d, J = 7.5 Hz, 1H), 7.83 (ddd, J = 8.2, 6.8, 1.2 Hz, 1H), 7.74 (ddd, J = 8.3, 6.8, 1.3 Hz, 1H), 6.83 – 6.78 (m, 1H).

##### 1-[4-(1H-Pyrazol-5-yl)naphthalen-1-yl]methanamine (Int-2b)

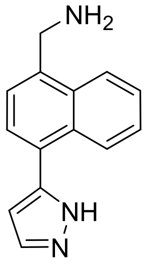


The reaction was carried out in a Büchi glass lab pressure reactor. To 4-(1H-pyrazol-5-yl)naphthalene-1-carbonitrile (287.00 mg; 1.24 mmol; 1.00 equiv.) was added MeOH (5 mL) and THF (3 mL). RaNi (approx. 0.30 g) was washed with MeOH (2x) and then added to the reaction mixture. To the mixture was then added a solution of $NH_3$ in MeOH (20%) (5 mL) and the pressure reactor sealed. The reaction was flushed with $N_2$ (3x), followed by $H_2$ (3x). The reaction was then stirred at 55°C and 5.5 bar ($H_2$) for 16 h. The reaction was then filtered over kieselguhr under vacuum and the kieselguhr washed with THF and MeOH. The filtrate was concentrated in vacuo at 50 °C to provide the crude material as a pale yellow-orange solid in a purity of 89% (LCMS at 220 nm) and a yield of 95% (296.00 mg; 1.18 mmol).

**LCMS**: Rt: 0.81 min; [M-$NH_3$]: 207.2 (Method B).

#### *tert*-Butyl *N*-[2-(3-methyl-3-phenylpiperidin-1-yl)-2-oxoethyl]carbamate (Int-7a)

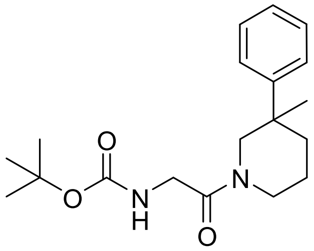

To a solution of 3-methyl-3-phenylpiperidine (110.00 mg; 0.60 mmol; 1.00 eq.) and Boc-glycine (109.90 mg; 0.63 mmol; 1.05 eq.) in anhydrous DMF (2 mL) was added DIPEA (96.00 µL; 0.54 mmol; 0.91 eq.). The mixture was cooled to 0°C. Next, HATU (216.00 mg; 0.57 mmol; 0.95 eq.) was added at 0 °C and the reaction mixture stirred at 0 °C for 10 min. Then, the reaction was allowed to warm to 25°C, and stirred at 25°C for 20 h. After 20 h, the mixture was diluted with EtOAc (20 mL) and water (20 mL) and the phases separated. The aqueous phase was extracted with EtOAc (2x 20 mL). The combined organic phase was washed with brine (2x80 mL), dried over anhydrous sodium sulfate, filtered and concentrated in vacuo at 50 °C to provide the crude material as a colorless to pale yellow oil / sticky residue (274.9 mg). The crude material was purified by flash column chromatography using a gradient of 0-10% MeOH (B) in DCM (A) (LC time program %B: 0.0 min-0%; 2.4 min-0%; 6.3 min-5%; 14.9 min-10%; 25.0 min-10%; flow rate: 30 mL/min). Product containing fractions were combined and concentrated in vacuo at 35°C to provide the product as a colorless oil in a purity of 81% (LCMS at 220 nm) and a yield of 70% (172 mg; 0.42 mmol).

**LCMS:** Rt: 1.81 min; [$M+2H^+-C_4H_9$]: 277.2 (Method A).

#### 2-Amino-1-(3-methyl-3-phenylpiperidin-1-yl)ethan-1-one hydrochloride (Int-7b)

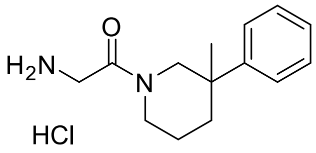

To a solution of *tert*-butyl *N*-[2-(3-methyl-3-phenylpiperidin-1-yl)-2-oxoethyl]carbamate (172.00 mg; 0.42 mmol; 1.00 eq.) in anhydrous DCM (2 mL) was added hydrogen chloride solution 4.0M in dioxane (523.34 µL; 2.09 mmol; 5.00 eq.) at 22°C. Upon addition the solution turned into a colorless suspension. The mixture was stirred at 22°C for 24 h. After 24 h, the mixture was concentrated in vacuo at 40 °C to provide the crude product as the HCl salt 2-amino-1-(3-methyl-3-phenylpiperidin-1-yl)ethan-1-one hydrochloride (177.70 mg; 0.42 mmol). The crude product was obtained as a yellow oil with a purity of approx. 63%. The yield was assumed to be quantitative. The crude was used directly in the next step without further purification.

**LCMS:** Rt: 0.89 min; [$M+H^+$]: 233.2 (Method A).

#### *tert*-Butyl *N*-[(3*R*)-1-(4-hydroxynaphthalene-1-carbonyl)pyrrolidin-3-yl]carbamate (Int-8a)

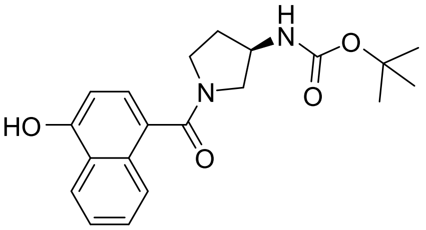

To a solution of 4-hydroxynaphthalene-1-carboxylic acid (101.10 mg; 0.51 mmol; 1.00 eq.) and (R)-(+)-3-(Boc-amino)pyrrolidine (97.20 mg; 0.52 mmol; 1.01 eq.) in anhydrous DMF (2 mL) was added DIPEA (89.70 µL; 0.50 mmol; 0.99 eq.). The mixture was cooled to 0°C. Next, HATU (204.20 mg; 0.54 mmol; 1.05 eq.) was added at 0 °C and the reaction mixture stirred at 0 °C for 10 min. Then, the reaction was allowed to warm to 25°C, and stirred at 25°C for 30 h. After 30 h, the mixture was diluted with EtOAc (20 mL) and brine (20 mL) and the phases separated. The aqueous phase was extracted with EtOAc (2x 20 mL). The combined organic phase was washed with brine (2x80 mL), dried over anhydrous sodium sulfate, filtered and concentrated in vacuo at 50 °C to provide the crude material as an orange oil (382.1 mg). The crude material was purified by flash column chromatography using a gradient of 0-10% MeOH (B) in DCM (A) (LC time program %B: 0.0 min-0%; 2.0 min-0%; 17.0 min-10%; 25.0 min-10%; flow rate: 30 mL/min). Product containing fractions were combined and concentrated in vacuo at 35°C to provide the product as a pale orange film in a purity of 97% (LCMS at 220 nm) and a yield of 52% (96.9 mg; 0.27 mmol).

**LCMS:** Rt: 1.38 min; $[M+H^+]$: 357.2 (Method A).

**$^1$H NMR (400 MHz, DMSO-d$_6$)** (mixture of 2 species; major:minor approx. 1:0.7) δ [ppm] 10.50 – 10.41 (m, 1.7 H, major+minor), 8.21 – 8.14 (m, 1.7 H, major+minor), 7.82 – 7.72 (m, 1.7H, major+minor), 7.57 – 7.42 (m, 3.4 H, major+minor), 7.33 (d, *J* = 7.8 Hz, 0.7 H, minor), 7.36 – 7.24 (m, 1.7 H, major+minor), 7.16 (d, *J* = 6.2 Hz, 1 H, major), 6.90 – 6.83 (m, 1.7 H, major+minor), 4.13 – 4.01 (m, 0.7 H, minor), 3.91 – 3.81 (m, 1H, major), 3.80 – 3.65 (m, 1.7 H, major+minor), 3.62 – 3.49 (m, 1H, major), 3.45 – 3.35 (m, 0.7 H, minor), 3.28 – 3.19 (m, 1.7 H, major+minor), 3.17 (d, *J* = 5.1 Hz, 0.7 H, minor), 3.14 – 3.04 (m, 0.7 H, minor), 2.96 – 2.84 (m, 1 H, major), 2.14 – 2.02 (m, 1 H, major), 2.01 – 1.89 (m, 0.7 H, minor), 1.89 – 1.77 (m, 1H, major), 1.74 – 1.62 (m, 0.7 H, minor), 1.41 (s, 6.3 H, minor), 1.28 (s, 9 H, major).

#### 4-[(3*R*)-3-Aminopyrrolidine-1-carbonyl]naphthalen-1-ol hydrochloride (Int-8b)

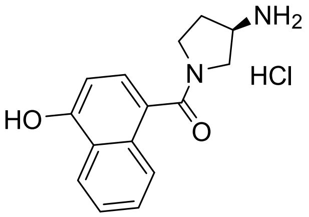

To a solution of *tert*-butyl *N*-[(3*R*)-1-(4-hydroxynaphthalene-1-carbonyl)pyrrolidin-3-yl]carbamate (96.90 mg; 0.27 mmol; 1.00 eq.) in anhydrous DCM (2 mL) was added hydrogen chloride solution 4.0M in dioxane (331.69 µl; 1.33 mmol; 5.00 eq.) at 22°C. Upon addition the solution turned into a yellow suspension. The mixture was stirred at 22 °C for 24 h. After 24 h, the mixture was concentrated in vacuo at 35 °C to provide the crude product as the HCl salt 4-[(3*R*)-3-aminopyrrolidine-1-carbonyl]naphthalen-1-ol hydrochloride (81.90 mg; 0.27 mmol). The crude product was obtained as a colorless solid with a purity of approx. 95%. The crude was used directly in the next step without further purification. The yield was assumed to be quantitative.

**LCMS:** Rt: 0.64 min; $[M+H^+]$: 257.2 (Method A)

**$^1$H NMR (400 MHz, DMSO-d$_6$)** δ [ppm] 10.61 (s, 1H), 8.44 (s, 1H), 8.25 (s, 1H), 8.22 – 8.15 (m, 1H), 7.87 – 7.80 (m, 1H), 7.57 – 7.44 (m, 2H), 7.44 – 7.28 (m, 1H), 6.97 – 6.87 (m, 1H), 3.97 – 3.59 (m, 3H), 3.46 – 3.34 (m, 1H), 3.24 – 3.10 (m, 1H), 2.37 – 1.86 (m, 2H).

#### *tert*-Butyl N-{2-[3-(3-chlorophenyl)morpholin-4-yl]-2-oxoethyl}carbamate (Int-9a)

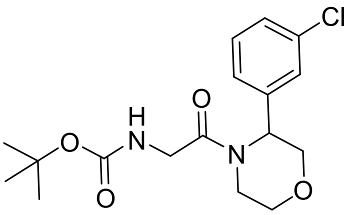


To a suspension of 3-(3-chlorophenyl)morpholine hydrochloride (100.30 mg; 0.41 mmol; 1.00 eq.) and Boc-glycine (94.30 mg; 0.54 mmol; 1.32 eq.) in anhydrous DMF (2 mL) was added DIPEA (72.0 µL; 0.41 mmol; 1.00 eq.). The mixture was cooled to 0°C. Next, HATU (159.70 mg; 0.42 mmol; 1.03 eq.) was added at 0 °C and the reaction mixture stirred at 0 °C for 10 min. Then, the reaction was allowed to warm to 25°C, and stirred at 25°C for 24 h. After 24 h, the mixture was diluted with EtOAc (20 mL) and brine (20 mL) and the phases separated. The aqueous phase was extracted with EtOAc (2x 20 mL). The combined organic phase was washed with brine (2x80 mL), dried over anhydrous sodium sulfate, filtered and concentrated in vacuo at 50 °C to provide the crude material as a pale orange oil (149.5 mg). The crude material was purified by flash column chromatography using a gradient of 0-10% MeOH (B) in DCM (A) (LC time program %B: 0.0 min-0%; 2.9 min-0%; 20.0 min-10%; 25.0 min-10%; flow rate: 30 mL/min). Product containing fractions were combined and concentrated in vacuo at 35°C to provide the product as a colorless oil in a purity of 91% (LCMS at 220 nm) and a yield of 54% (85.0 mg; 0.22 mmol).

**LCMS:** Rt: 1.64 min; m/z [$M+2H^+-C_4H_9$] for $C_{13}H_{16}ClN_2O_4^+$: 299.1 (Method A).

#### 2-Amino-1-[3-(3-chlorophenyl)morpholin-4-yl]ethan-1-one hydrochloride (Int-9b)

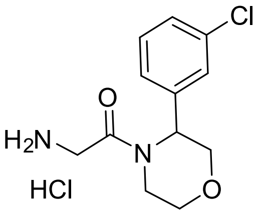


To a solution of (*tert*-butyl *N*-{2-[3-(3-chlorophenyl)morpholin-4-yl]-2-oxoethyl}carbamate (139.40 mg; 0.36 mmol; 1.00 eq.)) in anhydrous DCM (2 mL) was added hydrogen chloride solution 4.0M in dioxane (444.0 µL; 1.78 mmol; 5.00 eq.) at 22°C. The mixture was stirred at 22°C for 20 h. After 20 h, the mixture was concentrated under reduced pressure at 35 °C to provide the crude product as the HCl salt 2-amino-1-[3-(3-chlorophenyl)morpholin-4-yl]ethan-1-one hydrochloride (116.10 mg; 0.36 mmol). The crude product was obtained as a colorless to pale yellow solid in a purity of approx. 89%. The yield was assumed to be quantitative. The crude hydrochloride salt was used directly in the next step without further purification.

**LCMS:** Rt: 0.78 min; [$M+H^+$]: 255.1 (Method A).

#### *tert*-Butyl *N*-[(3*R*,5*R*)-1-(3-chloro-4-fluorobenzoyl)-5-methylpyrrolidin-3-yl]carbamate (Int-10a)

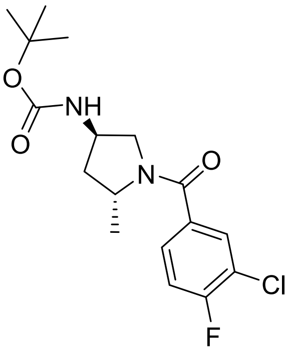

To a suspension of tert-butyl *N*-[(3*R*,5*R*)-5-methylpyrrolidin-3-yl]carbamate (122.30 mg; 0.59 mmol; 1.15 eq.) and 3-chloro-4-fluorobenzoic acid (95.00 mg; 0.52 mmol; 1.00 eq.) in anhydrous DMF (4 mL) was added DIPEA (540 µL; 3.04 mmol; 5.88 eq.). The mixture was cooled to 0°C. Next, HATU (300.3 mg; 0.79 mmol; 1.53 eq.) was added at 0 °C and the reaction mixture stirred at 0 °C. After 10 min at 0°C, the reaction was allowed to warm to 25°C and stirred at 25°C for 4 h. After 4 h, the mixture was diluted with DCM (20 mL) and water (20 mL) and the phases separated. The aqueous phase was extracted with DCM (2x 20 mL). The combined organic phase was washed with brine (2x80 mL), dried over anhydrous sodium sulfate, filtered and concentrated in vacuo at 50 °C to provide the crude material as a dark brown oil (368.4 mg). The crude material was purified by flash column chromatography using a gradient of 0-5% MeOH (B) in DCM (A) (LC time program %B: 0.0 min-0%; 3.0 min-0%; 20.0 min-5%; 22.9min-5%; flow rate: 30 mL/min). The product containing fraction was concentrated in vacuo at 40°C to provide the product as an orange solid in a purity of 98% (LCMS at 220 nm) and a yield of 83% (156.9 mg; 0.43 mmol).

**LCMS:** Rt: 1.68 min; [M+Na$^+$] [$C_{17}H_{22}ClFN_2NaO_3^+$]: 379.1 (Method A).

**$^1$H NMR (400 MHz, DMSO-d$_6$)** δ [ppm] 7.71 – 7.62 (m, 1H), 7.56 – 7.42 (m, 2H), 7.14 (s, 1H), 4.28 – 4.17 (m, 1H), 3.90 (s, 1H), 3.72 – 3.55 (m, 1H), 3.24 – 3.10 (m, 1H), 2.09 – 1.94 (m, 1H), 1.76 – 1.66 (m, 1H), 1.32 (s, 9H), 1.27 – 1.22 (m, 3H).

**Note:** Residual amounts of DCM (<1% by weight) and DMF (3% by weight) observed. Also, unknown minor species observed in $^1$H NMR.

#### (3*R*,5*R*)-1-(3-Chloro-4-fluorobenzoyl)-5-methylpyrrolidin-3-amine hydrochloride (Int-10b)

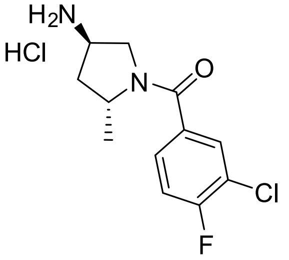

To a solution of *tert*-butyl *N*-[(3*R*,5*R*)-1-(3-chloro-4-fluorobenzoyl)-5-methylpyrrolidin-3-yl]carbamate (156.90 mg; 0.43 mmol; 1.00 eq.) in anhydrous DCM (2 mL) was added hydrogen chloride solution 4.0M in dioxane (536.90 µL; 2.15 mmol; 5.00 eq.) at 22°C. The mixture was stirred at 22°C for 23 h.

After 23 h, the mixture was concentrated in vacuo at 35 °C to provide the crude product as the HCl salt (3*R*,5*R*)-1-(3-chloro-4-fluorobenzoyl)-5-methylpyrrolidin-3-amine hydrochloride. The crude product was obtained as a yellow solid in a purity of approx. 89% (assigned based on $^1$H NMR analysis) and a yield of 96% (136.30 mg; 0.41 mmol). The crude material was used directly in the next step without further purification.

**LCMS**: Rt: 0.80 min; [M+H$^+$] found: 257.1 (Method A).

**$^1$H NMR (400 MHz, DMSO-d$_6$)** δ [ppm] 8.46 (s, 1H), 8.22 (s, 2H), 7.78 – 7.45 (m, 3H), 4.38 – 4.17 (m, 1H), 3.91 – 3.72 (m, 2H), 3.42 – 3.31 (m, 1H), 2.35 – 2.21 (m, 1H), 1.97 – 1.82 (m, 1H), 1.34 – 1.18 (m, 3H).

Note: Residual 1,4-dioxane (approx. 11% by weight) in $^1$H NMR spectrum observed.

#### *tert*-Butyl *N*-{[(3*S*)-1-(3,5-difluorobenzoyl)pyrrolidin-3-yl]methyl}carbamate (Int-11a)

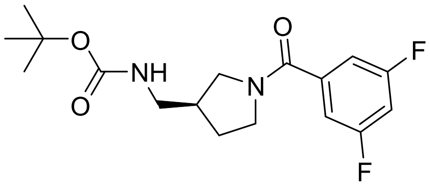


To a suspension of *tert*-butyl (3*R*)-3-pyrrolidinylmethylcarbamate hydrochloride (237.40 mg; 1.00 mmol; 1.12 eq.) and 3,5-difluorobenzoic acid (144.50 mg; 0.90 mmol; 1.00 eq.) in anhydrous DMF (4 mL) was added DIPEA (573 µL; 3.22 mmol; 3.60 eq.). The mixture was cooled to 0 °C. AT 0 °C, HATU (361.30 mg; 0.95 mmol; 1.06 eq.) was added and the reaction mixture stirred at 0 °C for 10 min. Then, the reaction was allowed to warm to 25°C and stirred at 25 °C for 23 h. After 23 h, the mixture was diluted with DCM (20 mL) and water (20 mL) and the phases separated. The aqueous phase was extracted with DCM (2x 20 mL). The combined organic phase was washed with brine (2x60 mL), dried over anhydrous sodium sulfate, filtered and concentrated in vacuo at 40 °C to provide the crude material as an orange oil (486.5 mg). The crude material was purified by flash column chromatography using a gradient of 0-10% MeOH (B) in DCM (A) (LC time program %B: 0.0 min-0%; 2.0 min-0%; 20.0 min-10%; 22.7 min-10%; 25.0 min-15%; 35.0 min-15%; flow rate: 30 mL/min). The product containing fraction was concentrated in vacuo at 40°C to provide the product as a pale orange oil in a purity of 69% (assigned based on $^1$H NMR analysis) and a yield of 69% (303.00 mg; 0.61 mmol).

**LCMS:** Rt: 1.54 min; [M+Na$^+$] [$C_{17}H_{22}F_2N_2NaO_3$]: 363.2 (Method A).

**$^1$H NMR (400 MHz, DMSO-d$_6$)** (mixture of 2 species; minor: major = 1.0:0.8) δ [ppm] 7.40 – 7.17 (m, 5.4H, major+minor), 7.03 (t, J = 5.8 Hz, 0.8H, minor), 6.92 (t, J = 5.9 Hz, 1H, major), 3.59 – 3.35 (m, 5.4H, major+minor), 3.20 – 3.08 (m, 2.6H, major+minor), 3.05 – 2.79 (m, 2.8H, major+minor), 2.38 – 2.18 (m, 1.8H, major+minor), 1.99 – 1.82 (m, 1.8H, major+minor), 1.68 – 1.51 (m, 1.8H, major+minor), 1.39 (s, 7.2H, minor), 1.33 (s, 9H, major).

Note: The product was observed to exist in solution as a mixture of at least 2 species. The $^1$H NMR spectrum displays residual amounts of DCM (1% by weight), DMF (<1% by weight) and an unidentified impurity.

#### 1-[(3*S*)-1-(3,5-Difluorobenzoyl)pyrrolidin-3-yl]methanamine hydrochloride (Int-11b)

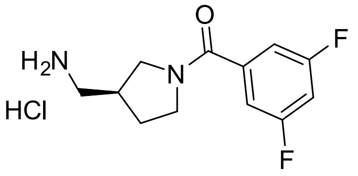


To a solution of *tert*-butyl *N*-{[(3*S*)-1-(3,5-difluorobenzoyl)pyrrolidin-3-yl]methyl}carbamate (303.00 mg; 0.61 mmol; 1.00 eq.) in anhydrous DCM (2 mL) was added hydrogen chloride solution 4.0M in dioxane (767.82 µL; 3.07 mmol; 5.00 eq.) at 22°C. The mixture was stirred at 22°C for 19 h. After 19 h, the mixture was concentrated in vacuo at 35 °C to provide the crude product as the HCl salt 1-[(3S)-1-(3,5-difluorobenzoyl)pyrrolidin-3-yl]methanamine hydrochloride (252.00 mg; 0.61 mmol). The crude product was obtained as a pale orange solid with a purity of approx. 68%. The yield was assumed to be quantitative. The crude was used directly in the next step without further purification.

**LCMS:** Rt: 0.63 min; $[M+H^+]$: 241.1 (Method A).

#### Preparation of products modified on the right-hand side of the Hyp-OH core

##### (2*S*,4*R*)-1-[(2*S*)-2-Acetamido-3,3-dimethylbutanoyl]-4-hydroxy-*N*-{[4-(1H-pyrazol-5-yl)naphthalen-1-yl]methyl}pyrrolidine-2-carboxamide (Compound 2)

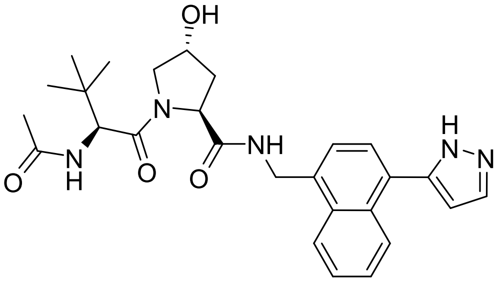


(2*S*,4*R*)-1-[(2*S*)-2-Acetamido-3,3-dimethylbutanoyl]-4-hydroxy-N-{[4-(1H-pyrazol-5-yl)naphthalen-1-yl]methyl}pyrrolidine-2-carboxamide was prepared following the literature procedure described by Tutter *et al.*[2] with minor modifications. To a suspension of (2*S*,4*R*)-1-[(2*S*)-2-acetamido-3,3-dimethylbutanoyl]-4-hydroxypyrrolidine-2-carboxylic acid (258.50 mg; 0.86 mmol; 1.00 eq.) and the crude 1-[4-(1H-pyrazol-5-yl)naphthalen-1-yl]methanamine (296.00 mg; 1.18 mmol; 1.38 eq.) in anhydrous DMF (4 mL) was added DIPEA (458.12 µL; 2.58 mmol; 3.00 eq.). The mixture was cooled to 0 °C. Next, HATU (454.40 mg; 1.20 mmol; 1.39 eq.) was added at 0 °C and the reaction mixture stirred at 0 °C for 15 min. Then, the reaction was allowed to warm to 25°C and stirred at 25°C for 24 h. After 24 h, the mixture was diluted with EtOAc (30 mL) and water (30 mL) and the phases separated. The aqueous phase was extracted with EtOAc (2x 30 mL). The combined organic phase was washed with water (1x 100mL) and brine (1x100 mL). The organic phase was dried over anhydrous sodium sulfate, filtered and concentrated in vacuo at 50 °C to provide the crude material as a pale yellow oil (511.7 mg). The crude was purified via preparative RP HPLC on a 40 g flash column (IR-C18, F0040) via liquid injection (solution in 4 mL DMSO) using a gradient of MeCN+0.1% FA (B) in water+0.1% FA (A) (LC-time-%B: 0 min-0%; 5 min-0%; 25 min-40%; 30 min-40%; 30 min-100%; 35 min-100%; 35 min-50%; 55 min-50%; flow rate: 40 mL/min). Product containing fractions were combined and concentrated in vacuo at 50°C to provide the product as a colorless solid in a purity of 92% (LCMS purity at 254 nm) and yield of 45% (207.9 mg; 0.39 mmol).

**LCMS**: 1.23 min; 492.3 $[M+H^+]$ (Method A).

**$^1$H NMR (700 MHz, DMSO-$d_6$)** δ [ppm] 13.22 (s, 1H (minor)), 13.05 (s, 1H (major)), 8.74 – 8.63 (m, 1H), 8.63 – 8.52 (m, 1H), 8.08 (s, 1H), 7.93 (d, *J* = 9.4 Hz, 1H), 7.90 (s, 1H), 7.71 – 7.43 (m, 4H), 6.65 – 6.49 (m, 1H), 5.11 (d, *J* = 3.7 Hz, 1H), 4.81 (dd, *J* = 15.5, 5.9 Hz, 1H), 4.75 – 4.69 (m, 1H), 4.54 (d, *J* = 9.5 Hz, 1H), 4.46 (t, *J* = 8.0 Hz, 1H), 4.38 – 4.32 (m, 1H), 3.69 (dd, *J* = 10.4, 4.2 Hz, 1H), 3.63 (dt, *J* = 10.5, 1.9 Hz, 1H), 2.04 – 1.98 (m, 1H), 1.95 – 1.89 (m, 1H), 1.88 (s, 3H), 0.94 (s, 9H).

Note: The product exists in solution as a mixture of 2 species. The $^1$H NMR spectrum is in agreement with literature.[2]

#### (2*S*,4*R*)-1-[(2*S*)-2-Acetamido-3,3-dimethylbutanoyl]-4-hydroxy-N-[2-(3-methyl-3-phenylpiperidin-1-yl)-2-oxoethyl]pyrrolidine-2-carboxamide (Compound 7)

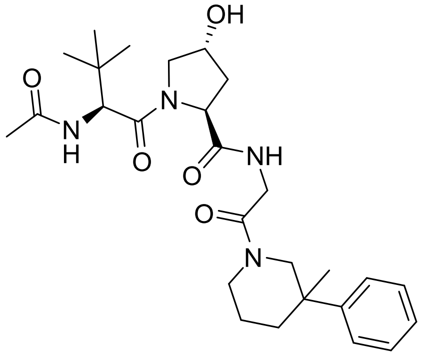

(2*S*,4*R*)-1-[(2*S*)-2-Acetamido-3,3-dimethylbutanoyl]-4-hydroxypyrrolidine-2-carboxylic acid (130.00 mg; 0.43 mmol; 1.03 eq.) and crude 2-amino-1-(3-methyl-3-phenylpiperidin-1-yl)ethan-1-one hydrochloride (177.70 mg; 0.42 mmol; 1.00 eq.) were suspended in anhydrous DMF (4 mL), followed by addition of DIPEA (595.53 µL; 3.35 mmol; 8.00 eq.). The mixture was cooled to 0°C. Next, HATU (205.40 mg; 0.54 mmol; 1.29 eq.) was added at 0 °C and the reaction mixture stirred at 0 °C for 10 min. Then, the reaction was allowed to warm to 25°C, and stirred at 25°C for 23 h. After 23 h, the mixture was diluted with EtOAc (20 mL) and water (20 mL) and the phases separated. The aqueous phase was extracted with EtOAc (2x 20 mL). The combined organic phase was washed with brine (2x80 mL), dried over anhydrous sodium sulfate, filtered and concentrated in vacuo at 50 °C to provide the crude material as a yellow oil (338.9 mg). The crude material was purified by flash column chromatography using a gradient of 0-10% MeOH (B) in DCM (A) (LC time program %B: 0.0 min-0%; 1.5 min-0%; 16.0 min-10%; 25.0 min-10%; flow rate: 30 mL/min). Product containing fractions were combined and concentrated in vacuo at 40°C to provide the product as a mixture of at least 2 species (likely diastereoisomers as indicated) as a colorless solid in a purity of 96% (assigned based on $^1$H NMR analysis) and a yield of 76% (166.1 mg; 0.32 mmol).

**LCMS:** Rt: 1.40 min; [M+H$^+$]: 501.3; [M+Na$^+$]: 523.3 (Method A).

**$^1$H NMR (400 MHz, DMSO-d$_6$)** δ [ppm] (mixture of at least 2 species in a ratio of approx. major:minor = 1.0:0.5) 8.08 – 7.94 (m, 1.5H, major+minor), 7.92 – 7.84 (m, 1.5H, major+minor), 7.50 – 7.13 (m, 7.5H, major+minor), 5.14 – 5.05 (m, 1.5H, major+minor), 4.55 – 4.49 (m, 1.5H, major+minor), 4.49 – 4.41 (m, 1.5H, major+minor), 4.33 – 4.23 (m, 1.5H, major+minor), 4.19 – 3.36 (m, 11.5H, major+minor), 3.06 – 2.89 (m, 0.5H, minor), 2.11 – 1.89 (m, 4.5H, major+minor), 1.89 – 1.83 (m, 4H, major+minor), 1.83 – 1.50 (m, 4H, major+minor), 1.44 – 1.31 (m, 1H, major), 1.25 – 1.18 (m, 1.5H, minor), 1.14 – 1.09 (m, 3H, major), 0.93 (s, 9H, major, partially overlaps with minor signal), 0.92 (s, 4.5H, minor, partially overlaps with major signal).

Note: Residual amount of DCM in NMR spectrum observed 4% (by weight).

Note: Product obtained as a mixture of at least 2 species according to $^1$H NMR spectrum. The ratio of the species is approx. 1.0:0.5. The species are likely diastereomers since 2-amino-1-(3-methyl-3-phenylpiperidin-1-yl)ethan-1-one hydrochloride (177.70 mg; 0.42 mmol; 1.00 eq.) is derived from a racemic building block.

#### (2*S*,4*R*)-1-[(2*S*)-2-Acetamido-3,3-dimethylbutanoyl]-4-hydroxy-*N*-[(3*R*)-1-(4-hydroxynaphthalene-1-carbonyl)pyrrolidin-3-yl]pyrrolidine-2-carboxamide (Compound 8)

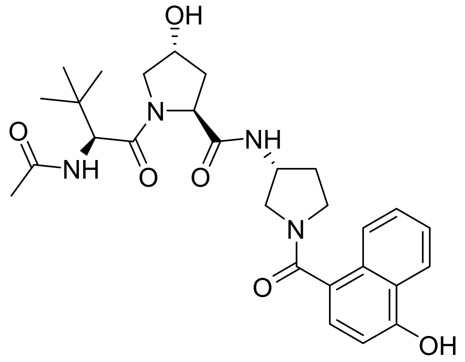


(2*S*,4*R*)-1-[(2*S*)-2-Acetamido-3,3-dimethylbutanoyl]-4-hydroxypyrrolidine-2-carboxylic acid (86.40 mg; 0.29 mmol; 1.08 eq.) and crude 4-[(3*R*)-3-aminopyrrolidine-1-carbonyl]naphthalen-1-ol hydrochloride (81.90 mg; 0.27 mmol; 1.00 eq.) were supsended in anhydrous DMF (2 mL), followed by addition of DIPEA (377.29 µL; 2.12 mmol; 8.00 eq.). The mixture was cooled to 0°C. Next, HATU (123.80 mg; 0.33 mmol; 1.23 eq.) was added at 0 °C and the reaction mixture stirred at 0 °C for 10 min. Then, the reaction was allowed to warm to 25°C, and stirred at 25°C for 21 h. After 21 h, the mixture was diluted with EtOAc (20 mL) and water (20 mL) and the phases separated. The aqueous phase was extracted with EtOAc (2x 20 mL). The combined organic phase was washed with brine (2x80 mL), dried over anhydrous sodium sulfate, filtered and concentrated in vacuo at 50 °C to provide a pale yellow oil (36.2 mg; primarily bis-acylated product based on LCMS analysis). The aqueous phase was acidifed with 1N aqueous HCl to pH 2 and then extracted with DCM (3x 40 mL) and the organic phase dried over anhydrous sodium sulfate, filtered and concentrated at 40°C. The aqueous phase was checked again by LCMS and still contained product. Therefore, the acidified aqueous phase was saturated with solid sodium chloride and the saturated aqueous solution extracted with EtOAc (3x30 mL). The organic phase was dried over anhydrous sodium sulfate, filtered and concentrated in vacuo at 40°C. The DCM-extracted phase- and EtOAc-extracted phase-derived crude materials were combined and concentrated in vacuo to provide the crude product as a yellow liquid (542.3 mg). The crude material (542.3 mg) was purified by flash column chromatography using a gradient of 0-15% MeOH (B) in DCM (A) (LC time program %B: 0.0 min-0%; 1.6 min-0%; 17.0 min-15%; 31.2 min-15%, 34.9 min - 20%; 39.9 min - 20%; flow rate: 40 mL/min). Product containing fractions were combined and concentrated in vacuo at 40°C to provide a colorless solid (37.9 mg). The product was repurified via reverse phase preparative HPLC using a linear gradient from 0-60% MeCN+0.1% FA (B) in water+0.1% FA (A) (LC-time-%B: 0 min-10%; 2 min-10%; 18 min-60%; 20 min-60%; 20 min-50%; 25 min-50%; flow rate: 50 mL/min). Product containing fractions were combined and concentrated under reduced pressure at 50°C temperature. The product (2*S*,4*R*)-1-[(2*S*)-2-acetamido-3,3-dimethylbutanoyl]-4-hydroxy-*N*-[(3*R*)-1-(4-hydroxynaphthalene-1-carbonyl)pyrrolidin-3-yl]pyrrolidine-2-carboxamide was obtained as a colorless solid in a purity of >99% and a yield of 17% (23.00 mg; 0.04 mmol).

**LCMS:** Rt: 1.08 min; [M+H$^+$]: 525.3 (Method A).

**$^1$H NMR (400 MHz, DMSO-d$_6$)** (mixture of rotamers: minor:major = 1: 1.3 ) δ [ppm] 10.58 – 10.32 (m, 2.3H, major+minor), 8.26 (d, J = 6.9 Hz, 1H, minor), 8.22 – 8.07 (m, 3.6H, major+minor), 7.93 (d, J = 9.3 Hz, 1H, minor), 7.86 – 7.70 (m, 3.6H, major+minor), 7.59 – 7.39 (m, 4.6H, major+minor), 7.37 – 7.23 (m, 2.3H, major+minor), 6.95 – 6.75 (m, 2.3H, major+minor), 5.17 – 5.02 (m, 2.3H, major+minor), 4.56 – 4.41 (m, 2.3H, major+minor), 4.38 – 4.06 (m, 6.9H, major+minor), 3.84 – 2.91 (m, 13.8H, major+minor), 2.17 – 1.65 (m, 9.2H, major+minor), 1.88 (s, 3H, minor), 1.84 (s, 3.9H, major), 0.96 (s, 9H, minor), 0.81 (s, 11.7H, major).

Note: The product was obtained as a mixture of at least 2 species according to the $^1$H NMR spectrum. To test whether the species are rotamers, variable temperature NMR was recorded at 303K, 330 K and 363 K. Coalescence of signals (*e.g.* Me) at high temperatures supports that the product exists as a mixture of rotamers in solution.

#### (2*S*,4*R*)-*N*-{2-[3-(3-Chlorophenyl)morpholin-4-yl]-2-oxoethyl}-1-[(2*S*)-2-acetamido-3,3-dimethylbutanoyl]-4-hydroxypyrrolidine-2-carboxamide (Compound 9)

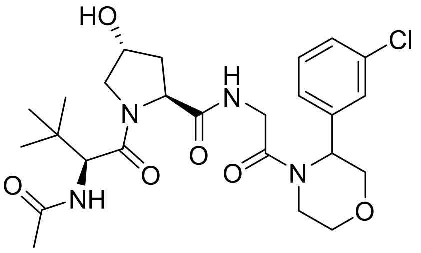


To a suspension of (2*S*,4*R*)-1-[(2*S*)-2-acetamido-3,3-dimethylbutanoyl]-4-hydroxypyrrolidine-2-carboxylic acid (110.50 mg; 0.37 mmol; 1.03 eq.) and 2-amino-1-[3-(3-chlorophenyl)morpholin-4-yl]ethan-1-one hydrochloride (116.10 mg; 0.36 mmol; 1.00 eq.) in anhydrous DMF (4 mL) was added DIPEA (505.16 µL; 2.84 mmol; 8.00 eq.). The mixture was cooled to 0°C. Next, HATU (167.50 mg; 0.44 mmol; 1.24 eq.) was added at 0 °C and the reaction mixture stirred at 0 °C for 10 min. Then, the reaction was allowed to warm to 25°C, and stirred at 25°C for 20 h. After 20 h, the mixture was diluted with EtOAc (20 mL) and water (20 mL) and the phases separated. The aqueous phase was extracted with EtOAc (2x 20 mL). The combined organic phase was washed with brine (2x80 mL), dried over anhydrous sodium sulfate, filtered and concentrated in vacuo at 50 °C to provide the crude material as a pale yellow film (122.5 mg). The crude material was purified by flash column chromatography using a gradient of 0-10% MeOH (B) in DCM (A) (LC time program %B: 0.0 min-0%; 1.0 min-0%; 16.0 min-10%; 25.0 min-10%; flow rate: 30 mL/min). Product containing fractions were combined and concentrated in vacuo at 40°C to provide the product as a colorless solid in a purity of 99% (LCMS at 220 nm) and a yield of 37% (68.4 mg; 0.13 mmol).

**LCMS:** Rt: 1.27 min; [M+H$^+$]: 523.2 (Method A).

**$^1$H NMR (400 MHz, DMSO-d$_6$)** δ [ppm] 8.24 – 8.09 (m, 1H), 7.88 (d, *J* = 9.9 Hz, 1H), 7.46 – 7.24 (m, 4H), 5.45 – 5.29 (m, 1H), 5.15 – 5.05 (m, 1H), 4.51 (d, *J* = 9.3 Hz, 1H), 4.48 – 4.11 (m, 4H), 4.12 – 3.99 (m, 1H), 3.93 – 3.38 (m, 6H), 3.27 – 3.09 (m, 1H), 2.06 – 1.89 (m, 2H), 1.89 – 1.83 (m, 3H), 0.93 (s, 9H).

Note: Residual amount of DCM in NMR spectrum observed 8% (by weight).

#### (2*S*,4*R*)-*N*-[(3*R*,5*R*)-1-(3-Chloro-4-fluorobenzoyl)-5-methylpyrrolidin-3-yl]-1-[(2*S*)-2-acetamido-3,3-dimethylbutanoyl]-4-hydroxypyrrolidine-2-carboxamide (Compound 10)

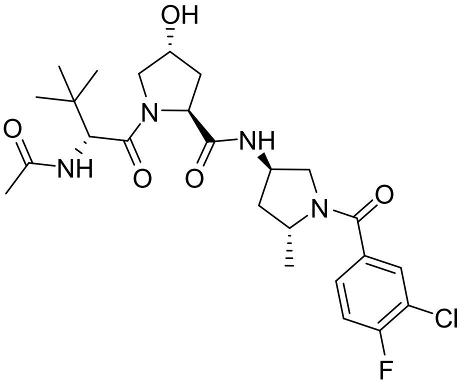


To a suspension of (2*S*,4*R*)-1-[(2*S*)-2-acetamido-3,3-dimethylbutanoyl]-4-hydroxypyrrolidine-2-carboxylic acid (135.30 mg; 0.45 mmol; 1.08 eq.) and (3*R*,5*R*)-1-(3-chloro-4-fluorobenzoyl)-5-methylpyrrolidin-3-amine hydrochloride (136.30 mg; 0.41 mmol; 1.00 eq.) in anhydrous DMF (4 mL) was added DIPEA (368 µL; 2.07 mmol; 5.00 eq.) and the mixture stirred at 25°C for 10 min. The mixture was cooled to 0°C. Next, HATU (175.30 mg; 0.46 mmol; 1.11 eq.) was added at 0 °C and the reaction mixture stirred at 0 °C for 5 min. Then, the reaction was allowed to warm to 25°C and stirred at 25°C for 18 h. After 18 h, the mixture was diluted with DCM (20 mL) and water (20 mL) and the phases separated. The aqueous phase was extracted with DCM (2x 20 mL). The combined organic phase was washed with brine (1x100 mL), dried over anhydrous sodium sulfate, filtered and concentrated in vacuo at 40 °C to provide the crude material as an orange oil (309.5 mg). The crude material was purified by flash column chromatography using a gradient of 0-10% MeOH (B) in DCM (A) (LC time program %B: 0.0 min -0%, 5.0 min-0%, 20.0 min-10%, 26.0 min-10%, 35.0 min-15%; flow rate: 30 mL/min). Product containing fractions were combined and concentrated in vacuo at 40°C to provide the product as a colorless solid in a purity of 99% (LCMS at 220 nm) and a yield of 68% (147.1 mg; 0.28 mmol).

**LCMS:** Rt: 1.30 min; $[M+H^+]$: 525.3 (Method A)

**$^1$H NMR (400 MHz, DMSO-d$_6$)** δ [ppm] 8.14 (d, *J* = 6.4 Hz, 1H), 7.69 (d, *J* = 7.2 Hz, 1H), 7.60 (d, *J* = 9.2 Hz, 1H), 7.56 – 7.38 (m, 2H), 5.12 – 5.00 (m, 1H), 4.43 (d, *J* = 9.2 Hz, 1H), 4.38 – 4.20 (m, 3H), 4.19 – 4.12 (m, 1H), 3.82 – 3.71 (m, 1H), 3.68 – 3.49 (m, 2H), 3.19 – 3.09 (m, 1H), 2.09 – 1.89 (m, 2H), 1.86 (s, 3H), 1.84 – 1.71 (m, 2H), 1.32 – 1.21 (m, 3H), 0.81 (s, 9H).

Note: Residual DCM 3% by weight in NMR spectrum observed.

#### (2*S*,4*R*)-*N*-{[(3*S*)-1-(3,5-Difluorobenzoyl)pyrrolidin-3-yl]methyl}-1-[(2*S*)-2-acetamido-3,3-dimethylbutanoyl]-4-hydroxypyrrolidine-2-carboxamide (Compound 11)

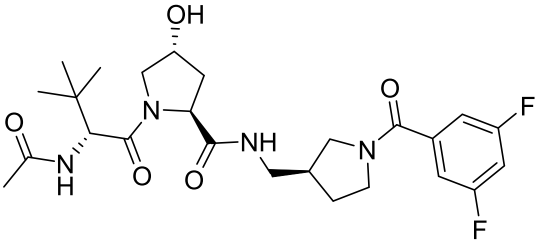


To a suspension of (2*S*,4*R*)-1-[(2*S*)-2-acetamido-3,3-dimethylbutanoyl]-4-hydroxypyrrolidine-2-carboxylic acid (179.10 mg; 0.59 mmol; 0.97 eq.) and 1-[(3*S*)-1-(3,5-difluorobenzoyl)pyrrolidin-3-yl]methanamine hydrochloride (252.00 mg; 0.61 mmol; 1.00 eq.) in anhydrous DMF (4 mL) was added DIPEA (546 µL; 3.07 mmol; 5.00 eq.) and the mixture stirred at 25°C for 10 min. The mixture was cooled to 0°C. Next, HATU (233.10 mg; 0.61 mmol; 1.00 eq.) was added at 0 °C and the reaction mixture stirred at 0 °C for 5 min. Then, the reaction was allowed to warm to 25°C and stirred at 25°C for 18 h. After 18 h, the mixture was diluted with DCM (20 mL) and water (20 mL) and the phases separated. The aqueous phase was extracted with DCM (2x 20 mL). The combined organic phase was washed with brine (1x100 mL), dried over anhydrous sodium sulfate, filtered and concentrated in vacuo at 40 °C to provide the crude material as an orange oil (374.1 mg). The crude material was purified by flash column chromatography using a gradient of 0-10% MeOH (B) in DCM (A) (LC time program %B: 0.0 min -0%, 5.0 min-0%, 25.0 min-10%, 35.0 min-10%.; flow rate: 30 mL/min). Product containing fractions were combined and concentrated in vacuo at 40°C to provide a colorless solid. The product was repurified via reverse phase preparative HPLC using a linear gradient from 0-60% MeCN+0.1% FA (B) in water+0.1% FA (A) (LC-time-%B: 0.0 min -0%, 2.0 min-0%, 19.0 min-60%, 21.3 min-60%, 21.3 min-100%, 23.0 min-100%, 23.0 min-50%, 25.0 min-50%; flow rate: 50 mL/min). Product containing fractions were combined and concentrated under reduced pressure at 50°C. The product (2S,4R)-N-{[(3S)-1-(3,5-difluorobenzoyl)pyrrolidin-3-yl]methyl}-1-[(2S)-2-acetamido-3,3-dimethylbutanoyl]-4-hydroxypyrrolidine-2-carboxamide (115.60 mg; 0.22 mmol) was obtained as a colorless solid in a yield of 37% and a purity of 98% (LCMS at 220 nm).

**LCMS:** Rt: 1.15 min; [M+H$^+$]: 509.3 (Method A).

**$^1$H NMR (400 MHz, DMSO-d$_6$) δ [ppm] (mixture of 2 species major+minor)** 8.13 – 7.95 (m, 1H, major+minor), 7.91 – 7.77 (m, 1H, major+minor), 7.41 – 7.29 (m, 1H, major+minor), 7.26 – 7.12 (m, 2H, major+minor), 5.14 – 5.00 (m, 1H, major+minor), 4.55 – 4.42 (m, 1H, major+minor), 4.37 – 4.19 (m, 2H, major+minor), 3.69 – 2.92 (m, 8H, major+minor), 2.44 – 2.21 (m, 1H, major+minor), 2.03 – 1.57 (m, 2H, major+minor), 1.88 – 1.85 (m, 3H, major+minor), 1.84 – 1.57 (m, 2H, major+minor), 0.95 – 0.87 (m, 9H, major+minor).

Note: Variable temperature $^1$H NMR spectra of the product were recorded at 298 K, 320 K, and 363 K. Coalescence of signals at higher temperature suggests the product to exist in solution as a mixture of rotamers. Residual DCM (10% by weight) was observed in the $^1$H NMR spectrum.

#### Preparation of products modified on the left-hand side of the Hyp-OH core

##### (2*S*,4*R*)-4-Hydroxy-1-[(2*S*)-2-(2-hydroxyacetamido)-3,3-dimethylbutanoyl]-*N*-{[4-(4-methyl-1,3-thiazol-5-yl)phenyl]methyl}pyrrolidine-2-carboxamide (Compound 3)

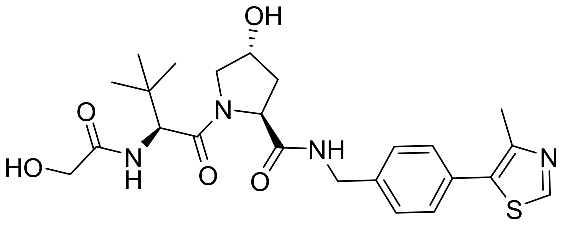


To a suspension of (2*S*,4*R*)-1-((*S*)-2-amino-3,3-dimethylbutanoyl)-4-hydroxy-*N*-(4-(4-methylthiazol-5-yl)benzyl)pyrrolidine-2-carboxamide hydrochloride (109.80 mg; 0.23 mmol; 1.00 eq) and glycolic acid (19.80 mg; 0.26 mmol; 1.13 eq.) in anhydrous DMF (2 mL) was added DIPEA (112.00 µL; 0.63 mmol; 2.73 eq.), and the mixture cooled to 0 °C. Next, HATU (81.10 mg; 0.21 mmol; 0.93 eq.) was added at 0 °C and the reaction mixture stirred at 0 °C for 10 min. Then, the reaction was allowed to warm to 25°C and stirred at 25°C for 5 h. After 5 h, the mixture was diluted with EtOAc (20 mL) and water (20 mL) and the phases separated. The aqueous phase was extracted with EtOAc (2x 20 mL). The combined organic phase was washed with brine (2x80 mL), dried over anhydrous sodium sulfate, filtered and concentrated in vacuo at 50 °C to provide the crude material as a heterogenous mixture of pale yellow oil with solid material (108.1 mg). The crude material was purified by flash column chromatography using a gradient of 0-15% MeOH (B) in DCM (A) (LC time program %B: 0.0 min-0%; 1.0 min-0%; 14.0 min-10%; 20.0 min-10%; 25.0 min-15%; 30.0 min-15%; flow rate: 30 mL/min). Product containing fractions were combined and concentrated in vacuo at 50°C to provide the product as a colorless solid in a purity of 95% (LCMS at 220 nm) and a yield of 42% (49.2 mg; 0.10 mmol).

**LCMS**: Rt: 1.14 min; [M+H$^+$]: 489.2 (Method A).

**1H NMR (400 MHz, DMSO-d$_6$)** δ [ppm] 8.98 (s, 1H), 8.60 (t, J = 6.0 Hz, 1H), 7.49 – 7.41 (m, 1H), 7.40 (s, 4H), 5.69 (t, J = 5.8 Hz, 1H), 5.15 (d, J = 3.5 Hz, 1H), 4.56 (d, J = 9.6 Hz, 1H), 4.47 – 4.37 (m, 2H), 4.37 – 4.33 (m, 1H), 4.25 (dd, J = 15.8, 5.6 Hz, 1H), 3.90 (dd, J = 16.0, 5.7 Hz, 1H), 3.81 (dd, J = 16.0, 5.9 Hz, 1H), 3.67 (dd, J = 10.7, 3.9 Hz, 1H), 3.60 (d, J = 10.7 Hz, 1H), 2.45 (s, 3H), 2.11 – 2.02 (m, 1H), 1.90 (ddd, J = 13.0, 8.9, 4.4 Hz, 1H), 0.94 (s, 9H).

Note: ~7% (by weight) DCM observed in $^1$H NMR spectrum.

##### (2*S*,4*R*)-4-Hydroxy-1-[(2*S*)-2-(2-methoxyacetamido)-3,3-dimethylbutanoyl]-*N*-{[4-(4-methyl-1,3-thiazol-5-yl)phenyl]methyl}pyrrolidine-2-carboxamide (Compound 4)

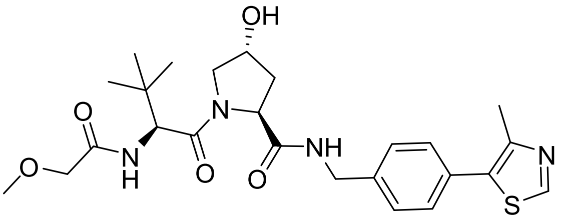


To a suspension of (2*S*,4*R*)-1-((*S*)-2-amino-3,3-dimethylbutanoyl)-4-hydroxy-*N*-(4-(4-methylthiazol-5-yl)benzyl)pyrrolidine-2-carboxamide hydrochloride (102.20 mg; 0.21 mmol; 1.00 eq.) in anhydrous DMF (2 mL) was added methoxyacetic acid, 98% (16.00 µL; 0.21

mmol; 0.97 eq.), and DIPEA (112.00 µL; 0.63 mmol; 2.94 eq.). The mixture was cooled to 0°C before addition of HATU (74.80 mg; 0.20 mmol; 0.92 eq.) and the reaction mixture stirred at 0 °C for 15 min. Then, the reaction was allowed to warm to 25°C and stirred at 25°C for 23 h. After 23 h, the mixture was diluted with EtOAc (20 mL) and water (20 mL), and the phases separated. The aqueous phase was extracted with EtOAc (2x 20 mL). The combined organic phase was washed with brine (2x80 mL), dried over anhydrous sodium sulfate, filtered and concentrated in vacuo at 50 °C to provide the crude material as a colorless film (109 mg). The crude material was purified by flash column chromatography using a gradient of 0-15% MeOH (B) in DCM (A) (LC time program %B: 0.0 min-0%; 1.0 min-0%; 14.0 min-10%; 20.0 min-10%; 25.0 min-15%; 29.0 min-15%; flow rate: 30 mL/min). Product containing fractions were combined and concentrated in vacuo at 50°C to provide the product as a colorless solid in a purity of approx. 92% (assigned based on $^1$H NMR analysis) and a yield of 58% (67.60 mg; 0.12 mmol).

**LCMS:** Rt: 1.26 min; [M+H$^+$]: 503.2 (Method A)

**$^1$H NMR (400 MHz, DMSO-d$_6$)** δ [ppm] 8.98 (s, 1H), 8.60 (t, J = 6.0 Hz, 1H), 7.40 (s, 4H), 7.37 (d, J = 9.6 Hz, 1H), 5.15 (d, J = 3.5 Hz, 1H), 4.57 (d, J = 9.5 Hz, 1H), 4.48 – 4.37 (m, 2H), 4.36 – 4.32 (m, 1H), 4.25 (dd, J = 15.8, 5.6 Hz, 1H), 3.95 – 3.82 (m, 2H), 3.66 (dd, J = 10.7, 3.9 Hz, 1H), 3.60 (d, J = 10.7 Hz, 1H), 3.33 (s, 3H), 2.44 (s, 3H), 2.10 – 2.01 (m, 1H), 1.90 (ddd, J = 13.1, 8.9, 4.5 Hz, 1H), 0.94 (s, 9H).

Note: approx. 8% unassigned impurity (peaks of a second species) observed in $^1$H NMR spectrum); Residual DCM (6% by weight) observed in $^1$H NMR spectrum.

#### (2*S*,4*R*)-1-[(2*S*)-2-(2-Butoxyacetamido)-3,3-dimethylbutanoyl]-4-hydroxy-*N*-{[4-(4-methyl-1,3-thiazol-5-yl)phenyl]methyl}pyrrolidine-2-carboxamide (Compound 5)

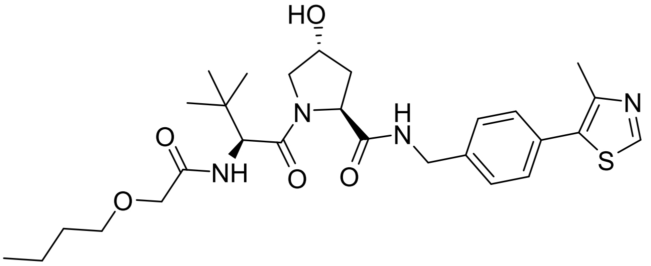


To a suspension of (2*S*,4*R*)-1-((*S*)-2-amino-3,3-dimethylbutanoyl)-4-hydroxy-*N*-(4-(4-methylthiazol-5-yl)benzyl)pyrrolidine-2-carboxamide hydrochloride (101.90 mg; 0.21 mmol; 1.00 eq.) in anhydrous DMF (2 mL) was added 2-butoxyacetic acid (28.60 µL; 0.21 mmol; 0.98 eq.), and DIPEA (224 µL; 1.26 mmol; 5.89 eq.). The mixture was cooled to 0°C followed by addition of HATU (78.10 mg; 0.21 mmol; 0.96 eq.) at 0 °C. The reaction mixture was stirred at 0 °C for 10 min, before it was allowed to warm to 25°C, and stirred at 25°C for 21 h. After 21 h, the mixture was diluted with EtOAc (20 mL) and water (20 mL) and the phases separated. The aqueous phase was extracted with EtOAc (2x 20 mL). The combined organic phase was washed with brine (2x80 mL), dried over anhydrous sodium sulfate, filtered and concentrated in vacuo at 50 °C to provide the crude material as a colorless to pale yellow oil (200.4 mg). The crude material was purified by flash column chromatography using a gradient of 0-10% MeOH (B) in DCM (A) (LC time program %B: 0.0 min-0%; 1.0 min-0%;

15.0 min-10%; 25.0 min-10%; flow rate: 30 mL/min). Product containing fractions were combined and concentrated in vacuo at 35°C. The product was obtained as a colorless solid in a purity of 91% (based on $^{1}$H NMR analysis) and a yield of 69% (87.80 mg; 0.15 mmol).

**LCMS:** Rt: 1.55 min; [M+H$^{+}$]: 545.3 (Method A).

**$^{1}$H NMR (400 MHz, DMSO-d$_6$)** δ [ppm] 8.98 (s, 1H), 8.62 (t, *J* = 6.0 Hz, 1H), 7.46 – 7.31 (m, 5H), 5.15 (d, *J* = 3.5 Hz, 1H), 4.55 (d, *J* = 9.6 Hz, 1H), 4.49 – 4.37 (m, 2H), 4.37 – 4.31 (m, 1H), 4.25 (dd, *J* = 15.8, 5.6 Hz, 1H), 3.91 (d, *J* = 1.2 Hz, 2H), 3.66 (dd, *J* = 10.7, 3.8 Hz, 1H), 3.60 (d, *J* = 10.7 Hz, 1H), 3.47 (td, *J* = 6.4, 0.9 Hz, 2H), 2.44 (s, 3H), 2.07 (ddd, *J* = 12.8, 5.4, 3.8 Hz, 1H), 1.90 (ddd, *J* = 13.0, 9.0, 4.5 Hz, 1H), 1.58 – 1.46 (m, 2H), 1.42 – 1.30 (m, 2H), 0.94 (s, 9H), 0.89 (t, *J* = 7.4 Hz, 3H).

#### (2*S*,4*R*)-1-[(2*S*)-2-[2-(3-Cyanophenoxy)acetamido]-3,3-dimethylbutanoyl]-4-hydroxy-*N*-{[4-(4-methyl-1,3-thiazol-5-yl)phenyl]methyl}pyrrolidine-2-carboxamide (Compound 6)

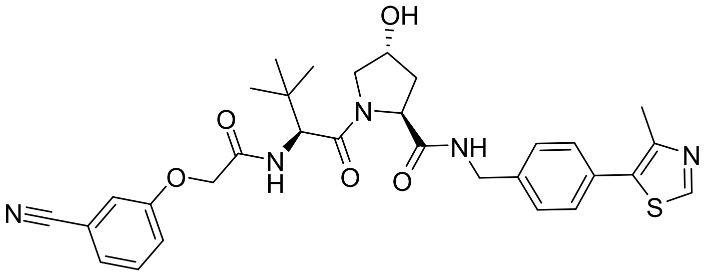


To a suspension of (2*S*,4*R*)-1-((*S*)-2-amino-3,3-dimethylbutanoyl)-4-hydroxy-*N*-(4-(4-methylthiazol-5-yl)benzyl)pyrrolidine-2-carboxamide hydrochloride (107.00 mg; 0.22 mmol; 1.00 eq.) and (3-cyanophenoxy)acetic acid (36.80 mg; 0.21 mmol; 0.93 eq.) in anhydrous DMF (2 mL) was added DIPEA (224 µL; 1.26 mmol; 5.61 eq.). The mixture was cooled to 0°C. Next, HATU (83.20 mg; 0.22 mmol; 0.97 eq.) was added at 0°C and the reaction mixture stirred at 0 °C for 10 min. Then, the reaction was allowed to warm to 25°C and stirred at 25°C for 3.5 h. After 3.5 h, the mixture was diluted with EtOAc (20 mL) and water (20 mL) and the phases separated. The aqueous phase was extracted with EtOAc (2x 20 mL). The combined organic phase was washed with brine (2x80 mL), dried over anhydrous sodium sulfate, filtered and concentrated in vacuo at 50 °C to provide the crude material as a heterogenous mixture of pale yellow film with solid material (146.3 mg). The crude material was purified by flash column chromatography using a gradient of 0-10% MeOH (B) in DCM (A) (LC time program %B: 0.0 min-0%; 1.0 min-0%; 15.0 min-10%; 25.0 min-10%; Flow rate: 30 mL/min). Product containing fractions were combined and concentrated in vacuo at 50°C to provide the product as a colorless solid in a purity of >99% and a yield of 82% (108.20 mg; 0.18 mmol).

**LCMS:** Rt: 1.483 min; [M+H$^{+}$]: 590.2 (Method A)

**$^{1}$H NMR (400 MHz, DMSO-d$_6$)** δ [ppm] 8.99 (s, 1H), 8.59 (t, J = 6.1 Hz, 1H), 8.02 (d, J = 9.3 Hz, 1H), 7.53 – 7.46 (m, 1H), 7.46 – 7.37 (m, 6H), 7.33 – 7.27 (m, 1H), 5.14 (d, J = 3.5 Hz, 1H), 4.83 – 4.67 (m, 2H), 4.58 (d, J = 9.4 Hz, 1H), 4.50 – 4.38 (m, 2H), 4.35 (s, 1H), 4.24 (dd, J = 15.8, 5.5 Hz, 1H), 3.67 (dd, J = 10.6, 4.0 Hz, 1H), 3.60 (d, J = 10.7 Hz, 1H), 2.45 (s, 3H), 2.10 – 2.01 (m, 1H), 1.95 – 1.85 (m, 1H), 0.94 (s, 9H).

Note: residual amounts of DCM (1% by weight) observed in $^{1}$H NMR spectrum. Also, aromatic multiplet 7.46 – 7.37 (m, 6H) integrates too high (6.9 should correspond to 6 Hs).

### Appendix

#### NMR spectra of intermediates

[1]H NMR (700 MHz, DMSO) δ 13.34 (s, 1H), 8.94 (dt, $J$ = 8.6, 1.0 Hz, 1H), 8.21 – 8.15 (m, 2H), 7.98 (dd, $J$ = 2.2, 1.0 Hz, 1H), 7.87 (d, $J$ = 7.5 Hz, 1H), 7.83 (ddd, $J$ = 8.2, 6.8, 1.2 Hz, 1H), 7.74 (ddd, $J$ = 8.3, 6.8, 1.3 Hz, 1H), 6.83 – 6.78 (m, 1H).

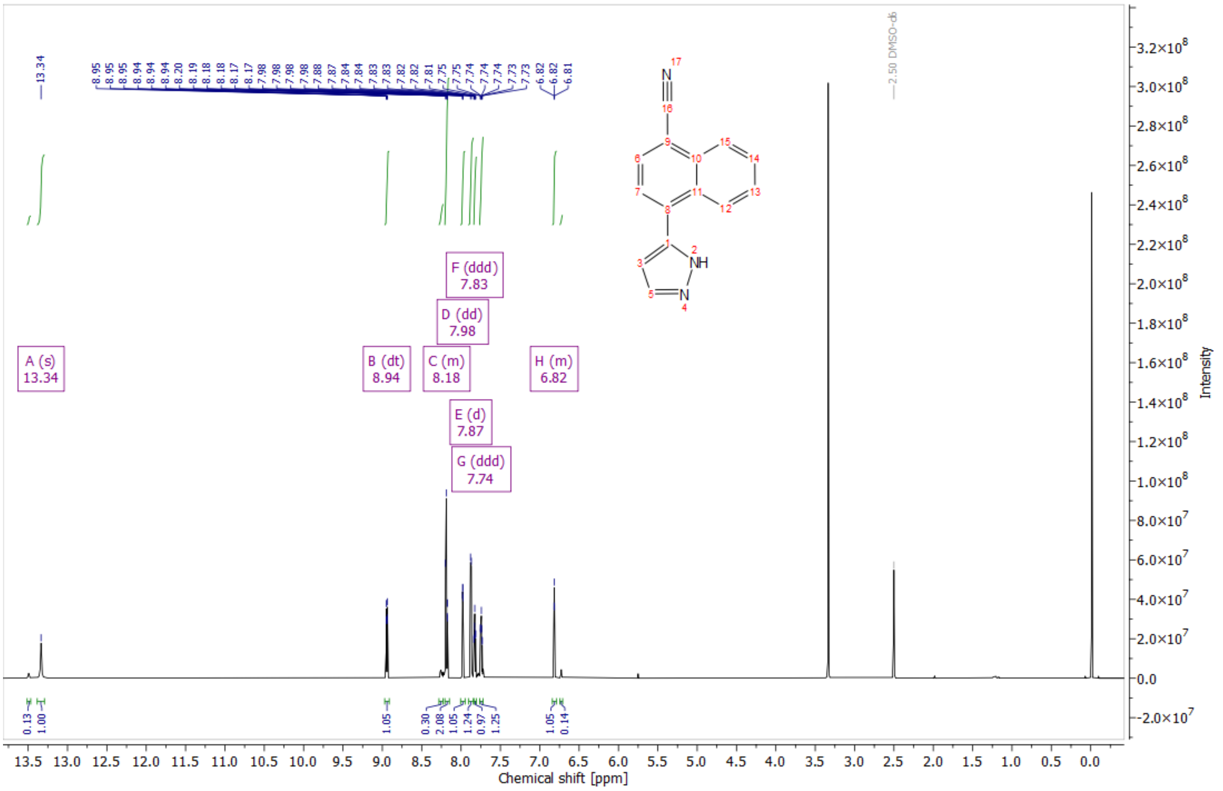


*Figure 1: [1]H NMR spectrum in DMSO-$d_6$ of 4-(1H-pyrazol-5-yl)naphthalene-1-carbonitrile (Int-2a)*

[1]H NMR (400 MHz, DMSO) (major:minor approx. 1:0.7) δ 10.50 – 10.41 (m, 1.7 H, major+minor), 8.21 – 8.14 (m, 1.7 H, major+minor), 7.82 – 7.72 (m, 1.7H, major+minor), 7.57 – 7.42 (m, 3.4 H, major+minor), 7.33 (d, $J$ = 7.8 Hz, 0.7 H, minor), 7.36 – 7.24 (m, 1.7 H, major+minor), 7.16 (d, $J$ = 6.2 Hz, 1 H, major), 6.90 – 6.83 (m, 1.7 H, major+minor), 4.13 – 4.01 (m, 0.7 H, minor), 3.91 – 3.81 (m, 1H, major), 3.80 – 3.65 (m, 1.7 H, major+minor), 3.62 – 3.49 (m, 1H, major), 3.45 – 3.35 (m, 0.7 H, minor), 3.28 – 3.19 (m, 1.7 H, major+minor), 3.17 (d, $J$ = 5.1 Hz, 0.7 H, minor), 3.14 – 3.04 (m, 0.7 H, minor), 2.96 – 2.84 (m, 1 H, major), 2.14 – 2.02 (m, 1 H, major), 2.01 – 1.89 (m, 0.7 H, minor), 1.89 – 1.77 (m, 1H, major), 1.74 – 1.62 (m, 0.7 H, minor), 1.41 (s, 6.3 H, minor), 1.28 (s, 9 H, major).

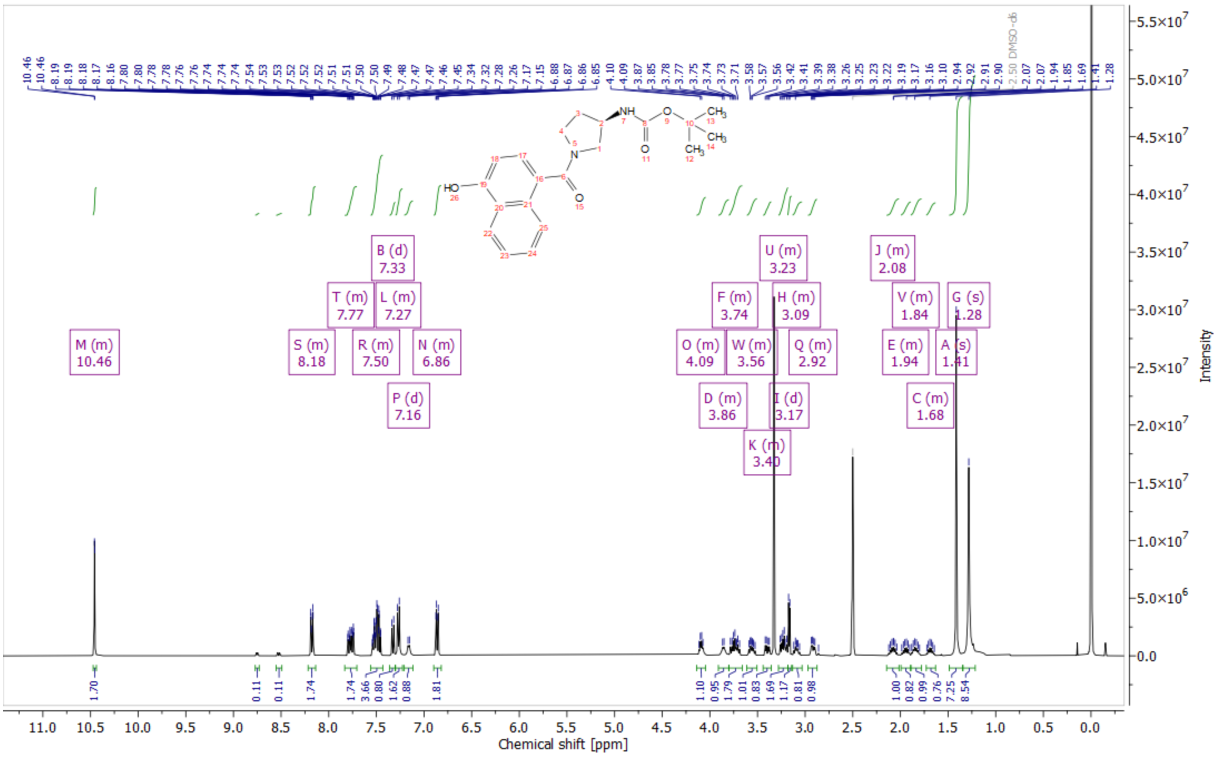


*Figure 2: [1]H NMR spectrum in DMSO-d$_6$ of tert-butyl N-[(3R)-1-(4-hydroxynaphthalene-1-carbonyl)pyrrolidin-3-yl]carbamate (Int-8a)*

[1]H NMR (400 MHz, DMSO) δ 10.61 (s, 1H), 8.44 (s, 1H), 8.25 (s, 1H), 8.22 – 8.15 (m, 1H), 7.87 – 7.80 (m, 1H), 7.57 – 7.44 (m, 2H), 7.44 – 7.28 (m, 1H), 6.97 – 6.87 (m, 1H), 3.97 – 3.59 (m, 3H), 3.46 – 3.34 (m, 1H), 3.24 – 3.10 (m, 1H), 2.37 – 1.86 (m, 2H).

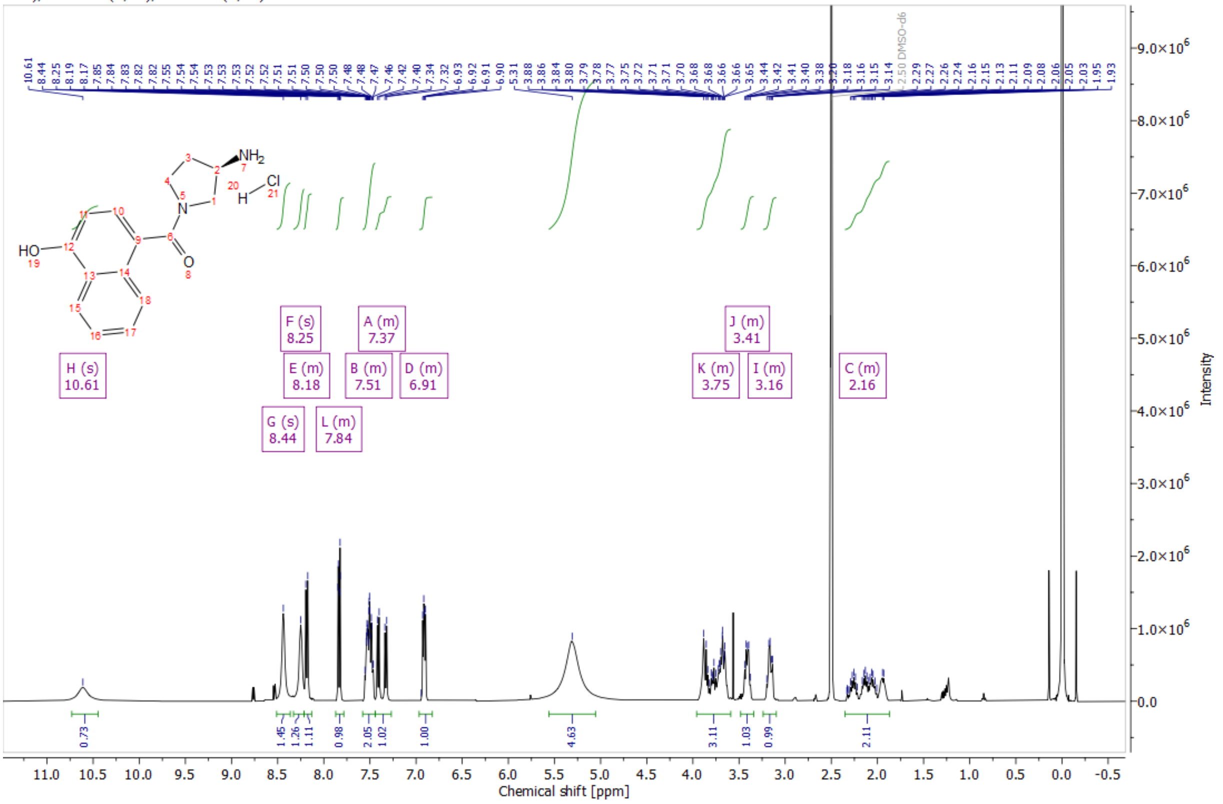


*Figure 3: [1]H NMR spectrum in DMSO-d$_6$ of crude HCl salt 4-[(3R)-3-aminopyrrolidine-1-carbonyl]naphthalen-1-ol hydrochloride (Int-8b)*

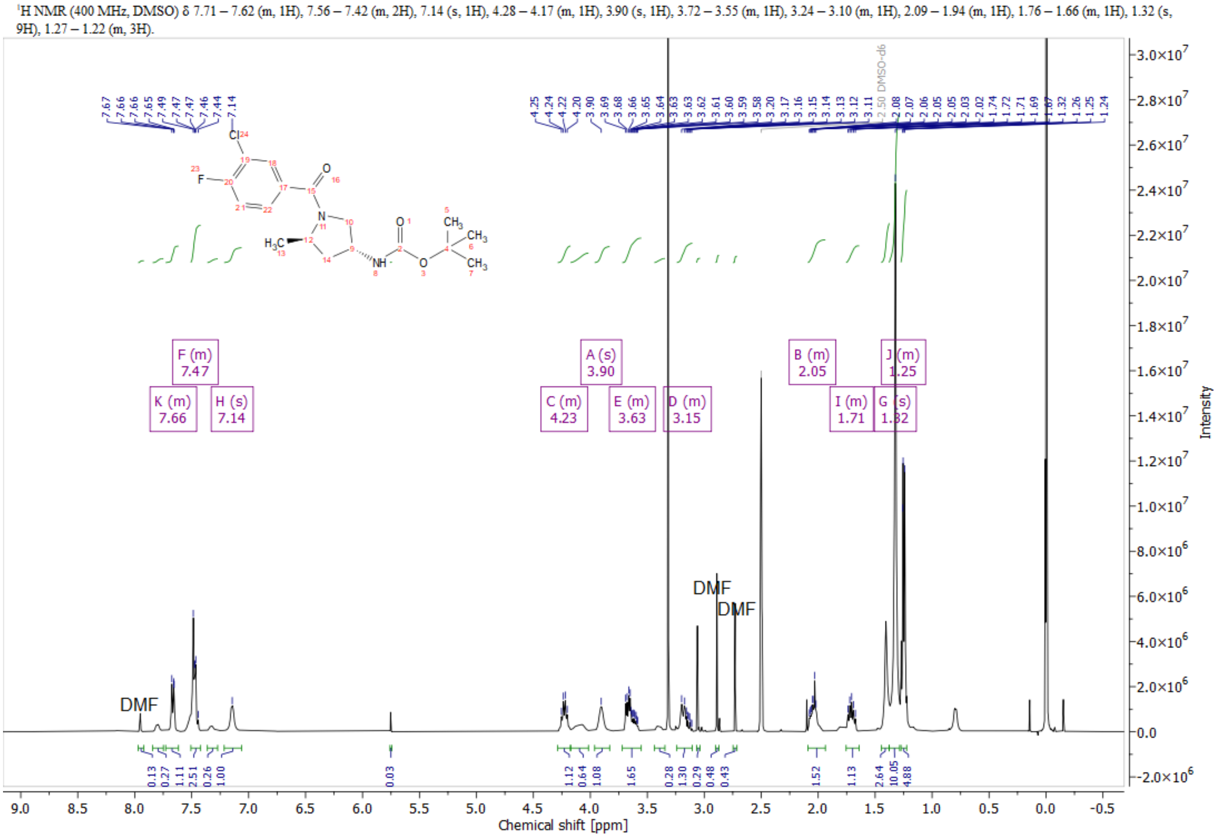


*Figure 4: [1]H NMR spectrum in DMSO-$d_6$ of tert-butyl N-[(3R,5R)-1-(3-chloro-4-fluorobenzoyl)-5-methylpyrrolidin-3-yl]carbamate (Int-10a)*

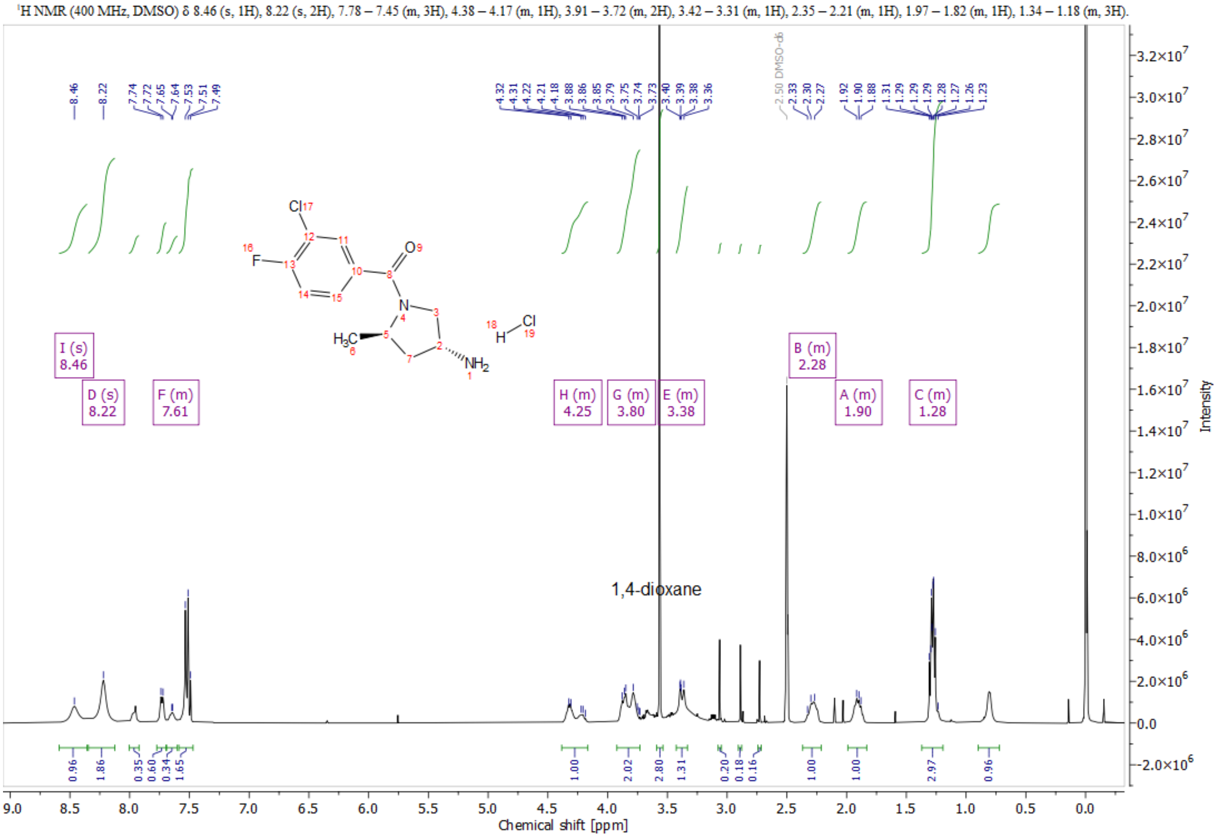


*Figure 5: [1]H NMR spectrum in DMSO-$d_6$ of (3R,5R)-1-(3-chloro-4-fluorobenzoyl)-5-methylpyrrolidin-3-amine hydrochloride (Int-10b)*

$^{1}H$ NMR (400 MHz, DMSO-$d_6$) (minor: major = 1.0:0.8) δ (ppm) 7.40 – 7.17 (m, 5.4H, major+minor), 7.03 (t, J = 5.8 Hz, 0.8H, minor), 6.92 (t, J = 5.9 Hz, 1H, major), 3.59 – 3.35 (m, 5.4H, major+minor), 3.20 – 3.08 (m, 2.6H, major+minor), 3.05 – 2.79 (m, 2.8H, major+minor), 2.38 – 2.18 (m, 1.8H, major+minor), 1.99 – 1.82 (m, 1.8H, major+minor), 1.68 – 1.51 (m, 1.8H, major+minor), 1.39 (s, 7.2H, minor), 1.33 (s, 9H, major).

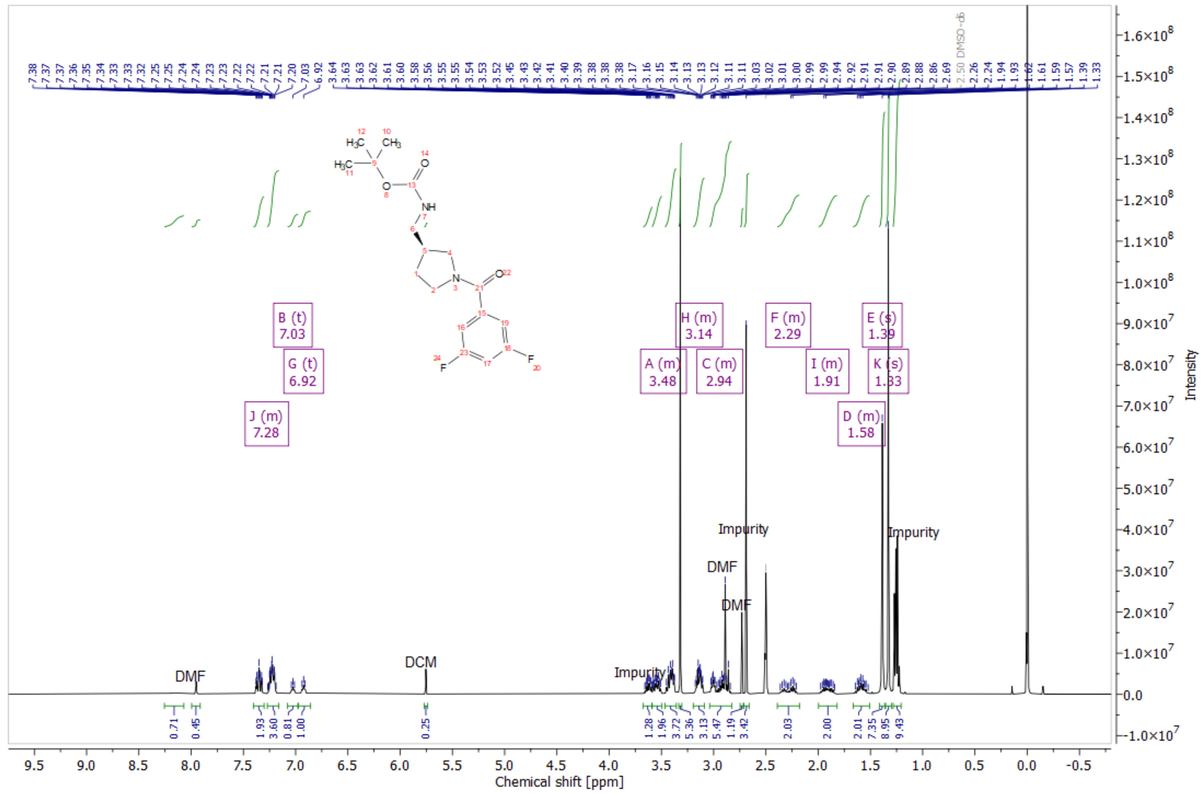


*Figure 6: 1H NMR spectrum in DMSO-d6 of tert-butyl N-{[(3S)-1-(3,5-difluorobenzoyl)pyrrolidin-3-yl]methyl}carbamate (Int-11a)*

### NMR Spectra of products modified at the right-hand side of Hyp-OH

$^1$H NMR (700 MHz, DMSO) δ 13.22 (s, 0H), 13.05 (s, 1H), 8.74 – 8.63 (m, 1H), 8.63 – 8.52 (m, 1H), 8.08 (s, 1H), 7.93 (d, *J* = 9.4 Hz, 1H), 7.90 (s, 1H), 7.71 – 7.43 (m, 4H), 6.65 – 6.49 (m, 1H), 5.11 (d, *J* = 3.7 Hz, 1H), 4.81 (dd, *J* = 15.5, 5.9 Hz, 1H), 4.75 – 4.69 (m, 1H), 4.54 (d, *J* = 9.5 Hz, 1H), 4.46 (t, *J* = 8.0 Hz, 1H), 4.38 – 4.32 (m, 1H), 3.69 (dd, *J* = 10.4, 4.2 Hz, 1H), 3.63 (dt, *J* = 10.5, 1.9 Hz, 1H), 2.04 – 1.98 (m, 1H), 1.95 – 1.89 (m, 1H), 1.88 (s, 3H), 0.94 (s, 9H).

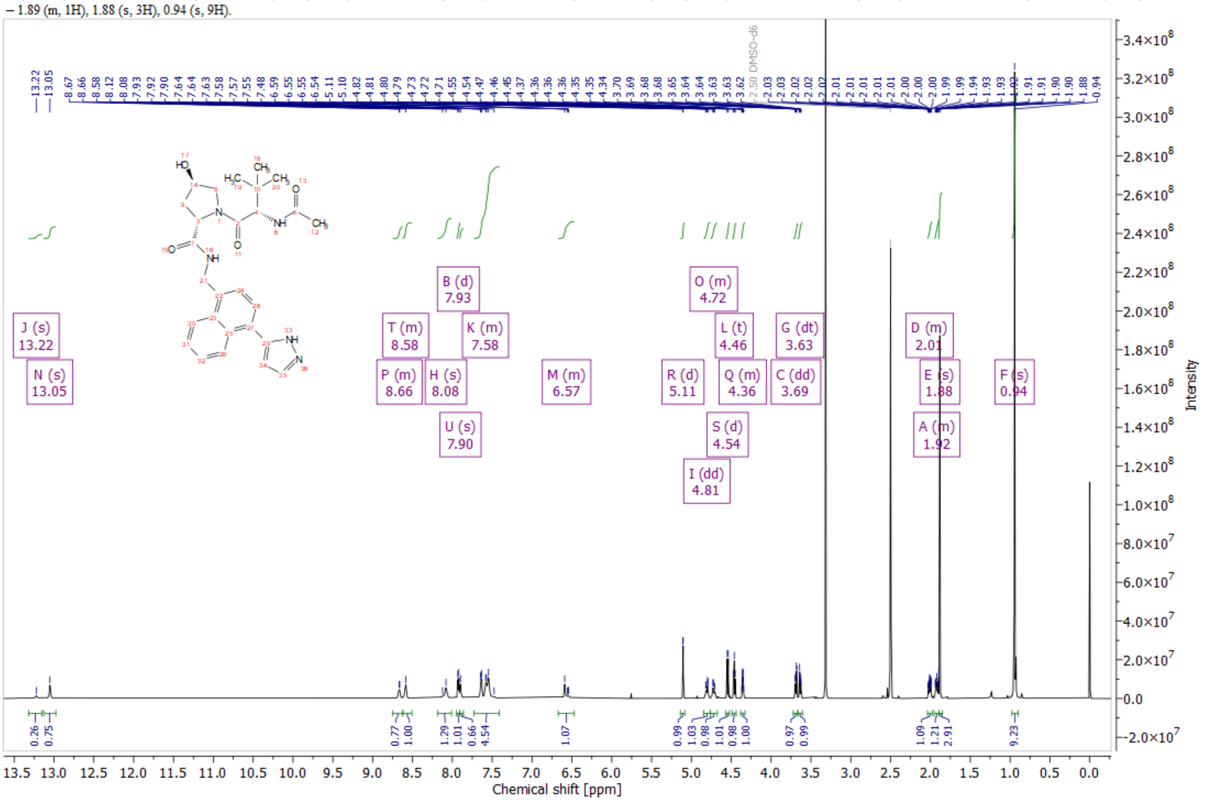


*Figure 7: $^1$H NMR spectrum in DMSO-$d_6$ of (2S,4R)-1-[(2S)-2-acetamido-3,3-dimethylbutanoyl]-4-hydroxy-N-{[4-(1H-pyrazol-5-yl)naphthalen-1-yl]methyl}pyrrolidine-2-carboxamide (Compound 2)*

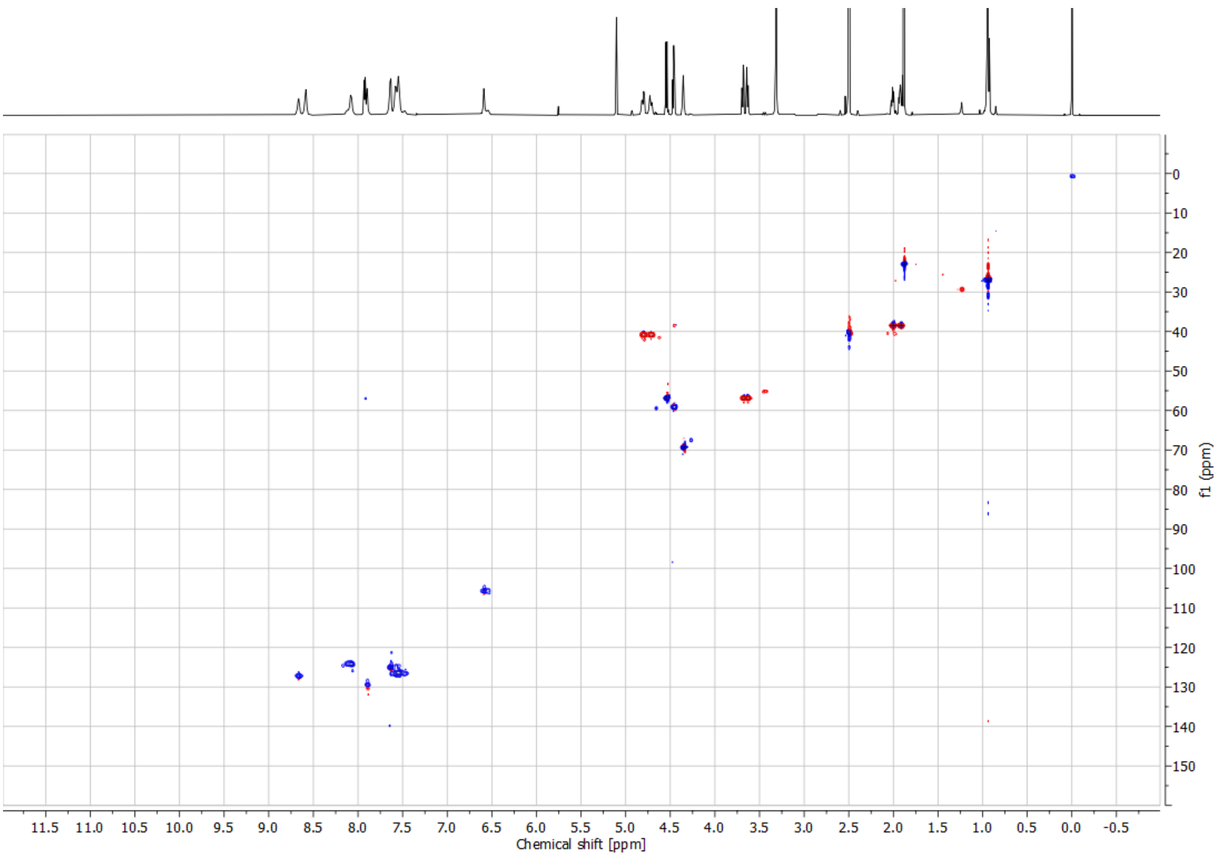

*Figure 8: HSQC NMR spectrum in DMSO-$d_6$ of (2S,4R)-1-[(2S)-2-acetamido-3,3-dimethylbutanoyl]-4-hydroxy-N-{[4-(1H-pyrazol-5-yl)naphthalen-1-yl]methyl}pyrrolidine-2-carboxamide (Compound 2)*

$^1$H NMR (400 MHz, DMSO) δ ppm (mixture of at least 2 specied in a ratio of approx. major:minor = 1.0:0.5 ) 8.08 – 7.94 (m, 1.5H, major+minor), 7.92 – 7.84 (m, 1.5H, major+minor), 7.50 – 7.13 (m, 7.5H, major+minor), 5.14 – 5.05 (m, 1.5H, major+minor), 4.55 – 4.49 (m, 1.5H, major+minor), 4.49 – 4.41 (m, 1.5H, major+minor), 4.33 – 4.23 (m, 1.5H, major+minor), 4.19 – 3.36 (m, 11.5H, major+minor), 3.06 – 2.89 (m, 0.5H, minor), 2.11 – 1.89 (m, 4.5H, major+minor), 1.89 – 1.83 (m, 4H, major+minor), 1.83 – 1.50 (m, 4H, major+minor), 1.44 – 1.31 (m, 1H, major), 1.25 – 1.18 (m, 1.5H, minor), 1.14 – 1.09 (m, 3H, major), 0.93 (s, 9H, major, partially overlaps with minor signal), 0.92 (s, 4.5H, minor, partially overlaps with major signal).

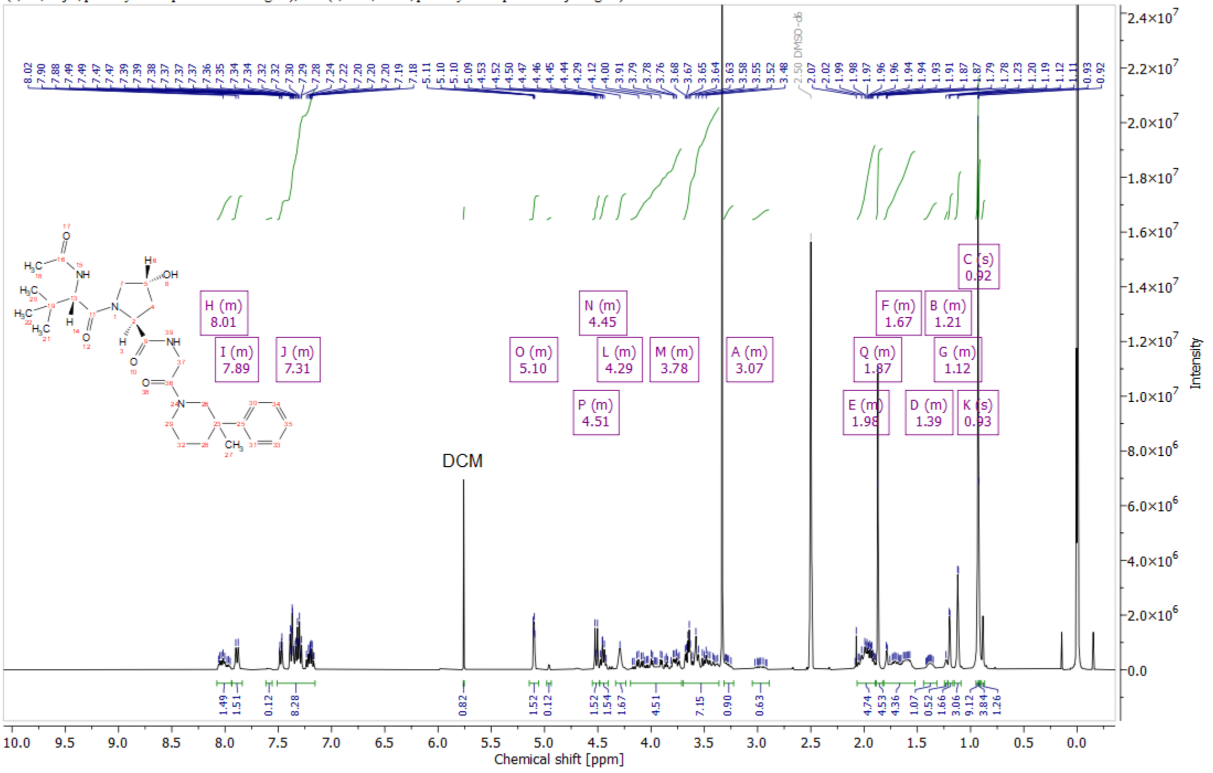

*Figure 9: $^1$H NMR spectrum in DMSO-$d_6$ of (2S,4R)-1-[(2S)-2-acetamido-3,3-dimethylbutanoyl]-4-hydroxy-N-[2-(3-methyl-3-phenylpiperidin-1-yl)-2-oxoethyl]pyrrolidine-2-carboxamide (Compound 7)*

[1]H NMR (400 MHz, DMSO) (mixture of rotamers: major:minor = 1: 1.3 ) δ 10.58 – 10.32 (m, 2.3H, major+minor), 8.26 (d, $J$ = 6.9 Hz, 1H, minor), 8.22 – 8.07 (m, 3.6H, major+minor), 7.93 (d, $J$ = 9.3 Hz, 1H, minor), 7.86 – 7.70 (m, 3.6H, major+minor), 7.59 – 7.39 (m, 4.6H, major+minor), 7.37 – 7.23 (m, 2.3H, major+minor), 6.95 – 6.75 (m, 2.3H, major+minor), 5.17 – 5.02 (m, 2.3H, major+minor), 4.56 – 4.41 (m, 2.3H, major+minor), 4.38 – 4.06 (m, 6.9H, major+minor), 3.84 – 2.91 (m, 13.8H, major+minor), 2.17 – 1.65 (m, 9.2H, major+minor), 1.88 (s, 3H, minor), 1.84 (s, 3.9H, major), 0.96 (s, 9H, minor), 0.81 (s, 11.7H, major).

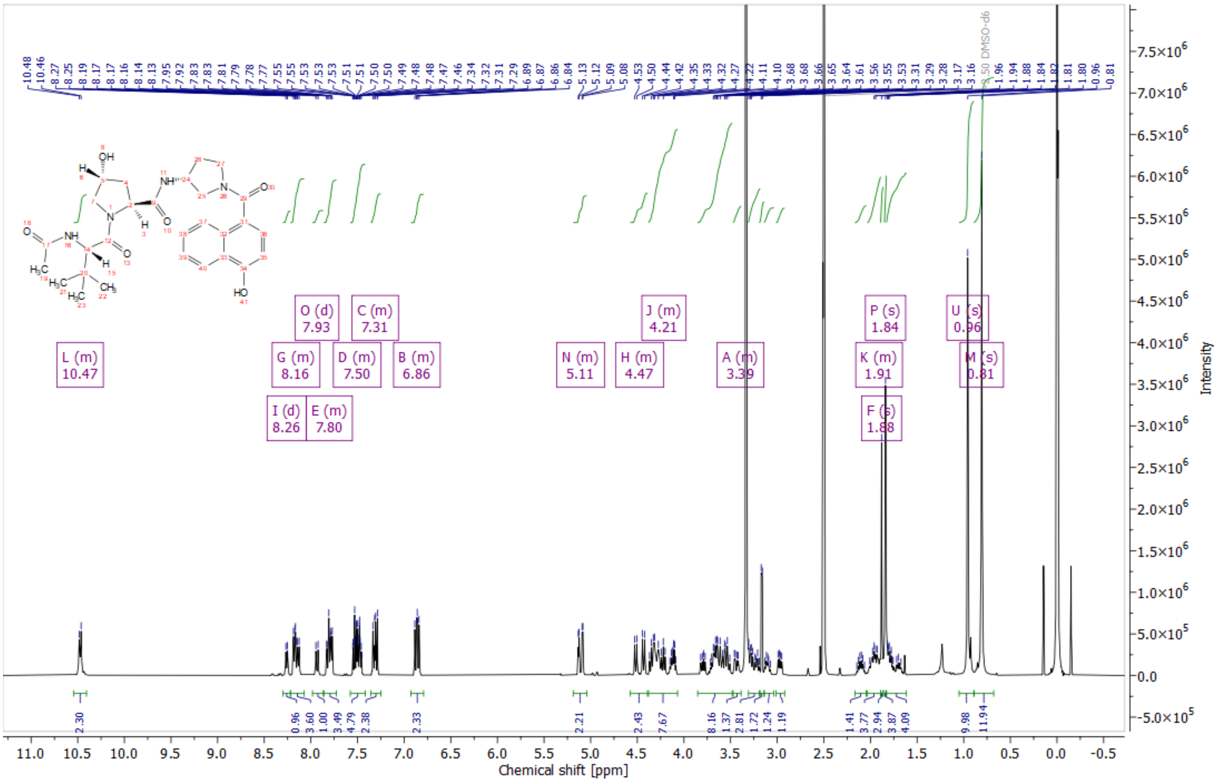


*Figure 10: [1]H NMR spectrum in DMSO-$d_6$ of (2S,4R)-1-[(2S)-2-acetamido-3,3-dimethylbutanoyl]-4-hydroxy-N-[(3R)-1-(4-hydroxynaphthalene-1-carbonyl)pyrrolidin-3-yl]pyrrolidine-2-carboxamide (Compound 8)*

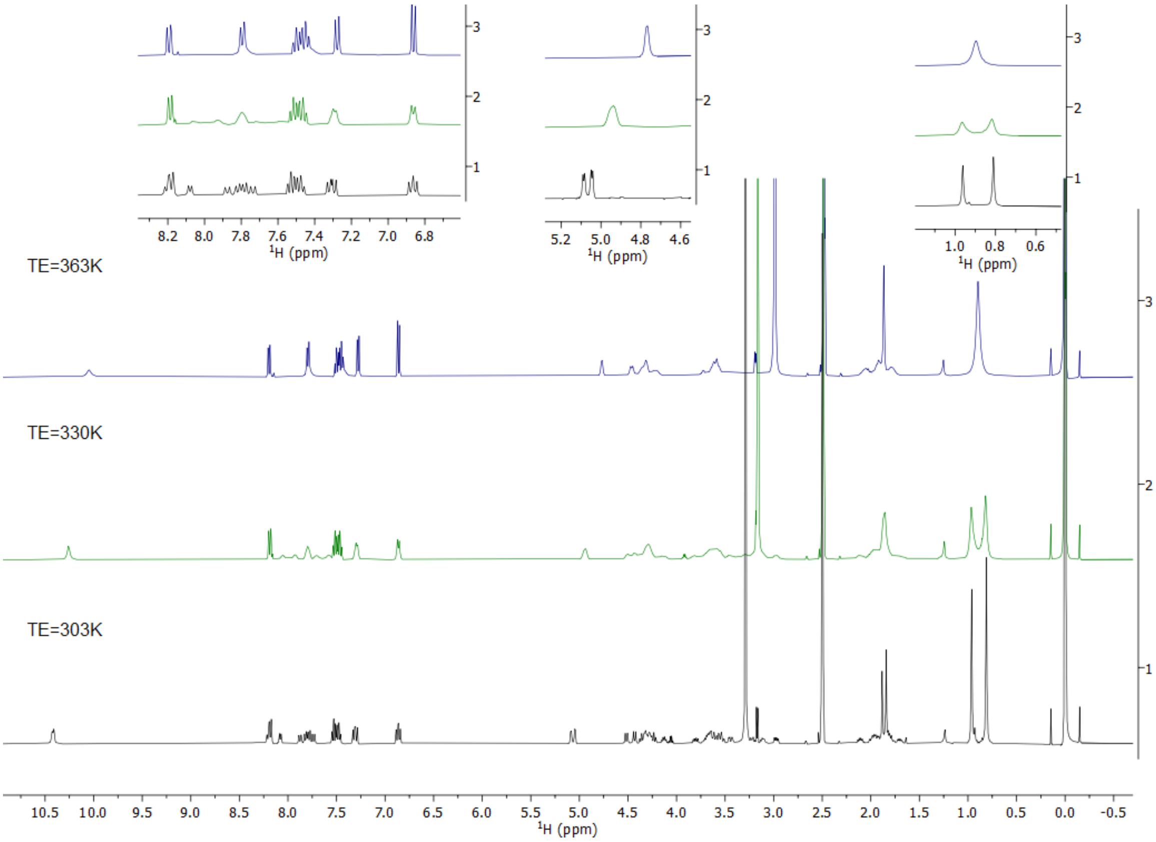


*Figure 11: Stacked [1]H vT NMR spectra in DMSO-$d_6$ at different T (TE) of (2S,4R)-1-[(2S)-2-acetamido-3,3-dimethylbutanoyl]-4-hydroxy-N-[(3R)-1-(4-hydroxynaphthalene-1-carbonyl)pyrrolidin-3-yl]pyrrolidine-2-carboxamide (Compound 8)*

$^1$H NMR (400 MHz, DMSO) δ 8.24 – 8.09 (m, 1H), 7.88 (d, $J$ = 9.9 Hz, 1H), 7.46 – 7.24 (m, 4H), 5.45 – 5.29 (m, 1H), 5.15 – 5.05 (m, 1H), 4.51 (d, $J$ = 9.3 Hz, 1H), 4.48 – 4.11 (m, 4H), 4.12 – 3.99 (m, 1H), 3.93 – 3.38 (m, 6H), 3.27 – 3.09 (m, 1H), 2.06 – 1.89 (m, 2H), 1.89 – 1.83 (m, 3H), 0.93 (s, 9H).

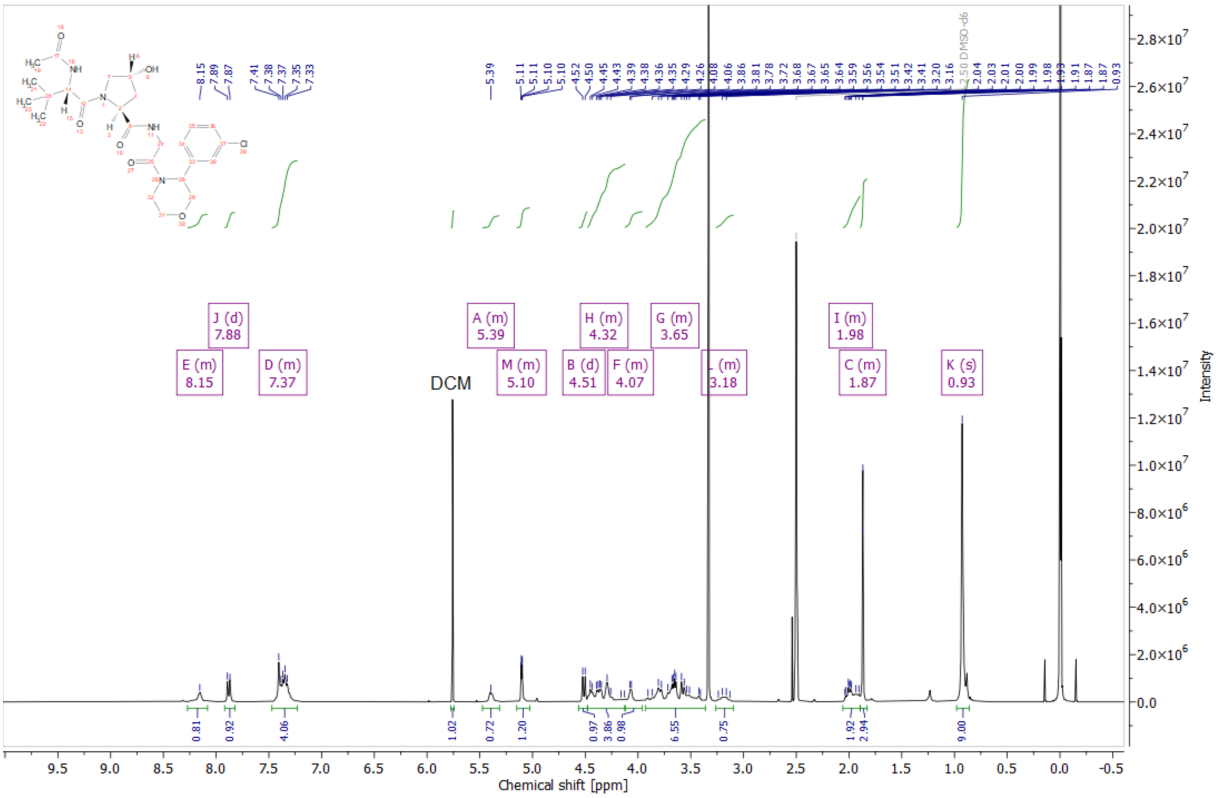


*Figure 12: $^1$H NMR spectrum in DMSO-$d_6$ of (2S,4R)-N-{2-[3-(3-Chlorophenyl)morpholin-4-yl]-2-oxoethyl}-1-[(2S)-2-acetamido-3,3-dimethylbutanoyl]-4-hydroxypyrrolidine-2-carboxamide (Compound 9)*

$^1$H NMR (400 MHz, DMSO) δ 8.14 (d, $J$ = 6.4 Hz, 1H), 7.69 (d, $J$ = 7.2 Hz, 1H), 7.60 (d, $J$ = 9.2 Hz, 1H), 7.56 – 7.38 (m, 2H), 5.12 – 5.00 (m, 1H), 4.43 (d, $J$ = 9.2 Hz, 1H), 4.38 – 4.20 (m, 3H), 4.19 – 4.12 (m, 1H), 3.82 – 3.71 (m, 1H), 3.68 – 3.49 (m, 2H), 3.19 – 3.09 (m, 1H), 2.09 – 1.89 (m, 2H), 1.86 (s, 3H), 1.84 – 1.71 (m, 2H), 1.32 – 1.21 (m, 3H), 0.81 (s, 9H).

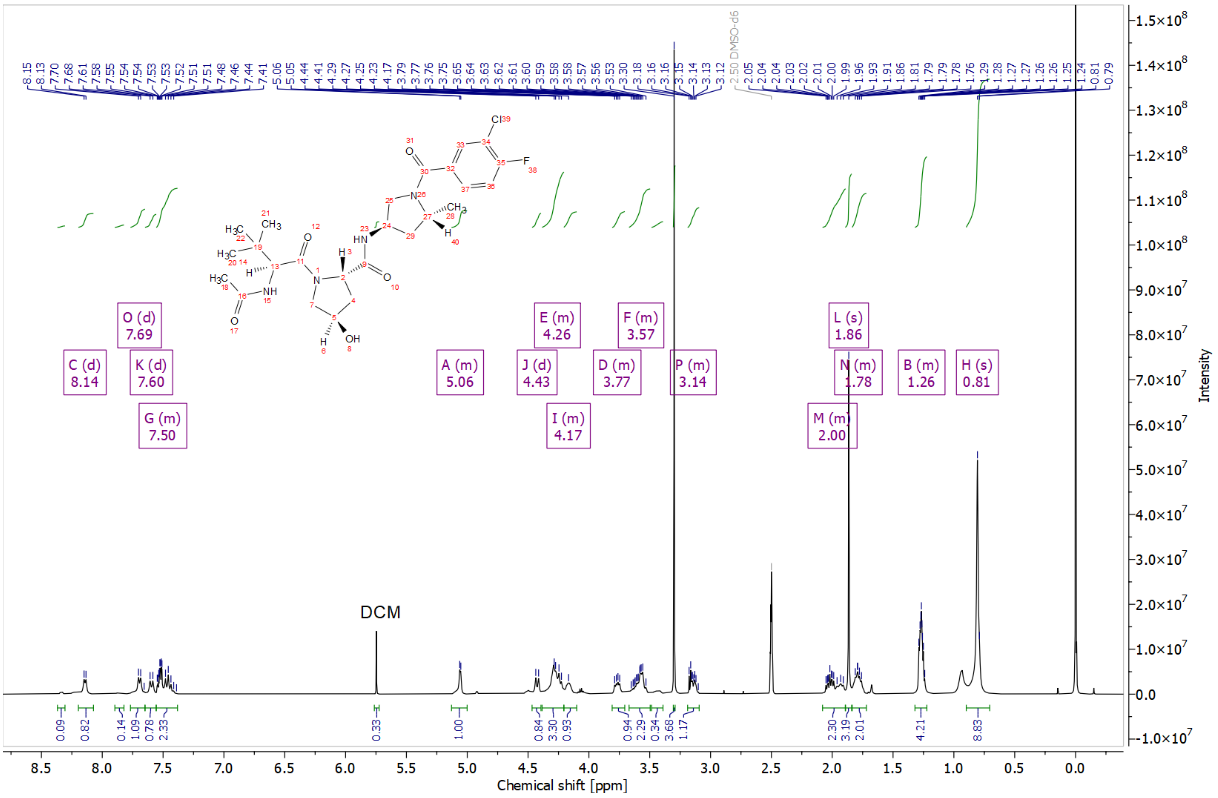


*Figure 13: $^1$H NMR spectrum in DMSO-$d_6$ of (2S,4R)-N-[(3R,5R)-1-(3-chloro-4-fluorobenzoyl)-5-methylpyrrolidin-3-yl]-1-[(2S)-2-acetamido-3,3-dimethylbutanoyl]-4-hydroxypyrrolidine-2-carboxamide (Compound 10)*

$^{1}$H NMR (400 MHz, DMSO) δ (mixture of 2 species (rotamers) major+minor) 8.13 – 7.95 (m, 1H, major+minor), 7.91 – 7.77 (m, 1H, major+minor), 7.41 – 7.29 (m, 1H, major+minor), 7.26 – 7.12 (m, 2H, major+minor), 5.14 – 5.00 (m, 1H, major+minor), 4.55 – 4.42 (m, 1H, major+minor), 4.37 – 4.19 (m, 2H, major+minor), 3.69 – 2.92 (m, 8H, major+minor), 2.44 – 2.21 (m, 1H, major+minor), 2.03 – 1.57 (m, 2H, major+minor), 1.88 – 1.85 (m, 3H, major+minor), 1.84 – 1.57 (m, 2H, major+minor), 0.95 – 0.87 (m, 9H, major+minor).

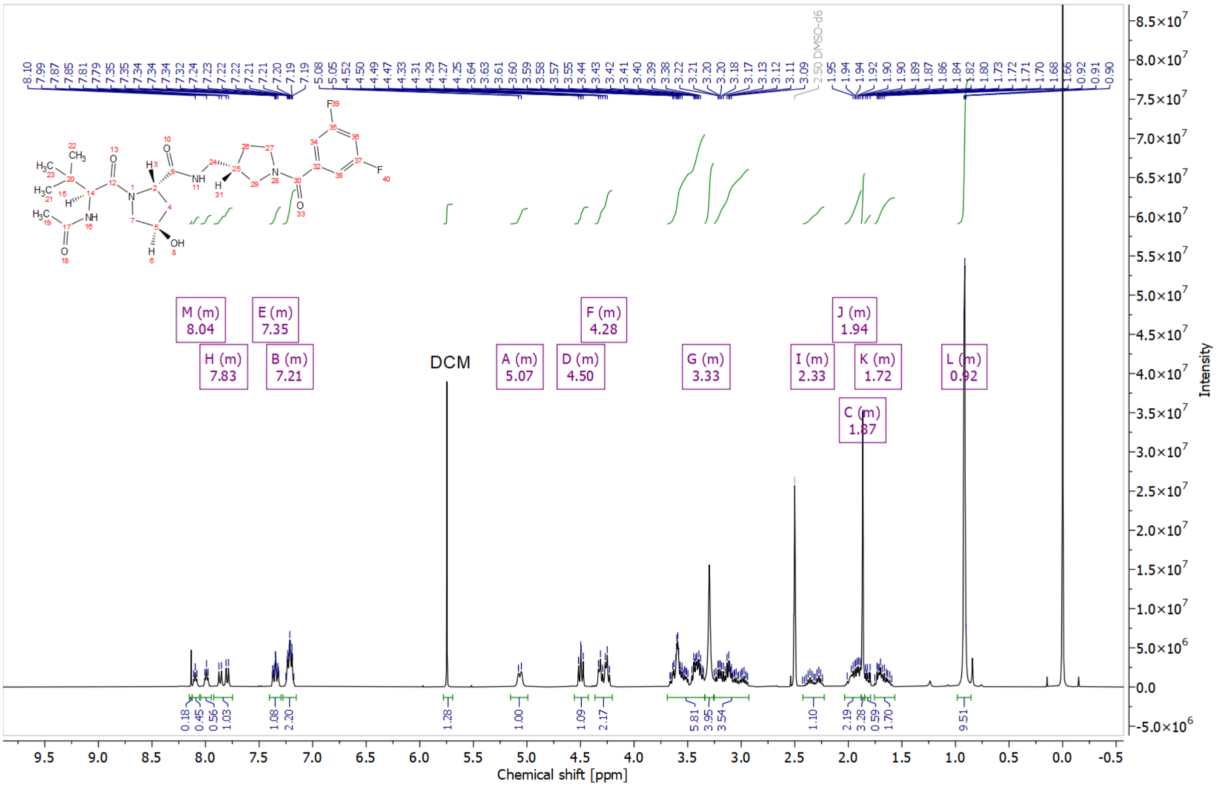


*Figure 14: $^{1}$H NMR spectrum in DMSO-$d_6$ of (2S,4R)-N-{[(3S)-1-(3,5-difluorobenzoyl)pyrrolidin-3-yl]methyl}-1-[(2S)-2-acetamido-3,3-dimethylbutanoyl]-4-hydroxypyrrolidine-2-carboxamide (Compound 11).*

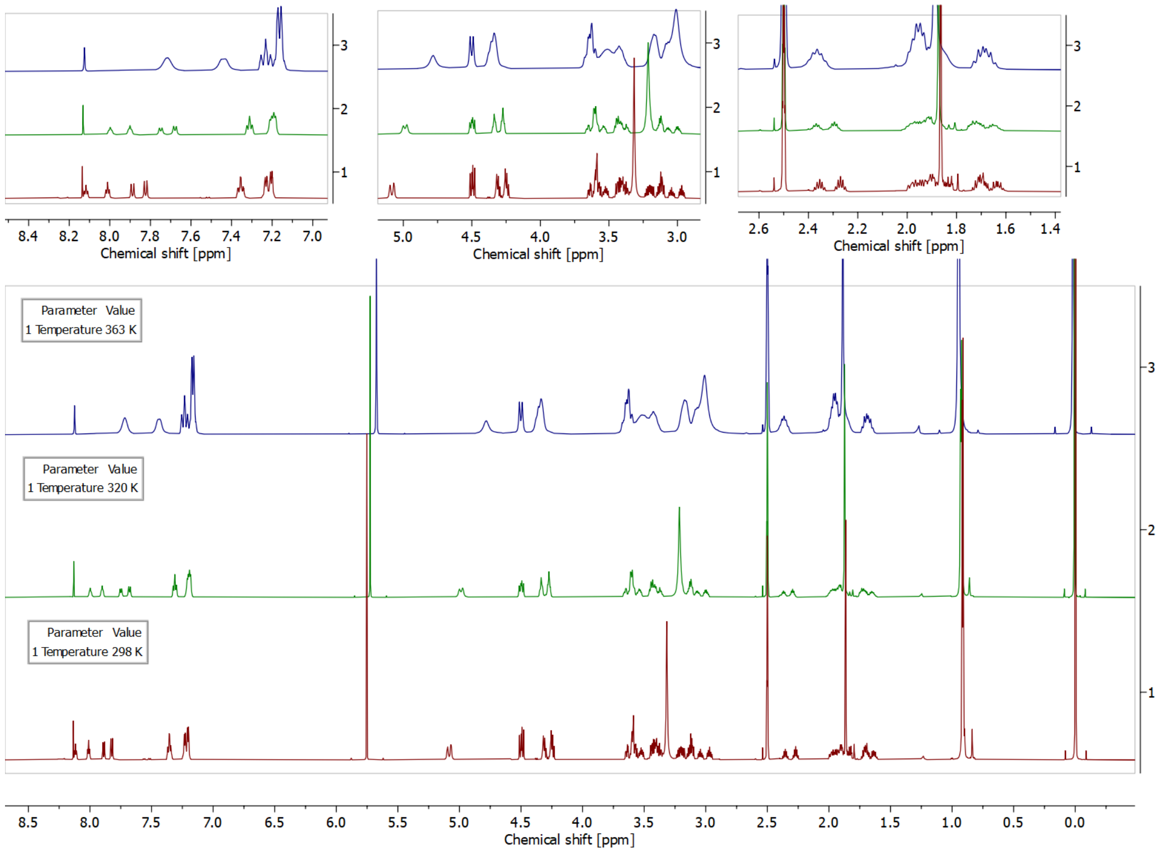


*Figure 15: [1]H NMR spectra of (2S,4R)-N-{[(3S)-1-(3,5-difluorobenzoyl)pyrrolidin-3-yl]methyl}-1-[(2S)-2-acetamido-3,3-dimethylbutanoyl]-4-hydroxypyrrolidine-2-carboxamide (Compound 11) in DMSO-$d_6$ at 298 K, 320 K, 363 K. Coalescence of signals at higher temperature suggest the product to exist in solution as a mixture of rotamers*

### NMR spectra of products modified at the left-hand side of Hyp-OH

$^1$H NMR (400 MHz, DMSO) δ 8.98 (s, 1H), 8.60 (t, *J* = 6.0 Hz, 1H), 7.49 – 7.41 (m, 1H), 7.40 (s, 4H), 5.69 (t, *J* = 5.8 Hz, 1H), 5.15 (d, *J* = 3.5 Hz, 1H), 4.56 (d, *J* = 9.6 Hz, 1H), 4.47 – 4.37 (m, 2H), 4.37 – 4.33 (m, 1H), 4.25 (dd, *J* = 15.8, 5.6 Hz, 1H), 3.90 (dd, *J* = 16.0, 5.7 Hz, 1H), 3.81 (dd, *J* = 16.0, 5.9 Hz, 1H), 3.67 (dd, *J* = 10.7, 3.9 Hz, 1H), 3.60 (d, *J* = 10.7 Hz, 1H), 2.45 (s, 3H), 2.11 – 2.02 (m, 1H), 1.90 (ddd, *J* = 13.0, 8.9, 4.4 Hz, 1H), 0.94 (s, 9H).

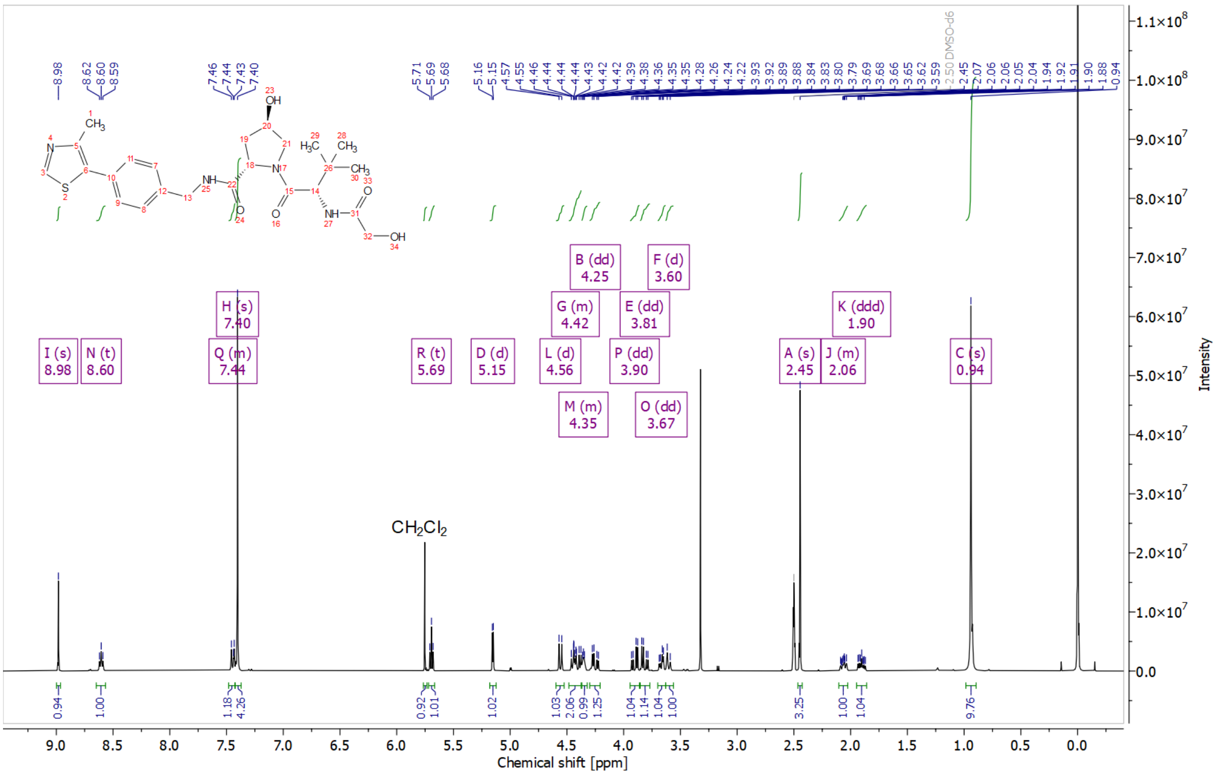


*Figure 16: $^1$H NMR spectrum in DMSO-$d_6$ of (2S,4R)-4-hydroxy-1-[(2S)-2-(2-hydroxyacetamido)-3,3-dimethylbutanoyl]-N-{[4-(4-methyl-1,3-thiazol-5-yl)phenyl]methyl}pyrrolidine-2-carboxamide (Compound 3)*

[1]H NMR (400 MHz, DMSO) δ 8.98 (s, 1H), 8.60 (t, $J$ = 6.0 Hz, 1H), 7.40 (s, 4H), 7.37 (d, $J$ = 9.5 Hz, 1H), 5.15 (d, $J$ = 3.5 Hz, 1H), 4.57 (d, $J$ = 9.6 Hz, 1H), 4.49 – 4.37 (m, 2H), 4.36 – 4.31 (m, 1H), 4.25 (dd, $J$ = 15.8, 5.6 Hz, 1H), 3.96 – 3.81 (m, 2H), 3.66 (dd, $J$ = 10.7, 3.9 Hz, 1H), 3.60 (d, $J$ = 10.7 Hz, 1H), 3.33 (s, 3H), 2.44 (s, 3H), 2.11 – 2.01 (m, 1H), 1.90 (ddd, $J$ = 13.0, 8.9, 4.5 Hz, 1H), 0.94 (s, 9H).

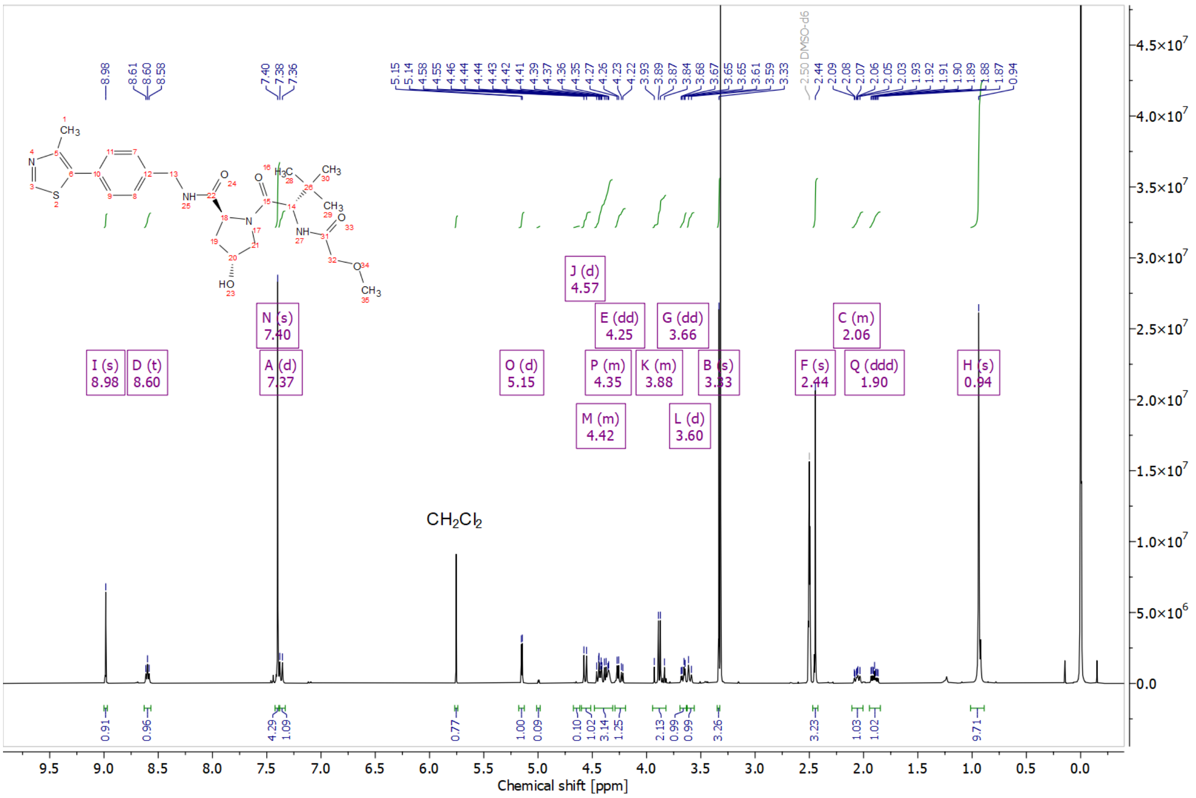


*Figure 17: [1]H NMR spectrum in DMSO-$d_6$ of (2S,4R)-4-hydroxy-1-[(2S)-2-(2-methoxyacetamido)-3,3-dimethylbutanoyl]-N-{[4-(4-methyl-1,3-thiazol-5-yl)phenyl]methyl}pyrrolidine-2-carboxamide (Compound 4)*

[1]H NMR (400 MHz, DMSO) δ 8.98 (s, 1H), 8.62 (t, $J$ = 6.0 Hz, 1H), 7.46 – 7.31 (m, 5H), 5.15 (d, $J$ = 3.5 Hz, 1H), 4.55 (d, $J$ = 9.6 Hz, 1H), 4.49 – 4.37 (m, 2H), 4.37 – 4.31 (m, 1H), 4.25 (dd, $J$ = 15.8, 5.6 Hz, 1H), 3.91 (d, $J$ = 1.2 Hz, 2H), 3.66 (dd, $J$ = 10.7, 3.8 Hz, 1H), 3.60 (d, $J$ = 10.7 Hz, 1H), 3.47 (td, $J$ = 6.4, 0.9 Hz, 2H), 2.44 (s, 3H), 2.07 (ddd, $J$ = 12.8, 5.4, 3.8 Hz, 1H), 1.90 (ddd, $J$ = 13.0, 9.0, 4.5 Hz, 1H), 1.58 – 1.46 (m, 2H), 1.42 – 1.30 (m, 2H), 0.94 (s, 9H), 0.89 (t, $J$ = 7.4 Hz, 3H).

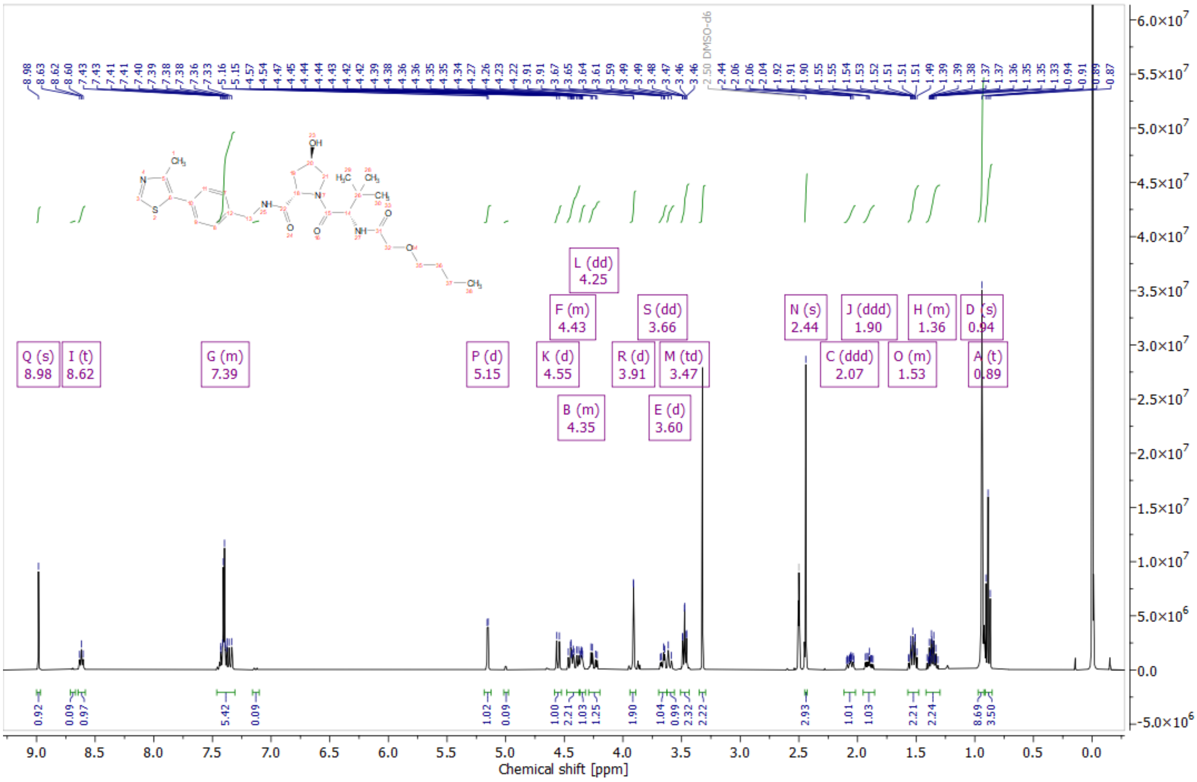


*Figure 18: [1]H NMR spectrum in DMSO-$d_6$ of (2S,4R)-1-[(2S)-2-(2-butoxyacetamido)-3,3-dimethylbutanoyl]-4-hydroxy-N-{[4-(4-methyl-1,3-thiazol-5-yl)phenyl]methyl}pyrrolidine-2-carboxamide (Compound 5)*

$^1$H NMR (400 MHz, DMSO) δ 8.99 (s, 1H), 8.59 (t, $J$ = 6.1 Hz, 1H), 8.02 (d, $J$ = 9.3 Hz, 1H), 7.53 – 7.46 (m, 1H), 7.46 – 7.37 (m, 6H), 7.33 – 7.27 (m, 1H), 5.14 (d, $J$ = 3.5 Hz, 1H), 4.83 – 4.67 (m, 2H), 4.58 (d, $J$ = 9.4 Hz, 1H), 4.50 – 4.38 (m, 2H), 4.35 (s, 1H), 4.24 (dd, $J$ = 15.8, 5.5 Hz, 1H), 3.67 (dd, $J$ = 10.6, 4.0 Hz, 1H), 3.60 (d, $J$ = 10.7 Hz, 1H), 2.45 (s, 3H), 2.10 – 2.01 (m, 1H), 1.95 – 1.85 (m, 1H), 0.94 (s, 9H).

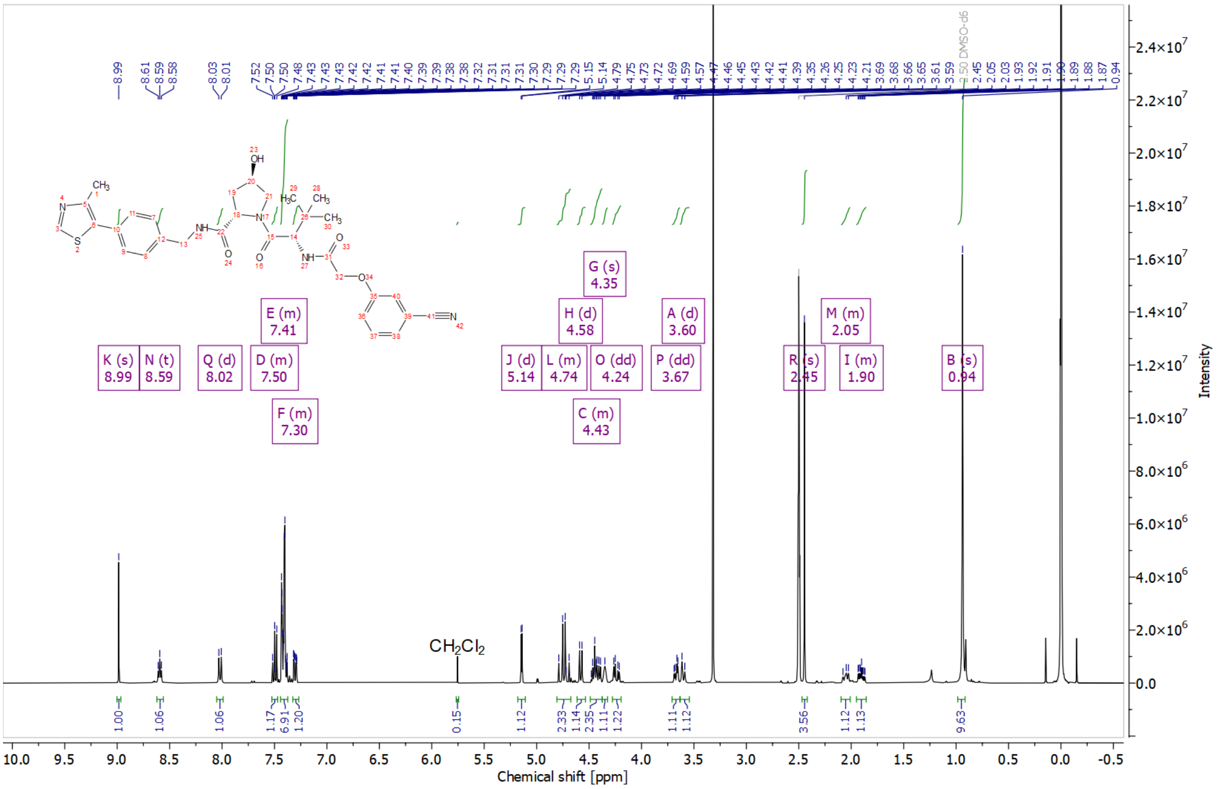


*Figure 19: $^1$H NMR spectrum in DMSO-$d_6$ of (2S,4R)-1-[(2S)-2-[2-(3-cyanophenoxy)acetamido]-3,3-dimethylbutanoyl]-4-hydroxy-N-{[4-(4-methyl-1,3-thiazol-5-yl)phenyl]methyl}pyrrolidine-2-carboxamide (Compound 6)*

### HPLCMS chromatograms of intermediates

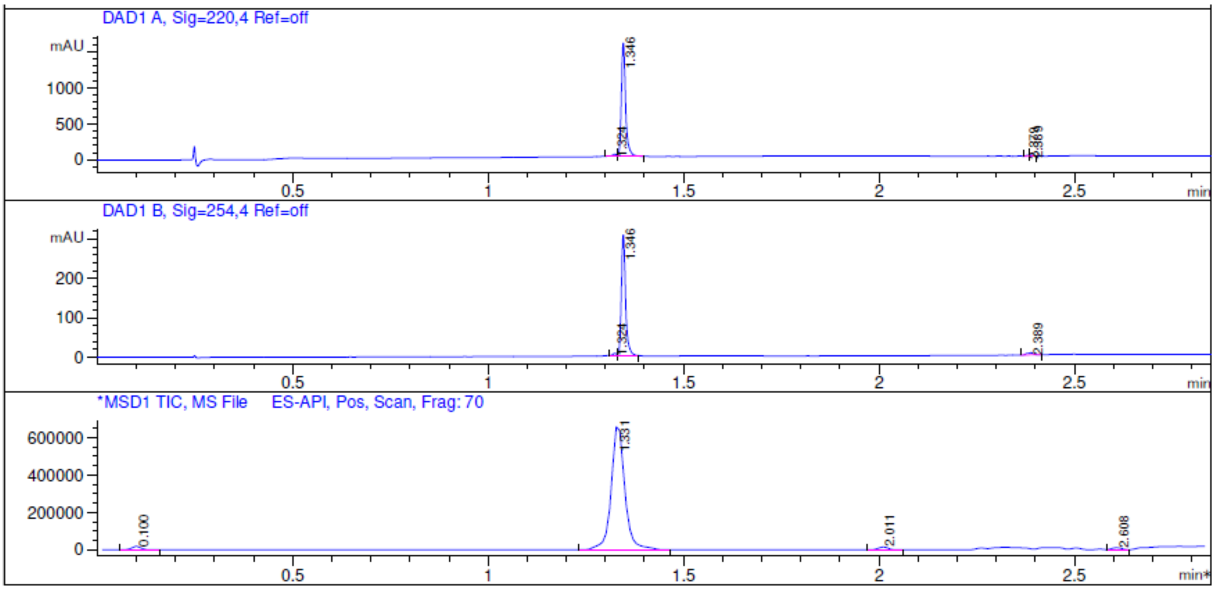


*Figure 20: LCMS chromatogram of 4-(1H-pyrazol-5-yl)naphthalene-1-carbonitrile (Int-2a)*

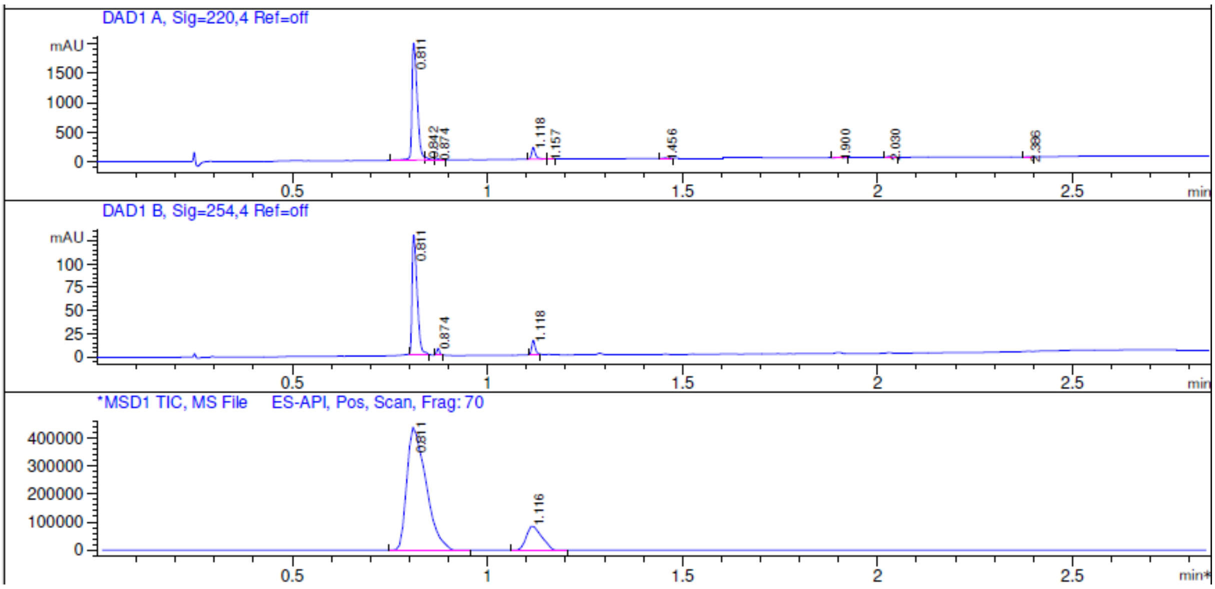


*Figure 21: LCMS chromatogram of 1-[4-(1H-pyrazol-5-yl)naphthalen-1-yl]methanamine (Int-2b)*

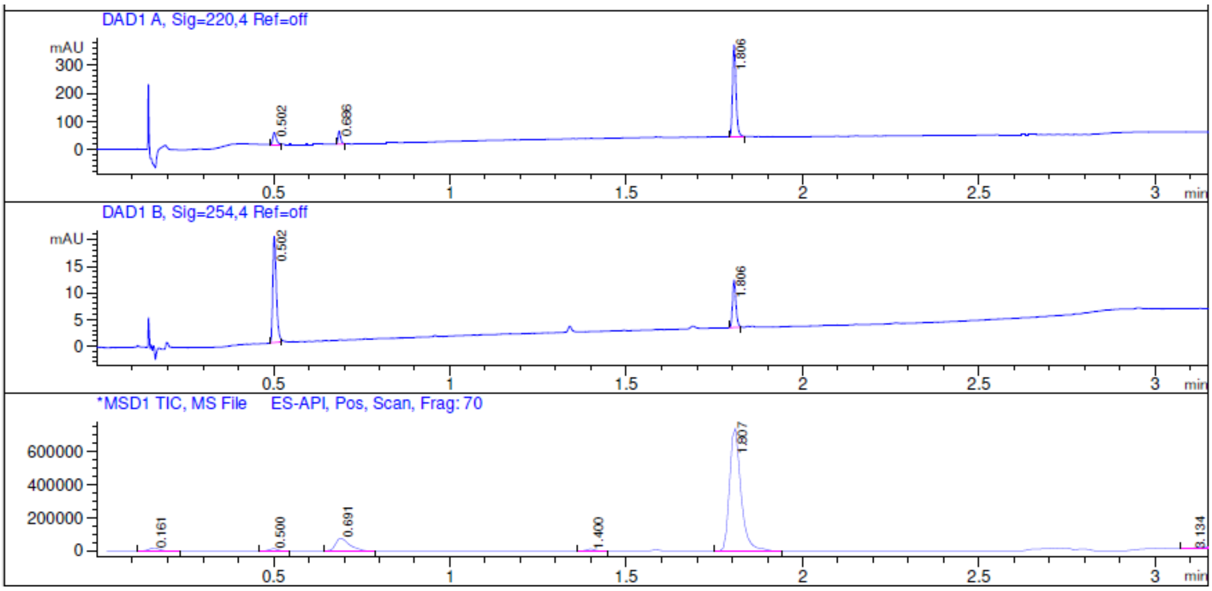


*Figure 22: LCMS chromatogram of tert-butyl N-[2-(3-methyl-3-phenylpiperidin-1-yl)-2-oxoethyl]carbamate (Int-7a)*

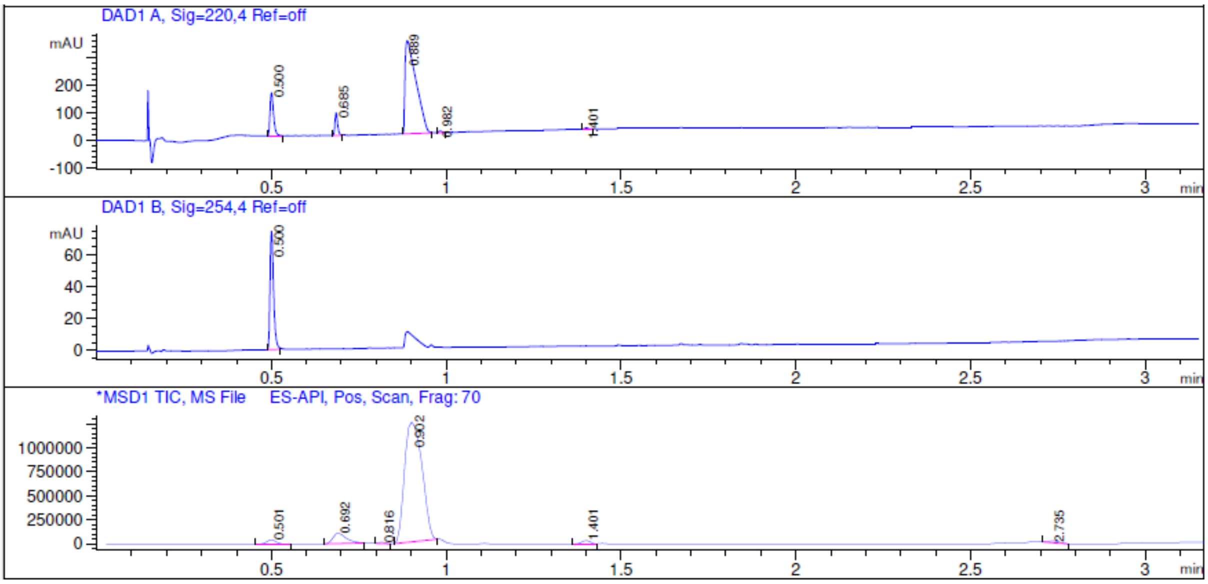


*Figure 23: LCMS chromatogram of crude HCl salt 2-amino-1-(3-methyl-3-phenylpiperidin-1-yl)ethan-1-one hydrochloride (Int-7b)*

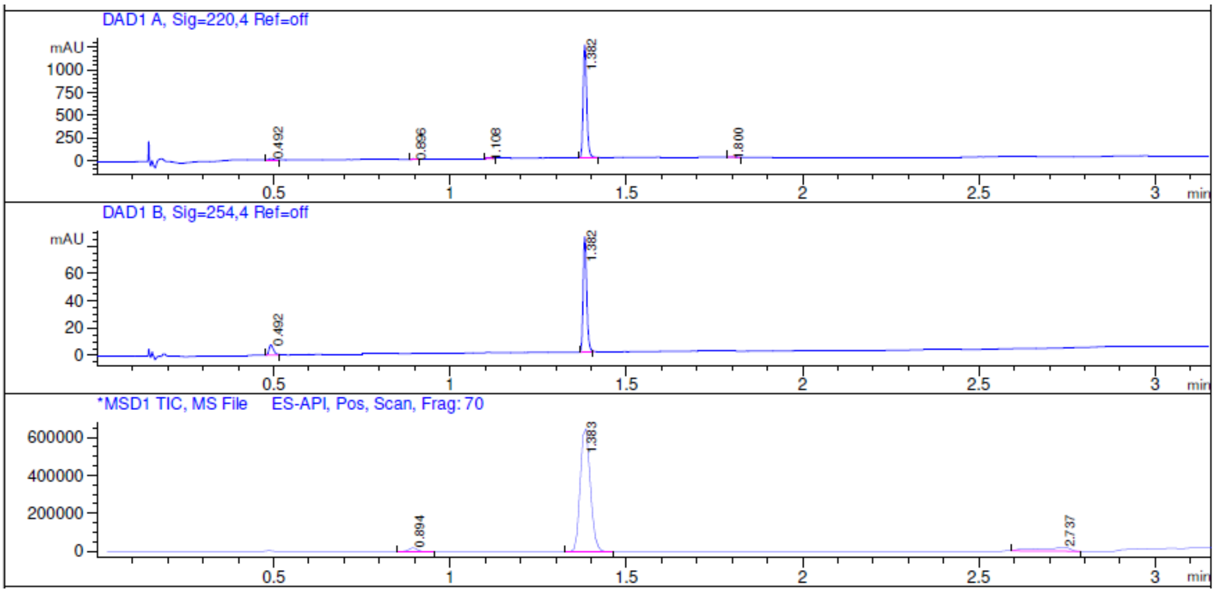


*Figure 24: LCMS chromatogram of tert-butyl N-[(3R)-1-(4-hydroxynaphthalene-1-carbonyl)pyrrolidin-3-yl]carbamate (Int-8a)*

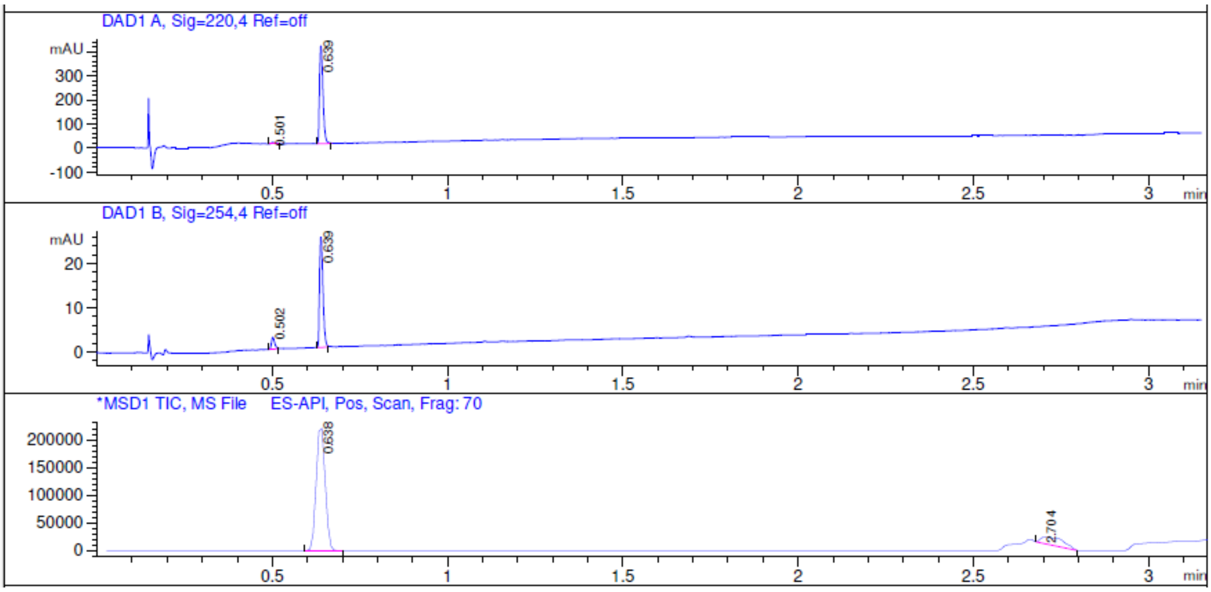


*Figure 25: LCMS chromatogram of crude HCl salt 4-[(3R)-3-aminopyrrolidine-1-carbonyl]naphthalen-1-ol hydrochloride (Int-8b)*

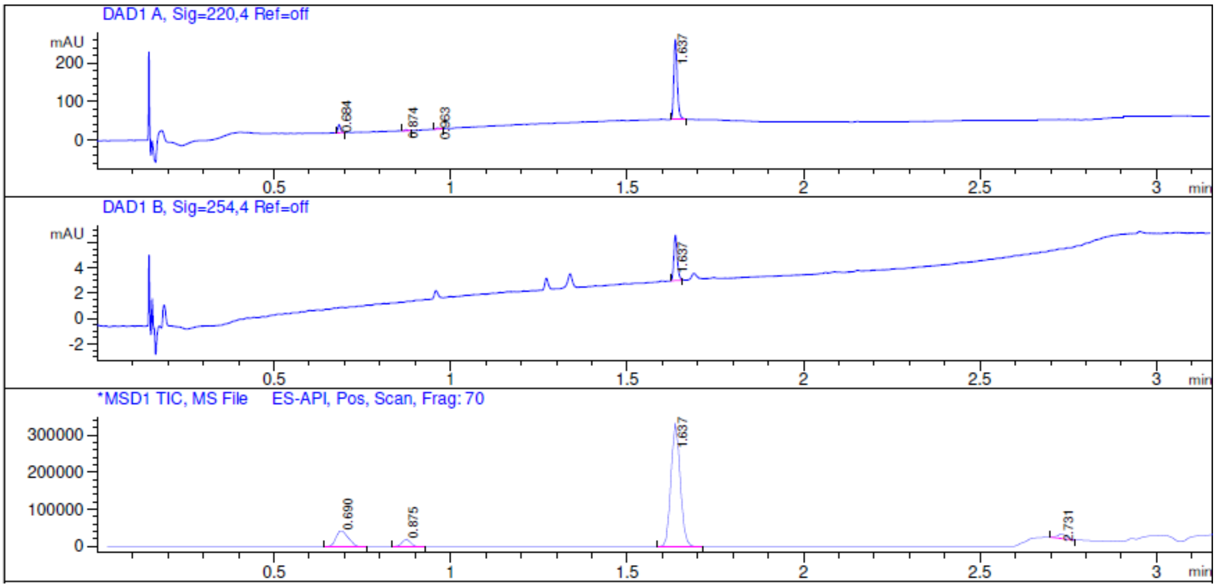


*Figure 26: LCMS chromatogram of tert-Butyl N-{2-[3-(3-chlorophenyl)morpholin-4-yl]-2-oxoethyl}carbamate (Int-9a)*

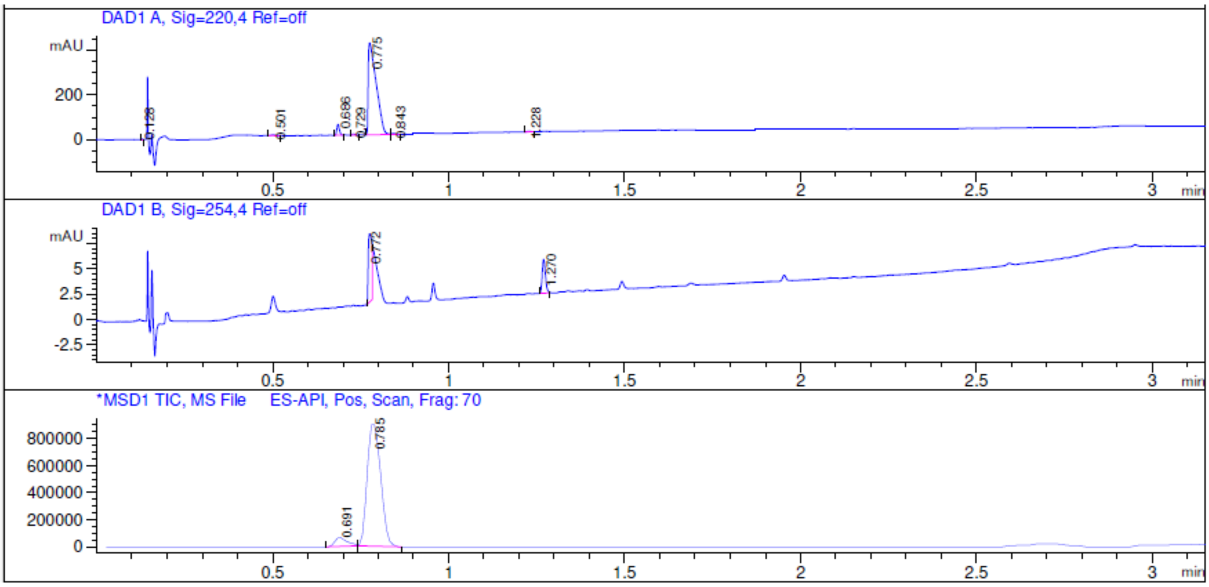


*Figure 27: LCMS chromatogram of 3-(3-chlorophenyl)morpholine hydrochloride (Int-9b)*

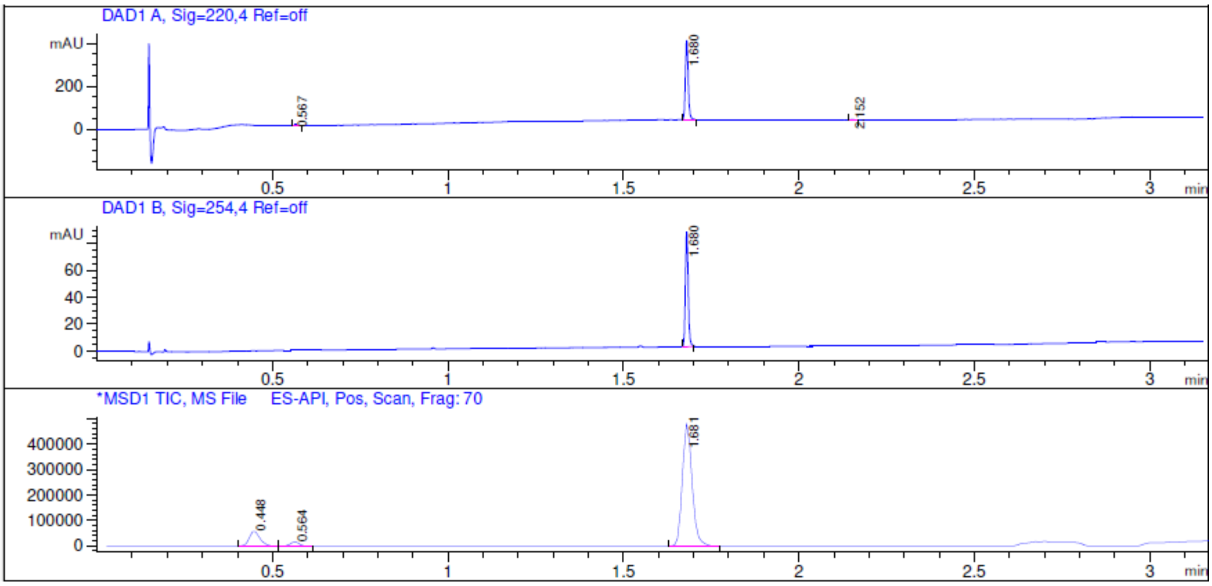


*Figure 28: LCMS chromatogram of tert-butyl N-[(3R,5R)-1-(3-chloro-4-fluorobenzoyl)-5-methylpyrrolidin-3-yl]carbamate (Int-10a)*

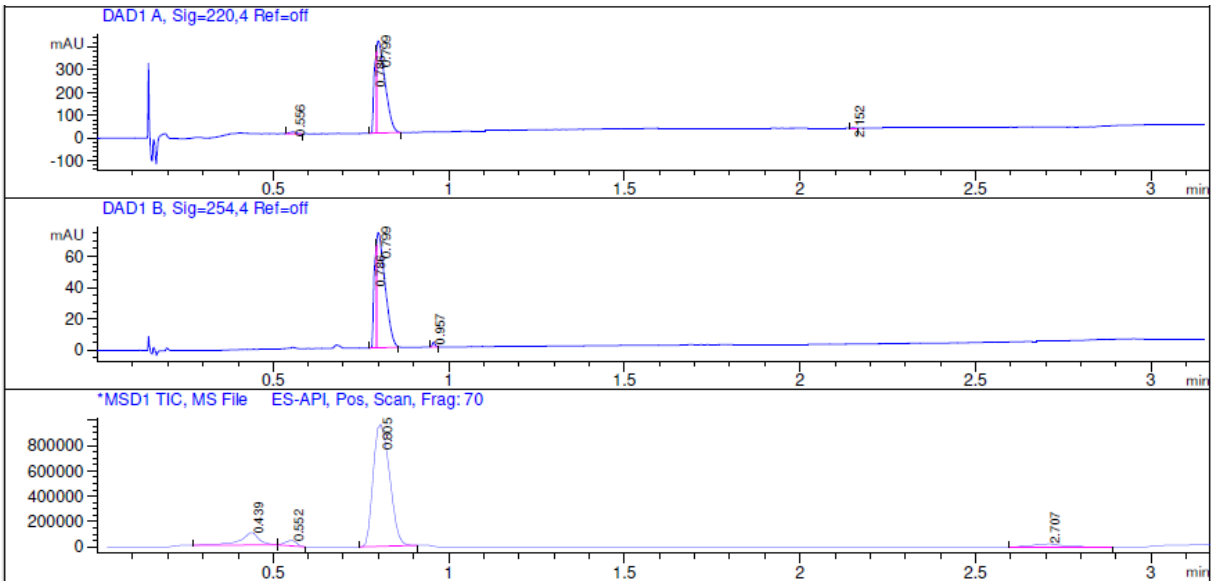


*Figure 29: LCMS chromatogram of (3R,5R)-1-(3-chloro-4-fluorobenzoyl)-5-methylpyrrolidin-3-amine hydrochloride (Int-10b)*

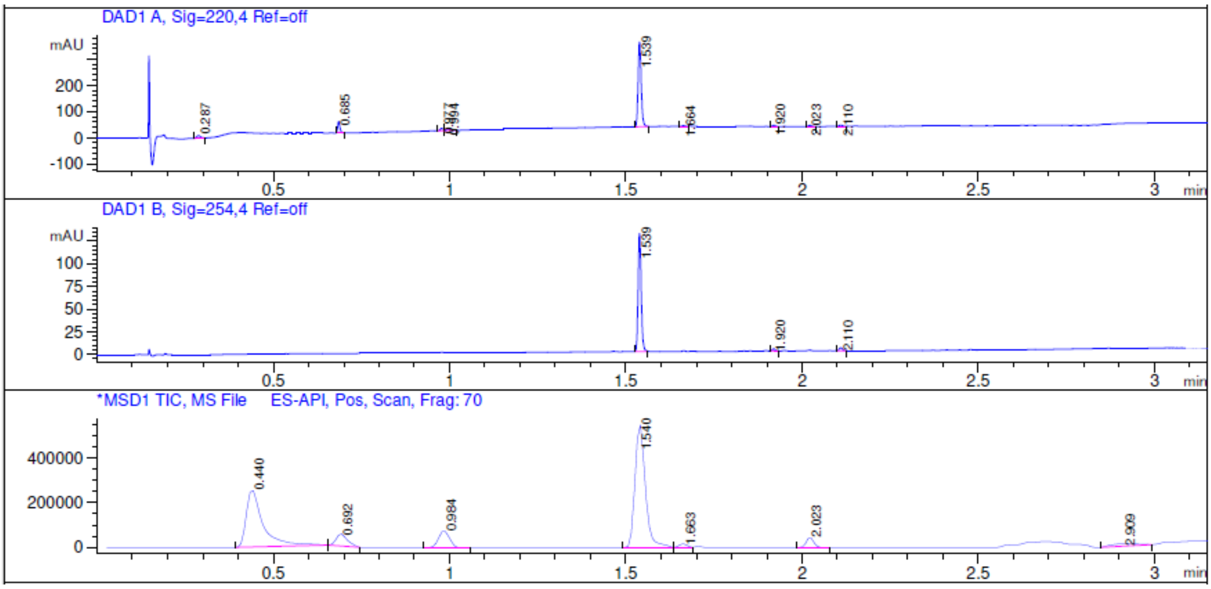


*Figure 30: LCMS chromatogram of tert-butyl N-{[(3S)-1-(3,5-difluorobenzoyl)pyrrolidin-3-yl]methyl}carbamate (Int-11a)*

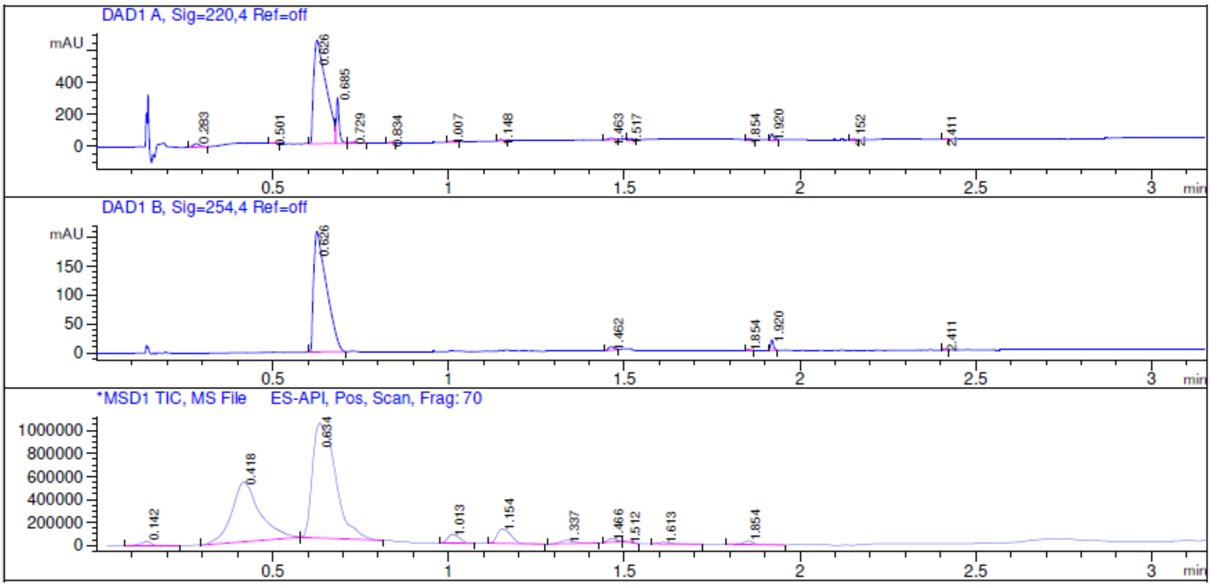


*Figure 31: LCMS chromatogram of 1-[(3S)-1-(3,5-difluorobenzoyl)pyrrolidin-3-yl]methanamine hydrochloride (Int-11b)*

### HPLCMS chromatograms of products modified at the right-hand side of Hyp-OH

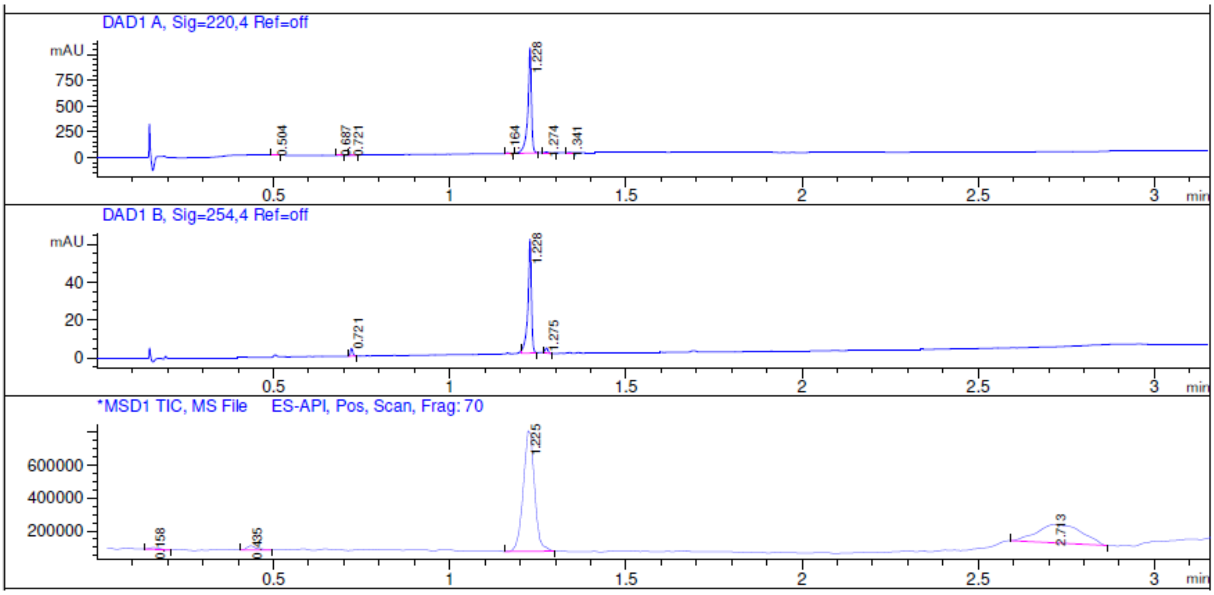


*Figure 32: LCMS chromatograms of (2S,4R)-1-[(2S)-2-acetamido-3,3-dimethylbutanoyl]-4-hydroxy-N-{[4-(1H-pyrazol-5-yl)naphthalen-1-yl]methyl}pyrrolidine-2-carboxamide (Compound 2)*

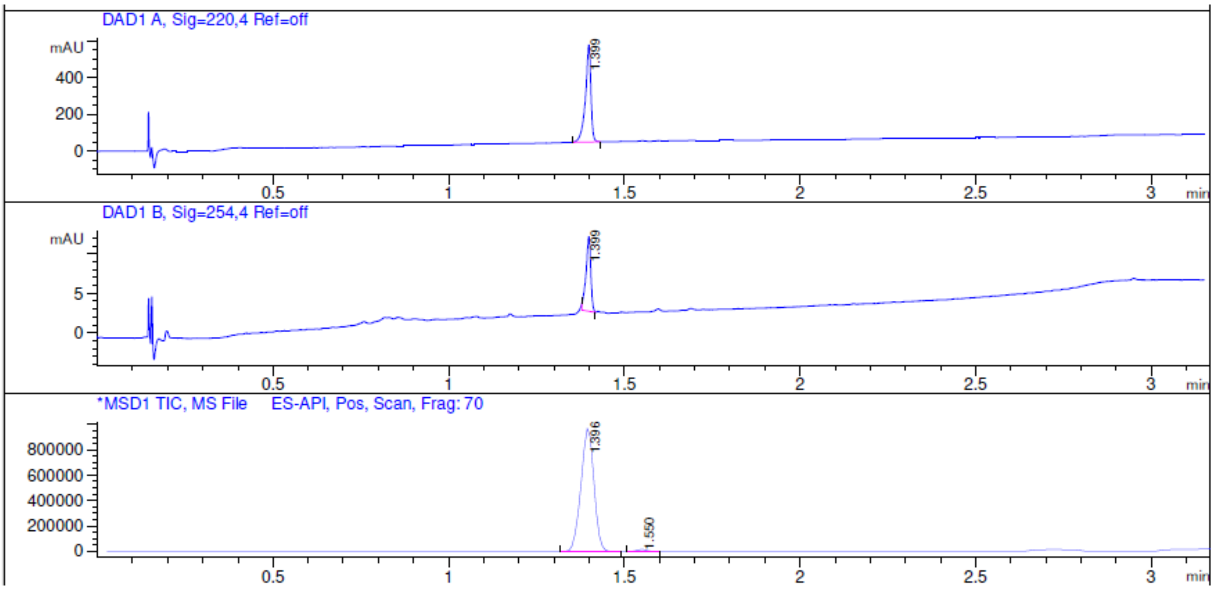


*Figure 33: LCMS chromatogram of (2S,4R)-1-[(2S)-2-acetamido-3,3-dimethylbutanoyl]-4-hydroxy-N-[2-(3-methyl-3-phenylpiperidin-1-yl)-2-oxoethyl]pyrrolidine-2-carboxamide (Compound 7)*

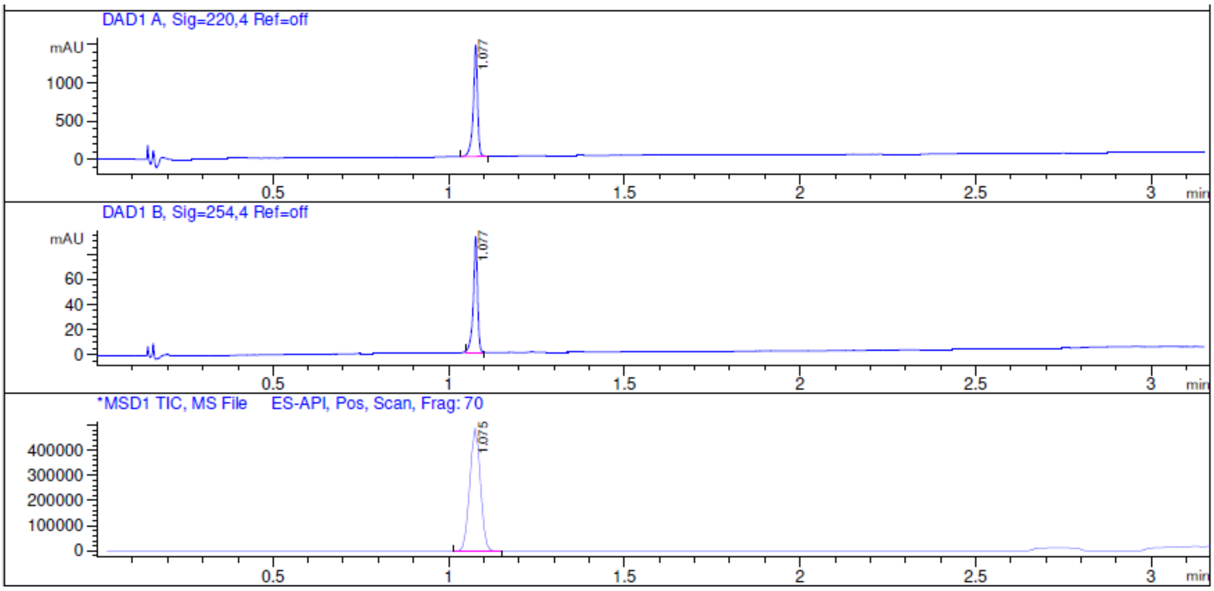


*Figure 34: LCMS chromatogram of (2S,4R)-1-[(2S)-2-acetamido-3,3-dimethylbutanoyl]-4-hydroxy-N-[(3R)-1-(4-hydroxynaphthalene-1-carbonyl)pyrrolidin-3-yl]pyrrolidine-2-carboxamide (Compound 8)*

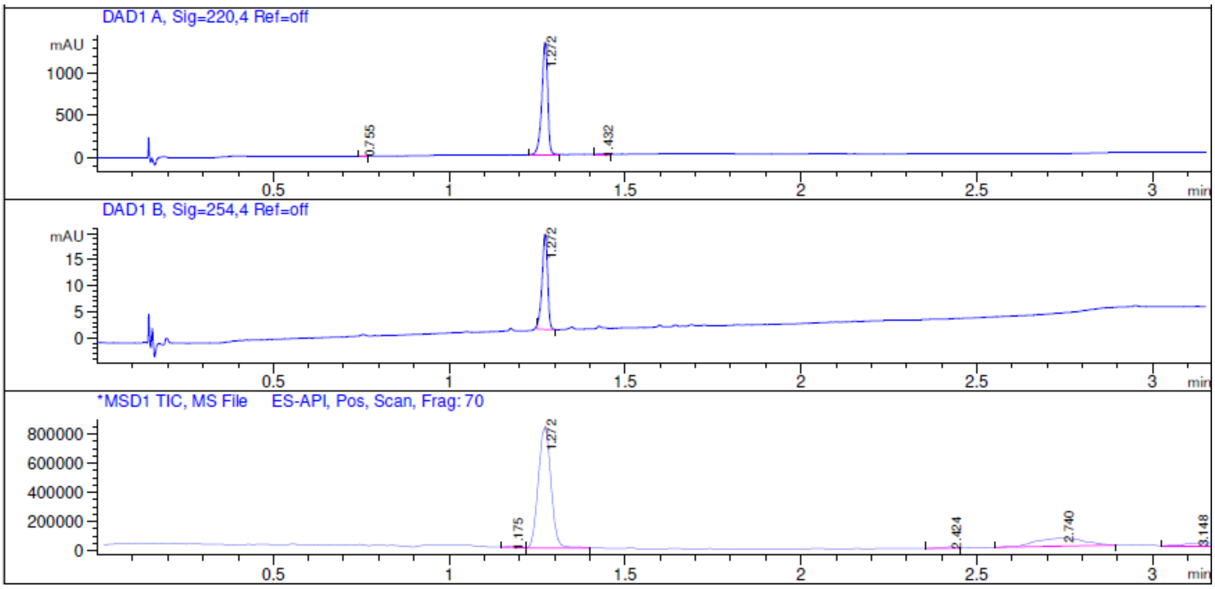


*Figure 35: LCMS chromatogram of (2S,4R)-N-{2-[3-(3-chlorophenyl)morpholin-4-yl]-2-oxoethyl}-1-[(2S)-2-acetamido-3,3-dimethylbutanoyl]-4-hydroxypyrrolidine-2-carboxamide (Compound 9)*

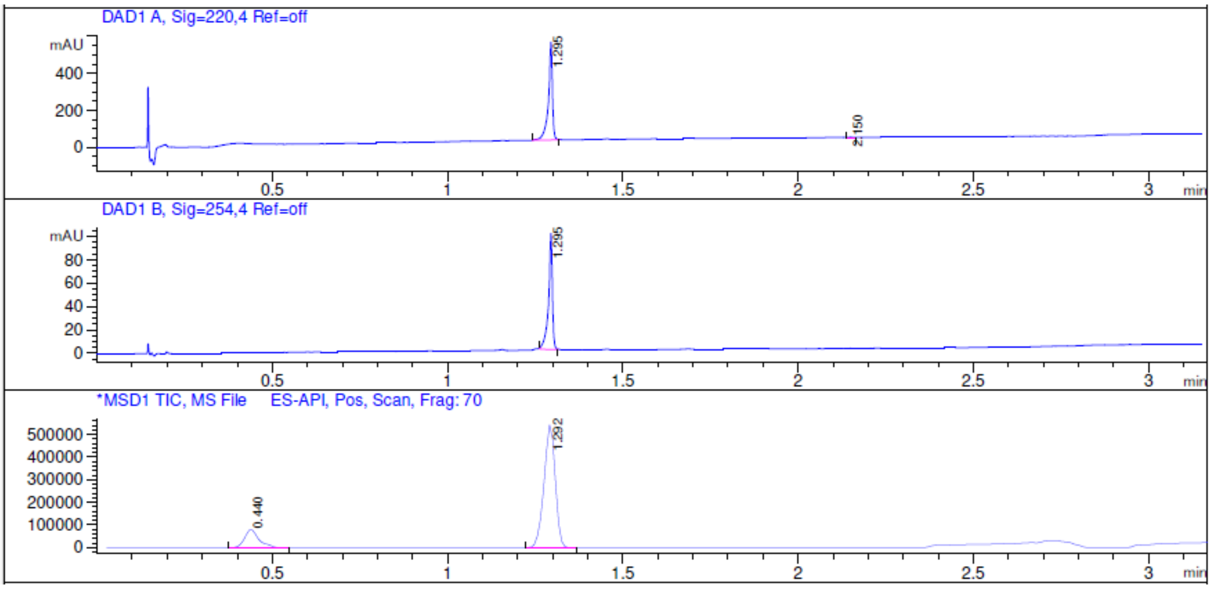


*Figure 36: LCMS chromatogram of (2S,4R)-N-[(3R,5R)-1-(3-chloro-4-fluorobenzoyl)-5-methylpyrrolidin-3-yl]-1-[(2S)-2-acetamido-3,3-dimethylbutanoyl]-4-hydroxypyrrolidine-2-carboxamide (Compound 10)*

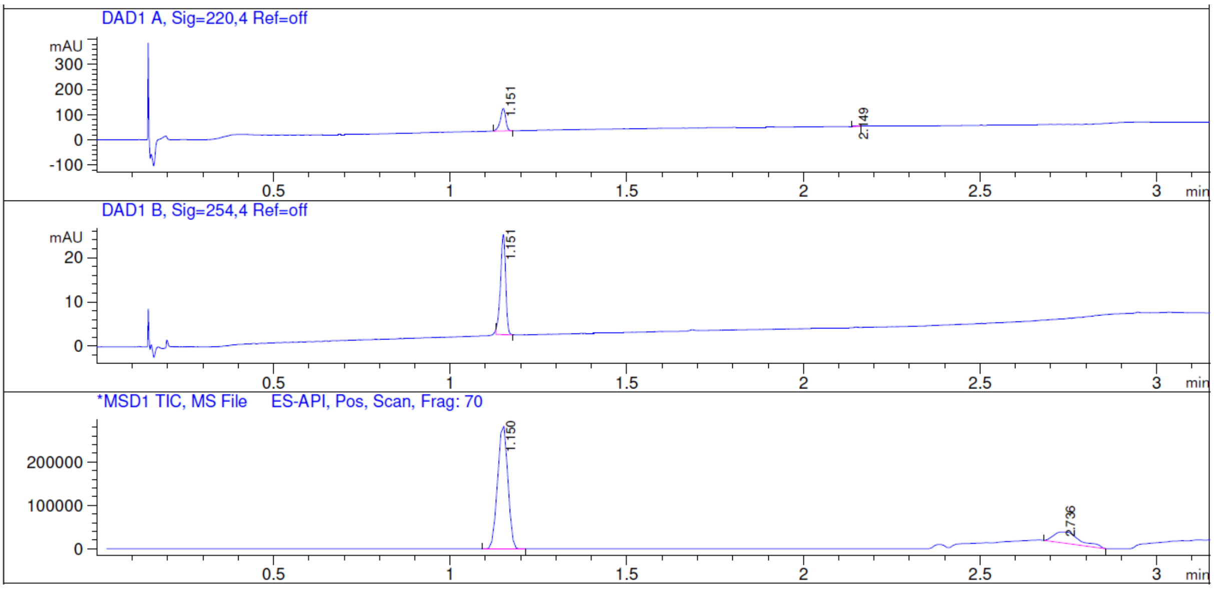


*Figure 37: LCMS chromatogram of (2S,4R)-N-{[(3S)-1-(3,5-difluorobenzoyl)pyrrolidin-3-yl]methyl}-1-[(2S)-2-acetamido-3,3-dimethylbutanoyl]-4-hydroxypyrrolidine-2-carboxamide (Compound 11)*

#### HPLC-MS chromatograms of products modified at the left-hand side of Hyp-OH

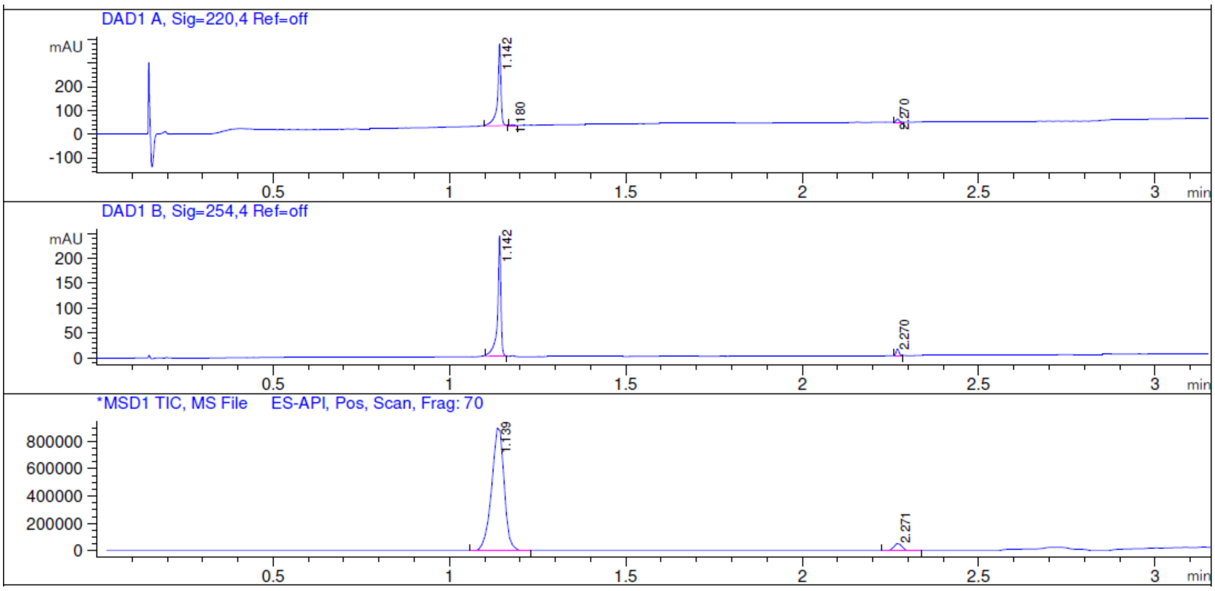


*Figure 38: LCMS chromatogram of (2S,4R)-4-hydroxy-1-[(2S)-2-(2-hydroxyacetamido)-3,3-dimethylbutanoyl]-N-{[4-(4-methyl-1,3-thiazol-5-yl)phenyl]methyl}pyrrolidine-2-carboxamide (Compound 3)*

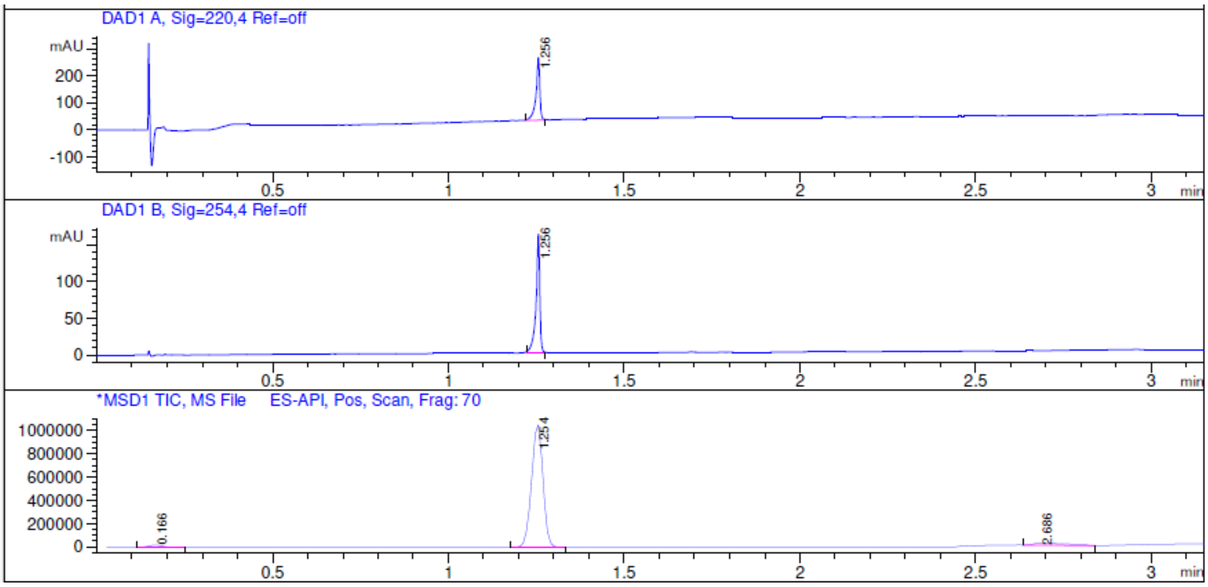


*Figure 39: LCMS chromatogram of (2S,4R)-4-hydroxy-1-[(2S)-2-(2-methoxyacetamido)-3,3-dimethylbutanoyl]-N-{[4-(4-methyl-1,3-thiazol-5-yl)phenyl]methyl}pyrrolidine-2-carboxamide (Compound 4)*

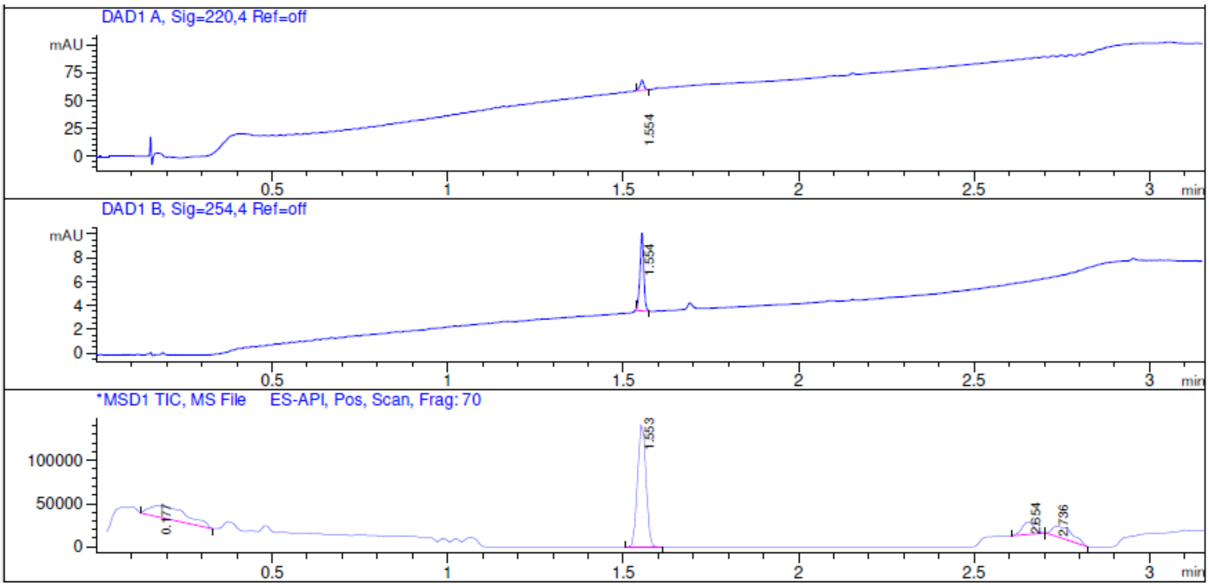


*Figure 40: LCMS chromatogram of (2S,4R)-1-[(2S)-2-(2-butoxyacetamido)-3,3-dimethylbutanoyl]-4-hydroxy-N-{[4-(4-methyl-1,3-thiazol-5-yl)phenyl]methyl}pyrrolidine-2-carboxamide (Compound 5)*

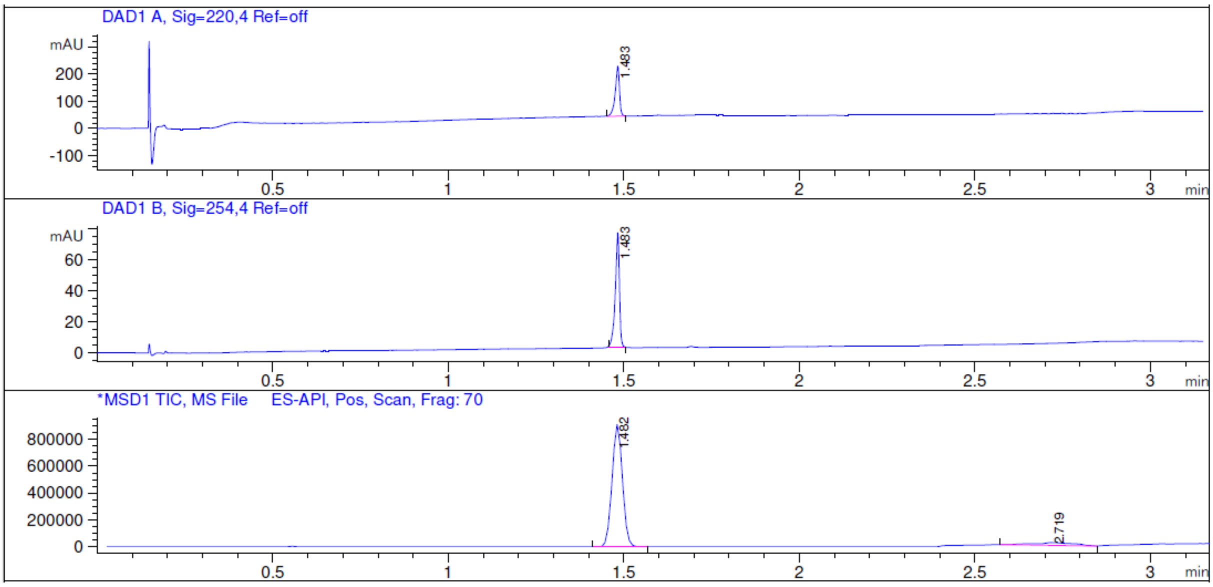


*Figure 41: LCMS chromatogram of (2S,4R)-1-[(2S)-2-[2-(3-cyanophenoxy)acetamido]-3,3-dimethylbutanoyl]-4-hydroxy-N-{[4-(4-methyl-1,3-thiazol-5-yl)phenyl]methyl}pyrrolidine-2-carboxamide (Compound 6)*
